# The dispersal of domestic cats from Northern Africa and their introduction to Europe over the last two millennia

**DOI:** 10.1101/2025.03.28.645893

**Authors:** M. De Martino, B. De Cupere, V. Rovelli, P. Serventi, M. Baldoni, T. Di Corcia, S. Geiger, F. Alhaique, P. C. Alves, H. Buitenhuis, E. Ceccaroni, E. Cerilli, J. De Grossi Mazzorin, C. Detry, M. Dowd, I. Fiore, L. Gourichon, I. Grau-Sologestoa, H. C. Küchelmann, G. K. Kunst, M. McCarthy, R. Miccichè, C. Minniti, M. Moreno, V. Onar, T. Oueslati, M. Parrag, B. Pino Uria, G. Romagnoli, M. Rugge, L. Salari, K. Saliari, A. B. Santos, U. Schmölcke, A. Sforzi, G. Soranna, N. Spassov, A. Tagliacozzo, V. Tinè, S. Trixl, S. Vuković, U. Wierer, B. Wilkens, S. Doherty, N. Sykes, L. Frantz, F. Mattucci, R. Caniglia, G. Larson, J. Peters, W. Van Neer, C. Ottoni

**Affiliations:** Centre of Molecular Anthropology for Ancient DNA studies, Department of Biology, University of Rome Tor Vergata; Rome, 00133, Italy; PhD Program in Evolutionary Biology and Ecology, Department of Biology, University of Rome Tor Vergata; Rome, 00133, Italy; Royal Belgian Institute of Natural Sciences; Brussels, B-1000, Belgium; Institute of Palaeoanatomy, Domestication Research and the History of Veterinary Medicine, Ludwig-Maximilians University Munich; Munich, 80539, Germany; Bioarchaeology Service, Museo delle Civiltà; Rome, 00144, Italy; CIBIO, Research Centre in Biodiversity and Genetic Resources, InBIO/BIOPOLIS Program in Genomics, Biodiversity and Land Planning, Universidade do Porto, Campus de Vairão; Vairão, 4485-661, Portugal; Departamento de Biologia, Faculdade de Ciências, Universidade Do Porto; 4099-002, Porto, Portugal; Estação Biológica de Mértola, EBM, Praça Luís de Camões; 7750-329, Mértola, Portugal; Groningen Institute of Archaeology, University of Groningen; NL-9712 ER Groningen, The Netherlands; Soprintendenza Archeologia, Belle Arti e Paesaggio per le province di L’Aquila e Teramo; L’Aquila, 67100, Italy; Independent researcher; Italy; Department of Cultural Heritage, University of Salento; Lecce, 73100, Italy; UNIARQ - Center of Archaeology of the University of Lisbon, Department of History, School of Arts and Humanities, University of Lisbon; Alameda da Universidade, 1600-214, Lisbon, Portugal; Faculty of Science, Atlantic Technological University; Sligo, F91YW50, Ireland; CEPAM, UMR 7264, Université Côte d’Azur, CNRS; Nice, 06000, France; IPNA/IPAS, Department of Environmental Sciences, University of Basel; Basel, 4055, Switzerland; Landesarchäologie Bremen; Bremen, D-28195, Germany; Vienna Institute for Archaeological Science, Research Network Human Evolution and Archaeological Sciences, University of Vienna; Vienna, 1090, Austria; Independent Consultant Zooarchaeologist.; Department of Biological, Chemical and Pharmaceutical Sciences and Technologies, University of Palermo; Palermo, 90123, Italy; Department of Science of Antiquities, Sapienza University of Rome; Rome, 00185, Italy; Spanish National Research Council, Institute of History; Madrid, 28037, Spain; Department of Anatomy, Milas Faculty of Veterinary Medicine, Muğla Sıtkı Koçman University, Milas, Muğla, Türkiye; CNRS, HALMA UMR 8164, University of Lille; Villeneuve d’Ascq 59653, France; Independent researcher; Vienna, Austria; Department of Modern Languages and Literatures, History, Philosophy and Law Studies, Tuscia University; Viterbo, 01100, Italy; Department of Cultural Heritage, Underwater Archaeology, University of Salento; Lecce, 73100, Italy; Department of History, Culture and Society, University of Rome Tor Vergata; Rome, 00133, Italy; Natural History Museum of Vienna; Vienna, 1010, Austria; Zentrum für Baltische und Skandinavische Archäologie (ZBSA), Leibniz Zentrum für Archäologie (LEIZA); Schloss Gottorf, D-24837 Schleswig, Germany; Natural History Museum of Maremma; Grosseto, 58100, Italy; National Museum of Natural History at the Bulgarian Academy of Sciences; Sofia, BG-1000, Bulgaria; Soprintendenza Archeologia Belle Arti e Paesaggio per l’area metropolitana di Venezia e le Province di Belluno, Padova e Treviso; Padua, 35139, Italy; State Office for Cultural Heritage Management Baden-Wuerttemberg; Constance, 78467, Germany; Laboratory for Bioarchaeology, Archaeology Department, University of Belgrade – Faculty of Philosophy; Belgrade, 11000, Serbia; Soprintendenza Archeologia, Belle Arti e Paesaggio per la città metropolitana di Firenze e le province di Pistoia e Prato; Florence, 50125, Italy; Department of Archaeology and History, University of Exeter; Exeter, UK; School of Biological and Chemical Sciences, Queen Mary University of London, London, UK; Unit for Conservation Genetics (BIO-CGE), Italian Institute for Environmental Protection and Research (ISPRA); Ozzano dell’Emilia, 40064, Italy; The Palaeogenomics & Bio-Archaeology Research Network, Research Laboratory for Archaeology and History of Art, The University of Oxford, Oxford, UK; Bavarian Natural History Collections, State Collection of Palaeoanatomy; Munich, 80333, Germany; ArchaeoBioCenter, Ludwig-Maximilians University Munich; Munich, 80539, Germany

**Author notes:** Corresponding author. (C.O.). Deceased during the course of this study (E. Cerilli, J. De Grossi Mazzorin).

## Abstract

The domestic cat (*Felis catus*) descends from the African wildcat subspecies *Felis lybica lybica*. Its global distribution alongside humans testifies to its successful adaptation to anthropogenic environments. Uncertainty remains regarding whether domestic cats originated in the Levant, Egypt or elsewhere in its natural range, and on the timing and circumstances of their dispersal into Europe. By analysing 87 ancient and modern cat genomes, we demonstrate that domestic cats did not spread to Europe with Neolithic farmers, as previously thought. Conversely, our results suggest that they were introduced to Europe over the last 2,000 years, most likely from North Africa. We also demonstrate that a separate earlier (1st millennium BCE) introduction of wildcats from Northwest Africa originated the present-day wild population in Sardinia.

## Introduction

The domestic cat (*Felis catus*) has a complex evolutionary history that ultimately resulted in a close association with humans (*1*). As one of the most successful mammalian domesticates, it has a worldwide distribution that includes remote islands. When including feral cats, the global population of *Felis catus* is approaching one billion (*2, 3*).

Despite their ubiquity, the timing and circumstances of cat domestication and dispersal remain uncertain. This is due to a variety of factors including: the paucity of felid remains in archaeological contexts, the difficulty of assigning species and domestication status to individual skeletal elements (since wild and domesticated forms overlap in size and morphology (*4*–*6*)), and the limited number of ancient and modern genomes analysed so far. As a result, current hypotheses regarding when, where and how cats were domesticated are poorly supported by empirical evidence.

Genetic findings from present-day wild and domestic cats have demonstrated that the African wildcat (*Felis lybica lybica*), currently distributed across North Africa and the Near East, is ancestral to all modern domestic cats (*7*). The burial of a cat in association with the complete skeleton of a man dated to ∼7,500 BCE in Cyprus (*8*) led to the hypothesis that cats had been domesticated in the Levant during the Pre-Pottery Neolithic (9,600-7,000 BCE), where they may have acted as pest-controllers in early farming communities (*9*). A more traditional view, built upon iconographic and funerary evidence, pointed to Pharaonic Egypt as the original place of cat domestication (*10, 11*). This process has been dated to the 2nd millennium BCE, but attempts to tame cats may date as far back as the Predynastic period ∼3,700 BCE (*12*).

A clear understanding of the dispersal of domestic cats is hampered by the limited knowledge of the natural distribution ranges of African (*F. l. lybica*) and European (*Felis silvestris*) wildcats in areas where the two species either overlap or were adjacent, especially in the past. Ancient mitochondrial DNA (mtDNA) evidence suggested that *F. l. lybica* naturally occurred in Anatolia and Southeast Europe ∼10,000 years ago (*13*). The spatiotemporal distribution of maternal lineages in Europe, Southwest Asia and North Africa, suggested that domestic cats carrying the haplogroup IV-A were initially dispersed by people from Anatolia via a southeastern route into Central Europe as early as the mid-5th millennium BCE, possibly alongside a Neolithic expansion (*13, 14*). No later than the 1st millennium BCE, during Classical Antiquity, domestic cats carrying the haplogroup IV-C from Egypt were dispersed across Europe and Southwest Asia (*13, 14*).

Past hybridization between populations of wild *F. l. lybica* and *F. silvestris* may have naturally occurred (*13, 15*), thus leading to discordant nuclear and mtDNA evolutionary reconstructions, as has been shown in present-day European wildcats (*7, 16*). Recent genome-wide data from ancient and modern cats have questioned the use of mtDNA to draw conclusions about former natural distributions of wildcats and the spread of domestic cats (*15*).

The reconstruction of cat dispersal is further complicated by the uncertain origins of wild *F. l. lybica* populations on the Mediterranean islands of Sardinia and Corsica (*17*). Though it is claimed that their origin can be traced to Near Eastern cats introduced by farmers during the Neolithic (*18*– *20*), archaeological evidence suggests a much later introduction (*21*), and an analysis of morphological markers concluded that the modern populations shared an affinity with North African wildcats (*22*). In addition, morphological (*23*) and genetic analyses (*20*) of present-day specimens have indicated that they are distinct from domestic cats. Combined, these lines of evidence suggest that wildcats in Sardinia and Corsica are not a feral form of domestic cats, but a separate wildcat lineage. The poor documentation of small carnivores in the archaeological record of Sardinia and Corsica (*21*), however, and the lack of genetic data from African and Near Eastern wildcats have made it difficult to test this hypothesis and to identify their source populations.

Within mainland Europe, a more recent zooarchaeological reassessment of cat remains combined with mtDNA signatures and direct radiocarbon dating has challenged the assumption that domestic cats were already present in Europe before the 2nd millennium BCE, thus pointing to Dynastic Egypt as the more likely cradle of cat domestication and dispersal (*24*).

Here, to assess the timing and possible routes of domestic cat dispersal into Europe, as well as their relationship with wildcats from Sardinia, we conducted paleogenomic analyses of 225 ancient cat specimens from 97 archaeological sites in Europe and Anatolia (Table S1, S2). We generated 70 low-coverage (∼0.07-fold to ∼1.39-fold) ancient genomes (Fig. 1A, Table S3) spanning a period of more than ten millennia from the 9th millennium BCE to the 19th century (c.) CE, and 17 low-to mid-range coverage (∼0.7-to ∼18-fold) genomes of present-day and museum specimens of wildcats from Italy (including Sardinia), Bulgaria and North Africa (Table S4). To ensure a high degree of temporal resolution for the reconstructed genetic variation, we directly radiocarbon-dated 37 cat remains from 30 archaeological sites (Table S5).

**Fig. 1.**
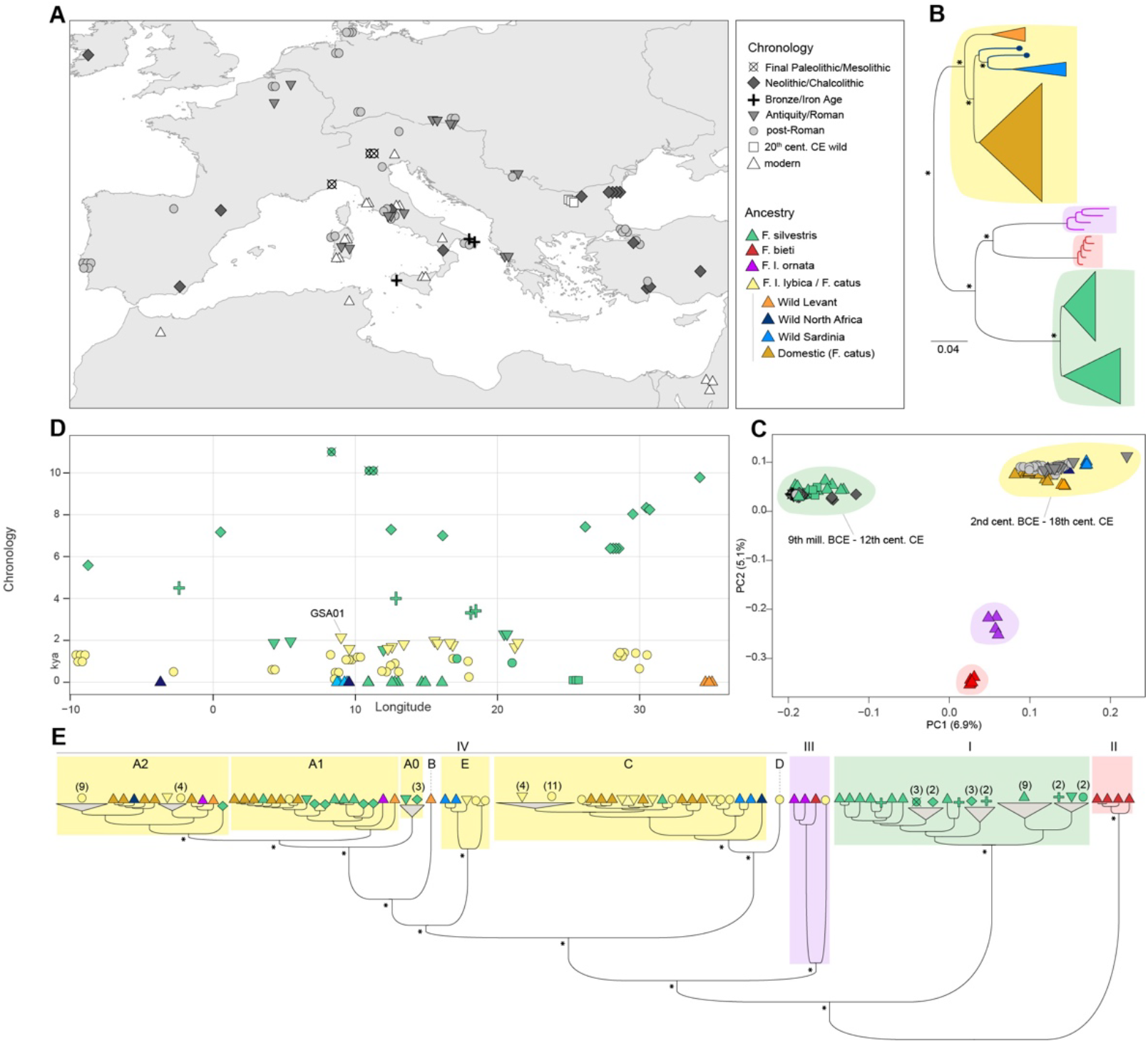
Samples and spatiotemporal distribution of cat ancestries. **(A)** Geographic provenance of cats for which ancient and modern genomes were generated in this study. The approximate chronology of each sample is coded with shapes and gray scale colored as in the legend panel (chronology). Three samples from Israel previously analyzed (*15*) are also reported in the map. **(B)** Genome-wide autosomal SNPs maximum-likelihood (ML) tree of high- and mid-range coverage modern cats from this study and the literature. Support values are based on 500 bootstrap replicates; the asterisks indicate bootstrap values=100. Main clades are colored as in the legend panel (ancestry) **(C)** Principal component analysis built by projecting low coverage samples onto the coordinate space defined by modern mid-to high-range coverage wild and domestic cats. Present-day samples are color-coded based on the clades of figure 1B, while symbols and colors of ancient samples are chronologically coded as in figure 1A. **(D)** Spatiotemporal representation of cat ancestries where each dot represents a cat. Symbols are as in the geographic map (Fig. 1A). Colors are as in the ML tree (Fig. 1B) for the present-day wildcats, and in the ancient cats they represent genome-wide ancestries as inferred by the clusters of the PCA (Fig. 1C). **(E)** Bayesian tree of complete mtDNAs of ancient cats for which genome-wide data were obtained in this study, and present-day wild and domestic cats from this study and the literature. Symbols are as in the geographic map (Fig. 1A). Colors are as in the autosomal ML tree for the present-day samples (Fig. 1B) and in the ancient cats they represent genome-wide ancestries as inferred by the clusters of the PCA (Fig. 1C). Clades with four or more cats were collapsed and the total number is reported in brackets. Posterior probabilities between 0.86 and 1 are indicated by an asterisk in the main nodes.

### Distinct North African ancestries in present-day domestic cats and Sardinian wildcats

To provide an updated phylogeny of present-day wild and domestic cats, we first reconstructed a Maximum-Likelihood (ML) autosomal tree (Fig. 1B, Fig. S1) using the genomes of present-day wildcats generated in this study along with samples from the literature (Table S6). We identified four main clades corresponding to the different wildcat taxa investigated, which corroborates previous studies illustrating that *F. l. lybica* is a sister group to *F. l. ornata, F. silvestris* and *F. bieti* (*15*). This analysis also showed that *F. l. lybica* populations are geographically structured with two distinct, well-supported (bootstrap=100) clades corresponding to Levantine and African wildcats. Domestic cats form a distinct clade that is sister to the African wildcats, thus suggesting a closer genetic proximity to these than to the modern Levantine population. The present-day Sardinian wildcats clustered with the African wildcats and formed a monophyletic group with the Moroccan specimen.

We then assessed population structure in wild and domestic populations using Multidimensional Scaling (MDS) based on identity-by-state (IBS) (Fig. S2) and an unsupervised ADMIXTURE analysis (Fig. S3). Both the MDS and ADMIXTURE at K=4 defined clusters that mirrored the main clades in the ML tree. Higher K values (K=5 to 7) identified substructure within the European wildcats and discriminated *F. l. lybica* from *F. catus* groups. Domestic cats shared more genetic affinities with wildcats from Africa, particularly with the Tunisian one, than with those from the Levant and Sardinia (K=5 to 7). *F. l. lybica* ancestry was better resolved at K=8, where a distinct component was assigned to the wildcats from the Levant, and the Sardinian wildcats shared more ancestry with the Moroccan sample. This was also illustrated by the MDS, in which Moroccan and Sardinian wildcats clustered together, consistent with the ML tree topology. Comparable results were found by conducting admixture analyses on several subpanels of populations including only *F. l. lybica* wildcats (n=8) and different random subsamples (n=3) of domestic cats (Fig. S4, Supplementary text), to account for potential biases due to uneven sampling.

To further test whether present-day domestic cats are genetically closer to African than to Levantine wildcats, we employed outgroup-*f*_*3*_ and *f*_*4*_ statistics using the jungle cat (*F. chaus*) as outgroup. Our results showed that all the present-day domestic cats shared more genetic drift with the present-day Tunisian wildcat (Fig. S5). Outgroup-*f*_*3*_ and *f*_*4*_ statistics conducted on the Sardinian wildcats showed that they shared more drift with the present-day Moroccan individual than with domestic cats (Fig. S6, S7). Although preliminary, due to the limited sample of wildcat genomes (three from the Levant and two from North Africa), these results allow hypothesizing that domestic cats and Sardinian wildcats were derived from two genetically distinct populations in North Africa (represented in our dataset by Tunisian and Moroccan wildcats, respectively).

### Domestic cats did not spread to Europe during the Neolithic

Previous studies that made use of diagnostic SNPs from a fragment of the mitochondrial genome suggested that clades IV-A and IV-B, typical of *F. l. lybica* populations from the Near and Middle East, were amongst those that dispersed from Anatolia to central via southeastern Europe during the Neolithic (*13, 14*). Recently, this wave of Neolithic cat dispersal into Europe has been questioned by the demonstration that Eastern European wildcat (*F. silvestris*) populations naturally possess some degree of Near Eastern wildcat ancestry due to admixture along a hybrid zone during the Late Pleistocene or Early Holocene (*15*). This study concluded that mtDNA haplogroups IV-A and B are thus not suitable diagnostic markers for tracing the dispersal of domestic cats. To address this, we analysed genome-wide transversions and tested whether samples from Neolithic/Chalcolithic Bulgaria (n=4) and Neolithic Anatolia (n=4) (ranging 9,900-4,300 BCE) in which mtDNA IV-A was previously detected (*13*), were European wildcats.

We found that, at the nuclear level, all the samples (n=22) that dated from the 9th millennium to the 3rd c. BCE project onto the European wildcat (*F. silvestris*) PCA cluster (Fig. 1C-D, Fig. S8). Crucially, this cluster includes Neolithic and Chalcolithic cats from Anatolia (n=4) and Bulgaria (n=4) dated from the 7th to the 5th millennium BCE that possessed a IV-A mtDNA (Fig. 1E, Fig. S9, S10), typically associated with *F. l. lybica*. All the other samples from across the rest of Europe dating from the 9th millennium to the 3rd c. BCE (n=14), as well as one cat from Chalcolithic Bulgaria possessed both nuclear European wildcat ancestry and a mtDNA haplogroup I that is characteristic of *F. silvestris* (*7*). The mtDNA clade IV-A was also found in wildcats dating to the Roman era (Belgium and Italy, n=2), and Modern periods (Germany, n=3, Bulgaria, n=1) (*13*), thus implying the persistence of IV-A in southeastern and central European wildcats for the past seven millennia.

Conflicting mitochondrial and nuclear evolutionary histories (i.e. mitonuclear discordance) may be due either to hybridization or incomplete lineage sorting (ILS) (*25*). In this case, given the clear biogeographic pattern of mitonuclear discordance as illustrated in the ancient European wildcats (Fig. S11), ILS can be ruled out (*26*). By considering the effective population size estimated with a Bayesian Skyline plot and from the literature (Supplementary text, (*27*)), assuming neutrality and a complete mtDNA turnover in Neolithic Anatolian wildcats (∼8,000 BCE), we computed the mean time required for haplogroup IV-A to reach fixation in the ancient Anatolian wildcat population (*28, 29*). Our results show that *F. l. lybica* IV-A mtDNA would reach fixation in not less than about 5,000 years in a small European wildcat population (N_e_=1000), thus placing the admixture event not earlier than the Late Pleistocene (Supplementary text, Supplementary table S8. Fig. S12). Additional genome data from modern wildcats in Anatolia and the Caucasus, which are currently lacking, will help to test different admixture scenarios.

These results demonstrate that cats previously found to possess a mtDNA clade IV-A in Neolithic Anatolia (∼8,000-6,000 BCE) and Neolithic, Chalcolithic Southeast Europe (∼5,500-4,000 BCE) (*13*) were not *F. l. lybica*/*F. catus* cats introduced by humans, but *F. silvestris* wildcats whose ancestors hybridized with *F. l. lybica*. This implies that European wildcats have been present in these regions for at least the last ten millennia (*17, 30*). Our genomic data thus corroborate previous evidence showing that haplogroup IV is not restricted to *F. l. lybica*, and that the matrilineage IV-A alone should not be used to infer either ancient wildcat distributions, or human-mediated translocations across continental Europe in the past (*15*). Although cats are traditionally viewed as commensals that frequented the human niche in the Neolithic Levant, recent zooarchaeological evidence suggests that wildcats were also intensively hunted and exploited for food and fur (*31*). Regardless of the nature of the Neolithic cat-human relationship, our data indicate that this did not lead to the spread of *F. l. lybica*/*F. catus* cats into Europe.

Of the 22 *F. silvestris* samples dating from the 9th millennium to 3rd c. BCE *F. silvestris* samples analyzed here, 20 originate from either archaeological settlements (i.e. Neolithic and Chalcolithic Anatolia and Bulgaria, Bronze Age Italy and Spain, Hellenistic Greece, n=15) or cave sites with evidence of anthropogenic activity (Late Paleolithic, Mesolithic and Neolithic Italy, n=5). The relationship between humans and wildcats in Europe was possibly based on exploitation for fur (*32, 33*), and food, as suggested by the Mesolithic samples from Galgenbühel/Dos de la Forca (*34*). More complex socio-cultural and symbolic relationships should not be discounted (*35*), considering the wildcat remains analyzed here from Bronze Age Partanna (Sicily, Italy) collected in a bell-shaped vase (*36*), and a feline clay head from Chalcolithic Bulgaria (*37*).

### Human-mediated dispersal of F. l. lybica from the 1st millennium BCE

The earliest sample in our dataset that possesses *F. l. lybica*/*F. catus* nuclear ancestry is a cat from the site of Genoni (GSA01) in Sardinia that has been directly radiocarbon dated to the 2nd c. cal. BCE (Fig. 1C-D). In a PCA conducted specifically on *F. lybica/F. catus* samples (Fig. 2A), this individual clusters with three present-day Sardinian wildcats, and their genetic proximity was confirmed by outgroup-*f*_*3*_ statistics that we used to estimate the shared drift between GSA01 and ancient and modern *F. l. lybica*/*F. catus* samples. We found the highest *f*_*3*_ values with present-day wildcats from Sardinia (Fig. S13). These results suggest that the origins of present-day Sardinian wildcats can be traced back over 2,000 years ago to an ancestral population to which GSA01 belonged.

**Fig. 2.**
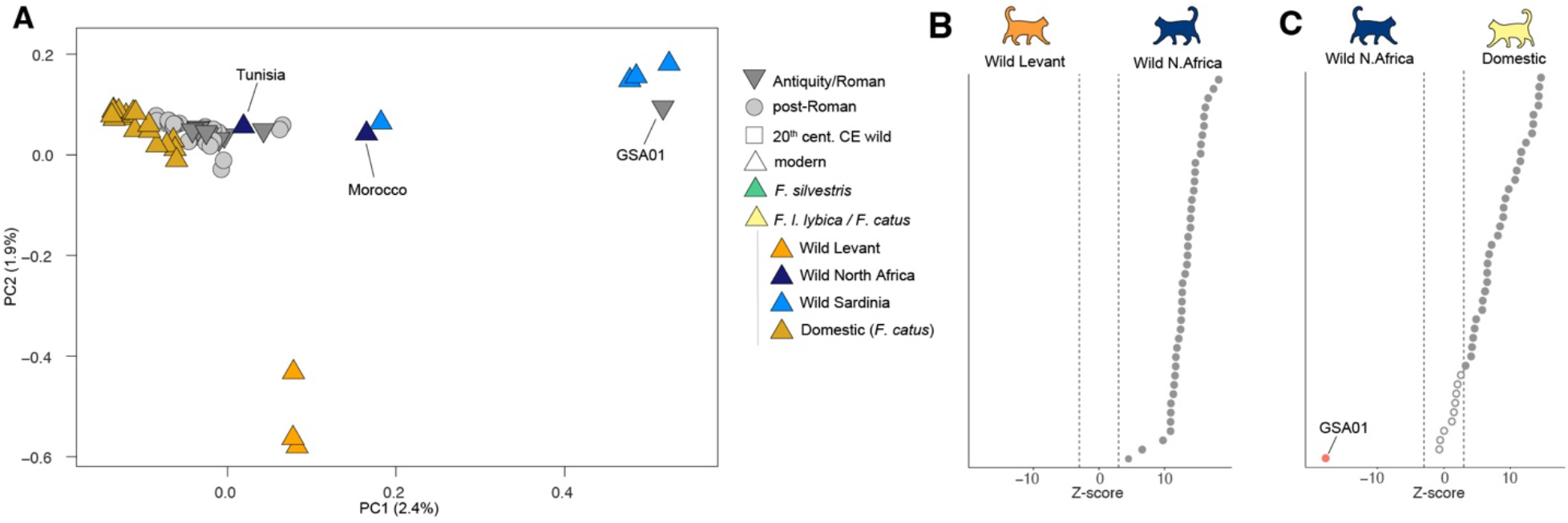
Genome-wide ancestries of ancient domestic cats. **(A)** Principal component analysis built by projecting low coverage samples onto the coordinate space defined by modern high- and mid-range coverage *F. l. lybica* and domestic (*F. catus*) cats. Present-day samples are color-coded based on the clades of figure 1B, while symbols and colors of ancient samples are chronologically coded as in figure 1A. **(B)** z-scores resulting from the test *f*_*4*_(*F. chaus*, ancient *F. l. lybica*; modern Levant *l. lybica* wildcats, modern African *F. l. lybica* wildcats). Each dot represents an ancient cat analyzed in this study. Significant values are reported as filled dots and non-significant values as empty dots. The dashed lines indicate the significance threshold (±3). **(C)** z-scores resulting from the test *f*_*4*_(*F. chaus*, ancient *F. l. lybica*; modern African *F. l. lybica* wildcats, modern domestic cats). Colors and significance are as in Fig. 1B; the negative significant value of sample GSA01 is reported in red.

All other archaeological cats from Europe and Anatolia (n=42) that cluster with *F. l. lybica*/*F. catus* are dated from the 1st c. CE onwards. These samples include three specimens from Roman to 19th century Sardinia and group together with present-day domestic cats (Fig. 2A). This was corroborated by *f*_*4*_ statistics that we used to assess patterns of shared drift in *F. l. lybica*/*F. catus* between ancient and present-day wild and domestic cats. Similarly to GSA01, all the ancient domestic samples share more affinities with modern African wildcats than with their Levantine relatives (Fig. 1D, Fig. S14). In contrast to GSA01, ancient domestic cats were more closely related to present-day domestic cats than to North African wildcats (Fig. 1E).

These results suggest that the human-mediated dispersal of *F. l. lybica*/*F. catus* into Europe occurred at least twice. The first translocation of *F. l. lybica* to Sardinia from a source population in Northwest Africa dates to the second half of the 1st millennium BCE at the latest, as is evidenced by the 200-50 cal. BCE GSA01 sample. It is probable that this first population introduced to Sardinia were wildcats that remained largely isolated from their domestic counterparts that were introduced from the Roman Imperial era onwards.

Phoenicians and later the Punic people established and maintained colonies in Northwest Africa, Sardinia and the southeastern Iberian Peninsula from the 9th c. BCE until their defeat by the Romans in the Third Punic War in 146 BCE (*38, 39*). Because Sardinia had been under Roman supremacy since 238 BCE (*40*), the translocation of *F. l. lybica* to Sardinia could have been carried out by either the Phoenicians and Punic people or by the Romans before the Imperial period starting 27 BCE (*41*).

Subsequently, since the Roman Imperial era, cats more genetically similar to present-day domestic cats were spread across Europe from a distinct North African population. The earliest sample carrying the ancestry found in present-day domestic cats was dated to 50 cal. BCE-80 cal. CE, from the site of Mautern, in Austria. Cats with this ancestry were then identified in Italy (n=4) and in Roman Imperial military sites along the Danube Limes in Austria and Serbia (n=6) (*42*–*45*), confirming previous claims that the Roman army and its entourage played a role in spreading domestic cats to central and eastern Europe (*1, 46*). This dispersal reached northern Europe relatively early in the Roman Imperial era, as testified by the genome of a cat from the site of Fishbourne, Britain, dated to 24-123 cal. CE (*15*). The ancestry typical of present-day domestic cats is then found continuously in all cats from Europe and Anatolia from the Byzantine era (n=7), the Medieval (n=23) and post-Medieval periods (n=2) until present-day (Fig. 1D).

The identification of the ancestral population that sourced the domestic cat dispersal remains challenging due to the lack of ancient and modern genetic data from African wildcats. Egypt is traditionally considered the core region for the domestication of cats and their subsequent spread (*11*–*13*). The increasingly intensive relationship between humans and cats in the Nile Valley is demonstrated by their role in the Egyptian state cult of *Bastet*, and as rodent-killers in the agricultural economy (*10*). This hypothesis is supported by the presence of the mtDNA haplogroup IV-C in eight out of 10 Roman Imperial cats, a haplogroup previously found in five Ptolemaic to Roman Egyptian mummies (*13*). Our observation that ancient and modern domestic cats share higher genetic drift with the Tunisian wildcat, however, may suggest an additional, if not alternative source population from the west/central North African coast of the Mediterranean. Large ports that served a fertile agricultural hinterland, as was the case in Phoenician-Punic Carthage (*47*), may have fostered the synanthropic association of cats with humans, and their subsequent dispersal across the Mediterranean. Crucially, genome data from modern and ancient cats from Egypt, which are currently lacking, will allow these two hypotheses to be tested.

Different factors may have driven the translocation of cats to new cultural settings. Arguably, the association between cats and the cult of the goddess *Bastet* elevated the species’ prominence beyond Pharaonic Egypt, influencing Phoenicians (*48*), Greeks and Romans (*49*) alike. As with the human-mediated dispersal of domestic chickens (*50, 51*) or fallow deer (*52*), the initial translocation of cats could have been religiously motivated (*24*). Dispersal trajectories may also have been driven by the economic value of cats as pest controllers in maritime voyages, in view of the extensive trade network of Carthage and the role of Egypt as a grain supplier to the Roman Empire (*53*).

Overall, the 1st millennium BCE marks a pivotal period for the introduction of cats into Europe. This was also corroborated by a Bayesian Skyline plot of *F. l. lybica* mtDNA lineages showing a rapid expansion starting 2,500 years ago (Supplementary text and Supplementary figures S15). Of three cats dated to this chronological range for which the genome was sequenced, two are *F. silvestris* from Greece, the only *F. l. lybica* being the 2nd c. BCE cat from Genoni. A cat with *F. l. lybica* IV-A1 mtDNA radiocarbon dated to the 8th-5th c. BCE from eastern France (Entzheim, Alsace) was recently suggested to be a European wildcat based on metric analysis (*24*). Yet, zooarchaeological evidence of five neonatal kittens buried in single deposit radiocarbon dated to 364–121 cal. BCE from Gussage All Saints, southern Britain, points to introductions of domestic cats in more northern parts of western Europe during the Iron Age (*15, 24*), which could be associated with Mediterranean trade routes during Classical-Antiquity (*54*). Genome data from these and other Iron Age contexts in Europe are needed to better understand the chronological, spatial, and cultural settings of their human-mediated dispersal.

### Ancient and recent gene flow in wild and domestic cats

A recent study revealed that following the introduction of domestic cats to Europe >2,000 years ago, gene flow between indigenous wildcats and introduced domestic cats was limited, and the proportion of wildcat ancestry in domestic cats ranged from 0 to 14% (*15*). Analyses of domestic introgression in present-day wildcats from Scotland, however, demonstrated that much greater rates of hybridization have taken place over the last 70 years (*55*), possibly owing to a decline in the wildcat population.

We assessed the interactions between wild and domestic cats across a temporal and geographic range stretching from western Europe to central Anatolia. In the PCA (Fig. 1C, Fig. S15), we observed that ancient cats from Anatolia and the Balkans and modern European wildcats from Scotland, Italy and Germany spread out of the core of the European wildcat cluster, which may reflect a history of admixture.

We tested for the incorporation of *F. l. lybica*/*F. catus* ancestry into ancient and modern European wildcats by computing D-statistics. As a reference for European wildcats, we used a group (n=5) of ancient samples from Spain and northern Italy dated from 9th to the 3rd millennium BCE (from the Mesolithic to the Chalcolithic) that we ascertained not to carry any *F. l. lybica*/*F. catus* ancestry (Supplementary text, Fig. S16). As admixture sources we used the wildcats from the Levant and a domestic cat (Ocicat breed) that was devoid of *F. silvestris* ancestry (Supplementary text, Fig. S17). In nine of the 11 samples from Anatolia and the Balkans dated from the Neolithic to the Iron Age (8th millennium to 3rd c. BCE), we detected gene flow when using Levantine wildcats (Fig. 3A). When using the domestic source, only five were found to yield detectable levels of gene flow (one from Turkey, three from Bulgaria and one from Greece). Regardless, D-values in ancient Anatolian and Balkans samples were found to be systematically higher in all instances when using the Levantine wildcats as the source of admixture. This pattern is less evident in present-day European wildcats, in which the levels of gene flow from either sources (Levant and domestic) were more similar.

**Fig. 3.**
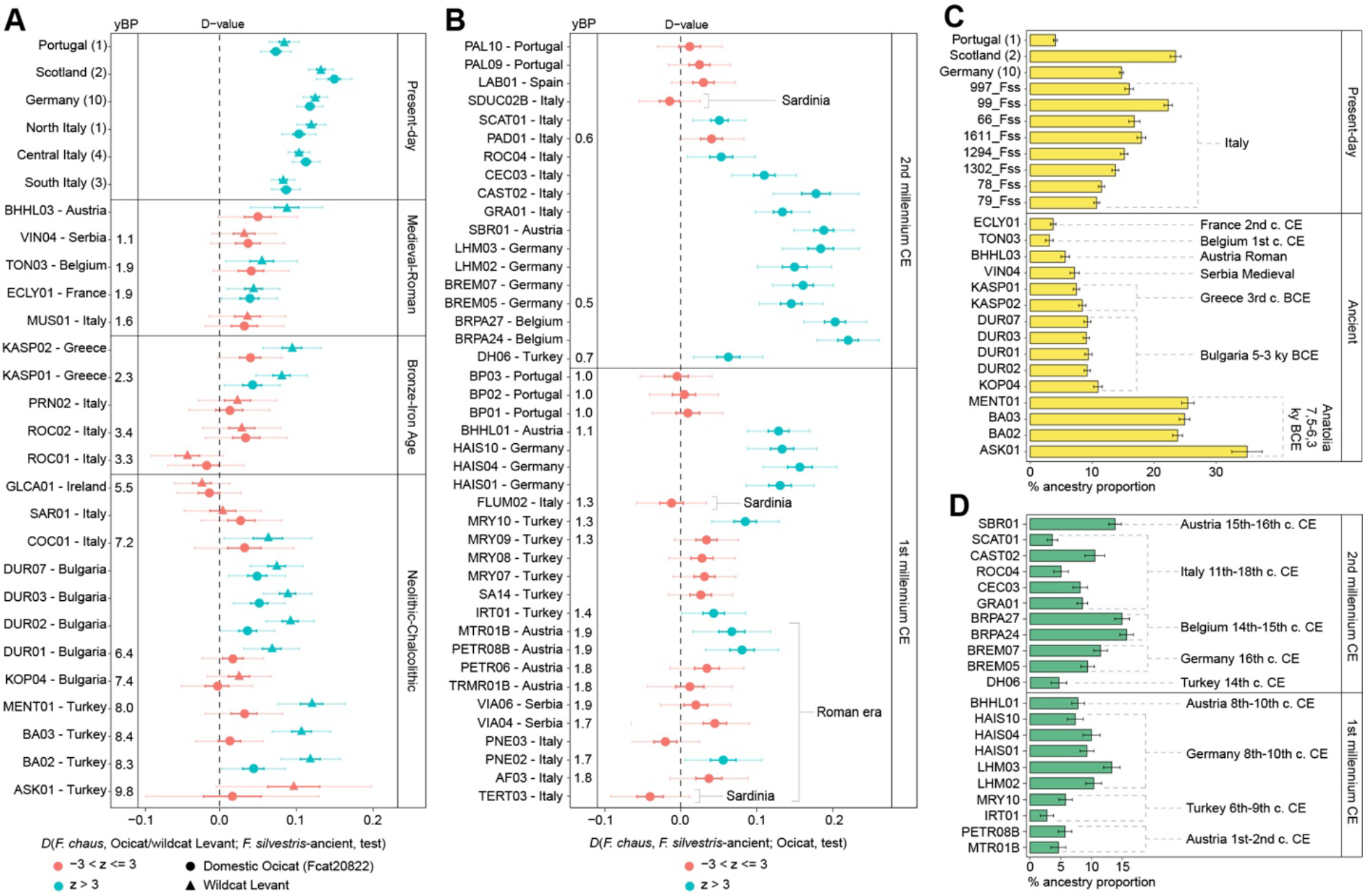
Patterns of gene flow between wild and domestic cats across time. **(A)** *D* statistics resulting from *D*(*F. chaus*, Ocicat/wildcat Levant; *F. silvestris*-ancient, test) where gene flow into ancient and modern European wildcats was tested against two distinct sources, a domestic one (*F. catus*, represented by the Ocicat breed), and a wild one from the Levant (*F. l. lybica*, represented by three wildcats from Israel). Colors indicate z-scores significance (±3) and symbols represent the source used, as in the legend. Error bars represent ±1 and ±3 (lighter blue/red color bar) standard deviations. In radiocarbon dated samples the age in years Before the Present (yBP) is reported. **(B)** *D* statistics resulting from *D*(*F. chaus, F. silvestris*-ancient; Ocicat, test) to test gene flow from the European wildcats into ancient domestic cats. Colors indicate z-scores significance (±3), as in the legend. Error bars represent ±1 and ±3 (lighter blue/red color bar) standard deviations. In radiocarbon dated samples the age in years Before the Present (yBP) is reported. **(C)** Proportion of *F. l. lybica* ancestry in ancient and modern European wildcats from this study and the literature returning significant values of gene flow in the *D* statistics. The proportions were computed with *f*_*4*_-ratio following the model described in Fig. S19. Cats from Scotland and Germany were pooled, the numerosity is reported in brackets. More details are in the Supplementary text. **(D)** Proportion of *F. silvestris* ancestry in the ancient domestic cats returning significant values of gene flow in the *D* statistics of panel 3B. The sample PNE02, which returned a slightly significant *D* value, is excluded from the figure as it did not show detectable proportions of *F. silvestris* ancestry. The proportions were computed with *f*_*4*_-ratio following the model described in the Fig. S19.

We quantified the degree of gene flow by estimating admixture proportions with *f*_*4*_-ratio (Fig. S19). They revealed ∼7% to ∼11% of *F. l. lybica*/*F. catus* ancestry in the ancient *F. silvestris* samples from the Balkans, and higher proportions in the Neolithic *F. silvestris* wildcats from western (∼24%) and central Anatolia (∼34%) (Fig. 3C). Modern wildcats from Italy exhibited ∼10 to ∼22% of *F. l. lybica*/*F. catus* ancestry, thus showing intermediate values between wildcats from Germany (∼14%) and Scotland (∼23%).

Overall, our data reveal two distinct sources of gene flow in European wildcats: an ancient one from *F. l. lybica* populations, and a more recent one from domestic cats following their dispersal into Europe. This supports that admixture first occurred in the Late Pleistocene, prior to the introduction of domestic cats in Europe. This likely stemmed from a contact zone between the two species located either in the Levant or in eastern Anatolia. This admixture led to the acquisition of *F. l. lybica* mtDNAs via mitochondrial capture, with possible unidirectional mating of local European wildcat males with Near Eastern wildcat females (*56*). As previously suggested (*15*), this scenario would explain the mitonuclear discordance detected in Neolithic and Chalcolithic cats from Southeast Europe and Anatolia, as well as the presence of a cat in possession of a IV-A1 mtDNA in Mesolithic Romania (7,700 BCE, (*13*)). Unidirectional mating is also supported by previous genetic research showing a high occurrence of European wildcats carrying a *F. l. lybica* mtDNA (26%) as opposed to much rarer occurrence of domestic cats with *F. silvestris* mtDNA (0.4%) (*7*).

Lastly, the *D* and *f*_*4*_-ratio tests for the incorporation of *F. silvestris* ancestry into ancient domestic cats (Fig. 3B and 3D) showed detectable levels of gene flow from European wildcats in only two of 10 cats dated to the Roman era (up to ∼6% in two cats from Austria dated to the 1st-2nd c. CE). In the Middle Ages, the degree of introgression increased slightly, and *F. silvestris* ancestry rose up to ∼15% in domestic cats from Mediterranean and continental Europe (n=16), apart from all the Iberian and Sardinian samples. Our analysis thus suggests that following their introduction to Europe in the Roman era, domestic cats started to incorporate increasing levels of wildcat ancestry over time. However, this pattern does not extend to the Iberian Peninsula, where we were unable to detect gene flow in any of six Medieval samples we investigated. Overall, the diachronic pattern of admixture that we detected supports the hypothesis that habitat degradation and an encroaching human presence disrupted the ecological and spatial separation between wild and early domestic cat populations in Europe (*15*). Conceivably, deforestation and the expansion of agriculture over time led to a greater overlap of the home ranges and thus to increased gene flow between wild and domestic cats.

### Conclusions

Our results demonstrate that the dispersal of present-day domestic cats can be traced back not to the Neolithic or from the Fertile Crescent, but instead several millennia later and most likely from North Africa. Mediterranean civilizations during the 1st millennium BCE were the main agents of *F. l. lybica* translocation, which occurred in two waves involving genetically distinct populations of North African origin. The first dispersal most likely featured wildcats from Northwest Africa that were introduced to Sardinia and founded the present-day *F. l. lybica* populations in the island. The second wave established the genetic pool of modern domestic cats.

Our results offer a new interpretative framework for the geographic origin of domestic cats, suggesting a broader and more complex process of domestication that may have involved multiple regions and cultures in North Africa. Efforts should continue to pinpoint the original source population(s) of present-day domestic cats, and to clarify the cultural and socio-economic processes that led to their domestication and global dispersal. Currently, the genetic diversity of wildcat populations in North Africa and the Levant is represented by five individuals (two from Morocco and Tunisia, three from Israel). For this reason, it is fundamental to generate more ancient and modern genomes from these regions in the future, and particularly from Egypt.

## Supporting information

Supplementary test and figures

## Acknowledgments

We are grateful to Joachim Schultze for granting permission to sample the collection at The Schleswig-Holstein State Museums Foundation Schloss Gottorf and Corinna Mayer for support in the sampling procedure; to Fiona Beglane (Atlantic Technological University) for her help in providing the cat bones from Glencurran cave; to José Carlos Brito (CIBIO, University of Porto) for granting the access to the North African wildcat DNA samples; to Luca Lapini ed Egidio Mallia for providing the wildcat samples from the Alps and southern Italy; to Eduard Pollhammer for permission for sampling bones from Petronell-Carnuntum and the collections of the province of Lower Austria (Hainburg); to Stefan Groh and Helga Sedlmayer for granting access to the material from Mautern/Danube (ÖAI/ÖAW Vienna). We also wish to thank Andrew Kitchener for his comments on a manuscript draft; to Gene Shev (University of Rome Tor Vergata) for comments and language revision; to Cristina Martinez for co-supervision of MDM’s doctoral project, and Gabriele Scorrano for discussion and feedback on data analysis and results. We thank Paolo Boscato, Francesco Boschin, Anna Gręzak, Giovanni Boschian, Marco Romboni, Maria João Valente, Vera Pereira, Lluís Lloveras, Jordi Nadal for providing samples reported in Table S1. We also wish to thank Soprintendenza Archeologia, belle arti e paesaggio per le province di Sassari e Nuoro for granting permission to analyze the archaeological samples from Sardinia. Bioinformatic analyses were performed on the Galileo100 high performance computing resources of CINECA, with the support of Elixir-Italy under the HPC@CINECA program and the CINECA award under the ISCRA initiative.

## Author contribution

Conceptualization: CO, WVN, JP. Data curation: WVN, BDC, CO. Formal analysis: MDM, CO. Funding acquisition: CO, NS. Investigation: MDM, VR, PS, MB, TDC, BDC, WVN, CO. Methodology: MDM, VR, PS, CO, SG. Project administration: TDC, CO. Sample provision: FA, PCA, HB, EC, ECer, JDGM, CD, MD, IF, LG, IGS, HCK, GCK, MM, RM, CM, MM, VO, TO, MP, BPU, GR, MR, LS, KS, ABS, US, AS, GS, NSp, AT, VT, ST, SV, UW, BW, FM, RC. Supervision: CO, WVN, JP. Visualization: CO, MDM, PS. Writing – original draft: MDM, CO. Writing – review & editing: GL, JP, WVN, LF, NS, SD, BDC, NSp, VR, PS, MB, FA, CD, MD, IGS, HCK, GKK, RM, CM, MR, KS, US, ST, SV, UW, BW, FM, RC.

## Data and materials availability

Genomic sequencing data are available through NCBI SRA Project accession no. PRJEB81815.

## Supplementary Materials

Supplementary Text

Materials and Methods

Figs. S1 to S20

References (*50*–*145*)

Datasets ST1 to ST8

