## Supplementary test and figures for "The dispersal of domestic cats from Northern Africa and their introduction to Europe over the last two millennia"

De Martino et al.

#### Supplementary Material

#### Supplementary text

This section describes the archaeological sites from which samples successfully used for genome-wide data analysis originate.

##### Turkey

**Aşıklı Höyük** (contact: Hijlke Buitenhuis). Aşıklı Höyük (AH) sits directly on a floodplain of the Melendiz River (elevation 1,119 m). AH is an artificial mound composed of 16 m of anthropogenic deposits. It is the earliest known preceramic Neolithic mound site in Central Anatolia. The oldest Levels, 4 and 5, spanning 8,200 to approximately 9,000 cal. BCE, associate with round-house architecture and arguably represent the birth of the Pre-Pottery Neolithic in the region (*57*). Of particular interest from a faunal viewpoint is the discovery of extensive dung layers at the site, implying that people held animals captive in the village from the earliest habitation onwards. One cat was analyzed in this study. Radiocarbon dating conducted on this cat at KIK-IRPA provided an age of 8,745±37BP (RICH-25375) corresponding to 7,950-7,610 cal. BCE (95.4%).

**Bademağacı** (contact: Bea De Cupere). Bademağacı Höyük is located in the south of the Lake District (Pisidia), southwest Anatolia, about 50 Km north of Antalya. The mound lies at an altitude of 780m above sea level (asl) just north of the pass (ancient Klimax) in the Taurus Mountains, which links this region with the coastal plain of Pamphylia. The site was excavated from 1993 until 2010, and a study of the faunal material was published (*58*). Three ancient cats identified as wild with traditional metric analyses were analyzed. They all yielded genome-wide data and were dated to the Neolithic based on radiocarbon dates or the associated archaeological context from which the bones were unearthed. Radiocarbon dating conducted on two cats provided an age of 7,496±32BP (RICH-34506) and 7398±32BP (RICH-3450), the first corresponding to 6,430-6,240 cal. BCE (95.4%,) and the second to 6,390-6,090 cal. BCE (94%).

**Menteşe** (contact: Lionel Gourichon). This tell is located in the northern part of Western Anatolia next to the dried-out lake Yenişehir, about 25km south of the archeological site of Ilıpınar. The mound is four meters in height and has a diameter of ~150m. It encompasses three different strata. The youngest layer is associated with the Roman Imperial period, the other two date to the Bronze Age and the Chalcolithic. The oldest stratum of this latest phase was dated to 6,400 BCE and is older than Ilıpınar. One cat specimen was analyzed and returned genome-wide data. It was identified as probably wild based on bone traditional metrics context (*59*) and radiocarbon dated to 7,163±30BP (RICH-34361) corresponding to 6,073-5,986 cal. BCE (95.4%).

**Sagalassos** (contact: Bea De Cupere). The antique site of Sagalassos is located 7 km north of the small town of Ağlasun on a steep, south-facing slope of the Ağlasun Dağları (Western Taurus range, Southwest Turkey) at an altitude of 1,450 to 1,650m asl. In Imperial times it was the main city of ancient Pisidia and was continuously inhabited from the 5th century BCE until the earlier 13th century AD. Large-scale excavations from 1990 onwards yielded an enormous amount of animal remains, mainly representing consumption refuse of domestic livestock, waste of artisanal activities and remains of animal carcasses (*60*, *61*). Two cats were analyzed in this study. Based on stratigraphic sequencing of the site, they are dated to the Early Byzantine period.

**Demircihüyük** (contact: Joris Peters, Bea De Cupere). This mound is located 25 km northwest of the town of Eskişehir, at 860m asl on the northwestern fringes of the central Anatolian plateau.

The site, which was inhabited in the Early and Middle Bronze Age, yielded more than 80,000 identifiable mammal remains, including 12 cat bones (*62*). Five of these bones, found in levels K2 to P, are from the skeleton of a juvenile animal. In this study, an ulna of this individual was analyzed. According to its stratigraphic position the bone would date to the second half of the Early Bronze Age period, between about 2,690 and 2,570 BCE (*63*). However, one cat was analyzed in this study and was radiocarbon dated to 608±22BP (RICH-35360) corresponding to 1,300-1,370 cal. AD (74.5%) / 1,375-1,400 cal. AD (20.9%).

**Iznik - Roman theatre** (contact: Vedat Onar). The Roman theater of Iznik, located in the Marmara region along the eastern shore of Lake Iznik, was built during the reign of Emperor Trajan (98-117 AD) and remained in use as a theatre for two centuries. There is evidence of use for religious purposes thereafter, especially with the dominance of Christianity in the region in the 4th century AD, and in the 12th century AD two churches were built in the area that was then used as a cemetery. Ceramic kilns were built in the late Byzantine and Ottoman periods that remained in use until the 17th century AD. During the 2019-2020 excavations (*64*), a single bone of a subadult cat was found in a grid near the western façade of the theatre. Radiocarbon dating resulted in 1,448±22BP (RICH-35359) corresponding to 575-650 cal. AD (95.4%).

**Yenikapı, Marmaray exc**. (contact: Vedat Onar). Rescue excavations that started in 2004 at Yenikapı, an area of the European side of Istanbul, yielded numerous shipwrecks, amphorae and animal remains (*65*). The archaeozoological material includes food refuse as well as carcasses of complete animals that were disposed of in the ancient harbor built by Theodosius I in the late 4th century AD. Portus Theodosiacus lost its importance in the 7th century AD when the grain imports from Egypt stopped. In the 12th century AD the harbor was blocked when it was silted up by sediments from the Lykos river. Four cats were analyzed in the present study. Two were radiocarbon dated and belong to the 7th-8th century AD: 1,313±26BP (RICH-34529) and 1,350±25BP (RICH-34509), the first corresponding to cal. 650-710 cal. AD (48.9%) / 720-780 cal. AD (46.5%), and the second to 640-690 cal. AD (75.4%) / 740-780 cal. AD (20.0%).

#### Italy

**Roca vecchia** (contact: Michela Rugge). The archaeological site of Roca (Melendugno, Lecce) is located in southern Italy, on the Salento Adriatic coast in the area between San Foca and Torre dell'Orso, southeast of the city of Lecce. The archaeological area is developed on a promontory, known as the “castello-carrare” area, whose attestation is known from the Bronze Age (2nd millennium BCE). To this phase is referable the imposing work of fortification, erected to defend the settlement during the Middle Bronze Age (18th-14th centuries BCE) is repeatedly rebuilt in the subsequent phases of the Recent Bronze Age (14th-13th centuries BCE) and the Final Bronze Age (13th-10th centuries BCE). The site's involvement in the circuit of long-distance Mediterranean trade is evident from the abundant presence of Minoan-Mycenaean-derived pottery from levels dated to the 14th-13th centuries BCE. Occupation of the site continued during the Iron Age (10th-8th centuries BCE). The Messapian phase (4th-3rd century BCE) is also of great importance. In the Roman phase the area was used as an agricultural zone and villas and furnaces were built near the Messapic walls. A period of apparent abandonment marked the following centuries, until the Middle Ages. Three cats were analyzed in this study. Two of them were radiocarbon dated to 3,145±29BP (RICH-35362) and 3,051±23BP (RICH-35363), the first corresponding to cal. 1,500-1,380 cal. BCE (84.2%) / 1,350-1,310 cal. BCE (11.2%), and the second to 1,410-1,220 cal. BCE (95.4%). The third cat was dated to the Middle Age based on stratigraphic evidence.

**Castiglione in Sabina** (contact: Claudia Minniti, Jacopo De Grossi Mazzorin). The rural castle of Castiglione is in the Turano valley (Rieti province), Latium, central-west Italy. The castle was built in stone at the beginning of the 11th century AD (second period), following an earlier wooden structure (first period). It was rebuilt and restored several times until it was finally abandoned in the middle of the 11th century AD (third period). The faunal assemblage mainly falls within the last period. It consists for the most of large-sized domesticates exploited for meat-production and testifying to high-status eating habits (*66*). Due to the high-status of the site, the abundance of young animals seems to indicate that the tenderness of the meat was a concern of people eating at the castle. The high status of the site is also reflected by the presence of wild bird remains belonging to various species of wildfowl, including a medium- to large-sized raptor. The radius of one cat originating from the third period (11th century AD) was investigated in this study.

**Alba Fucens** (contact: Eugenio Cerilli, Emanuela Ceccaroni). Alba Fucens (Massa d'Albe, L'Aquila), a Roman colony under Latin law, was founded in 303 BCE in the territory of the Equi. Located in an elevated position (ca. 1,000 m above sea level) north of Lake Fucino, it occupied a strategic position at the foot of Mount Velino. During the Gothic War (535-553 AD), Alba Fucens was occupied by a garrison of the Byzantine army. Several excavation campaigns have been carried out between 2006 and 2011 in the court of the Sanctuary of Hercules. In particular, in 2011 a large circular cistern was identified. The cistern (4 m. diameter, 6 m. depth) was filled with the voluntary accumulation of architectural elements, mixed to a large quantity of pottery, glass and marble fragment as well as faunal elements. The materials recovered mainly come from the destruction of the Sanctuary of Hercules that occurred between the end of the 5th and the beginning of the 6th century AD. after a seismic event. The faunal assemblage is composed for the most part of domestic animals and minimally of wild taxa, an indication that hunting was an activity extremely secondary and occasional. Among the remains of wild animals, the majority belong to red deer (*Cervus elaphus*) followed, with very few elements, by the wild boar (*Sus scrofa*) and the chamois (*Rupicapra sp*.) (*67*). Remains of a left pelvis of a cat found in the cistern were genetically analyzed in this study and radiocarbon dated to 1,783±24BP (RICH-29170) corresponding to 210-340 cal. AD (95.4%).

**Arene Candide** (contact: Francesca Alhaique, Antonio Tagliacozzo). Arene Candide is a large cave located at 90 m above current sea level on the slopes of Monte Caprazoppa (Finale Ligure, Savona, northern Italy). The site has been renowned since the second half of the 19th century AD for the archaeological richness of its Holocene and Pleistocene layers. Several excavations occurred during 1940-42, 1948-50, 1970-72 and 2008-2011. Human occupation is testified since the Upper Paleolithic, which is divided in two groups of strata divided by a gap. The first is dated from about 25,000 BP to 18,000 BP (layers P). The second is dated to 11,000-10,000 BP (layers M). All the materials were sieved during the excavations resulting in an accurate collection of the bones, as indicated also by the presence in the sample of abundant microfauna. In the period corresponding to the P layers both humans and carnivores used the cave as a shelter: in the lower strata, the high carnivore/ungulate ratio, together with the almost complete absence of cut marks and other traces of human activity, point to an almost exclusive carnivore frequentation, while in the upper layers, also humans contributed to the accumulation of the bones. In the M layers, humans are instead the main agent of bone collection and almost all the species, including carnivores, display traces of human activity (*68*). Two wildcats originating from the M layers (*69*) and therefore stratigraphically dated to 10,000-11,000 BP were analyzed in this study, one of which resulted in genome-wide data.

**Galgenbühel/Dos de la Forca** (contact: Ursula Wierer). The Early Mesolithic rock shelter Galgenbühel/Dos de la Forca is located at Salurn/Salorno, in the Province of Bozen/Bolzano (Italy), in the Eastern Alps. Excavation campaigns were conducted between 1999 and 2002. Radiocarbon dates indicate a repeated human frequentation of the shelter approximately between 8,500 BCE and 7,500 BCE by Sauveterrian groups whose economy was centered on the exploitation of nearby wetlands and the forested valley bottom. Faunal species ecology indicates a habitat of slack and slow flowing waters bordered by shore and submerged vegetation and surrounded by forest (*70*). The abundant fish assemblage is dominated by pike (suggesting specialization in pike fishing) and by several species of the Cyprinidae, including the rudd and the roach. Seasonality data indicate that fishing was carried out during spring and summer (*71*, *72*). The collection of freshwater molluscs (*Unio*) and European pond turtles is attested. Mammals are dominated by beaver, wild boar and red deer. About 9% of the bones attributable to nearly all species indicate human manipulation of the carcasses through the presence of cut marks, including small mammals (Crezzini et al., in press). Among the carnivores, represented by a fair amount of bones, the wildcat is the best represented species (83 elements from at least 9 individuals). The analysis of cut-mark distribution in the wildcat assemblage of Galgenbühel/Dos de la Forca showed that they are only partly related to skinning. The localisation and the features of some marks attest disarticulation and therefore support the use of *F. silvestris* as food (*34*). Two wildcats were analyzed in this study, one of which radiocarbon dated to 8,899±35BP (RICH-34534), corresponding to 8,240-7,940 cal. BCE (95.4%).

**Genoni Santu Antine (Sardinia)** (contact: Barbara Wilkens). The Nuragic well of Santu Antine in Genoni, located in the central-western part of Sardinia near the Giara plateau, has a depth of approximately 40 meters and contains a rich assemblage of artifacts spanning multiple historical periods. Initially constructed during the Nuragic era (circa 1900–730 BCE), the well served both ritualistic and practical functions. Excavations conducted by the Superintendence for Archaeological Heritage of Sassari and Nuoro between the 1980s and early 1990s unearthed significant artifacts from the Nuragic, Punic, and Roman periods, illustrating its prolonged use (*73*). Additionally, a faunal assemblage of 377 specimens was collected from the well, with bone remains found in the upper layers of the sediment. Key finds from these excavations include two bronze figurines: one possibly depicting a deity in ritual nudity and another with a raised hand, suggesting a gesture of blessing or greeting. These items were found alongside Roman coins, bronze and lead vessels, and an ivory dagger handle, indicating local craftsmanship and extensive trade networks. Additionally, evidence of votive offerings and bronze pulleys, likely related to water collection, supports the well's dual role as both a functional and religious site, remaining active into the Roman period, particularly until about the 2nd century CE.
In this study, remains from two cats were analyzed, represented by two femurs from an adult and a right tibia from a juvenile, likely less than a year old. The elements exhibited relatively large dimensions. The adult sample underwent radiocarbon dating, yielding 2,121 ± 25 BP (RICH-34524), corresponding to 340–320 cal. BCE (3.9%) / 200–50 cal. BCE (91.5%).

**Graffignano** (contact: Francesca Alhaique, Giuseppe Romagnoli). Two underground refuse dumps, previously used as cisterns or silos but also containing animal remains, have been discovered within the Baglioni-Santacroce castle in Graffignano (Viterbo, northern Latium, Central Italy). The two pits are dated respectively to the first half of the 15th century AD and the second half of the 15th and the beginning of the 16th century AD based on typology and decoration of ceramics. Remains of the cat analyzed in this study were found in the most recent pit. These garbage pits had been used not only for discarding food debris, but also for disposing of pets (including cat) and pests, or weeds, together with other materials (*74*).

**Grotta del Cocci** (contact: Antonio Tagliacozzo, Leonardo Salari). Grotta dei Cocci is located on the right bank of the Nera River, facing the historic center of Narni (Terni, Umbria, Central Italy). The cave was excavated between 1989 and 2000. An articulated stratigraphic sequence was identified in which two principal moments of human frequentation are recognizable: the oldest dated to the last few centuries of the 6th millennium BCE, the most recent dated to the Early and Middle Bronze Age. More than 2000 faunal remains were unearthed from the Neolithic layers, the majority of which is represented by small mammals (*75*). The wildcat is represented by a phalanx and a femur attributed to a single individual, which was analyzed in this study. The sample was radiocarbon dated to 6,324±31BP (RICH-34517) corresponding to 5,370-5,210 cal. BCE (95.4%).

**Musarna** (contact: Antonio Tagliacozzo, Beatriz Pino Uria). Discovered in the middle of the 19th century AD, the Etrusco-Roman city of Musarna, located between Viterbo and Tuscania (Latium), was founded at the end of the 4th century BCE as a military colony, and abandoned at the beginning of the 7th century AD (*76*). Excavations were conducted between 1983 and 2003, and in 2011. Over 2,500 faunal remains originating from the public baths in contexts dated from the 3rd century BCE and the 1st century AD were analyzed, mostly belonging to domestic animals. Pig, sheep, goat, and rarely bovine remains were found in cisterns used for waste disposal after the abandonment of the baths (*77*). The cat sample analyzed in this study was radiocarbon dated to 1,644±25BP (RICH-34516) corresponding to 360-540 cal. AD (95.4%).

**Nardò "torre santa Caterina" (Sardinia)** (contact: Claudia Minniti, Jacopo De Grossi Mazzorin). The sea watchtower of Santa Caterina is located on the Ionian coast, at Nardò, about 10 km north of Gallipoli. It was probably built at the end of the 16th century AD. Its ground floor was likely used as animal shelter. The archaeological finds collected at the site have been dated from the 18th to the first half of the 19th century AD. Faunal remains testified to reliance of the inhabitants of the tower on sheep and goat husbandry and the exploitation of the nearby seacoast (fish remains and seashells). Few remains attributed to two cats (analyzed in this study) and a dog were found, probably used as companion animals. Among the wild species only small mammals were found, such as fox, hare and hedgehog (*78*).

**Nuraghe Flumenelongu (Sardinia)** (contact: Barbara Wilkens). The recent and Final Bronze/Early Iron Age (Nuragic Age) is a very important period for Sardinia, which sees the introduction of numerous domestic and wild species. Phoenician colonies were established in some coastal localities. In association with Phoenician, and later Punic, colonization or influence, there is an increase in the remains of deer, used for feeding and for the processing of antlers. Nuragic stratigraphic units at Nuraghe Flumenelongu see a clear predominance of sheeps and goats (80%) over pigs (9.62%). The cattle are scarcely present (*79*). The largest number of bone findings comes from stratigraphic units attributed to the Roman Imperial period. The numerically most represented animals are goats (35%) followed by pigs 23% and cattle 12%. Eight bone fragments belong to the dog and only one to the cat. Sheep are more numerous than goats. The pig is the typical meat animal, slaughtered at a young age. The bovine remains also present a poor state of preservation. The identification of anatomical parts belonging to individuals of all age stages suggests that these animals were not used exclusively for plowing or heavy labor. Given the high proportion of deer, hunting activity must have played an important role at this site. A cat was analyzed in this study and radiocarbon dated to 1,365±25BP (RICH-34518) corresponding to 600-620 cal. AD (2.7%) / 630-690 cal. AD (86.2%) / 740-780 cal. AD (6.6%).

**Padua Via Cesare Battisti** (contact: Antonio Tagliacozzo, Ivana Fiore). From January 25 to April 16, 1993, an archaeological excavation was carried out at Via Cesare no. 132, Padua. The street, a connecting section between the area of the hospitals and the Riviera dei Ponti Romani, has a SE-NW course. The excavation area is located approximately halfway along its course, not far from the intersection with Via Santa Sofia. Since the 8th century BCE, the development of the proto-urban settlement of Padua has been contained within the first large bend of the river Brenta (ancient Meduacus). Only sporadic settlements outside to the east as far as Via Santa Sofia are reported. Human occupation is observed since the 7th-6th century BCE (phase 0) until the 3rd century BCE (phase 1), abandonment of the site in association with an alluvial layer). The area was then reclaimed with drainage trenches between the 3rd and 1st centuries BCE (phase 2), with occupation during the Roman era until the 2nd century AD (phases 3-4) (*80*). The area was then abandoned again until reedification in the Medieval era. A cat from this latter period radiocarbon dated to 515±23BP (RICH-34527) corresponding to 1,400-1,440 cal. AD (95.4%) was analyzed in this study.

**Palatino Northeast slope (Rome)** (contact: Gabriele Soranna). The faunal assemblage, currently under study, was excavated on the NE slope of the Palatine Hill by the University of Rome La Sapienza since 2001. It relies on a well-defined stratigraphic chronology obtained from pottery and chronologically spans the Mid-Republic (4th-3rd c, BCE) to the Middle Ages (12th-13th AD). Frequency and ratio of identified taxa, in particular those of the domestic species mainly exploited for meat, show a framework fitting well into the urban context of Rome. Wild taxa appear marginal as well as birds and fish. Domestic species usually non-consumed in the diet such as dogs, cats and equids appeared rarely in the faunal record. Five bone fragments have been identified as domestic cat in contexts referred to the 4th-6th century AD, corresponding to four individuals, including one juvenile (*81*). Three specimens were analyzed in this study. For two of them we obtained genome-wide data. One cat was radiocarbon dated to 1,764±24BP (RICH-35364), corresponding to 230-370 cal. AD (95.4%).

**Partanna C. da Capo d'acqua (Sicily)** (contact: Roberto Micchichè). The specimen, represented by an almost complete skull without a mandible belonging to the species *Felis silvestris*, was discovered in the area of Contrada Stretto in Partanna (Trapani) (Long. 12°54'57.94"E – Lat. 37°43'28.43"N). The skull was found inside a large bell-shaped vessel (*36*), which constituted the pivotal element of a specific prehistoric cave context interpreted by archaeologists as significant for water cult rituals (*82*). The dating of this context is based on the discovery, within the same vessel containing the cat remains, of a dipper cup attributable to the Castelluccio facies, thereby linking it to the Early Bronze Age (*83*). Recent studies based on radiometric analyses (*84*) have provided a more precise chronological placement for the Sicilian Early Bronze Age, proposing a date range between 2,200 and 1,500 cal. BCE.

**S. Cecilia in Trastevere (Rome)** (contact: Claudia Minniti, Jacopo De Grossi Mazzorin). The faunal assemblage from the church of S. Cecilia in Trastevere originate from the Medieval layers of baptistery, between the end of the 12th and the beginning of the 13th century AD. The majority of the faunal remains identified (about 96%) belong to typical domestic animals (bovines, sheep, goat, pig and chicken). Wild animals are mostly represented by deer, wild boar, marten, hedgehog, hare and various bird species. Other domestic species observed, such as horse, donkey, dog and cat, were probably not used as food source (lack of butchery marks). Fourteen bone elements belonging to at least five cats were found, with no evidence of skinning or butchery (*85*). An adult cat was analyzed in this study.

**Saracena (Cosenza, Calabria)** (contact: Antonio Tagliacozzo, Vincenzo Tinè). The cave of San Michele lies on the right side of the narrow Garga valley, facing the modern settlement of Saracena (Cosenza, Southern Italy) at an altitude of 750 m a.s.l. Research conducted from 1998 to 2009 revealed an uninterrupted stratigraphic sequence from the Neolithic period to the Bronze Age (*86*). The faunal assemblage originates mostly from combustion pits from the Neolithic and Eneolithic layers. Domestic species are dominant (sheep and goat in particular), and, among the rare wild species, wild boar, deer and small mammals were found (hare, marten and wildcat). Remains attributed to two wildcats, one of which analyzed in this study, were found. It remains unclear whether they were used as food sources (*87*).

**Tertenia, Sardinia** (contact: Barbara Wilkens). Excavations conducted at Fusti 'e Carca, in Tertenia (Sardinia, Italy) showed evidence of human frequentation from the beginning of the 4th century BCE until the mid 6th century AD. Wall structures unearthed at the site are mostly attributable to the 4th-6th century AD. The faunal assemblage is chronologically dated to this later period and originates from a well in the area XXVII. A total of 6,028 bone fragments were found, of which 2813 identified and attributed to food waste. The remains of a cat, analyzed in this study, indicated the age of more than 11 months. The evidence of healed traumatic fractures in both the femurs testify to its domestic status and close relationship with humans (*88*).

##### Austria

**Bernhardsthal** (contact: Konstantina Saliari). This Germanic settlement, a farmstead, was located about 60 km north of the Danube that formed the Roman limes in this region. The site was occupied from the first half of the 2nd century AD until the mid-3rd century AD by the Marcomanni, a Germanic tribe that established a kingdom during the peak of power of the nearby Roman Empire. In total, 14,783 faunal remains have been identified and further analyzed from the settlement of Bernhardsthal (*89*, *90*). The composition of the faunal material indicates a peasant economy based on the domesticated animals, which dominated with almost 95%. According to the archaeozoological analysis, cattle were the prevalent domesticated taxon, comprising 54% of the total material, based on NISP data. Among the various species that were documented in Bernhardsthal, two partial skeletons, one of a juvenile and one of an adult cat were found. The adult animal was identified as a wildcat on the basis of its measurements. Of the juvenile cat, a tibia was analyzed and radiocarbon dated to 1,165±26BP (RICH-34515), corresponding to 770-980 cal. AD (95.4%). Of the adult animal, a radius was taken for analysis.

**Mautern an der Donau - Eastern Vicus** (contact: Günther Karl Kunst). The Roman auxiliary fort of Mautern and its associated *vicus* are located in northeastern Austria, province of Lower Austria. Mautern/Favianis saw a continuous Roman presence from the 1st until the 5th century

AD as one of the main military and civilian sites along the Danubian limes in the province of Noricum. An auxiliary fort was established in the early imperial period, and a civilian settlement, the *vicus*, developed around it from the 1st century onwards. The site gained regional importance in the Late Roman period (5th century AD), when it was converted into a fortified settlement. Among the numerous faunal remains studied from the eastern vicus, the presence of cat was attested in only one context, the filling of a pit-house (Grubenhütte) that was dated to 100-140 AD (*43*). The bone material was mainly composed of butchery refuse but with the presence of partial skeletons of a juvenile domestic pig, a small dog and a subadult cat. The cat skeleton, which was almost completely preserved, was identified as a wildcat on the basis of its relatively large size. The mandible that was analyzed in the present study was radiocarbon dated to 2,007±25BP (RICH-34526) corresponding to 50 cal. BCE-80 cal. AD (95.4%) which is slightly older than the assumed archaeological date.

**Petronell-Carnuntum** (contact: Günther Karl Kunst). Ancient Carnuntum is located about 40 km east of Vienna, along the southern shore of the Danube. The civilian town of Carnuntum was located directly on the Pannonian *limes* and developed parallel to the legionary camp that was about 2 km further east. This town developed since the 1st century AD and became the administrative centre of the Roman province *Pannonia superior* from the beginning of the 2nd century AD. It was important as a crossroad for two transcontinental main trade and transport routes. By the mid-5th century AD, the town was deserted. The cat remains analyzed in the present study are isolated finds from two different contexts. One of the bones was a femur (PETRcat06) retrieved from the latest walking horizon of the western street (so-called *Weststraße*) of *insula VI* (*91*). It has been radiocarbon dated 1,830±26BP (RICH-34523), corresponding to 120-260 cal. AD (87.6%) / 290-320 cal. AD (7.8%). From the same insula, a cat mandible (PETRcat08) was found inside the baths (*92*). This specimen was radiocarbon dated 1,919±24BP (RICH-35370), corresponding to 20-50 cal. AD (2.6%) / 60-210 cal. AD (92.8%).

**Salzburg - Residenz, Wirtshaus Schinagl** (contact: Konstantina Saliari). Since the 12th century AD, the prince-archbishops of Salzburg had lived in Salzburg's historic centre. In the early 17th century AD, a number of houses next to the bishop's palace were demolished to make way for an additional building added to the palace. During excavations in the 1980s, a cesspit was found in the courtyard of one of these houses that served as a tavern in the last phase of its occupation. The archaeological material (ceramics, glass, etc.) made it possible to date the filling of this pit to the 16th century AD. In total, 11,127 faunal remains have been identified and further analyzed (*93*). The faunal material represents mainly kitchen and table refuse from the tavern. Although mammals prevailed with 87.7% (and especially cattle with 47%, NISP data), a big variety of domesticated and wild birds as well as fish were found there. The presence of exotic species (e.g. salt-water fish) indicates suppliers from distant areas and a demand for high quality food. In addition to the kitchen refuse, the partial skeletons of a dwarf dog and three cats were found. Besides two young cats, there was an adult individual that was believed to possibly be from an angora cat. This hypothesis was based on the large size and stocky build of the skeletal elements compared to those of medieval Haithabu and modern cat breeds. The possible presence of a cat with Persian origins was suggested to be linked to the nearby presence of the bishop’s palace from where the cat may have come. The cesspit also yielded evidence for the presence of another special breed, i.e. a crested chicken that was sporadically depicted in western Europe since the 15th-16th century AD.

**Traismauer** (contact: Konstantina Saliari). The civil settlement (vicus) Traismauer/Augustiana, about 50 km west of Vienna, was located on Roman territory, just south of the Danube, which formed the limes in Roman times. The faunal assemblage consists of 12,792 identified faunal remains that are dated between the 1st to the 4th centuries AD (*44*). Domesticated species dominated in Traismauer with more than 95%; the most important taxon for the economy was cattle, which prevailed with 69.4% (NISP data). Among the faunal remains, a total of 17 cat bones were found, belonging to five individuals. They show a relatively large variation in size and were believed to represent both wild and domestic animals. The mandible analyzed in the present study was considered domestic. It was radiocarbon dated 1,897±27BP (RICH-34530), corresponding to 60-220 cal. AD (95.4%).

#### Germany

**Bremen, Marktplatz & Stadtgraben, Am Wall** (contact: Hans Christian Küchelmann). Bremen is located along the River Weser at about 60 km of the North Sea. It became a bishop's seat in 782 AD and, gained substantial political and economic influence as merchant town in medieval times. In 1260 AD Bremen became part of the Hanseatic League. During excavations carried out in 2002 on the Marktplatz (market place) of the old town center, over 5700 faunal remains were collected from contexts dating to medieval and postmedieval times, six of which were cat bones (*94*). A cat femur was found in the cellar of a house (Baleersches Haus) that was dated between the 16th and 18th century AD on the basis of ceramics, glass and coinage. A radiocarbon dating was carried out showing that the cat specimen needs to be placed between the mid-15th to mid-17th century AD: 381±24BP (RICH-34505) which corresponds to 1,440-1,530 cal. AD (65.0%) / 1570-1630 cal. AD (30.4%). The second bone sampled, a juvenile right femur, was found in the medieval moat of the town, which was filled during the enlargement of the town's defenses. More than 32,500 animal bones were recovered from the ditch, among these 86 cat bones. According to accompanying archaeological finds and dendrochronological data, the context is dated to the 2nd half of the 16th century AD (*95*).

**Haithabu, settlement area** (contact: Ulrich Schmölcke). The site is located in the municipality of Busdorf near Schleswig, at the mouth of the river Schlei that empties into the Baltic Sea. During the Viking Age, from the 9th to the mid-11th century AD, Haithabu was an important trading centre and a main hub for long-distance trade between Scandinavia, Western Europe, the North Sea region and the Baltic. The extensive excavations of Haithabu yielded no less than 220,000 identifiable mammal bones of which 1030 are from cat (*96*). With an MNI of 129, based on the tibiae, this is the largest available collection of archaeological cat bones known thus far. All bones, except two, were identified as domestic on the basis of the relatively small size of the remains. In the present study, three mandibles of adult animals from the settlement area were analyzed.

**Lauchheim-Mittelhofen** (contact: Simon Trixl). The site of Lauchheim-Mittelhofen is located about 1 km west from the small town of Lauchheim in Baden-Württemberg. Besides the Merovingian cemetery of Lauchheim-Wasserfurche, there is also the nearby site of Lauchheim-Mittelhofen with both an Early to High Medieval settlement as well as some burials directly associated with those farmsteads. In the present study, a bone was analyzed of a juvenile and of an adult animal coming from the Mittelhofen settlement. Both remains are from the same find context, i.e. the filling of a pit house dating to the local "Siedlungsstrukturphase II" (settlement phase II), covering the time span between ca. 825 AD and the 10th century AD (*97*).

**Spain**

Cova de Els Trocs, Huesca (contact: Marta Moreno). The cave of Els Trocs, located at more than 1500 m asl on the Central Iberian Pyrenees, was seasonally occupied by herders throughout the Neolithic (*98*). The fauna consists mainly of domestic animals, and in particular sheep, whereas wild taxa are rare. In the lowest level of the cave deposit occurred a single cat bone, a complete ulna, that was analyzed here. The specimen was radiocarbon dated to 6,207±32BP (RICH-34519), corresponding to 5,300-5,250 cal. BCE (9.4%) / 5,230-5,040 cal. BCE (86.0%).

**Castillo de Labastida** (contact: Idoia Grau-Sologestoa). Labastida is a town in the province of Álava (Basque Country, northern Spain), located between the River Ebro and the Sierra de Toloño mountain range. The construction of the castle of Labastida started in the 10th-11th centuries AD on the top of the hill dominating the town (*99*). The cat bone analyzed here, a metatarsus, was found in a big levelling/construction package that is dated to the abandonment of the castle, at the end of the 15th or perhaps the very early 16th century AD.

**Terrera Ventura, Tabernas** (contact: Joris Peters, Bea De Cupere). The site of Terrera Ventura is in Andalusia, in Tabernas, about 30 km north of Almería. This Chalcolithic settlement was inhabited for about a millennium and subsistence was mainly based on domestic animals, supplemented by hunting mainly red deer and hare (*100*). Two cat bones, belonging to the same individual, were found in a layer belonging to Phase III, which was dated between 2,700 and 2,400 BCE.

**Bulgaria**

**Durankulak** (contact: Joris Peters, Bea De Cupere). The site of Durankulak is located in northeastern Bulgaria, close to the Romanian border. When it was settled in Neolithic times, it was still on an island of the freshwater lake of Durankulak that was separated from the Black Sea by sand dunes and a beach strip. The area was inhabited intermittently until the Middle Ages. Among the faunal remains studied from the Chalcolithic levels at Durankulak are 7 bones of 3 individuals and two bones from Late Bronze Age contexts (*101*). The mandible of an adult animal (DURcat01) that was sampled for the present study came from the Late Aeneolithic horizon V and its Chalcolithic date was confirmed by the radiocarbon date of 5,535±32BP (RICH-34522), corresponding to 4,450-4,330 cal. BCE (95.4%). The second individual analyzed here is also represented by a mandible (DURcat04), depicted in Manhart (1998; Fig. 82) and labeled as Chalcolithic. However, it appears to be late medieval after radiocarbon dating: 414±33BP (RICH-21622), corresponding to 1,420-1,530 cal. AD (85.4%) / 1,570-1,630 cal. AD (10.0%).

**Koprivec** (contact: Joris Peters, Bea De Cupere). This inland site is located in northern Bulgaria. It lies at the confluence of the three headwaters of the Lom, which flows into the Danube at Ruse, 60 kilometers to the north. Excavations carried out in the 1990s yielded material dating between the early Neolithic and the Chalcolithic. Eight cat bones from two individuals were found among the faunal remains, consisting mainly of slaughter and consumption waste from domestic and hunted animals. All were found in levels dated as Neolithic B (*101*). In this study, we analyzed an ulna (KOPcat03) and a pelvis (KOPcat04), both from an adult animal. The respective radiocarbon dates are 7,100±34BP (RICH-21733) or 6,050-5,900 cal. BCE (95.4%), and 6,452±31BP (RICH-34528) or 5,480-5,360 cal. BCE (95.4%).

#### Portugal

**Bank of Portugal (contact: Cleia Detry).** Excavations in Lisbon, Portugal, were directed by Artur Rocha and took place at the former headquarters of the Bank of Portugal to create the Money Museum, which now also features a small archaeology exhibition. Located in today's downtown Lisbon, the site revealed a stratigraphic sequence starting from Roman times and continuing to the present day.

During the Roman period, the site was submerged but close to the shore, situated far from the city center, which was concentrated around the castle area a few kilometers away. This location was in the lower part near the Tagus River. By medieval times, the area had become dry land, and a 40-meter section of a defensive wall was built by King D. Dinis towards the end of the 13th century AD. This wall was later destroyed and buried by the great earthquake of 1755 AD. Today, 40 meters of this wall are on display at the Money Museum. In the more recent layers, around 300 human skeletons were discovered beneath the current location of the Church of São Julião, which served as a necropolis during the 19th century before being adapted as the headquarters of the Bank of Portugal. Zooarchaeological remains from the site are being studied by Ana Beatriz Santos for the Roman levels (Santos, PhD thesis, in preparation) and by C. Detry (in preparation) for the medieval and modern levels. So far, the results have revealed approximately 8,000 bones and teeth, and almost 2,000 shells. Of these, 143 remains were identified as belonging to cats.

We analyzed long bones from three adult cats, all were radiocarbon dated and gave Medieval dates: the first resulted in 1,084±23BP (RICH-35365) corresponding to 890-930 cal. AD (32.4%) / 940-1,020 cal. AD (63.0%); the second dated to 1,049±23BP (RICH-35366) corresponding to 900-920 cal. AD (4.6%) / 970-1040 cal. AD (90.8%); the third to 1,059±23BP (RICH-35367) corresponding to 890-920 cal. AD (11.0%) / 950-1030 cal. AD (84.4%).

**Palmela Castle** (contact: Cleia Detry). The castle of Palmela was built at an altitude of 250 m a. s. l., in a small town of the same name in the district of Setúbal (south-western Portugal). It is located on the Setúbal peninsula between the Tagus River and the mouth of the Sado River. In the 8th/9th centuries AD, construction started as a primitive fortification under Muslim rule. Afterwards, several building phases occurred, and the castle was occupied until modern times. The excavations occurred during the 1990´s and directed by Isabel Cristina Fernandes. In the course of a recent archaeozoological study conducted on more than 10,000 identified vertebrate remains from the different occupation phases, a total of 32 cat remains were found corresponding to a minimum of seven individuals (*102*). In the present study we analyzed a tibia of an adult cat from a 11th-12th century AD context (Almoravid period) and a humerus of a juvenile animal from a context dated to the 12th-13th century AD (transition Almohad period/1st Portuguese phase).

#### Belgium

**Parking 58, Brussels** (contact: Bea De Cupere). This site is located in the historical centre of Brussels between three main squares (Places de la Bourse, de Brouckère and Sainte-Catherine) (*103*). The excavations carried out in 2019 documented several ancient beds of the River Zenne between the 8th and 12th centuries AD, which yielded relatively small amounts of archaeological material. From the second half of the 13th century, the course of the Senne appears to have been altered by river regulation, leading to more debris being deposited. In the late 14th or early 15th century a quay was built on the right bank. Numerous faunal remains were found in the river bed, dating to the end of the 14th-15th century AD and mainly consisting of butchery and consumption refuse as well as of skeletons of animal carcasses thrown in the river. Two cat mandibles, dating to the late medieval period, have been retained for analysis here.

**Tongeren, Industrie Oost** (contact: Bea De Cupere). In the 1st century BCE, a Roman military encampment was installed in Tongeren, which developed into a civilian settlement a few decades later. This ultimately became the only Roman city in Flanders (northern Belgium). Archaeological traces are ubiquitous in today's city and are frequently found during infrastructure work. In 2021, evidence for late Iron Age and Roman occupation was found on the eastern edge of town (*104*). Among the Roman traces was a well that must have been associated with one of the poorly preserved buildings nearby. In the upper part of the shaft, below a layer of construction debris, numerous faunal remains were found. Besides a small number of bones representing human food refuse, several skeletons were found of animals that were not consumed. There were 8 dogs, several of which were still in anatomical connection, a partial cow skeleton and a more or less complete cat. The animal was 11-12 months of age when it died. The radiocarbon date of 1,953±27BP (RICH-34531) obtained on one of the bones corresponds to 40-10 cal. BCE (4.0%) / 10BC-130 cal. AD (91.4%).

##### France

**Ecly** (contact: Tarek Oueslati). The site of Écly – Les Septiers is located in the Ardennes department in northern France. It was a rural settlement occupied in the 2nd and 3rd century AD. Architectural traces are rare and poorly preserved, and the majority of the archaeological finds come from the lower part of a 9 m deep shaft. Besides consumption refuse consisting of the traditional domestic food animals, the shaft yielded remains of at least 10 fallow deer, suggesting that the villa was occupied by a well-to-do landowner that possessed a game park. A relatively high social status is also suggested by the presence of peacock and of two dwarf chickens. Wild species include hare, fox and several birds including raven. Cat is represented by the skeleton of an immature animal that died between 8,5 and 10 months of age (*105*). The wild or domestic status of the individual could not be established on an osteological basis. The cat was radiocarbon dated to 1,915±25BP (RICH-34507) corresponding to 20-50 cal. AD (2.1%) / 60-210 cal. AD (93.3%).

##### Serbia

**Viminacium Amphitheatre, Nad Klepačkom** (contact: Sonja Vuković). Viminacium is located on the right bank of the river Mlava, close to its confluence with the Danube, near the present-day town of Kostolac in eastern Serbia. Originally, in the 1st century AD, it was a legionary camp built to defend the northern border of the Roman Empire. The town that arose next to the camp, became the capital of the province of Moesia Superior and later Moesia Prima. It was a very prosperous town especially in the 2nd and 3rd centuries AD. Excavations in and around the amphitheatre of Viminacium, located at the north-eastern corner of the town, yielded more than 20,000 identifiable faunal remains (*45*). For the present genetic study, two cat bones that we also radiocarbon dated were retained. A humerus (VIAcat06) was dated to 1,923±25BP (RICH-34512), corresponding to 20-210 cal. AD (95.4%), whereas a mandible (VIAcat04) was dated to 1,697±25BP (RICH-34521), corresponding to 250-280 cal. AD (16.8%) / 330-420 cal. AD (78.6%). We also analyzed a cat bone derived from the nearby site of Nad Klepečkom, located at about 3km to the east of Viminacium. Excavations yielded evidence for occupation during Chalcolithic, Roman and medieval times. The radiocarbon date obtained on the bone from Nad Klepečkom showed that it is medieval: 992±25BP (RICH-34510) corresponding to 990-1050 cal. AD (49.5%) / 1080-1160 cal. AD (45.9%).

#### Ireland

**Glencurran Cave** (contact: Marion Dowd). This cave is located in the Burren, near the village of Kilnaboy. Excavations documented funerary and ritual practices from the Middle and Late Bronze Age, in addition to early medieval habitation (*106*). Deep in the cave, the partial skeleton of a wildcat was found which is assumed to have died naturally. This deposit seems to predate human activity in the cave (McCarthy, M. unpublished. Unpublished report on the faunal remains from Glencurran Cave). A canine was radiocarbon dated to 4778±24 BP (UBA-45133), corresponding to 3634-3526 cal. BCE (95.4%), indicating that the animal was Neolithic in age. For the genetic analysis, a mandible fragment was used.

#### Greece

**Kassope** (contact: Cornelia Becker). Kassope is located in Epirus, on a hill overlooking the Gulf of Ambracia, around 25 km west of Ambracia (present-day Arta) and 10 km from the Ionian coast. This ancient town was founded in the mid-4th century BCE and was occupied until the end of the 1st century BCE. More than 28,000 animal remains were identified from this site (*107*). A mandible and a femur of two adult cats, both identified as wild on the basis of size by the archaeozoologist, were analyzed. One cat was radiocarbon dated to 2,253±25BP (RICH-34520) corresponding to 400-340 cal. BCE (33.6%) / 310-200 cal. BCE (61.8%).

### Materials and Methods

#### Radiocarbon dating

37 samples from 30 archaeological sites (Table S6), were submitted for radiocarbon dating at the KIK-IRPA AMS laboratory, Royal Institute for Cultural Heritage, Brussels (*108*). Samples were prepared following pretreatment protocols described in (*109*). Dates were calibrated using OxCal 4.4 and the IntCal20 calibration curve (*110*).

#### Sample preparation and DNA extraction

Pre-PCR laboratory analyses of the archaeological samples were performed in the dedicated ancient DNA (aDNA) facility of the Center of Molecular Anthropology for Ancient DNA Studies of the University of Rome Tor Vergata, Villa Mondragone, Monte Porzio Catone (Rome, Italy). The Centre features state-of-the-art laboratories for aDNA consisting of individual 'cleanroom quality' working spaces (ancient sample pre-treatment, milling, DNA extraction, genomic library and PCR set-up fully equipped rooms).

The workflows adopted followed the standard precautions for access to the facilities and decontamination described in the literature (*111*). In particular, access to the pre-PCR laboratory was restricted to a limited number of people and only after wearing clean overalls, two pairs of gloves, over-shoes, surgical facemasks, plastic spectacles. Access to the aDNA facilities was not permitted if amplified libraries had been handled the same day. The aDNA working spaces were routinely cleaned with bleach and RNAse Away (Molecular BioProducts, San Diego, CA, USA) and every item entering the room was extensively washed with bleach or RNAse Away and UV-irradiated.

Modern DNA and post-PCR laboratory analyses were conducted in physically separate buildings at the Department of Biology of the University of Rome Tor Vergata.

*Archaeological Samples*

To minimize contamination, samples were UV-irradiated for 15 minutes on each side using a crosslinker set to 254 nm. Approximately 1 mm of the outer surface of each bone element was removed with a drill (Proxxon), while teeth were wiped with a tissue moistened with a 2% bleach solution, followed by a rinse with ultrapure water. Fine bone powder (80-150 mg) was collected using either a drill or a mortar and pestle, then stored in UV-irradiated vials at 4°C until DNA extraction. After processing each sample, the drilling room was thoroughly cleaned with a 2% bleach solution and UV-irradiated for 15 minutes. Drill bits were decontaminated by cleaning with a 2% bleach solution, rinsed with ultrapure water and 80% ethanol, and UV-irradiated for 30 minutes.

To maximize the recovery of endogenous ancient DNA, a two-step decontamination extraction protocol was applied as previously described (*112*), with minor modifications. Briefly, bone powder was washed in 1 mL of 0.5% bleach solution in a rotating incubator for 15 minutes at room temperature. Following a 30-second centrifugation at 13,000 rpm, the supernatant was discarded, and the pellet was washed three times with 1 mL of ultrapure water. The second decontamination step involved incubating the pellet in 1 mL of 0.5M EDTA (pH 8) for 30 minutes at 37°C in a rotating incubator. After pelleting the powder, the supernatant was removed, and a digestion was performed by incubating the pellet in 1 mL of lysis buffer (0.5M EDTA, pH 8, and 0.25 mg/mL proteinase K) for 24-48 hours at 37°C in a rotatory wheel. Silica-based purification of the extracts was performed as described elsewhere (*113*), with a few modifications: 3M sodium acetate was added to the binding buffer preparation, and the extract (~1 mL) was combined with 12 mL of binding buffer using the High Pure Viral Nucleic Acid Large Volume Kit (Roche). Purified DNA was eluted in 62 µL of TET buffer (1M Tris-HCl pH 8.0, 0.5M EDTA, 10% Tween-20, ultrapure water) and stored at -20°C until genomic library preparation. Extractions were performed in batches of eight samples, with two negative controls placed in positions 5 and 10.

*Historic Samples*

Claws and skin samples of wildcats from Bulgaria (Table S4) from the collection of the National Museum of Natural History in Sofia were already sampled for a previous study (*13*). Extraction of DNA and library preparation of these samples were done in dedicated facilities for the genetic analysis of historic samples (dated to the 19th and 20th century CE) at the Department of Biology of the University of Rome Tor Vergata. A piece of tissue was collected for each specimens using a sterile surgical blade and weighed with a precision scale. DNA extraction was carried out using the Tissue and Hair extraction Kit and DNA IQ™ System (Promega), following the manufacturer's instructions, and including a negative control.

#### Library preparation

*Archaeological and Historic Samples*

Illumina double-stranded genomic libraries were prepared as described in the literature (*114*), with a few modifications: 20 µL of DNA input was used, adapter concentration was reduced to 0.15 µM, purification steps were performed using the MinElute PCR Purification Kit (Qiagen), and DNA was eluted into 1.5 mL DNA LoBind® Tubes (Eppendorf). The final cleanup step was replaced by enzyme thermal deactivation (20 minutes at 80°C). All PCR tubes were UV-irradiated for 15 minutes with lids open before use. A library negative control (20 µL ultrapure water) was included in each preparation batch. Finally, genomic libraries were set up for PCR amplification using a double indexing strategy (*115*). In an experimental phase of the project where single-stranded library protocols were tested, one sample from Belgium (BRPAcat24) was processed with the Santa Cruz protocol (*116*), along with other samples from other projects conducted in the lab. Special SCR adapters and splints were ordered from IDT as suggested in the reference protocol.

*Modern Samples*
Present-day samples were processed in the dedicated facilities at the Department of Biology of the University of Rome Tor Vergata. High-quality DNA extracted from skin tissues opportunistically collected from found dead animals between 1998 and 2010 from Italy (n=12) and North Africa (n=2) were selected from the Italian Institute for Environmental Protection and Research (ISPRA) *Felis* DNA biobank (*117*). Sampled individuals had been previously identified morphologically by collectors according to phenotype, life history traits, and biometric indices. Genomic libraries were constructed using the NEBNext® Ultra™ II FS DNA Library Prep Kit with Sample Purification Beads (New England Biolabs Inc.), following the manufacturer's instructions. DNA extracts were quantified using the Invitrogen™ Qubit™ 4 Fluorometer (Fisher Scientific) to ensure the appropriate adapter concentration. A library negative control (20 µL ultrapure water) was included in each batch. Indexing-PCR was set up using the NEBNext® Multiplex Oligos for Illumina®, supplied in a 96-well plate format with pre-combined forward (i7) and reverse (i5) primers, according to the manufacturer's instructions.

#### Sequencing

Genomic libraries (ancient, historical, and modern) were amplified in the post-PCR facility of the Department of Biology at the University of Rome Tor Vergata using a dual-indexing amplification strategy (*115*). Genomic libraries of archaeological and historic samples underwent 15 PCR cycles, while modern libraries underwent 4 to 8 cycles, depending on the initial extract concentration. PCR cleanup for ancient and historical samples was performed using AMPure XP magnetic beads (Beckman Coulter Inc.), following the manufacturer's instructions. Genomic library profiles were analyzed using the 2100 Bioanalyzer instrument (Agilent). Amplified libraries were diluted to 2 nM, pooled in batches of 30-40, and subjected to shallow shotgun sequencing on the Illumina NextSeq550 platform (75 SE, High Output kit) at the University of Rome Tor Vergata. Libraries with at least 10% endogenous DNA were selected for deeper sequencing on the Illumina HiSeqX or NovaSeq6000 (Macrogen Inc.), with few exceptions: the samples TERT03 (Tertenia, Sardinia, 8.4%), MENT01 (Menteşe, Turkey, 6.2%), and ASK01 (Aşıklı Höyük, Turkey, 5.9%) were deep sequenced given their chorological and geographic relevance. Differently, the sample SCAT03 (Torre Santa Caterina, Nardò, Italy), was not further processed because a sample from the site and chronology (SCAT01, 17-18th c. AD) was already deep-sequenced. The sample DUR06 (Durankulak, Bulgaria, 5th millennium BCE) was not further sequenced due to the lower degree of *post-mortem* damage compared with the other samples from the same site (~8% vs ~30-40%). Similarly, the samples BAY01-03 (Bayraklı, Turkey, 4th century BCE) were not further sequenced due to the low degree of *post-mortem* damage (~1-4%) detected, which suggested inconsistency with the stratigraphic evidence. To confirm that, the sample BAY01 was radiocarbon dated to 103.26±0.27 pMC (uncal BP).
Modern specimens were sequenced on the Illumina NovaSeq6000 platform (Macrogen Inc.) without prior screening. The number of reads generated are reported in Supplementary Table 2 and 3.

#### Raw Data Pre-Processing and Alignment to the Reference Genomes

*Modern samples*

Read quality was inspected using FastQC v0.11.9 (*118*). Quality filtering and adapter trimming of the reads was performed using AdapterRemoval v.2.3.3 with "--minlenghts 30", "--minquality 25", and "--trims" options (*119*). Separate alignments were generated for nuclear and mitochondrial genomes. Full-genomes were aligned against the latest domestic cat reference genomes F.catus_Fca126_mat1.0 (GCF_018350175.1), by using the *mem* algorithm in bwa v0.7.17 (*120*) with default parameters. Aligned reads were sorted and indexed using samtools v0.1.19 (*121*). We assigned read groups with GATK v3.8 (*122*). PCR duplicates were removed using dedup v0.12.8 (*123*) and local realignment around indels performed using GATK v3.8 (*122*). The same pipeline was utilized for aligning reads to the mitochondrial reference sequence of *Felis catus* NC_001700.1 with few differences since we used a pseudo-circular version of the mtDNA reference sequence as done for the ancient data (see next paragraph).

*Ancient and historic samples*

We inspected raw fastq files with FastQC and performed quality filtering and adapter trimming of the reads with AdapterRemoval using the same parameters as for the modern samples but adding the option "--collapse". Collapsed fastq files were then used to generate full-genome alignment following the procedure used for the modern samples with only one difference: we used the bwa *aln* algorithm, instead of bwa *mem,* with relaxed parameters ("-o 2", "-n 0.01") and seed disabled ("-l 1024") as typically done for ancient DNA (*124*). For the alignment to the mitochondrial reference sequence of *Felis catus* NC_001700.1, we prepared a pseudo-circular reference version using the tool CircularGenerator of CircularMapper v.1.93.5 (*125*). Reads were then aligned to the elongated mtDNA reference with bwa *aln* again with options "-o 2", "-n 0.01", "-l 1024", and then realigned to the original reference using the tool RealignSAMFile of CircularMapper. Finally, we assessed *post-mortem* damage patterns and performed rescaling of the bam files using mapDamage2.0 (*126*) for both nuclear and mitochondrial alignments.

#### Mitochondrial DNA consensus sequences generation

Previous work has found mtDNA insertions in the nuclear genome (NUMTs) of domestic cats (*127*) that can negatively impact mtDNA alignments and subsequent consensus sequence generation by introducing variants stemming from the nuclear inserts when mapping to the mtDNA. Recent analysis of wild and domestic cats from northern Europe showed that using high-coverage mitogenomes (>10X) generated from genomic libraries that present a high ratio between reads mapped against the mitochondrial and the nuclear genomes and a majority consensus call (*15*) does not result in incorporation of NUMTs variants in the mtDNA consensus sequence. We therefore computed the ratio between the reads aligned to the mtDNA and nuDNA genomes scaled by the respective genome lengths (Table S7). We decided to exclude three historic samples from Bulgaria due to the low coverage (<10X) and/or the low ratio between molecules aligned to mitochondrial and nuclear genomes scaled by the length of their reference sequences (ratio<35). One sample, ASK01 from Neolithic central eastern Anatolia, was retained given its relevance and since it was showing a high ratio between reads mapped against the mitochondrial and the nuclear genomes (>700) despite a mean depth of coverage of ~8X. All other samples showing a mean depth of coverage higher than 10X and mitochondrial versus nuclear aligned reads ratio from 40 to 2,300 were retained.

For each sample we first generated a raw VCF containing only variant sites using bcftools v.1.15 (*121*) (bcftools mpileup -BI -q 25 -Q 25 --annotate FORMAT/AD, FORMAT/ADF,

FORMAT/ADR, FORMAT/DP, FORMAT/SP, INFO/AD, INFO/ADF, INFO/ADR sample.bam -f reference.fasta | bcftools call -mv -Ov -o sampleX_raw.vcf). From the raw VCF we generated a raw consensus sequence using bcftools *consensus* (cat reference.fasta | bcftools consensus -H I sampleX_raw.vcf > sampleX.cns.fa). From the same raw VCF we also extracted, using bcftools *view* and *filter* functions, low-quality variants (QUAL < 29) and positions with multiallelic calls for which the majority allele has a frequency lower than 0.8. We use the function *vcf2bed* from BEDOPS v2.4.41 (*128*) to store these variants in a bed file that we used to mask the raw consensus sequence by setting such positions as "N", using the function *maskfasta* from bedtools v.2.30.0 (*129*). We finally computed coverage statistics using the tool *genomecov* of bedtools v.2.30.0 and extracted all positions covered by less than 5 reads. This list of low-covered positions was used for a second round of masking, setting again such positions as "N" in the final consensus sequence. We then used a custom script to concatenate all consensus sequences to use for phylogenetic analyses.

**Genome-wide SNP panels curation**

To date, a curated panel of Single Nucleotide Polymorphisms (SNPs) isn't available for the wild and domestic cat species complex. We therefore adopted the GATK v.4 short variant discovery pipeline (*130*) and then performed hard filtering following GATK best practices recommendations (*131*). We used modern >10-fold coverage genomes from this study (n=12) and from the literature (n=44). Briefly, we first called variants separately for each sample bam files using the tool HaplotypeCaller in GVCF mode (option --ERC GVCF); then per-sample variants were collected in the so called "GenomicsDB datastore" using the tool GenomicsDBImport; finally, we used the joint genotyping tool GenotypeGVCFs to generate the raw multisample (n=56) VCF. Hard filtering was performed using the GATK tool VariantFiltration to mark low-quality SNPs by read depth (--filter-expression "QD < 2.0" --filter-name "QD2"), poor mapping quality (--filter-expression "MQRankSum < -12.5" --filter-name "MQRankSum-12.5" --filter-expression "ReadPosRankSum < -8.0" --filter-name "ReadPosRankSum-8"), or strand bias (--filter-expression "SOR > 3.0" --filter-name "SOR3" --filter-expression "FS > 60.0" --filter-name "FS60"). Marked variants were then filtered out with GATK tool SelectVariants (--exclude-filtered). We further filtered the variants with vcftools v.0.1.16 (*132*) retaining only biallelic SNPs with minimum quality of 40 (QUAL>40) and for which we have genotype information for all individuals (--max-missing 1 --minQ 40 --remove-indels --min-alleles 2 --max-alleles 2 --recode --recode-INFO-all), resulting in 61,999,591 variants. We finally converted the filtered VCF to plink binary format using the software plink v1.9 (*133*).

For the ancient low-coverage genomes (n=70) as well as historic (n=3) and modern samples (n=2) up to middle-coverage (~8X), we called pseudo-haploid genotypes on the 61,999,591 positions obtained from modern genomes. Pseudo-haploid genomes were generated using *mpileup* function of samtools v.1.15 (*121*) and pileupCaller from sequenceTools (https://github.com/stschiff/sequenceTools) by selecting a single read randomly ('--randomHaploid' parameter) for each individual at each of the targeted SNP positions. We then merged in plink the pseudo-haploid panel with the modern reference panel (n=131). Finally, the merged panel was filtered in plink for minor allele frequencies (maf) and linkage disequilibrium (LD) (--maf 0.02 --indep-pairwise 50 5 0.2) resulting in 3,954,446 SNPs with a genotypic call rate of ~56%. From this combined and filtered dataset, we prepared three panels for downstream analyses. The first consists only of modern diploid genomes (n=56), thus excluding all ancient and pseudohaploid data (PANEL-A). To reduce computational burden of lengthy phylogenetic and ADMIXTURE analyses, PANEL-A was thinned in plink by imposing a minimum distance between SNPs of at least 5kb, resulting in a total of 403,868 SNPs (PANEL-B). The third panel included also the ancient genomes (n=131) thus, to limit potential biases from deaminated cytosines, we removed all transitions resulting in 919,415 transversions (PANEL-C).

#### Data analysis

*Nuclear phylogeny and population structure of present-day wild and domestic cats*

Phylogenetic analysis of a SNP-based supermatrix was conducted using RAxML-NG v.1.2.2 (*134*). The supermatrix was prepared by converting PANEL-B to VCF format in PLINK 1.9. The VCF was then converted to PHYLIP format using the python script vcf2phylip.py (*135*). A Maximum Likelihood tree was constructed under the GTR+GAMMA model with default parameters, using 500 bootstrap replicates and *Felis chaus* as an outgroup. The resulting tree was visualized using Figtree v1.4.4 (available at http://tree.bio.ed.ac.uk/software/figtree/) (Fig. 1B, Fig. S1). Population structure was first examined using Multidimensional Scaling (MDS) analysis of PANEL-A (Fig. S2). For this we applied the Classical Metric Multidimensional Scaling algorithm using cmdscale function in R v.4.2.0 (*136*), which operates on an Identity-By-State (IBS) pairwise distance matrix that we generated in plink (plink --bfile PANEL-A --distance square --out distance.matrix).

We further explored population structure with unsupervised ADMIXTURE analysis using PANEL-B. We used Admixture v.1.3 software (*137*) for K values between 2 and 13, with 200 bootstrap replicates and a 5-fold cross-validation (CV) procedure. The lowest CV values (Fig. S3a) were found for K=3 and K=4 (0.37868 and 0.37955, respectively). Starting from K=9 CV errors increased sharply possibly due to overfitting. Results were visualized with pong v.1.4.9 (*138*) (Fig. S3b). At K=3, one component was assigned to *F. l. lybica* wildcats and domestic cats (*F. catus*), a second one to *F. silvestris* wildcats, and a third one was shared by the Asian wildcat (*F. l. ornata*) and the Chinese wildcat (*F. bieti*), which clustered separately starting from K=4. From K=5 to K=7 distinct ancestry components were assigned to *F. lybica* wildcats and domestic cats (*F. catus*), with the latter sharing more ancestry with the Tunisian wildcat. Furthermore, at these K values, Sardinian wildcats did not share any ancestry with domestic cats. At K=8, the analysis discriminated between Levantine and Sardinian wildcats. At this K value domestic cats are shown to share the highest amount of ancestry with the Tunisian wildcat whilst the Sardinian wildcats are modelled more similar to the Moroccan wildcat.

To account for uneven sampling and to focus more specifically on *F. l. lybica*/*F. catus*, we extracted all the *F. l. lybica* wildcats (from Morocco, Tunisia, Israel and Sardinia, n=8) and domestic cat (n=18) data from PANEL-A and prepared eight subpanels, each including all the wildcats and three randomly chosen domestic cats. Therefore, each subpanel consisted of a total of 11 individuals and, after imposing a minor allele count of 3, resulted in ~950,000 SNPs. We run the analysis for K values between 2 and 6. Given the fewer ancestries represented in these subpanels compared to the run made with PANEL-B, distinct African wildcat ancestries were already resolved at K=3 (Fig. S4). At K=2 North African and Levantine wildcat ancestries were discriminated, and domestic cats were modelled as mostly North African with varying levels of Levantine ancestry (27% to 42%). At K=3, one ancestry was maximized in the Sardinian wildcats, another in the Tunisian individual, and the third in the Levantine wildcats. Domestic cats possessed the same component as the Tunisian wildcat, while the Moroccan was the only one sharing ancestry with the Sardinian wildcats. Overall, the results showed the genetic distinctiveness of the Levantine wildcats, confirming their low affinity with domestic cats. The analysis also confirmed that the Moroccan individual is the one sharing more ancestry with the Sardinian wildcats (Fig. S4).

*F statistics of modern wild and domestic cats.*
Both phylogenetic and population structure analyses highlighted the affinity between Moroccan and Sardinian wildcats and suggested that North African wildcats have a closer relationship with modern domestic cats than to Levantine wildcats. To corroborate these results, we assessed patterns of shared genetic drift using F-statistics in Admixtools v.2.0.4 (*139*), *Felis chaus* as the outgroup. We used PANEL-A for all F-statistics involving only modern wild and domestic cats. Outgroup-$f_3$ statistics in the form of $f_3$(*Felis chaus*, Levantine/Sardinian/African wildcats, domestic cats) were computed to assess the amount of shared drift between domestic cats (n=18) and *F. l. lybica* wildcats. Results showed that all present-day domestic cats share more drift with the Tunisian wildcat although standard error (s.e.) bars overlapped with the Levantine wildcats in six instances (Fig. S5a). Therefore, we also run $f_4$ statistics in the form of $f_4$(*Felis chaus*, Domestic cats; Tunisian wildcat, Levantine/Moroccan/Sardinian wildcats), which showed that all 18 domestic cats are significantly more closely related to the Tunisian wildcat than to the other wildcats (Fig. S5b).
Both outgroup-$f_3$ and $f_4$ statistics also highlighted that domestic cats share the smallest amount of drift with the Sardinian wildcats. To better evaluate relationships between Sardinian wildcats and all other wild and domestic cats we used outgroup-$f_3$ statistics in the form of $f_3$(*Felis chaus*, Levantine/African wildcats/domestic, Sardinian wildcats). Results indicated that the Sardinian wildcats share the highest amount of drift with North African wildcats (firstly with the Moroccan wildcat and secondly with the Tunisian) and the lowest with the Levantine ones, with domestic cats showing intermediate values of shared drift (Fig. S6a). However, given the overlapping of s.e. bars we run $f_4$ statistics in the form of $f_4$(*Felis chaus*, Sardinian wildcats; Moroccan wildcat, Levantine/Tunisian/Domestic cats), which showed that Sardinian wildcats are significantly more closely related to the Moroccan wildcat than to any other wild and domestic cats (Fig. S6b). The distinctive relationship of domestic cats and Sardinian wildcats with North African wildcats (i.e. the Moroccan and the Tunisian individuals, respectively), was further corroborated by $f_4$ statistics in the form of $f_4$(*Felis chaus,* Domestic/Sardinian wildcats; Morocco, Tunisia) (Fig. S7). Future works, including more African and Middle/Near East genomes, possibly with more than one sample per region, are needed and expected to refine relationships between cat populations, both wild and domestic.

*Principal component analysis of modern and ancient cats*
For ancient data analyses, PANEL-C was utilized. First, we converted this panel to eigenstrat format using the *convertf* function of eigensoft v.7.2.1 (*140*). We then applied a principal component analysis using smartpca v.16000 (*141*), with the option LSQproject set to YES, so as to project low- to middle-coverage samples to the coordinate space built with modern high-coverage individuals. Based on the PCA results (Fig. 1C), we were able to assign the taxonomic status of the ancient samples and to refine the chronology of *F. l. lybica*/*F. catus* introduction throughout Europe and Anatolia (Fig. S8). A zoom of the wildcat cluster of Fig. 1C is provided in Fig. S16. To increase resolutions, we extracted all the modern and ancient *F. l. lybica* and *F. catus* individuals (n=70) from PANEL-C and ran smartpca with the same option used above (LSQproject set to YES) (Fig. 2A).

*F and D statistics of modern and ancient cats*
We explored patterns of shared genetic drift between ancient and modern wild and domestic cats (*F. l. lybica* and *F. catus*) using F-statistics and PANEL-C in Admixtools2. For most of the

analyses, modern samples were grouped according to the clades detected in the phylogenetic ML tree (Fig. 1B), distinguishing Levantine, North African (Morocco and Tunisia), Sardinian and domestic cats. In the PCA restricted to *F. l. lybica*/*F. catus* individuals (Fig. 2A), the sample GSA01 clustered with present-day Sardinian wildcat. Therefore, we ran outgroup-$f_3$ statistics in the form of $f_3$(outgroup; ancient/modern wild/domestic cats, GSA01/modern Sardinia) which confirmed the strong affinity between the present-day Sardinian wildcat and GSA01 (Fig. S13a and S13b).

Using $f_4$ statistics we tested which of the three modern groups (Levantine and North African wildcats and domestic cats) are genetically closer to the ancient samples (n=43) identified as *F. l. lybica*/*F. catus* from our PCA. The test $f_4$(outgroup, ancient *F. l. lybica*/*F. catus*; Levantine wildcats, African wildcats) showed that our ancient samples share more genetic drift with North African wildcats than with Levantine wildcats (Fig. 2B). By repeating this test using the two North African wildcats separately $f_4$(outgroup, ancient *F. l. lybica*/*F. catus*; Levantine wildcats, Tunisian/Moroccan wildcats), we found that the Tunisian wildcat has a much greater affinity with ancient domestic cats, as indicated by higher z-scores compared with the Moroccan individual (Fig. S14a and S14b). Subsequently, we tested whether the ancient samples were more closely related to modern domestic cats or North African wildcats using $f_4$(outgroup, ancient *F. l. lybica*; African wildcats, Domestic). The results showed that the ancient samples, except GSA01, share more genetic drift with domestic cats. GSA01 showed a distinct ancestry, more related to North African wildcats than to domestic cats (Fig. 2C), similar to patterns observed in modern Sardinian wildcats (see outgroup-$f_3$ in Fig. S6, MDS in Fig. S2, and admixture in Fig. S3-S4).

We used *D* statistics to assess patterns of gene flow between European wild and domestic cats over time. First, we tested whether ancient and modern *F. silvestris* experienced gene flow from *F. l. lybica*/*F. catus* using the form *D*(*Felis chaus*, Levantine/Moroccan/Tunisian wildcat; modern Portuguese *F. silvestris*, other ancient and modern *F. silvestris*). We used *F. l. lybica* wildcats from the Levant and North Africa as source of gene flow since admixture analysis (K=2-4 in Fig. S3) suggested that most modern domestic cats possess some degree of European wildcat ancestry. The modern Portuguese wildcat, previously identified as non-admixed (*15*), served as European wildcat reference. The results highlighted that the Portuguese wildcat carries some *F. l. lybica*/*F. catus* ancestry when compared with Neolithic and Chalcolithic western European wildcats, as well as with Mesolithic and Neolithic wildcats from Italy and Ireland (Fig. S17). Hence, these ancient samples represent a better proxy for the European wildcat ancestry than the Portuguese wildcat. For this reason, we repeated the test by selecting the most ancient of these samples (three Mesolithic samples from northern Italy and two Neolithic/Chalcolithic from Spain) as European wildcat reference instead of the Portuguese wildcat. We pooled the five ancient samples to minimize data loss due to missing data.

Afterwards, to test if we could also use the modern domestic cats of our comparative dataset as source of gene flow into the European wildcats we assessed if they were devoid of *F. silvestris* ancestry by running *D* statistics in the form *D*(*Felis chaus,* ancient pool *F. silvestris*; Tunisian wildcat, domestic cat). The Tunisian wildcat was modelled in all the previous analyses (see MDS, admixture and outgroup-$f_3$ statistics) as the closest wildcat genome to the ones of domestic cats, so it was selected as the non-admixed *F. l. lybica* reference for this test. The results (Fig. S18) showed that almost all the present-day domestic cats (16 of 18) share an excess of alleles with European wildcats. The domestic Ocicat sample showed the lowest non-significant z-score. Therefore, we repeated the test of gene flow from *F. l. lybica*/*F. catus* into ancient and modern European wildcats using the form *D*(*Felis chaus, F. l. lybica Levant*/*F. catus* Ocicat breed, ancient pool *F. silvestris,* other modern and ancient *F. silvestris*), with results shown in Fig. 3A. Finally, we explored patterns of gene flow from European wildcats into our ancient domestic cats, using the test *D*(*Felis chaus*, ancient pool *F. silvestris*; *F. catus* Ocicat breed, ancient domestic cats), with results shown in Fig. 3B.

*$f_4$-ratio*

We estimated admixture proportions in all the ancient and modern European wildcats (*F. silvestris*) that resulted positive to gene flow (see *D* statistics in previous paragraph and Fig. 3A). To do so, we used qpf4ratio test in admixtools2. This test uses the ratio (α) of two $f_4$ statistics under the phylogenetic model illustrated in Fig. S19: $\alpha = f_4(A,O;X,C)/f_4(A,O;B,C)$, where A = Mesolithic to Chalcolithic *F. silvestris* from Italy and western Europe, B = Bronze Age *F. silvestris* from Italy, C = *F. l. lybica* from the Levant (n=3) or domestic cats (Ocicat Breed), O = *F. chaus,* X = test. Results are in Fig. 3C.

We used the same test to estimate admixture proportions (1 – α) of the ancient domestic cats that tested positive to gene flow from the European wildcat, given the same phylogenetic model used above (Fig. S19) but using only the modern Ocicat domestic breed as population C. Results are in Fig. 3D.

*Mitochondrial phylogeny and demography*

We used the multi sequence mtDNA alignments fasta file (see paragraph 5.3 for details) to build maximum-likelihood (ML) and Bayesian phylogenetic trees using the software IQ-TREE v.2.2.0-beta (*142*) and BEAST v.2.7.6 (*143*) respectively.

For ML phylogenetic analysis we generated a tree using 1,000 bootstrap replicates in IQ-TREE (-B 1000) and 1,000 SH approximate likelihood ratio test replicates (--alrt 1000) for assessing branch supports and the ModelFinder Plus option for model selection (-m MFP). The tree was rooted by using the *F. margarita* mtDNA (NC_028308.1) as outgroup (iqtree2 -s Cats_multisequence_alignement.fasta -B 1000 -alrt 1000 -m MFP -o FMA08). The resulting tree (Fig. S9) was visualized in Figtree v1.4.4 (http://tree.bio.ed.ac.uk/software/figtree/).

For the Bayesian phylogenetic analysis, the multi sequence fasta file mtDNA alignment was converted to the nexus format with Aliview v.1.28 (*144*) and different partitions were specified based on the mtDNA *F. catus* NCBI annotation (accession number: NC_001700.1) for the D loop, RNAs, 1st, 2nd, and 3rd codon sites. The nexus file was then loaded into BEAUTi (which is part of the BEAST package) to convert it into the BEAST input file and using the following parameters: a strict molecular clock and a lognormal distribution with a mean in real space of 1.0E-8; a HKY+ G substitution model with gamma category count of 4; tip dates were specified using the mean radiocarbon date, when available, or alternatively the dates based on the archaeological contexts. The resulting tree (Fig. 1E and Fig. S10) was visualized using Figtree v1.4.4 (available at http://tree.bio.ed.ac.uk/software/figtree/).

Both phylogenetic reconstructions (i.e., Maximum-likelihood and Bayesian) resulted in concordant topologies (with few minor exceptions at the terminal nodes). The higher resolution achieved using complete mtDNA genomes revealed the presence of three well-supported (posterior probabilities >0.86) subclades in the topology of *F. l. lybica*/*F. catus* haplogroup IV-A, while confirming its distribution in both ancient Anatolia and Southeast Europe. We named these subclades IV-A0, IV-A1 and IV-A2 (Fig. 1E). Of the four Neolithic cats from Anatolia, three possessed lineages belonging to the haplogroup IV-A0, and one IV-A2. Four of the five specimens from Neolithic to Chalcolithic Bulgaria possessed the haplogroup IV-A1. All the other samples from across the rest of Europe dating from the 9th to the 2nd millennium BCE (n=12) as well as one cat from Chalcolithic Bulgaria had haplogroup I mtDNA, characteristic of *F. silvestris*. These results, coupled with the analyses of the nuclear genomes, highlighted several instances of mitonuclear discordance mostly regarding the European wildcat *F. silvestris* (Fig. S11).

Together with IV-A, IV-C was the most represented *F. l. lybica* haplogroup in our ancient dataset (48% and 44%, respectively). It occurred continuously in European domestic cats starting from the Roman era (n=8) through Medieval and post-Medieval times (n=15) until present-day (n=7). It was also found in two wildcats from Sardinia, and in a wildcat from Morocco, which carried a basal lineage. The other *F. l. lybica* clades were less frequent in the overall dataset that we compiled. IV-B was carried by a modern wildcat from the Levant, whereas IV-E was present in two modern Sardinian wildcats and in an ancient cat (GSA01) from the site of Genoni, in Sardinia, as well as in two domestic cats from Byzantine Turkey.

A domestic cat from Byzantine Turkey (MRY09) was found to carry a haplotype which clusters with the ones of present-day Asian wildcat (*F. l. ornata*). We tested this sample for gene flow from *F. l. ornata* by computing *D*(*F. chaus*, *F. l. ornata* (n=4), ancient and modern domestic cats, MRY09). The results (Fig. S20) showed that this sample did not carry any significant excess of ornata alleles compared to any other ancient domestic cat. Hence, the mitonuclear discordance may be the result of incomplete lineage sorting.

A modern domestic cat carried IV-D, which was typical of ancient cats from Northern Europe dated to the Viking era (*15*). Finally, all cats originating from Italy and Spain dated from the Mesolithic to the Bronze Age carried clade I mtDNAs typical of *F. silvestris*. Clade I was composed of two main clusters. One included ancient and modern cats from the Balkans, central Europe and northeast Italy. The other encompassed ancient and modern wildcats from Italy and Spain, a Neolithic wildcat from Ireland and a modern wildcat from Scotland.

We made use of mtDNA data to infer demographic histories of both domestic cats (clade IV, n= 70) and European wildcats (clade I, n=33) by estimating the effective population size ($N_e$) with the Coalescent Bayesian Skyline (BSP) method implemented in Beast v.2.7.6 (*143*). For both analyses we created two partitions, for the D-loop and the coding region; we used the HKY+ G substitution model with gamma category count of 4 for both partitions; we set tip dates using the mean radiocarbon date, when available, or alternatively the dates based on archaeological contexts; we did not fixed a mutation rate; we set a strict molecular clock; we ran three independent MCMC with 100,000,000 iterations each. The independent runs were merged using LogCombiner v.2.7.6 after discarding 10% of the states as burn-in, resulting in a total of 270,000,000 iterations for both analyses. Finally, we visualized the skyline plots with Tracer v.1.7.2.

The substitution rate estimated by BSP for the European wildcat clade I resulted in 4.91E-8 (95% HPD interval 3.36E-8 to 6.46E-8) and 1.84E-7 (95% HPD interval 1.18E-8 to 2.53E-8) substitution per site per year for the coding regions and the D-loop, respectively. The resulting skyline plot of the European wildcat clade I (Fig. S12) from across Europe showed a post-LGM (Last Glacial Maximum) population expansion starting about 11,000 years ago, possibly coinciding with the recolonization of Europe from glacial refugia, as previously suggested (*145*). By using a mean generation time of 3 years (*55*), the $N_e$ of the overall European wildcat population ranged from ~8,000 to ~20,000 individuals in the last 40,000 years. At a more regional level, recent estimates of $N_e$ based on nuclear data from present-day wildcat populations from Germany returned values ranging ~1,000-2,000 individuals in the last 40,000 years (*27*).

Our results showed that in ancient Anatolia four out of four *F. silvestris* wildcats dated 8,400-10,000 years ago possessed the *F. l. lybica* lineage IV-A. Assuming a complete mtDNA turnover, we used the effective population size estimates as upper ($N_e$=20,000) and lower ($N_e$=1,000) bounds to compute the mean time required for haplogroup IV-A to reach fixation in the ancient Anatolian wildcat population at varying *F. l. lybica* introgression proportions. This was computed using the formula from Kimura and Ohta (*29*) modified as in Posth *et al.* for the mtDNA (*28*). Under the assumption of neutrality, we modelled that even at fairly high introgression proportions (30%), *F. l. lybica* IV-A mtDNA would reach fixation in not less than 5,000 years in a small European wildcat population ($N_e$=1,000) (Table S8). This would place the admixture event not earlier than ~13,000 years ago, in the Late Pleistocene. As of today, genetic data of wildcats from Anatolia are restricted to the four Neolithic individuals presented here. Additional genome data from ancient and modern wildcats in Anatolia and the Caucasus, will help to infer more robust $N_e$ in European wildcats and more elaborated admixture scenarios in the future.

For the domestic cat clade IV, the BSP estimates of substitution rates resulted in 4.53E-8 (95% HPD interval 1.90E-8 to 7.40E-8) and 2.94E-7 (95% HPD interval 1.16E-7 to 4.87E-7) substitutions per site per year for the coding regions and the D-loop respectively. The resulting skyline plot of the domestic cat clade IV showed a population decline occurring between ~3,000 and ~2,000 years ago and a subsequent sharp increase of population size (Fig. S15). Such a trend can be consistent with a scenario of domestication process involving few mtDNA lineages (leading to the observed population decline) and then a rapid expansion following the human-mediated dispersal of domestic cats.

#### Supplementary figures

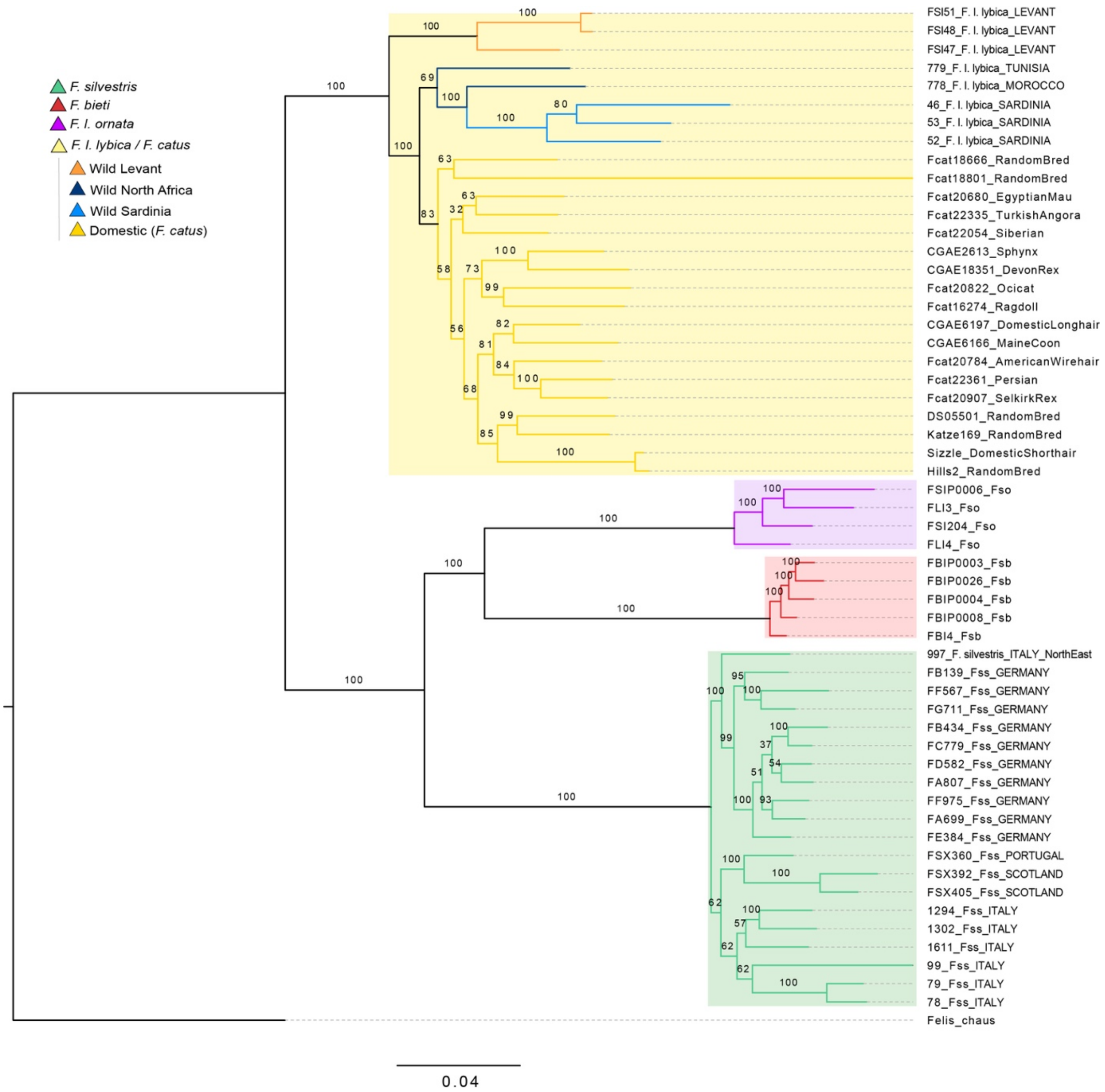

**Fig. S1. Nuclear phylogenetic tree.** Maximum-likelihood phylogenetic tree of autosomal SNPs of present-day wild and domestic cats. The tree was built from 403,868 SNPs of 55 high-coverage genomes and with 500 bootstrap replicates. The jungle cat, *F. chaus* was used as outgroup. Values at nodes indicate bootstrap supports. The four *Felis* taxonomic groups investigated are highlighted with different colors: purple for *F. l. ornata* (Asian wildcat); red for *F. bieti* (Chinese wildcat); green for *F. silvestris* (European wildcat); yellow for *F. l. lybica*/*F. catus* (North African wildcat and domestic cats). Within the *F. l. lybica*/*F. catus* clade different colors were used for the branches of the Levantine (orange), the North African and Sardinian wildcats (dark and light blue, respectively), and domestic cats (yellow).

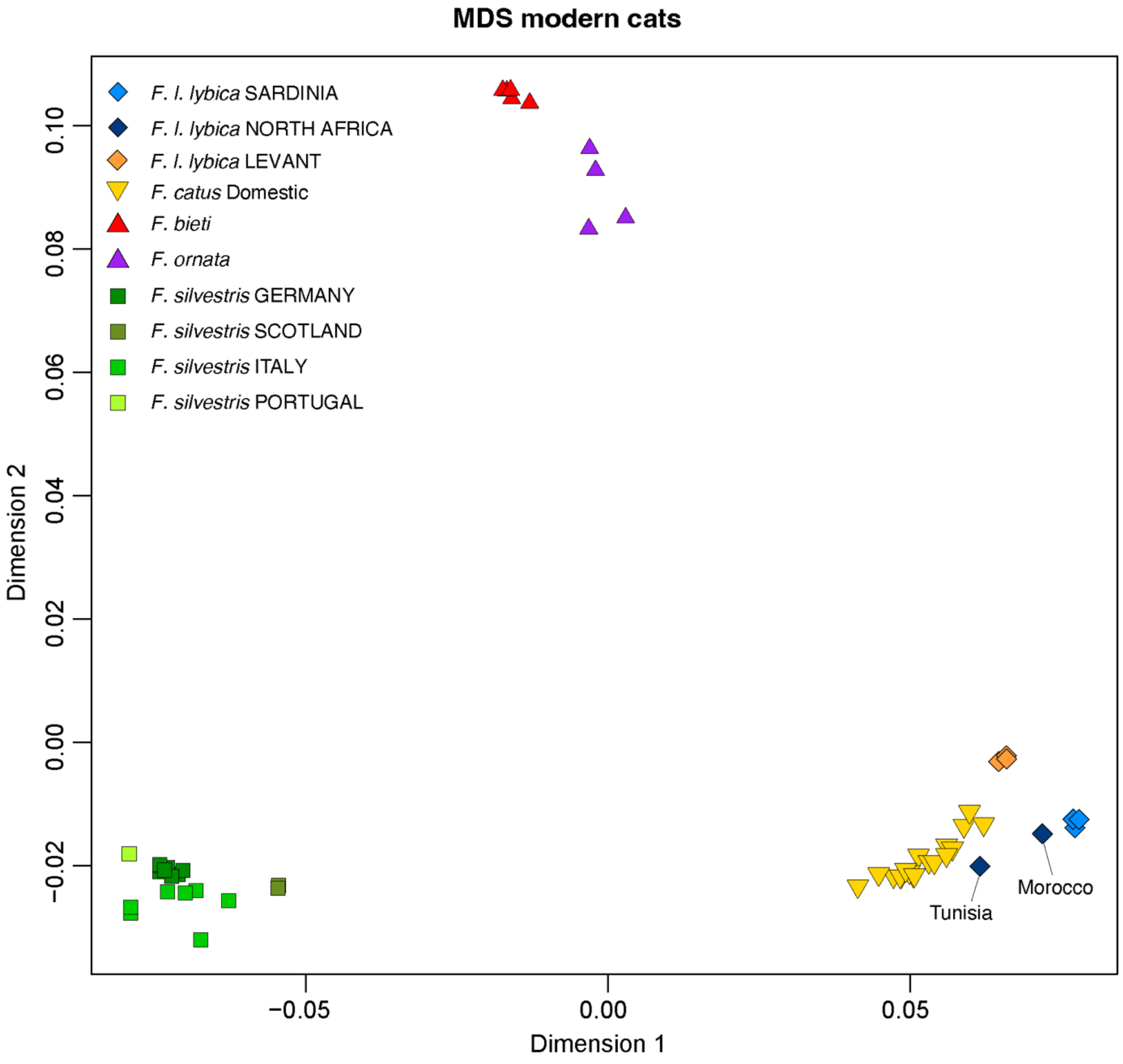

**Fig. S2. Multidimensional scaling.** Classic metric multidimensional scaling built from an Identity-by-State pairwise distance matrix of 3,954,446 autosomal SNPs from 55 present-day wild and domestic cat genomes. Each shape represents a single individual with shapes and colours as in the top-left legend.

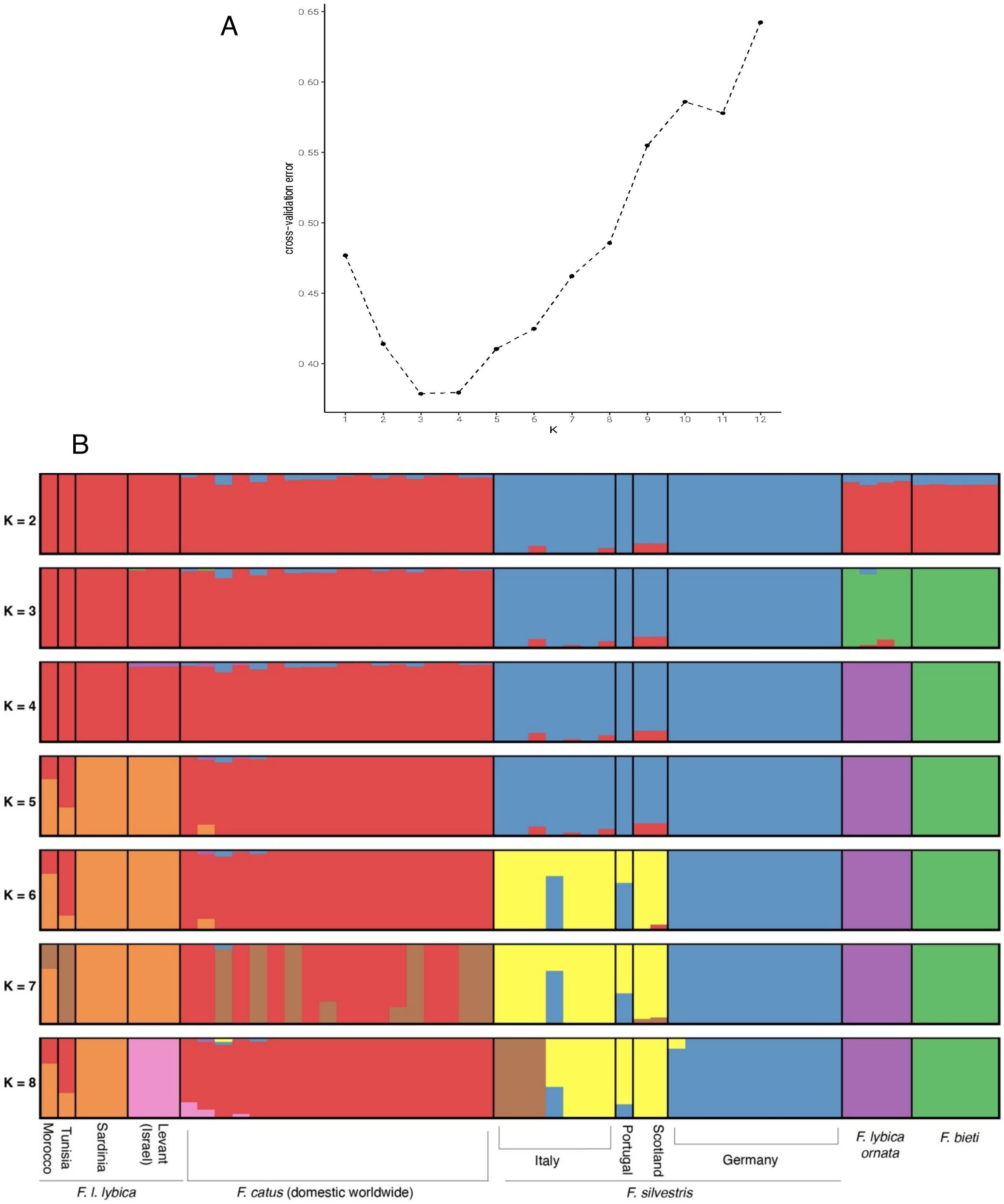

**Fig. S3. Admixture analysis of present-day wild and domestic cats. (A)** 5-fold cross-validation errors. **(B)** Admixture estimates computed from 403,868 autosomal SNPs of 55 present-day wild and domestic cat genomes. Barplot for K values from 2 to 8. Individuals are grouped based on taxonomy (*Felis catus*, *F. l. ornata, F. bieti*) and/or geography (Sardinia and Levant for *F. l. lybica;* Italy, Germany, Portugal and Scotland for *F. silvestris*).

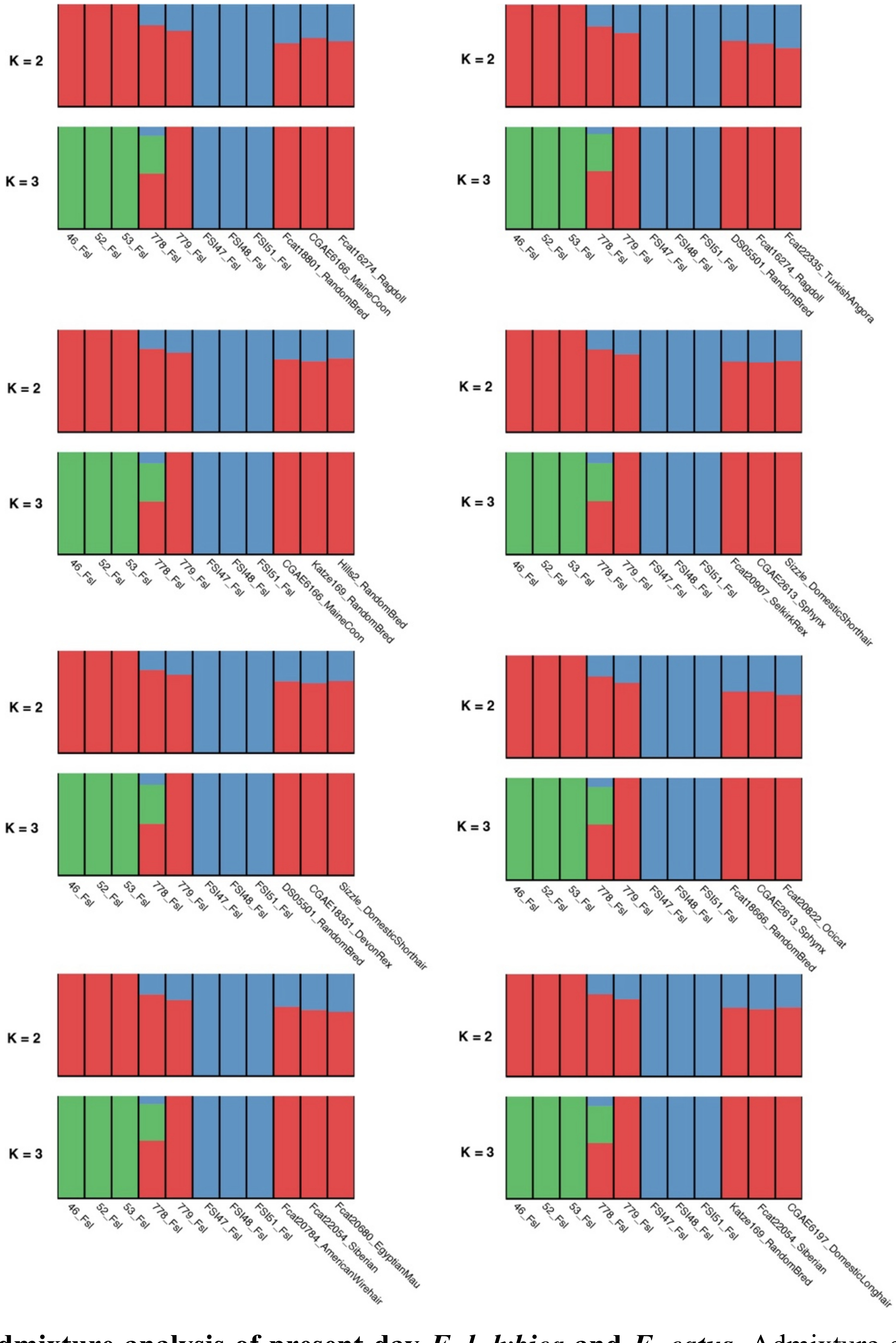

**Fig. S4. Admixture analysis of present-day *F. l. lybica* and *F. catus*.** Admixture estimates at K=2 and K=3 of eight subsamples consisting of 11 individuals of *F. l. lybica* and *F. catus*. In each barplot the first eight individuals are always the three Sardinian wildcat, the Moroccan, the Tunisian wildcat and lastly the three Levantine wildcats, while the last three bars represent three distinct domestic cats trios for each of the eight panels.

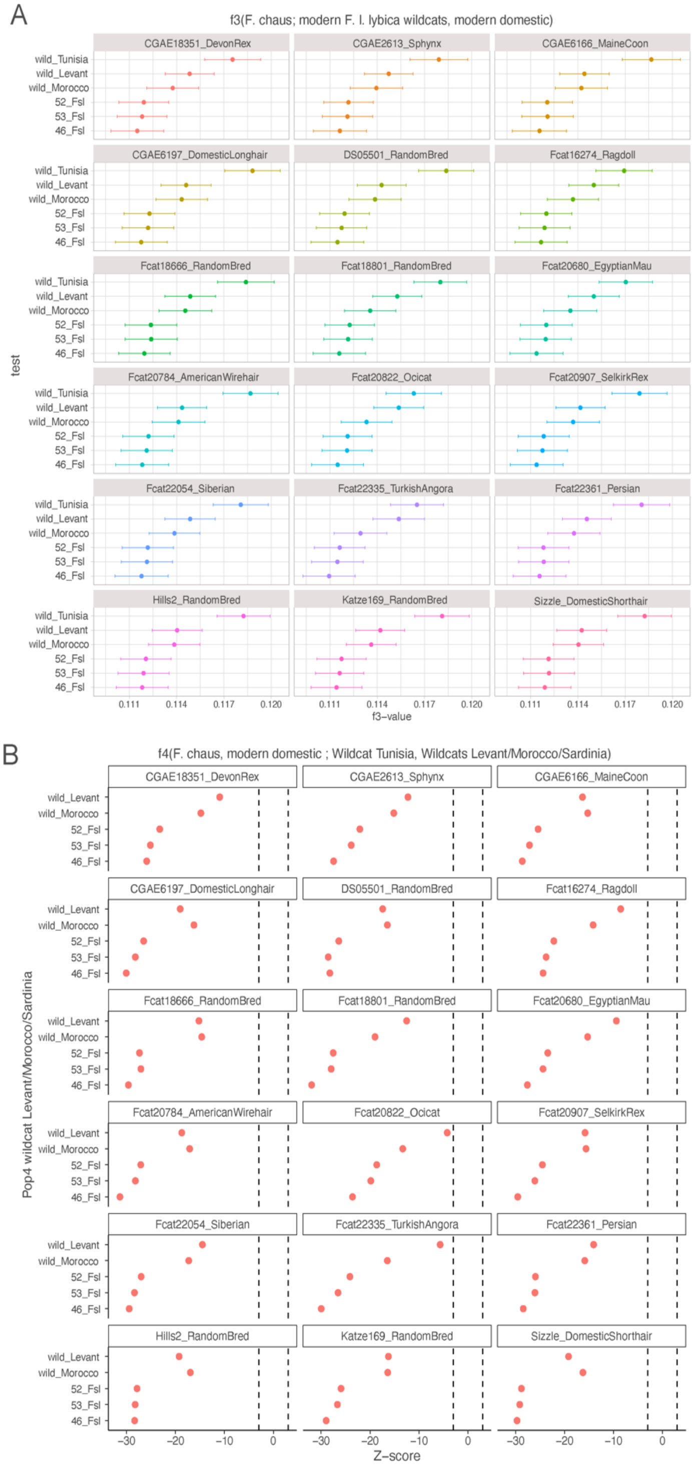

**Fig. S5. F-statistics of present-day domestic cats**. **(A)** Outgroup-$f_3$ statistics of domestic cats used to test the amount of shared drift with the three Sardinian, one Tunisian, one Moroccan, and three Levantine wildcats. The latter were grouped. The $f_3$ values is plotted with 3 standard errors. Colours were assigned automatically in R for different wildcat populations. **(B)** $f_4$ statistics of domestic cats used to test whether they share significantly more drift with the Tunisian wildcat compared to all other *F. l. lybica* wildcats. Levantine wildcats (n=3) were grouped. Dashed lines indicate significance threshold of +/- 3.

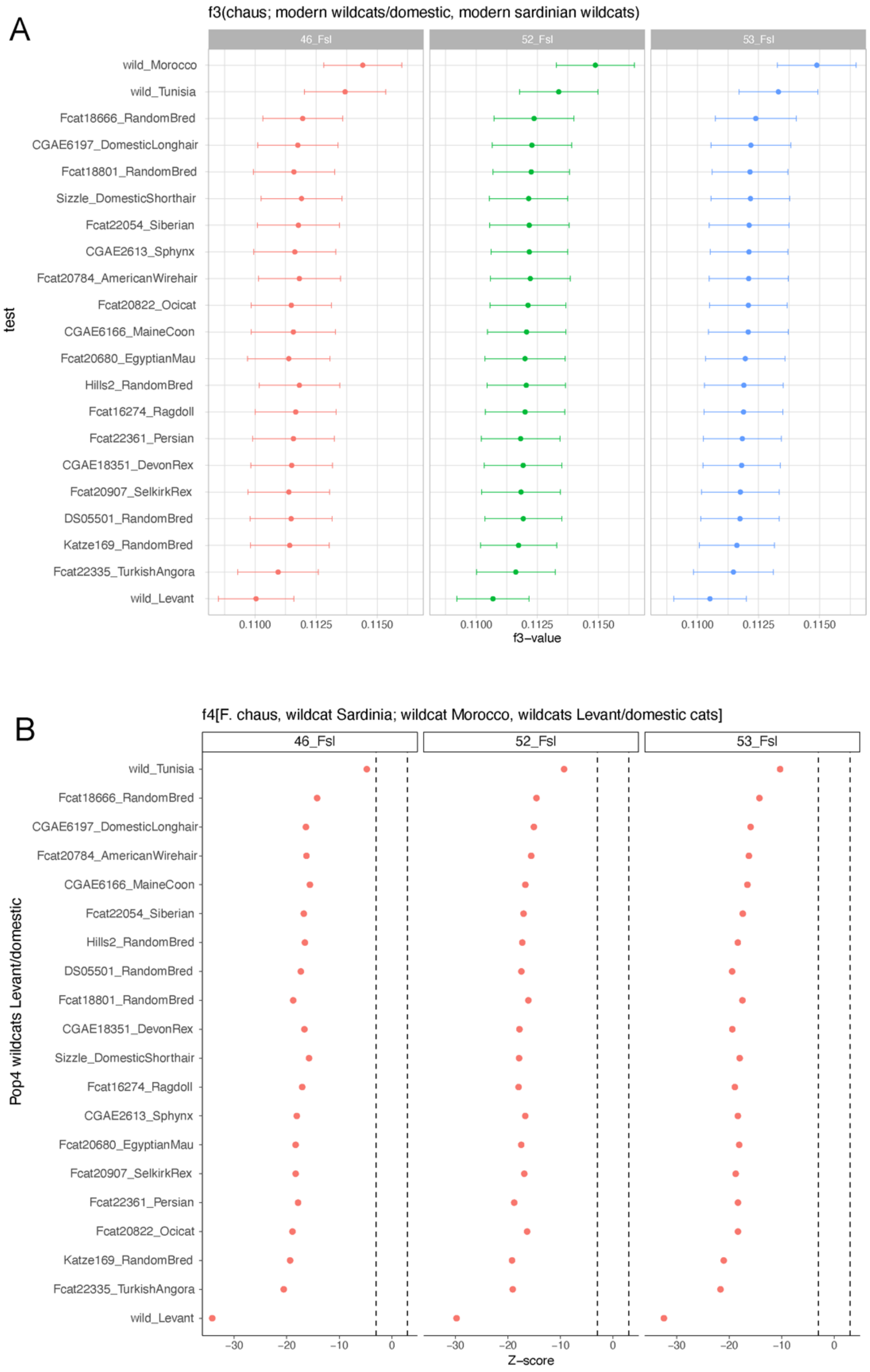

**Fig. S6. F-statistics of present-day Sardinian wildcats. (A)** Outgroup-$f_3$ of modern Sardinian wildcats (n=3) used to assess patterns of shared genetic drift with all others modern wild and domestic individuals in our dataset (only the Levantine wildcats were grouped, n=3). $f_3$ values were plotted with 3 standard errors. Colours were assigned automatically in R for different individuals. **(A)** $f_4$ statistics of Sardinian wildcats used to test whether they shared significantly more drift with the Moroccan wildcat compared to all other *F. l. lybica* wildcats. Levantine wildcats (n=3) were grouped. Dashed lines indicate significance threshold of +/- 3.

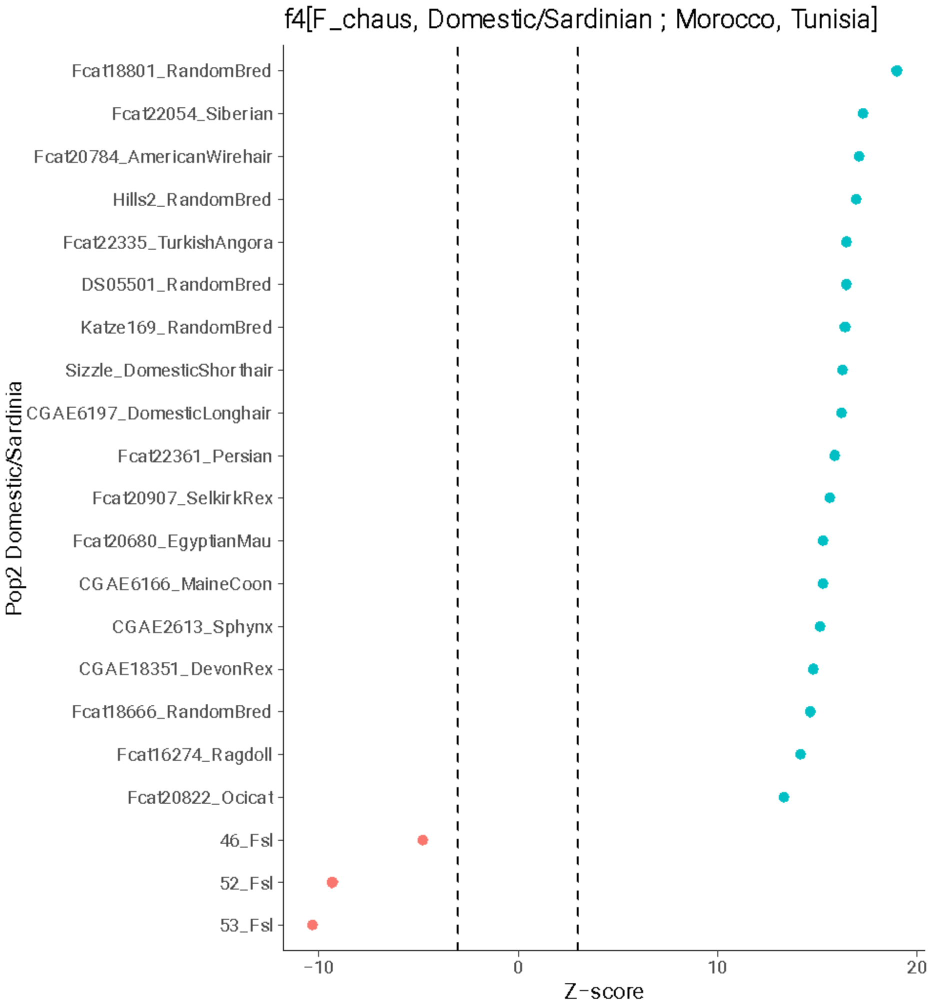

**Fig. S7. $f_4$ statistics of modern Sardinian wildcats and domestic cats.** Negative z-scores indicate an excess of alleles shared with the Moroccan wildcat while positive z-scores indicate excess of alleles shared with the Tunisian wildcat. All domestic and Sardinian specimens tested are on the y-axis. Red and blue dots indicate significant results (z-score > |3|). Dashed lines indicate significance threshold of +/- 3. Each dot represents an individual as reported on y-axis (for each domestic cat the ID-name and the breed is reported).

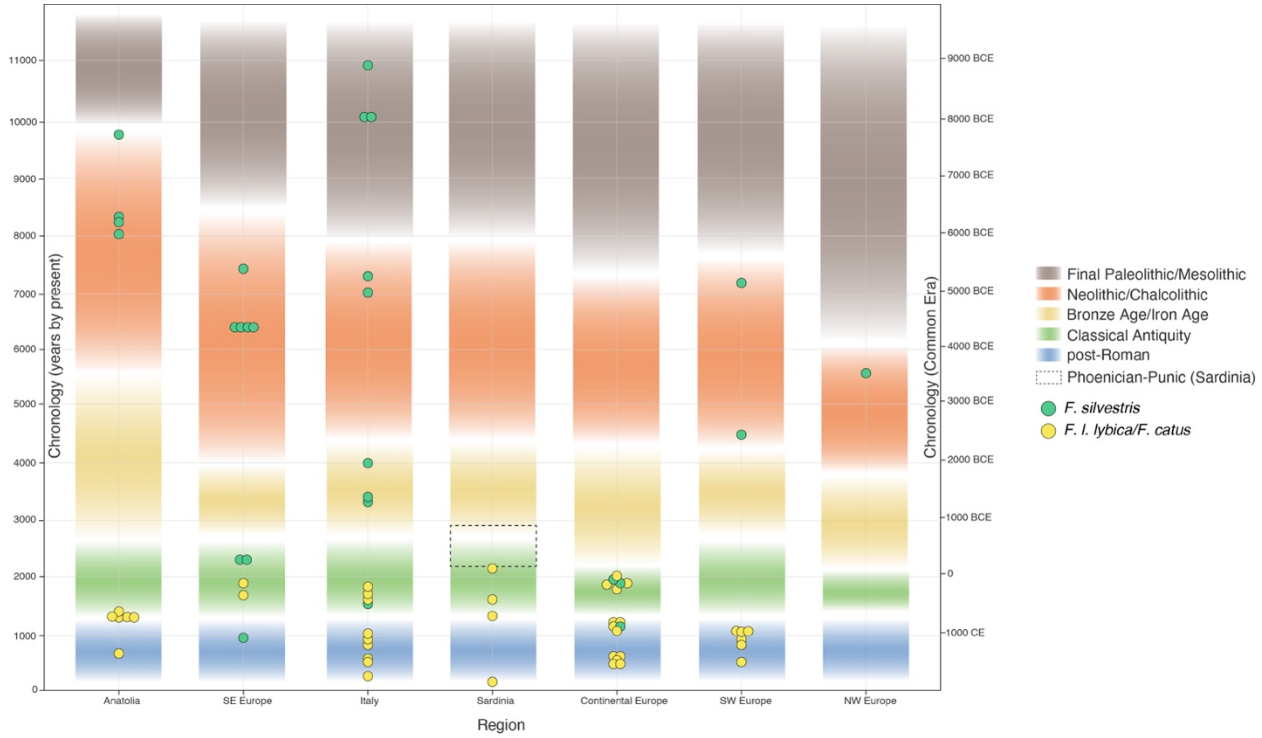

**Fig. S8. Cultural timeline and cat genomes.** Approximate cultural timeline showing the occurrence of European wildcats (green dots) and domestic cats (yellow dots) across different regions as revealed by our genome-wide analysis. Y-axis reports samples age in years before the present.

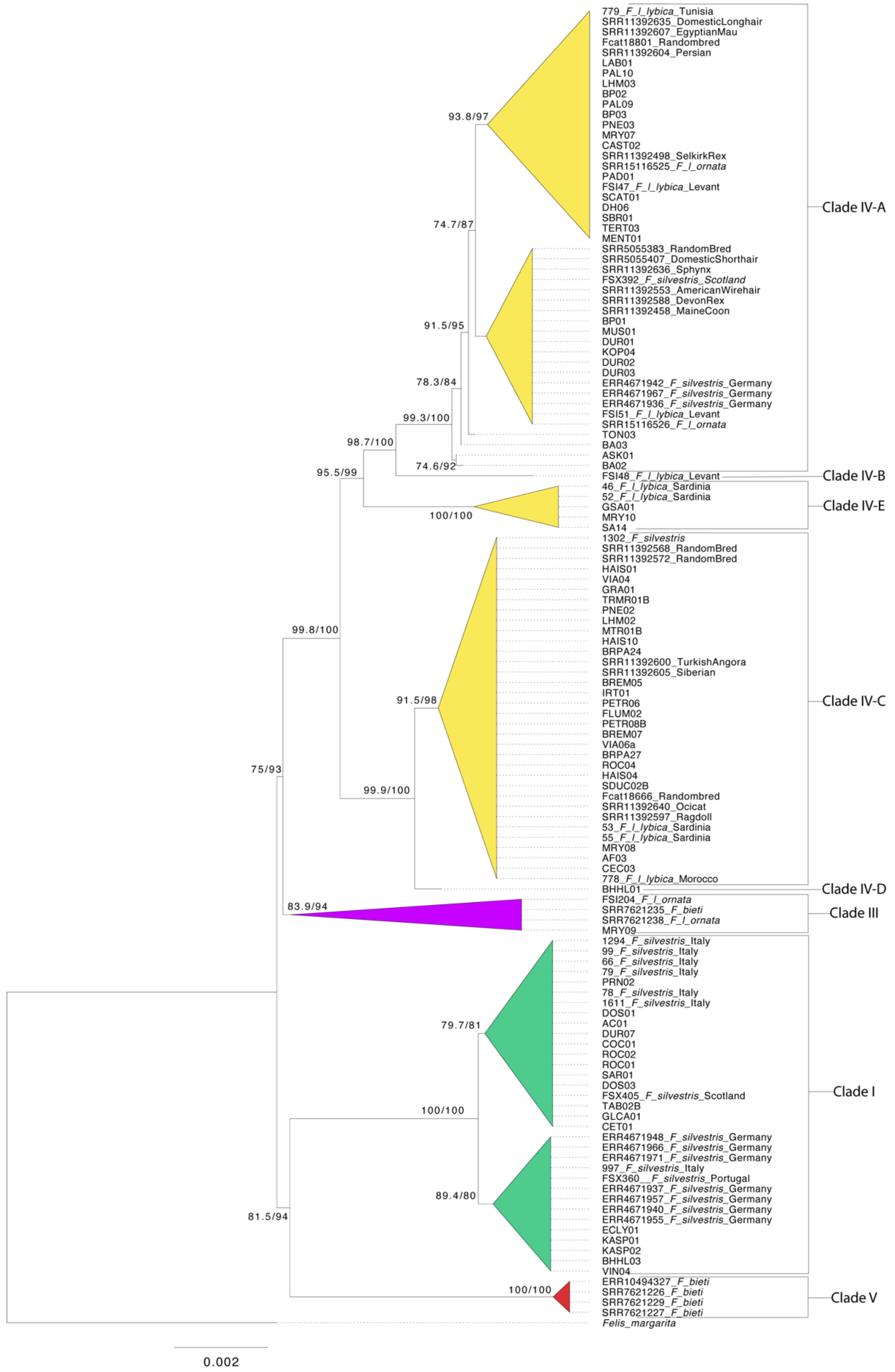

**Fig. S9. Maximum-Likelihood tree of complete mtDNAs.** Node values indicate SH-aLRT (first value) and bootstrap supports (second value). The *F. bieti* mtDNA clade is reported in red, *F. silvestris* in green, *F. l. ornata* in purple and *F. l. lybica* and *F. catus* in yellow.

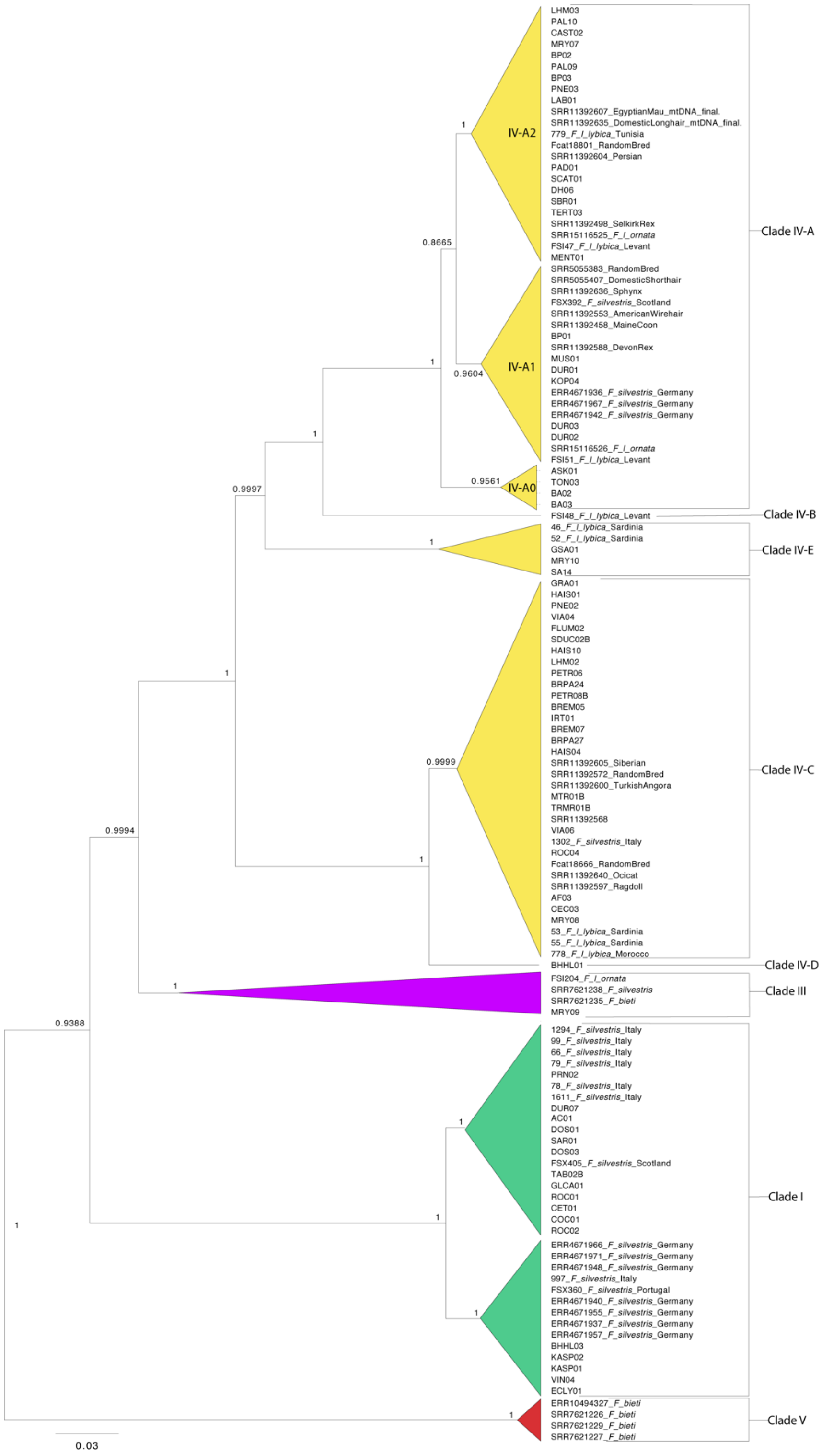

**Fig. S10. Bayesian phylogenetic tree of complete mtDNAs.** Detailed representation with individual samples names of the schematic tree reported in Fig. 1F. The main clades were collapsed in FigTree. The *F. bieti* mtDNA clade is reported in red, *F. silvestris* in green, *F. l. ornata* in purple and *F. l. lybica* and *F. catus* in yellow. Posterior probabilities are reported in the main nodes.

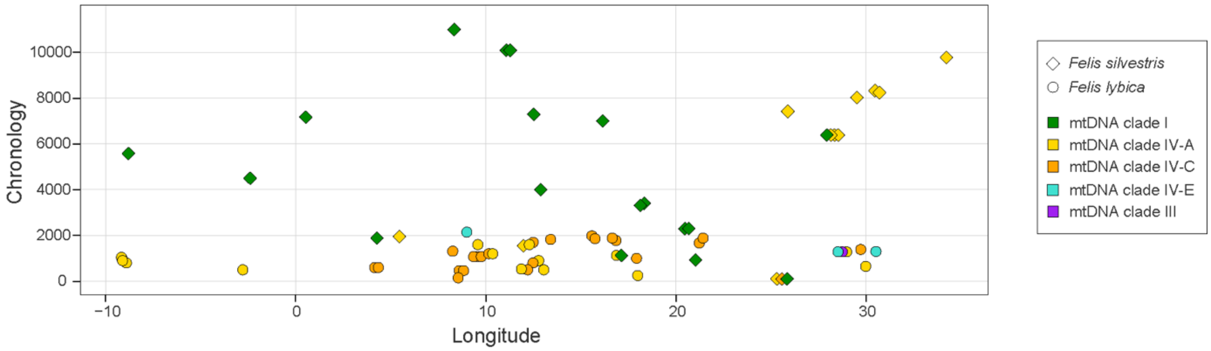

**Fig. S11. Spatiotemporal distribution of mitochondrial haplogroups.** Sample age is indicated in the y-axis in years before the present. Each symbol is an ancient cat. The shapes are based on the nuclear-based taxonomic assignment (*F. l. lybica and F. silvestris*) as inferred in the PCA from Fig. 1C.

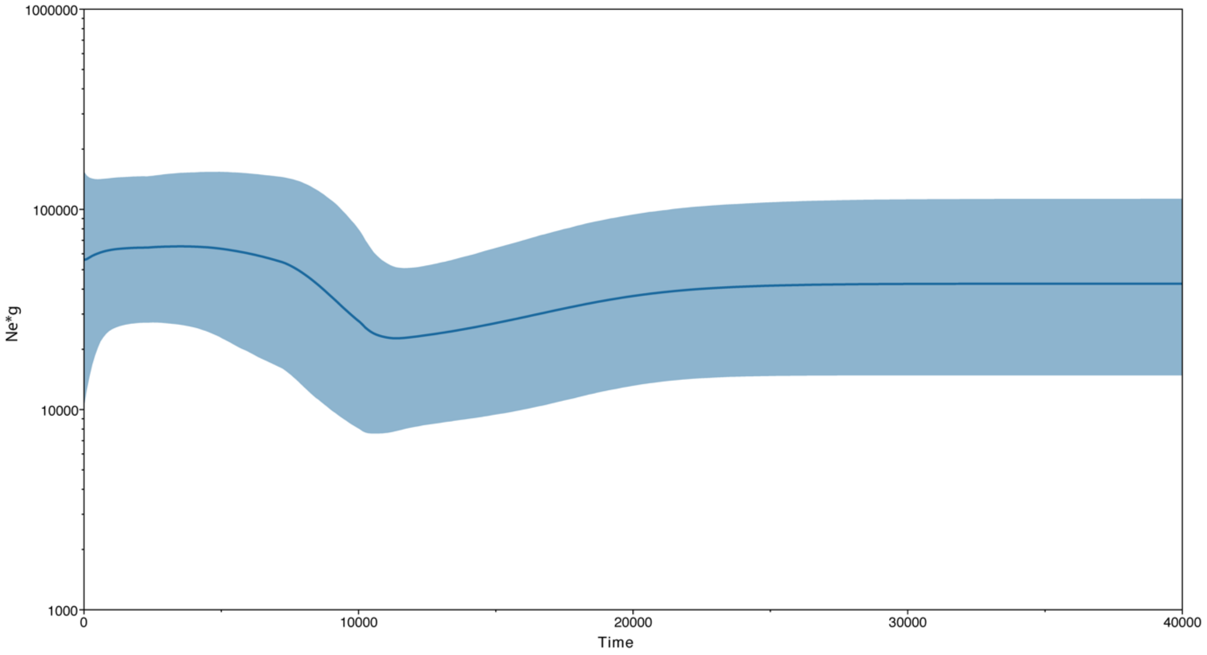

**Fig. S12. Bayesian Skyline plot of *F. silvestris* mtDNA.** Bayesian skyline plot derived from 15 modern and 17 ancient complete mtDNAs of *F. silvestris* clade I. The x-axis represents time in years before the present, and the y-axis is equal to $N_e$*g (the product of the effective population size and the generation time). The dark blue line indicates the mean estimate, while the blue area shows the 95% highest posterior density intervals.

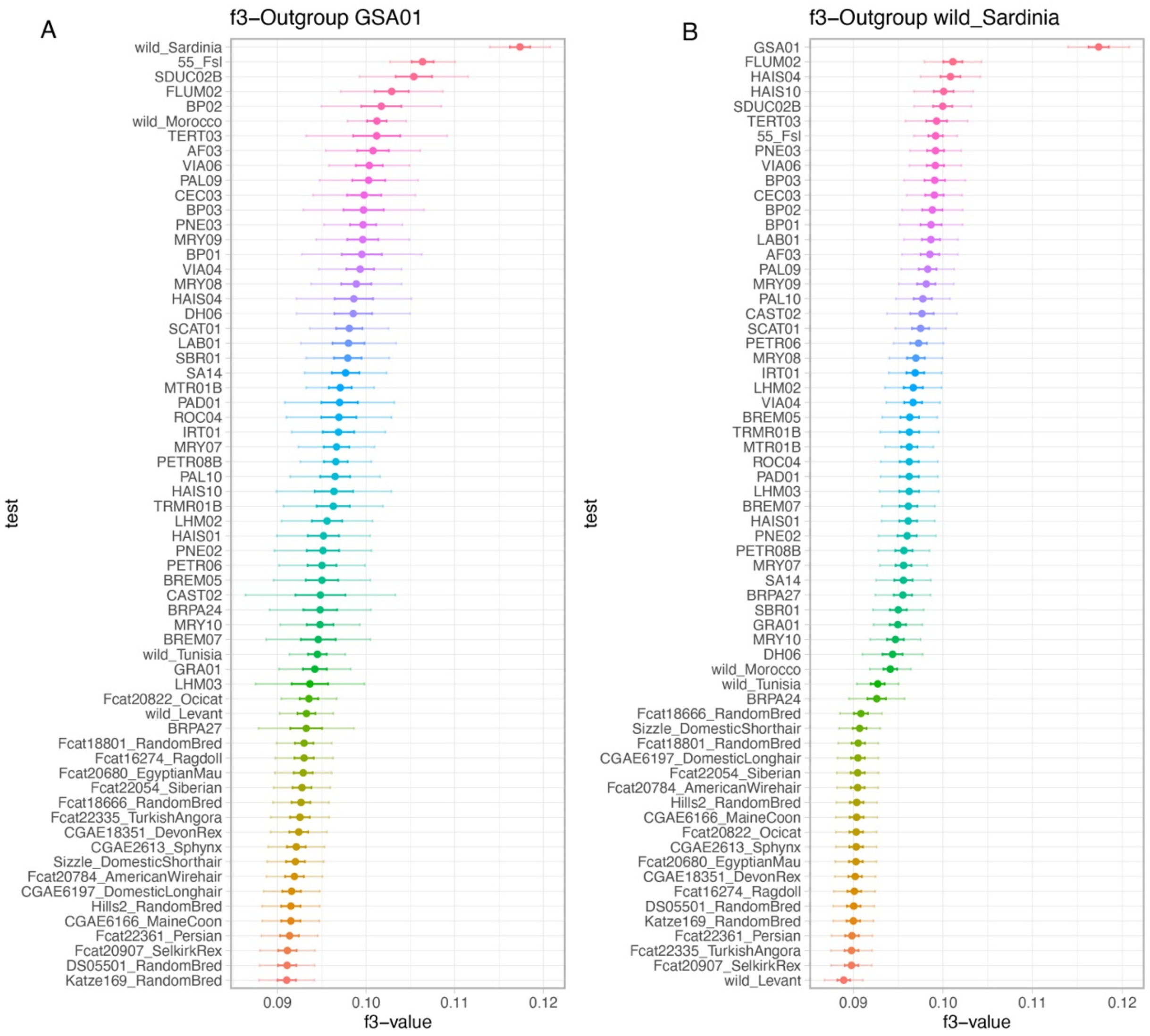

**Fig. S13. Outgroup-$f_3$ of GSA01 and modern wildcats from Sardinia.** The sample GSA01 **(A)** and modern **(B)** Sardinian wildcats (n=3, grouped) were tested for assessing patterns of shared genetic drift with all others modern and ancient wild and domestic *F. l. lybica*/*F. catus* individuals (y-axis) in our dataset. $f_3$ values were plotted with 3 standard errors. Colours represent different individuals and were assigned automatically in R.

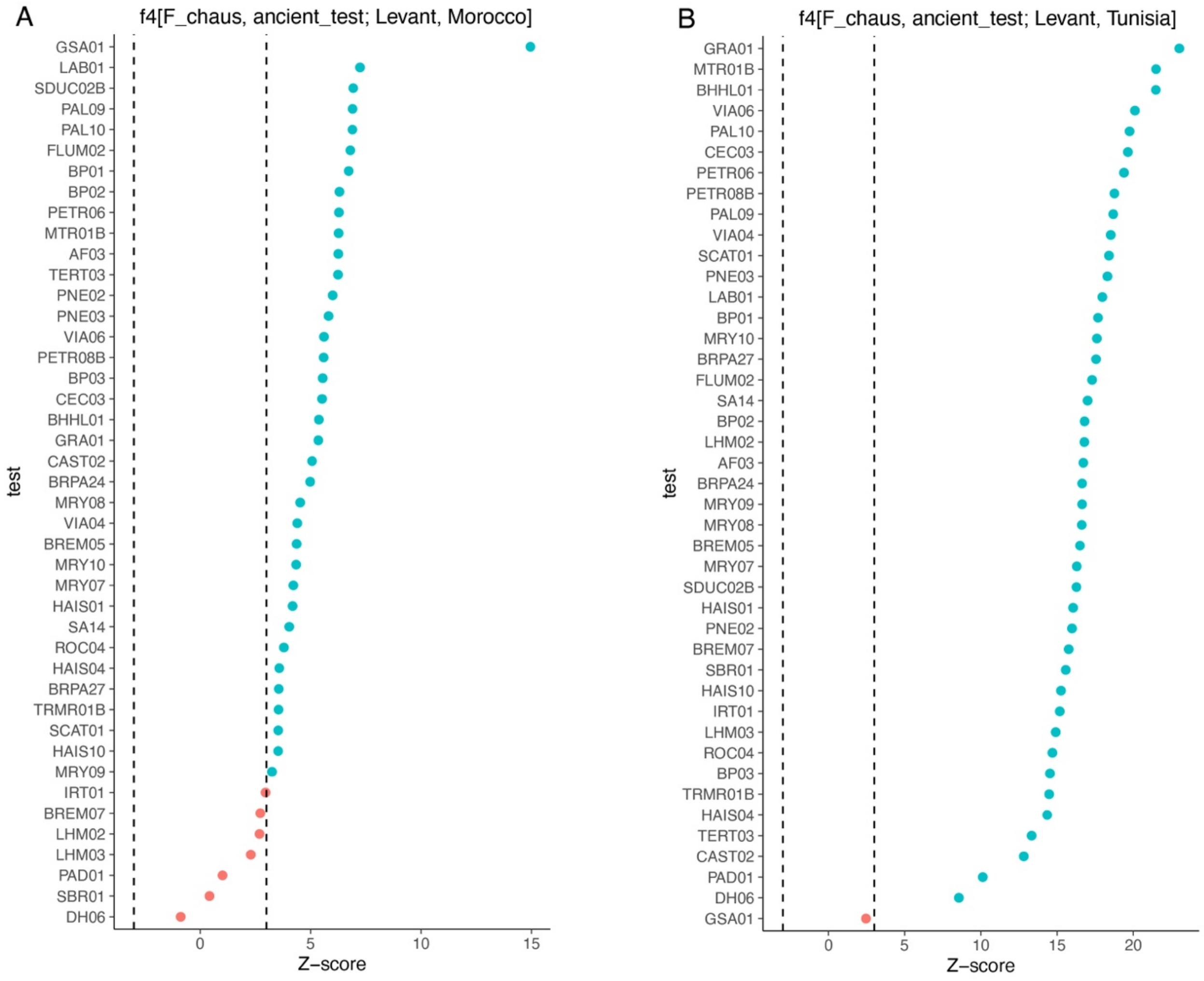

**Fig. S14. $f_4$ statistics of ancient *F. l. lybica*/*F. catus*.** Negative z-scores indicate an excess of alleles shared with the Levantine wildcats while positive z-scores indicate excess of alleles shared with the Moroccan wildcat (A) or the Tunisian wildcat (B). Dashed lines indicate significance threshold of +/- 3.

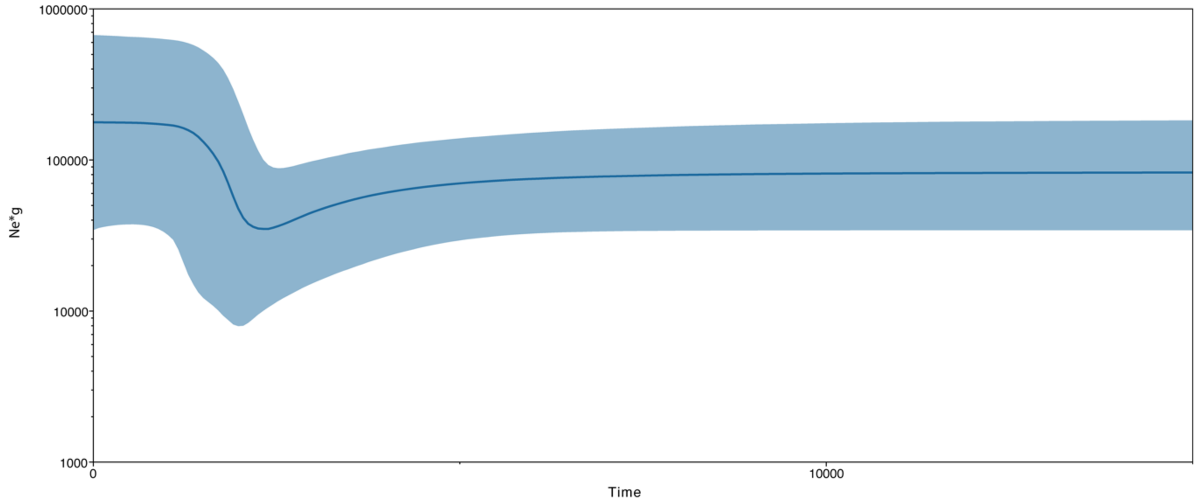

**Fig. S15. Bayesian Skyline plot of *F. catus* mtDNA.** Bayesian skyline plot derived from 18 modern and 42 ancient complete mtDNAs from *F. catus* clade IV. The x-axis is in units of years before present, and the y-axis is equal to $N_e$*g (the product of the effective population size and the generation time). The black line is the mean estimate, and the blue area shows the 95% highest posterior density intervals.

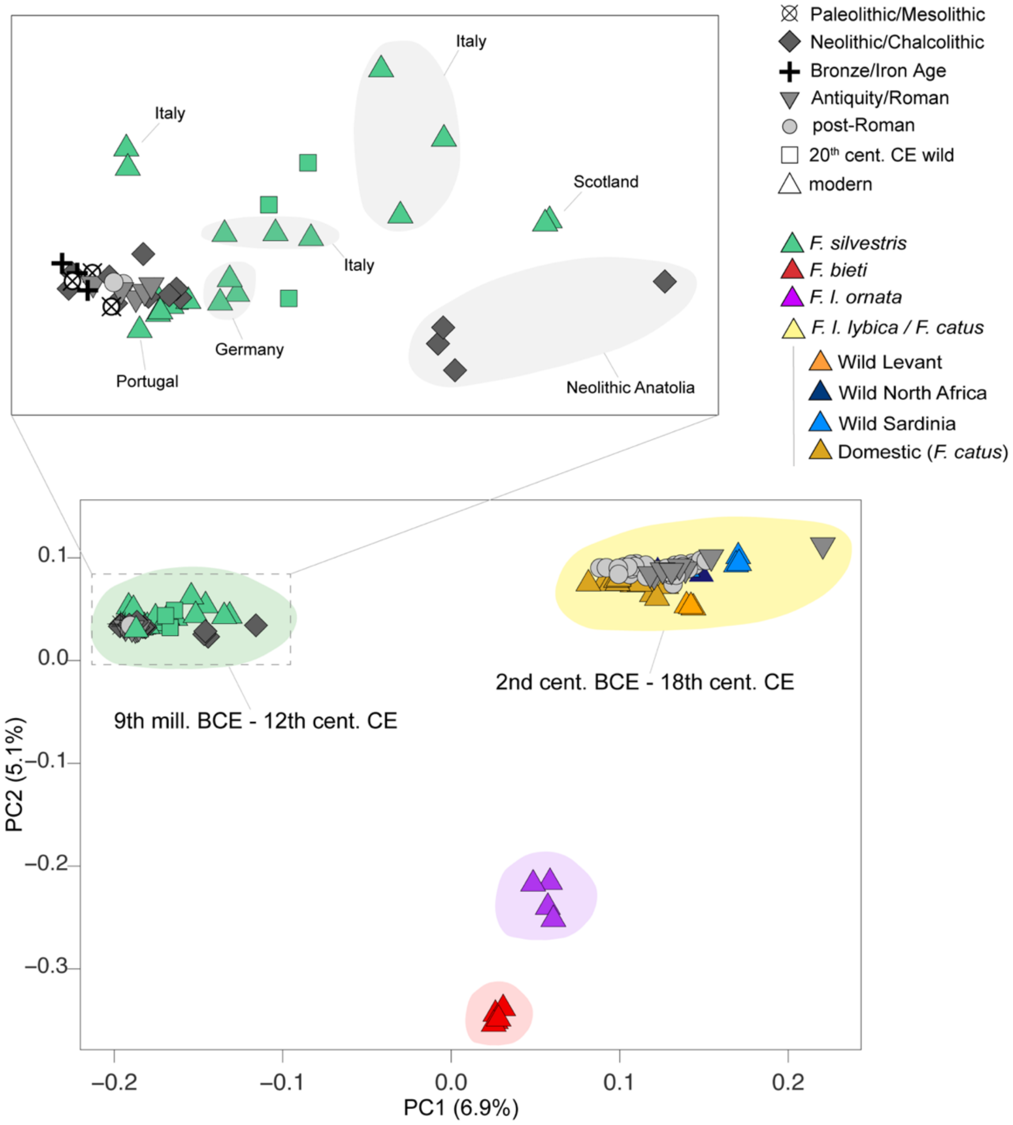

**Fig. S16. PCA of ancient and modern cats.** Principal component analysis built by projecting low coverage samples onto the coordinate space defined by modern mid-range coverage wild and domestic cats as in Figure 1C. with a zoom (inset) of the European wildcat cluster (*F. silvestris*). Colours and shapes of the symbols are as in the legend.

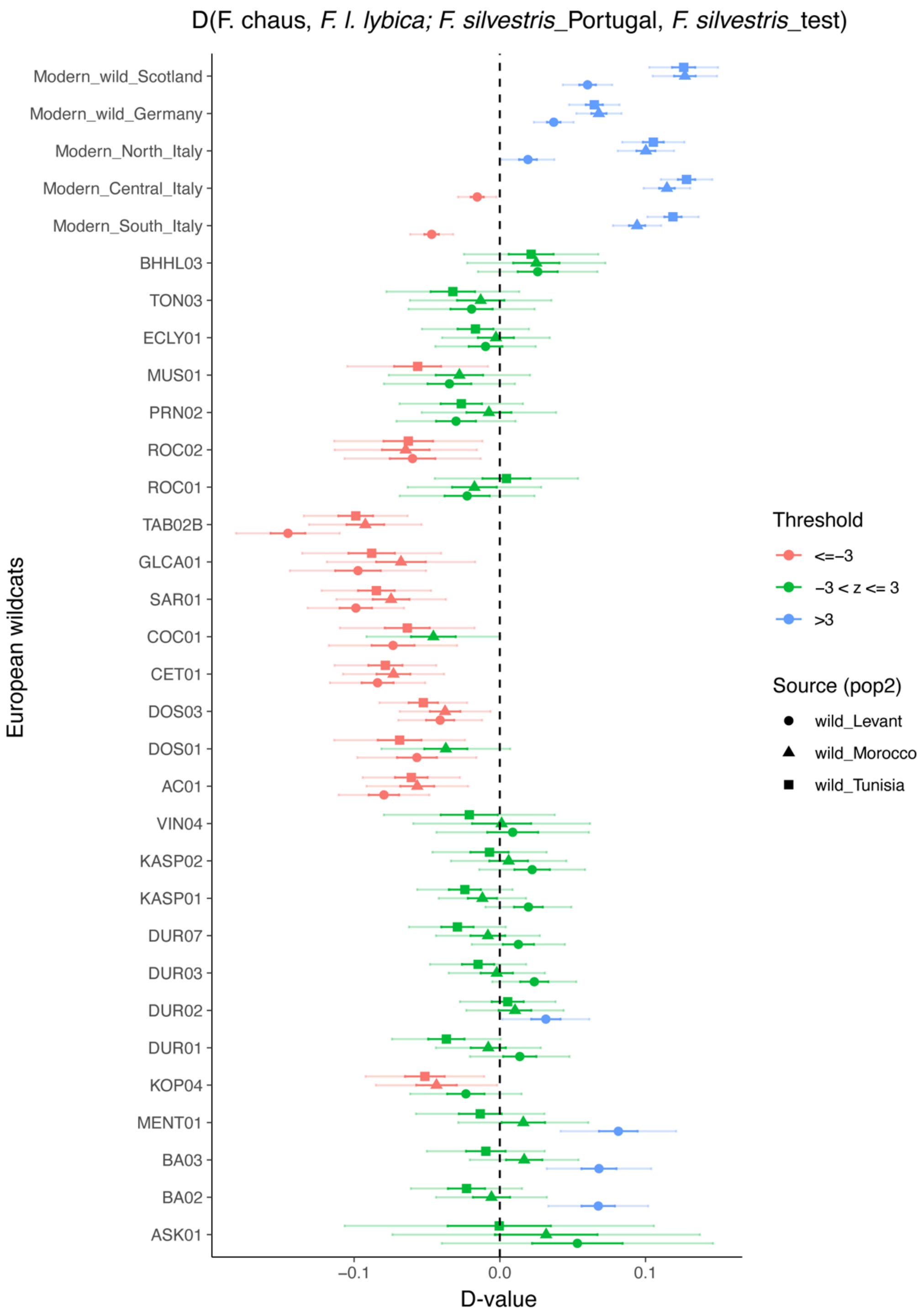

**Fig. S17.** ***D*** **statistics to test gene flow from** ***F. l. lybica*** **into** ***F. silvestris*****.** Individuals resulting positive to gene flow from *F. l. lybica* wildcats (z-scores higher than 3) are indicated by blue dots. Negative and significant z-scores (red dots) indicate gene flow from *F. l. lybica/F. catus* into the modern Portuguese wildcat (z-scores higher than -3). Green dots indicate non-significant gene flow. One standard error bar (darker) and three standard error bars (lighter) are plotted for each point. Shapes indicate the source used, as illustrated in the legend.

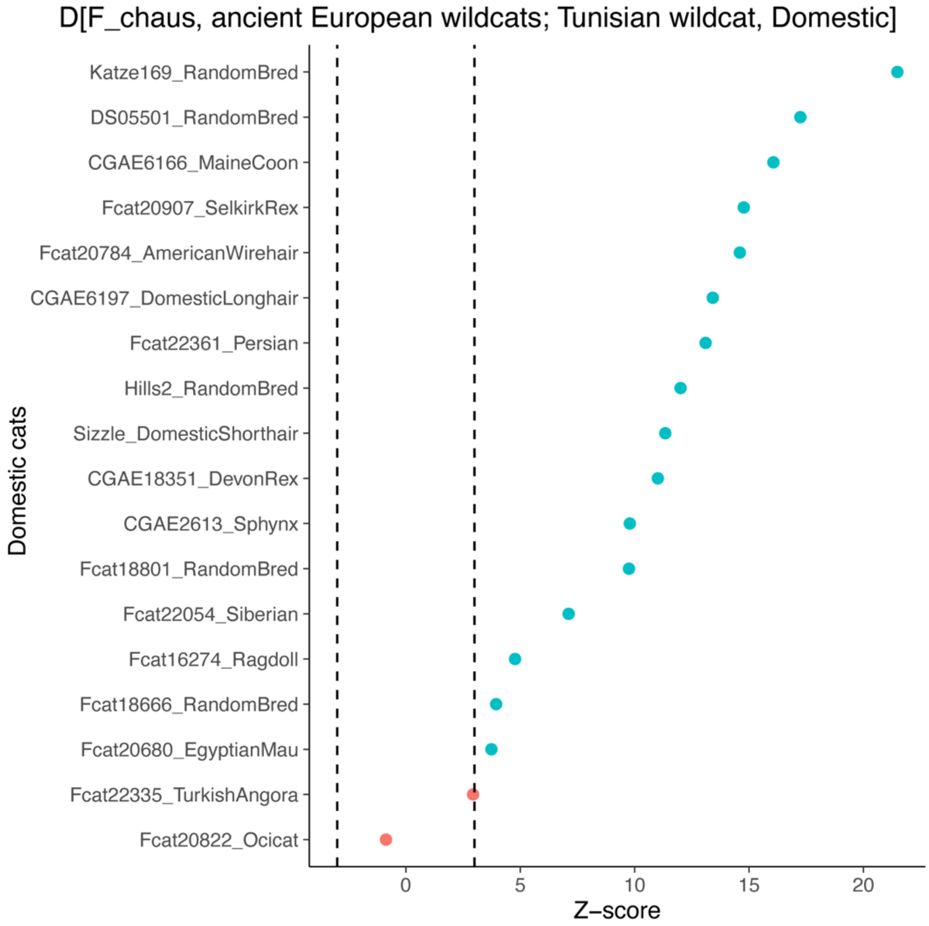

**Fig. S18. *D* statistics to test gene-flow from European wildcats into modern domestic cats.** Dashed lines indicate significance threshold of +/- 3. Individuals resulting positive to gene flow from European wildcats (z-scores higher than 3) are indicated by blue dots. Samples showing non-significant z-scores are indicated by red dots.

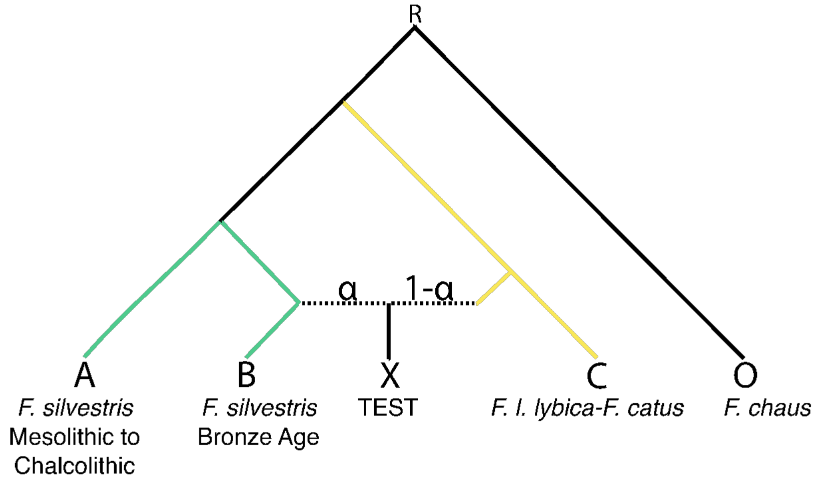

**Fig. S19. $f_4$-ratio phylogenetic model.** The tree model was used to compute admixture proportions in European wildcats and domestic cats and was designed following the nuclear phylogeny shown in Fig. 1B and Fig. S1, which placed *F. l. lybica*/*F. catus* closer to the tree root than *F. silvestris*.

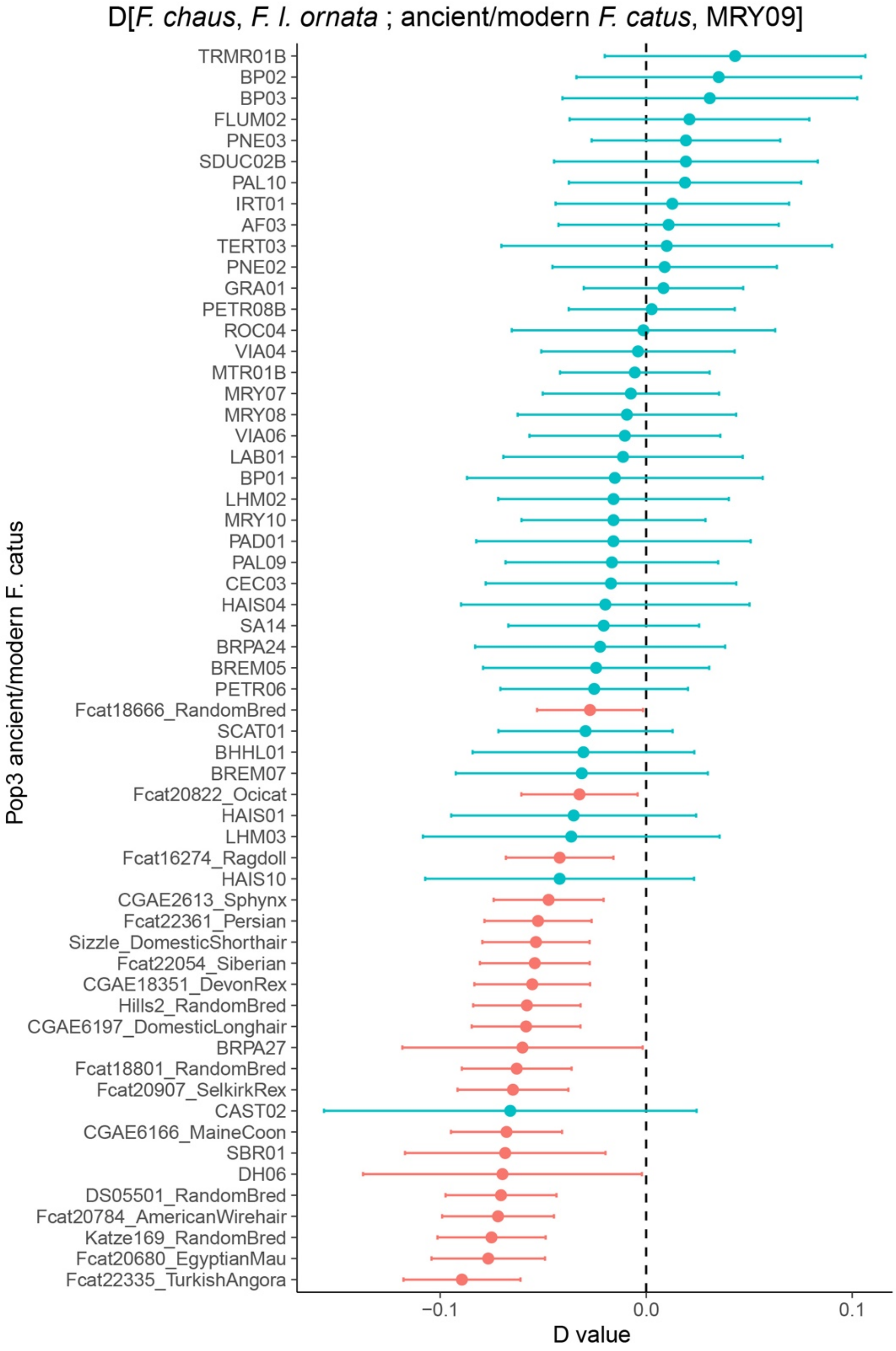

**Fig. S20. *D* statistics to test gene-flow from *F. l. ornata* into MRY09.** We tested whether MRY09 carries an excess of *F. l. ornata* alleles compared to all the other ancient and modern domestic cats. No positive significant D values were found (z-scores higher than 3) meaning that MRY09 did not show any signal of gene flow with *F. l. ornata* with. Negative and significant z-scores (red dots) indicate gene flow from *F. l. ornata* into the ancient or modern domestic cats to which MRY09 was compared. Three standard error bars are plotted for each point.

24. S. Doherty, M. Krajcarz, A. Carmagnini, E. Dimopoulos, A. Jamieson, J. M. Alves, A. Bălăşescu, P. Baker, P. Baranowski, F. Beglane, J. Bitz-Thorsen, A. Bolton, W. Bowden, H. Britton, B. De Cupere, M. De Martino, B. Donnelly-Symes, A. Evin, T. Fowler, M. Fulford, A. B. Gotfredsen, Y. Han, S.-J. Luo, I. Mainland, D. Maričević, S. Moody, T. Nesnídalová, T. O'Connor, D. Orton, J. Peters, F. Sadebeck, A. B. Siegenthaler, R. Smallman, Z. Sůvová,

R. Symmons, J. Sudds, G. Thomas, W. Van Neer, J. Wilczyński, C. Ottoni, D. Popović, L. A. F. Frantz, G. Larson, N. Sykes, The recent domestication and European dispersal of the cat. (in review, preprint available in bioRxiv).

33. C. Detry, "Paleoecología e paleoeconomía do baixo tejo no mesolítico final: o contributo do estudo dos mamíferos dos concheiros de muge," thesis, Universidad de Salamanca (2007).

34. J. Crezzini, F. Boschin, P. Boscato, U. Wierer, Wild cats and cut marks: Exploitation of Felis silvestris in the Mesolithic of Galgenbühel/Dos de la Forca (South Tyrol, Italy). *Quat. Int.* **330**, 52–60 (2014).

35. N. J. Overton, More than skin deep: Reconsidering isolated remains of 'fur-bearing species' in the British and European Mesolithic. *Camb. Archaeol. J.* **26**, 561–578 (2016).

36. R. Miccichè, P. Valenti, A. D. Messina, L. Sìneo, N. Yamaguchi, G. Catalano, S. Reale, M. Cosenza, Cat taming in the western Mediterranean. Issues, problematics and unpredictability in the light of bio-archaeological approaches to a museum specimen. *Conservation Science in Cultural Heritage* **15**, 13–25 (2017).

37. N. Spassov, D. Raychev, Late Wurm *Panthera pardus* remains from Bulgaria: the European fossil leopards and the question of the probable species survival until the Holocene. *Hist. Nat. Bulg.* **7**, 71–96 (1997).

38. S. Moscati, *The Phoenicians* (Bompiani, 1988).

39. Y. Le Bohec, "The 'third Punic war': The siege of Carthage (148–146 BC)" in *A Companion to the Punic Wars*, D. Hoyos, Eds. (Wiley Online Library, 2011), pp. 430–445.

40. P. Van Dommelen, Punic persistence: colonialism and cultural identities in Roman Sardinia. *Cultural identity in the Roman Empire* (2002).

41. J. B. Bury, *A History of the Roman Empire: From Its Foundation to the Death of Marcus Aurelius (27 B.C. – 180 A.D.)* (E-Artnow, 2019).

42. J. Peters, *Römische Tierhaltung und Tierzucht. Eine Synthese aus archäozoologischer Untersuchung und schriftlich-bildlicher Überlieferung, Passauer Universitätsschriften zur Archäologie 5* (Verlag Marie Leidorf, Rahden / Westfalen, 1998).

43. G. K. Kunst, "Tierreste aus ausgewählten Befunden der Grabungen 1997-1999 im Vicus Ost von Mautern a. d. Donau" in *Forschungen im Vicus Ost von Mautern-Favianis*, S. Groh, H. Sedlmayer, Eds. (Austrian Academy of Sciences ÖAW, 2006), pp. 636–708.

44. A. Riedel, Die Tierknochenfunde des römerzeitlichen Lagervicus von Traismauer/Augustiana in Niederösterreich. *Ann. Naturhist. Mus. Wien. Ser. A Mineral. Petrogr. Geol. Palaontol. Anthropol. Prähist.* **95**, 179–294 (1991).

45. S. Vuković-Bogdanović, Meat diet at the Upper Moesian limes: Archaeozoological evidences from the city of Viminacium and its surroundings. *Vivere Militare est. From Populus to Emperors – living on the Frontier* **II**, 269–310 (2018).

46. E.-M. Geigl, T. Grange, "Of cats and men: Ancient DNA reveals how the cat conquered the ancient world" in *Population Genomics* (Springer International Publishing, Cham, 2018), pp. 307–324.

47. D. Hoyos, *The Carthaginians* (Routledge, London, England, 2010) *Peoples of the Ancient World*.

48. B. Quillard, Les pendentifs porte-amulettes puniques: une mise à jour. Antiquités africaines. *L'Afrique du Nord de la protohistoire à la conquête arabe*, 5–60 (2023).

49. D. W. Engels, *Classical Cats* (Routledge, London, England, 2018).

50. J. Best, S. Doherty, I. Armit, Z. Boev, L. Büster, B. Cunliffe, A. Foster, B. Frimet, S. Hamilton-Dyer, T. Higham, O. Lebrasseur, H. Miller, J. Peters, M. Seigle, C. Skelton, R. Symmons, R. Thomas, A. Trentacoste, M. Maltby, G. Larson, N. Sykes, Redefining the timing and circumstances of the chicken's introduction to Europe and north-west Africa. *Antiquity* **96**, 868–882 (2022).

51. J. Peters, O. Lebrasseur, E. K. Irving-Pease, P. D. Paxinos, J. Best, R. Smallman, C. Callou, A. Gardeisen, S. Trixl, L. Frantz, N. Sykes, D. Q. Fuller, G. Larson, The biocultural origins and dispersal of domestic chickens. *Proc. Natl. Acad. Sci. U. S. A.* **119**, e2121978119 (2022).

52. K. H. Baker, H. Miller, S. Doherty, H. W. I. Gray, J. Daujat, C. Çakırlar, N. Spassov, K. Trantalidou, R. Madgwick, A. L. Lamb, C. Ameen, L. Atici, P. Baker, F. Beglane, H. Benkert, R. Bendrey, A. Binois-Roman, R. F. Carden, A. Curci, B. De Cupere, C. Detry, E. Gál, C. Genies, G. K. Kunst, R. Liddiard, R. Nicholson, S. Perdikaris, J. Peters, F. Pigière, A. G. Pluskowski, P. Sadler, S. Sicard, L. Strid, J. Sudds, R. Symmons, K. Tardio, A. Valenzuela, M. van Veen, S. Vuković, J. Weinstock, B. Wilkens, R. J. A. Wilson, J. A. Evans, A. R. Hoelzel, N. Sykes, The 10,000-year biocultural history of fallow deer and its implications for conservation policy. *Proc. Natl. Acad. Sci. U. S. A.* **121**, e2310051121 (2024).

59. L. Gourichon, D. Helmer, Etude de la faune néolithique de Menteşe (Turquie). *Life and death in a prehistoric settlement* (2008).

60. B. De Cupere, *Animals at Ancient Sagalassos. Evidence of the Faunal Remains (Studies in Eastern Mediterranean Archaeology IV)* (Brepols, Turnhout, 2001).

61. B. T. Fuller, B. De Cupere, E. Marinova, W. Van Neer, M. Waelkens, M. P. Richards, Isotopic reconstruction of human diet and animal husbandry practices during the Classical-Hellenistic, imperial, and Byzantine periods at Sagalassos, Turkey. *Am. J. Phys. Anthropol.* **149**, 157–171 (2012).

62. H. Rauh, “Knochenfunde von Säugetieren aus dem Demircihüyük (Nordwestanatolien),” thesis, Munich (1981).

76. V. Jolivet, Civita Musarna tra passato, presente e futuro. *Fasti Online* (2013).

77. H. Broise, V. Jolivet, P. Petitti, F. Catalli, A. Tagliacozzo, “Musarna (Viterbe)” in *Mélanges de l'école Française de Rome, Antiquité T. 102, N°1.* (1990), pp. 477–480.

78. C. Abatino, T. Scarano, I reperti faunistici di Torre Santa Caterina, Nardò (Lecce), University of Salento (2019).

79. B. Wilkens, G. Carenti, *Archeozoologia: Il Mediterraneo, La Storia, La Sardegna* (Editrice Democratica Sarda, 2012).

80. M. L. Bianco, R. Gregnanin, R. Caimi, J. Manning Press, 1997-Lo scavo urbano pluristratificato di via C. Battisti 132 a Padova. *Archeologia Veneta, XIX-XX*, 7–150 (1996).

81. G. Soranna, “Analisi di un campione faunistico proveniente dal Colle Palatino - Roma (Italia)” in *Atti 8° Convegno Nazionale Di Archeozoologia (Lecce, 2015),* (2019), pp. 171–181.

82. L. Conte, A. K. Ingoglia, A. M. Lucena Martìn, L. Riolo, Il culto dell'acqua e la civiltà dei fossati nel territorio di Partanna (Trapani). *Ann. Online, Sez. Museol. Sci. Nat.* (2007).

83. S. Tusa, S. M. Puglisi, *La Sicilia Nella Preistoria* (Sellerio, 1983).

91. S. Radbauer, G. K. Kunst, "Case study: Earlier Studies Done Well and What We Can Learn – Carnuntum" in *Roman Toilets. Their Archaeology and Cultural History* (ucrisportal.univie.ac.at, 2011).

92. G. K. Kunst, N. I. Kirchengast, "Street, backyard, baths-urban bonescapes from the Roman civilian city of Carnuntum (Lower Austria)" in *Über Speisen, Getränke und Macht zwischen Spätantike und Karolingerzeit: Akten der 29. und 30. Internationalen Symposien "Grundprobleme Der frühgeschichtlichen Entwicklung im mittleren Donauraum" in Mistelbach, 8.-11. November 2017 und in Brünn, 14.-16. November 2018* (Verlag Bernhard Albert Greiner, 2022), pp. 149–163.

93. E. Pucher, Ed., "Der frühneuzeitliche Knochenabfall eines Wirtshauses neben der Salzburger Residenz" in *Jahresschrift Des Salzburger Museum Carolino Augusteum 35/36 (1989/90)* (Salzburg, 1991), pp. 71–135.

94. Küchelmann, H. C. 2014a. Mit Knochen gepflastert – Tierknochenfunde vom Bremer Marktplatz (Fundstelle 201-Altstadt 2002). Schriften des Naturwissenschaftlichen Vereins für Schleswig-Holstein 73, 23–64.

95. Küchelmann, H.C. 2014b. Frühneuzeitliche Tierknochen aus dem Bremer Stadtgraben, Grabung 253-Altstadt 2011, Am Wall. Bericht für die Landesarchäologie Bremen, Bremen.

96. F. Johansson, H. Hüster, *Untersuchungen an Skelettresten von Katzen aus Haithabu (Ausgrabung 1966-1969) Berichte über die Ausgrabungen in Haihtabu 24* (Neumünster, 1987).

97. Sichert, B. M., Wilk, A. & Stephan, E. (in preparation). Die Tierknochenfunde aus der früh- bis hochmittelalterlichen Siedlung Lauchheim-Mittelhofen (Ostalbkreis). Forschungen und Berichte zur Archäologie in Baden-Württemberg.

98. C. Tejedor-Rodríguez, M. Moreno-García, C. Tornero, A. Hoffmann, Í. García-Martínez de Lagrán, H. Arcusa-Magallón, R. Garrido-Pena, J. I. Royo-Guillén, S. Díaz-Navarro, L. Peña-Chocarro, K. W. Alt, M. Rojo-Guerra, Investigating Neolithic caprine husbandry in the Central Pyrenees: Insights from a multi-proxy study at Els Trocs cave (Bisaurri, Spain). *PLoS One* **16**, e0244139 (2021).

99. J. A. Quirós Castillo, Castillo de Labastida. *Arkeoikuska 13. Eusko Jaurlaritza/Gobierno Vasco. Vitoria-Gasteiz*, 83–87 (2014).

100. A. von den Driesch, A. Morales, Los restos animales del yacimiento de Terrera Ventura (Tabernas, Almería). *Cuadernas de Prehistoria et Arqueología* **4**, 15–34 (1977).

101. H. Manhart, Die vorgeschichtliche Tierwelt von Koprivec und Durankulak und anderen prähistorischen Fundplätzen in Bulgarien aufgrund von Knochenfunden aus archäologischen Ausgrabungen: Dissertation der Fakultät für Biologie der Ludwig-Maximilians-Universität München. (1998).

102. C. Detry, Medieval and postmedieval fauna from the Palmela Castle (Setúbal, Portugal). *(submitted to International Journal of Osteoarchaeology)*.

103. J. Timmermans, Y. Devos, V. Ghesquière, C. B. De, K. Deforce, L. Speleers, Van Bellingen S., Brucity/parking 58: Un ancien quai de Senne (XVe siècle)(RBC/BHG). *Archaeologia Mediaevalis* **43**, 80–81 (2020).

104. B. De Cupere, N. De Winter, J. Steegmans, W. Vanaenrode, E. Wesemael, Allemaal beestjes. Studie van het dierljk bot uit een Romeinse waterput van de site Tongeren-Oost. *Tongerse Annalen*, 68–77 (2022).

105. L. Pastor, T. Oueslati, Écly – Les Septiers (Ardennes): un établissement rural d'époque romaine à l'identité cachée au fond du puits. In: Calbris M., Sadourny E., Brkojewitsch G., Bollard-Raineau, I. (eds), XVIe colloque de l'association AGER: Les marqueurs des statuts sociaux dans les campagnes romaines des Gaules et de leurs périphéries, Nantes, La Manufacture, 13-16 novembre 2024: pré-actes. (2024).

106. M. A. Dowd, Middle and late bronze age ritual activity at glencurran cave, co. Clare. *From Bann Flakes to Bushmills: Papers in Honour of Professor Peter Woodman*, 89–100 (2009).

107. H. Friedl, Tierknochenfunde aus Kassope/Griechenland (4.-1. Jh. v. Chr.). Dissertation (Ludwig-Maximilians-Universität München, Institut für Paläoanatomie, Domestikationsforschung und Geschichte der Tiermedizin, München). (1984).

108. M. Boudin, M. Van Strydonck, T. van den Brande, H.-A. Synal, L. Wacker, RICH – A new AMS facility at the Royal Institute for Cultural Heritage, Brussels, Belgium. *Nucl. Instrum. Methods Phys. Res. B* **361**, 120–123 (2015).

109. M. Wojcieszak, T. Van den Brande, G. Ligovich, M. Boudin, Pretreatment protocols performed at the royal institute for Cultural Heritage (rich) prior to ams 14C measurements. *Radiocarbon* **62**, e14–e24 (2020).

110. P. J. Reimer, W. E. N. Austin, E. Bard, A. Bayliss, P. G. Blackwell, C. Bronk Ramsey, M. Butzin, H. Cheng, R. L. Edwards, M. Friedrich, P. M. Grootes, T. P. Guilderson, I. Hajdas, T. J. Heaton, A. G. Hogg, K. A. Hughen, B. Kromer, S. W. Manning, R. Muscheler, J. G. Palmer, C. Pearson, J. van der Plicht, R. W. Reimer, D. A. Richards, E. M. Scott, J. R. Southon, C. S. M. Turney, L. Wacker, F. Adolphi, U. Büntgen, M. Capano, S. M. Fahrni, A. Fogtmann-Schulz, R. Friedrich, P. Köhler, S. Kudsk, F. Miyake, J. Olsen, F. Reinig, M. Sakamoto, A. Sookdeo, S. Talamo, The IntCal20 Northern hemisphere radiocarbon age calibration curve (0–55 cal kBP). *Radiocarbon* **62**, 725–757 (2020).

120. H. Li, Aligning sequence reads, clone sequences and assembly contigs with BWA-MEM. 2013. (2013).

121. P. Danecek, J. K. Bonfield, J. Liddle, J. Marshall, V. Ohan, M. O. Pollard, A. Whitwham, T. Keane, S. A. McCarthy, R. M. Davies, H. Li, Twelve years of SAMtools and BCFtools. *Gigascience* **10** (2021).

122. A. McKenna, M. Hanna, E. Banks, A. Sivachenko, K. Cibulskis, A. Kernytsky, K. Garimella, D. Altshuler, S. Gabriel, M. Daly, M. A. DePristo, The Genome Analysis Toolkit: a MapReduce framework for analyzing next-generation DNA sequencing data. *Genome Res.* **20**, 1297–1303 (2010).

123. A. Peltzer, *DeDup: A Merged Read Deduplication Tool Capable to Perform Merged Read Deduplication on Single End Data* (Github; https://github.com/apeltzer/DeDup).

135. E. M. Ortiz, *Vcf2phylip: Convert SNPs in VCF Format to PHYLIP, NEXUS, Binary NEXUS, or FASTA Alignments for Phylogenetic Analysis* (Github; https://github.com/edgardomortiz/vcf2phylip).
