## Supplementary datasets for "The dispersal of domestic cats from Northern Africa and their introduction to Europe over the last two millennia"

**Supplementary table 1.** List of archaeological samples analysed in this study.

| ID | ID-Lab | Individual | Site | Country | Latitude | Longitude | Material sampled WVN edited | Age criteria* | Measurements (mm)** | Chronology / C14 date | Cultural time bin Fig. 1 | mtDNA | PCA Fig. 2 cluster | Collector |
|---|---|---|---|---|---|---|---|---|---|---|---|---|---|---|
| BHHLcat01 | BHHL01 | cat01 | Bernhardsthal | Austria | 48.69 | 16.87 | left tibia shaft | prox NF, dist NF | GL without epiphyses 95 | 770AD (95.4%) 980AD | post-Roman | Felis l. lybica/Felis catus clade IV-D | Felis l. lybica/Felis catus | Konstantina Saliari |
| BHHLcat03 | BHHL03 | cat02? | Bernhardsthal | Austria | 48.69 | 16.87 | right radius, distal | dist F | SD 6.6; Bd 14.9 | Roman | post-Roman | Felis silvestris clade I | Felis silvestris | Konstantina Saliari |
| HTBcat01 | HTB01 | cat01 | Hinterbrühl | Austria | 48.08 | 16.23 | left radius | prox F, dist F | GL 121.1; Bp 9.4; Bd 14.3 | Holocene unspecified | | | | Günther Karl Kunst |
| MTRcat01 | MTR01B | cat01 | Mautern - vicus Ost | Austria | 48.39 | 15.58 | right mandible with C, P3, P4 & M1 | dentition fully erupted | (1) 66.0; (2) 61.7; (3) 58.7; (4) 54.5; (5) 19. | 50BC (95.4%) 80AD | Antiquity/Roman | Felis l. lybica/Felis catus clade IV-C | Felis l. lybica/Felis catus | Günther Karl Kunst |
| NBLcat01 | NBL01 | cat01 | Nebellöcher, Baden district | Austria | 47.97 | 16.21 | left humerus | prox fusing, dist F | | Holocene unspecified, prob. medieval/modern | | | | Günther Karl Kunst |
| NBLcat02 | NBL02 | cat02 | Nebellöcher, Baden district | Austria | 47.97 | 16.21 | left humerus | prox F, dist F | GL 94.1; Dp 19.5; SD 6.7; Bd 17.6 | Holocene unspecified, prob. medieval/modern | | | | Günther Karl Kunst |
| NBLcat03 | NBL03 | cat03 | Nebellöcher, Baden district | Austria | 47.97 | 16.21 | left humerus | prox F, dist F | GL 90.5; Dp 19.2; SD 7.2; Bd 17.2 | Holocene unspecified, prob. medieval/modern | | | | Günther Karl Kunst |
| NBLcat04 | NBL04 | cat04 | Nebellöcher, Baden district | Austria | 47.97 | 16.21 | right calcaneus | prox F | GL 28.9 | Holocene unspecified, prob. medieval/modern | | | | Günther Karl Kunst |
| NBLcat05 | NBL05B | cat05 | Nebellöcher, Baden district | Austria | 47.97 | 16.21 | right mandible with C, P3 & P4 | dentition fully erupted | (1) 59.8; (2) 56.0; (3) 51.9; (4) 48.8; (5) 20.2 | Holocene unspecified, prob. medieval/modern | | | | Günther Karl Kunst |
| PETRcat03 | PETR03 | cat01 | Petronell-Carnuntum | Austria | 48.12 | 16.85 | right humerus, distal | dist F | SD 7.7; Bd 20.7 | Roman | | | | Günther Karl Kunst |
| PETRcat04 | PETR04 | cat02 | Petronell-Carnuntum | Austria | 48.12 | 16.85 | right radius, proximal | prox F | Bp 8.4 | Roman | | | | Günther Karl Kunst |
| PETRcat06 | PETR06 | cat03 | Petronell-Carnuntum | Austria | 48.12 | 16.85 | left femur, proximal | prox F | Bp 21.2; DC 9.4; SD 7.9 | 120AD (87.6%) 260AD / 290AD (7.8%) 320AD | Antiquity/Roman | Felis l. lybica/Felis catus clade IV-C | Felis l. lybica/Felis catus | Günther Karl Kunst |
| PETRcat07 | PETR07B | cat04 | Petronell-Carnuntum | Austria | 48.12 | 16.85 | left mandible with C, P3, P4 & M1 | dentition fully erupted | (1) 61.4; (2) 57.5; (3) 53.8; (4) 50.1; (5) 19.4 | Roman | | | | Günther Karl Kunst |
| PETRcat08 | PETR08B | cat05 | Petronell-Carnuntum | Austria | 48.12 | 16.85 | left mandible with C, P3, P4 & M1 | dentition fully erupted | (1) 70.4; (2) 66.5; (3) 62; (4) 58.4; (5) 20.6; | 20AD ( 2.6%) 50AD / 60AD (92.8%) 210AD | Antiquity/Roman | Felis l. lybica/Felis catus clade IV-C | Felis l. lybica/Felis catus | Günther Karl Kunst |
| SBRcat01 | SBR01 | cat01 | Salzburg - Residenz | Austria | 47.80 | 13.05 | left humerus | prox F, dist F | GL 103.3; Dp 20.2; SD 7.9; Bd 18.5 | 15/16th centuries | post-Roman | Felis l. lybica/Felis catus clade IV-A2 | Felis l. lybica/Felis catus | Günther Karl Kunst |
| TRMRcat01 | TRMR01B | cat01 | Traismauer | Austria | 48.35 | 15.74 | left mandible with I1, P3 & M1 | dentition fully erupted | (1) 58.2; (2) 54.4; (3) 50.7; (4) 47.3; (5) 20.1 | 60AD (95.4%) 220AD | Antiquity/Roman | Felis l. lybica/Felis catus clade IV-C | Felis l. lybica/Felis catus | Konstantina Saliari |
| TRMRcat02 | TRMR02 | cat01? | Traismauer | Austria | 48.35 | 15.74 | right humerus, proximal | prox NF | | Roman | | | | Konstantina Saliari |
| BRPAcat01 | BRPA01 | cat01 | Parking 58, Brussels | Belgium | 50.85 | 4.35 | left tibia | prox F, dist F | GL 102.6; Bp 18.6; SD 6.0; Bd 13.8; Dd 9.4 | end 14th - 15th cent. AD | | | | Bea De Cupere |
| BRPAcat16 | BRPA16 | cat16 | Parking 58, Brussels | Belgium | 50.85 | 4.35 | left tibia | prox fusing, dist F | GL 106.6; Bp 16.9; SD 6.2; Bd 12.5; Dd 8.2 | 12th - 13th cent. AD | | | | Bea De Cupere |
| BRPAcat22 | BRPA22 | cat02B | Parking 58, Brussels | Belgium | 50.85 | 4.35 | left mandible with P3 and M1 in situ | dentition fully erupted | (1) 49.9; (2) 46.7); (3) 43.9; (4) 40.4); (5) 18 | end 14th - 15th cent. AD | | | | Bea De Cupere |
| BRPAcat24 | BRPA24 | cat04B | Parking 58, Brussels | Belgium | 50.85 | 4.35 | left mandible with P3 and M1 in situ | dentition fully erupted | (1) 51.7; (2) 48.2; (3) 45.3; (4) 41.6; (5) 18.5 | end 14th - 15th cent. AD | post-Roman | Felis l. lybica/Felis catus clade IV-C | Felis l. lybica/Felis catus | Bea De Cupere |
| BRPAcat27 | BRPA27 | cat07B | Parking 58, Brussels | Belgium | 50.85 | 4.35 | left mandible with P3 and M1 in situ | dentition fully erupted | 1) 47.4; (2) 45.1; (3) 41.8; (4) 39.7; (5) 17.2 | end 14th - 15th cent. AD | post-Roman | Felis l. lybica/Felis catus clade IV-C | Felis l. lybica/Felis catus | Bea De Cupere |
| TONcat03 | TON03 | cat02 | Tongeren, Industrie Oost | Belgium | 50.78 | 5.46 | os petrosum | adult | | 40BC (4.0%) 10BC / AD (91.4%) 130AD | Antiquity/Roman | Felis l. lybica/Felis catus clade IV-A0 | Felis silvestris | Bea De Cupere |
| BUD01 | BUD01 | cat01 | Budjaka (near Sozopol) | Bulgaria | 42.41 | 27.71 | shaft fragments of ulna and of humerus | -; cf. adult | | Late Neolithic | | | | Nicolai Spassov |
| DOLcat01 | DOL01 | cat01 | Dolnoslav | Bulgaria | 41.93 | 24.98 | humerus, distal | dist F | | 4510-4330 BC (cal AMS) | | | | Nicolai Spassov |
| DOLcat02 | DOL02 | cat01 | Dolnoslav | Bulgaria | 41.93 | 24.98 | right ulna, proximal | prox F | | Chalcolithic | | | | Nicolai Spassov |
| DOLcat03 | DOL03 | cat02 | Dolnoslav | Bulgaria | 41.93 | 24.98 | left maxilla with C (root), P3, P4 & M1 | dentition fully erupted | | Chalcolithic | | | | Nicolai Spassov |
| DURcat01 | DUR01 | cat01 | Durankulak | Bulgaria | 43.70 | 28.54 | right mandible with C, P3, P4 & M1 | dentition fully erupted | (1) 70.6; (2) (68); (3) 62; (4) (59); (5) 22.7; ( | 4450BC (95.4%) 4330BC | Neolithic/Chalcolithic | Felis l. lybica/Felis catus clade IV-A1 | Felis silvestris | Joris Peters, Bea De Cupere |
| DURcat02 | DUR02 | cat02 | Durankulak | Bulgaria | 43.70 | 28.54 | right mandible with C, P3, P4 & M1 | dentition fully erupted | (5) 23.7); (6) L 9.6; (7) 9.4; (9) 11.3; (10) 11 | Aeneolithic 4200-4100 cal BC | Neolithic/Chalcolithic | Felis l. lybica/Felis catus clade IV-A1 | Felis silvestris | Joris Peters, Bea De Cupere |
| DURcat03 | DUR03 | cat03 | Durankulak | Bulgaria | 43.70 | 28.54 | left mandible with P3 & P4 | young, M1 not yet erupted | | Aeneolithic 4200-4100 cal BC | Neolithic/Chalcolithic | Felis l. lybica/Felis catus clade IV-A1 | Felis silvestris | Joris Peters, Bea De Cupere |
| DURcat06 | DUR06 | cat04 | Durankulak | Bulgaria | 43.70 | 28.54 | right femur | dist fusing | GL 86.2; Dp 17.2; SD 5.8 | Aeneolithic 4200-4100 cal BC | | | | Joris Peters, Bea De Cupere |
| DURcat07 | DUR07 | cat05 | Durankulak | Bulgaria | 43.70 | 28.54 | left metatarsus III | dist F | GL 61.4; Bd 6.5 | Aeneolithic 4200-4100 cal BC | Neolithic/Chalcolithic | Felis silvestris clade I | Felis silvestris | Joris Peters, Bea De Cupere |
| KOPcat02 | KOP02 | cat02 | Koprivec | Bulgaria | 43.41 | 25.88 | left pelvis | all fused; adult | GL(96); LA 12.9; SH 13.9; SB 5.1 | Neolithic B 5800-4800 cal BC | | | | Joris Peters, Bea De Cupere |
| KOPcat03 | KOP03 | cat03 | Koprivec | Bulgaria | 43.41 | 25.88 | left ulna | prox F, dist F | GL 136; BPc 10.0; DPA 12.8; SDO 11.1 | 6050-5900 BC (cal AMS) | | | | Joris Peters, Bea De Cupere |
| KOPcat04 | KOP04 | cat04 | Koprivec | Bulgaria | 43.41 | 25.88 | right pelvis | all fused; adult | SH 13.3; SB 4.8 | 5480BC (95.4%) 5360BC | Neolithic/Chalcolithic | Felis l. lybica/Felis catus clade IV-A1 | Felis silvestris | Joris Peters, Bea De Cupere |
| BG07 | BG07 | modern wild cat0 | #N/A | Bulgaria | #N/A | | | | | modern (museum collection) | | Felis l. lybica/Felis catus clade IV-A1 | Felis silvestris | Nicolai Spassov |
| BG01 | BG01 | modern wild cat0 | #N/A | Bulgaria | #N/A | | | | | modern (museum collection) | | | | Nicolai Spassov |
| BG5 | BG5 | modern wild cat0 | #N/A | Bulgaria | #N/A | | | | | modern (museum collection) | | Felis l. lybica/Felis catus clade IV-C | Felis silvestris | Nicolai Spassov |
| BG16 | BG16 | modern wild cat1 | #N/A | Bulgaria | #N/A | | | | | modern (museum collection) | | Felis silvestris clade I | Felis silvestris | Nicolai Spassov |
| BG2 | BG2 | modern wild cat0 | #N/A | Bulgaria | #N/A | | | | | modern (museum collection) | | | | Nicolai Spassov |
| ECLYcat01 | ECLY01 | cat01 | Ecly | France | 4.28 | 49.54 | left mandible with M1 | dentition fully erupted | (6) 8.8/3.6; (7) 8.7; (9) 10.4; (10) 10.7 | 20AD (2.1%) 50AD / 60AD (93.3%) 210AD | Antiquity/Roman | Felis silvestris clade I | Felis silvestris | Tarek Oueslati |
| BREMcat05 | BREM05 | cat04 | Bremen; 201-Altstadt, Marktplatz | Germany | 53.10 | 8.83 | left femur | prox F, dist F | GL 95.3; Bp 19.5; DC 9.3; SD 7.8; Bd 17.2 | 1440AD (65.0%) 1530AD / 1570AD (30.4%) 1630AD | post-Roman | Felis l. lybica/Felis catus clade IV-C | Felis l. lybica/Felis catus | Hans Christian Küchelmann |
| BREMcat07 | BREM07 | cat06 | Bremen; 253-Altstadt, Am Wall | Germany | 53.10 | 8.83 | right femur | prox fusing, dist NF | GL without epiphyses 96.1 | 16th | post-Roman | Felis l. lybica/Felis catus clade IV-C | Felis l. lybica/Felis catus | Hans Christian Küchelmann |
| CLWHcat01 | CLWH01 | cat01 | Calw-Hirsau | Germany | 48.74 | 8.74 | left humerus | prox F, dist F | GL 85.4; Dp 17.0; SC 5.7; Bd 15.4 | 12th/13rd century CE | | | | Simon Trixl |
| CLWHcat02 | CLWH02 | cat02 | Calw-Hirsau | Germany | 48.74 | 8.74 | left humerus | prox NF, dist F | SC 6.0; Bd 15.3 | 12th/13rd century CE | | | | Simon Trixl |
| HAIScat01 | HAIS01 | cat01 | Haithabu, settlement area | Germany | 9.56 | 54.50 | right mandible with C, P3, P4 & M1 | dentition fully erupted | (1) 53.3; (2) 51.2; (3) 48.7; (4) 45.9; (5) 18.4 | 800-1050 AD | post-Roman | Felis l. lybica/Felis catus clade IV-C | Felis l. lybica/Felis catus | Ulrich Schmölke |
| HAIScat03 | HAIS03 | cat03 | Haithabu, settlement area | Germany | 9.56 | 54.50 | right mandible with C, P3, P4 & M1 | dentition fully erupted | (1) 54.3; (2) 51.6; (3) 48.4; (4) 45.4; (5) 17.8 | 800-1050 AD | | | | Ulrich Schmölke |
| HAIScat04 | HAIS04 | cat04 | Haithabu, settlement area | Germany | 9.56 | 54.50 | right mandible with P3, P4 & M1 | dentition fully erupted | (1) 54.0; (2) 50.7; (3) 47.7; (4) 44.2; (5) 17.7 | 800-1050 AD | post-Roman | Felis l. lybica/Felis catus clade IV-C | Felis l. lybica/Felis catus | Ulrich Schmölke |
| HAIScat10 | HAIS10 | cat10 | Haithabu, settlement area | Germany | 9.56 | 54.50 | right mandible with P3 & P4 | dentition fully erupted | (1) 54.6; (2) 51.4; (3) 48.0; (4) 48.0; (5) 19.1 | 800-1050 AD | post-Roman | Felis l. lybica/Felis catus clade IV-C | Felis l. lybica/Felis catus | Ulrich Schmölke |
| HDRcat01 | HDR01 | cat01 | Hochdorf-Reps | Germany | 9.02 | 49.02 | right metatarsus IV | dist F | GL 54.5 | 450 to 250 BCE (La Tène Period Early) | | | | Simon Trixl |
| HDRcat02 | HDR02 | cat02 | Hochdorf-Reps | Germany | 9.02 | 49.02 | left ulna, proximal | prox F | DPA 11.5; BPC 10.0 | 450 to 250 BCE (La Tène Period Early) | | | | Simon Trixl |
| LHMcat01 | LHM01 | cat01 | Lauchheim-Mittelhofen | Germany | 10.16 | 49.13 | left radius | prox F | Bp 7.6 | 6th/7th century CE | | | | Simon Trixl |
| LHMcat02 | LHM02 | cat02 | Lauchheim-Mittelhofen | Germany | 10.16 | 49.13 | right humerus | juv., prox NF, dist NF | GL without epiphyses 47 | 8th/9th century CE | post-Roman | Felis l. lybica/Felis catus clade IV-C | Felis l. lybica/Felis catus | Simon Trixl |
| LHMcat03 | LHM03 | cat03 | Lauchheim-Mittelhofen | Germany | 10.16 | 49.13 | left ulna, proximal | prox F | SDO 7.8; DPA 8.8; BPC 6.0 | 8th/9th century CE | post-Roman | Felis l. lybica/Felis catus clade IV-A2 | Felis l. lybica/Felis catus | Simon Trixl |
| TBScat01 | TBS01 | cat01 | Tauberbischofsheim-Dittigheim | Germany | 49.61 | 9.67 | right mandible with C, P3, P4 & M1 | dentition fully erupted | (2) 64.1; (4) 56.4; (5) 22.8; (6) 9.4/3.6; (7) 9. | 26th/25th century BCE, Chalcolithic (Corded Ware Culture) | | | | Simon Trixl |
| KAcat01 | KA01 | cat01 | Kastanas | Greece | 40.82 | 22.66 | right radius, proximal | prox F | Bp 9.3; SD 6.0 | ca. 1250 BC | | | | Cornelia Becker |
| KAcat02 | KA02 | cat02 | Kastanas | Greece | 40.82 | 22.66 | right pelvis | all fused; adult | | ca. 1250 BC | | | | Cornelia Becker |
| KAcat03 | KA03B | cat03 | Kastanas | Greece | 40.82 | 22.66 | left mandible with C, P3, P4 & M1 | dentition fully erupted | | ca 200-400 BC | | | | Cornelia Becker |
| KAcat06 | KA06 | cat02 | Kastanas | Greece | 40.82 | 22.66 | right pelvis | all fused; adult | LA 14.1; SH 14.0; SB 6.2; LFo 23.4 | late phase of the Late Bronze Age (c. 1300 BC) | | | | Cornelia Becker |
| KAcat07 | KA07 | cat03 | Kastanas | Greece | 40.82 | 22.66 | left humerus, distal | dist F | SD 7.7; Bd 23.3 | transitional phase between Late Bronze Age and Iron Age (c. 1200-1100 BC) | | | | Cornelia Becker |
| KASPcat01 | KASP01 | cat01 | Kassope | Greece | 39.15 | 20.67 | left mandible with P3, P4 & M1 | dentition fully erupted | (5) 23.4; (7) 9.7; (9) 12.7; (10) 10.6 | 400BC (33.6%) 340BC / 310BC (61.8%) 200BC | Antiquity/Roman | Felis silvestris clade I | Felis silvestris | Cornelia Becker |
| KASPcat02 | KASP02 | cat02 | Kassope | Greece | 39.15 | 20.67 | left femur, distal | dist F | SD 9.3; Bd 21.8 | 4th-1st cent BC | Antiquity/Roman | Felis silvestris clade I | Felis silvestris | Cornelia Becker |
| GLCAcat01 | GLCA01 | cat01 | Glencurran Cave, Co. Clare | Ireland | -8.79 | 52.97 | right mandible, with P3 preserved | dentition fully erupted | | 4778±24 BP (UBA-45133), 3634-3526 cal BC (Neolithic) | Neolithic/Chalcolithic | Felis silvestris clade I | Felis silvestris | Marion Dowd |
| ACcat01 | AC01 | cat01 | Arene Candide (Finale Ligure, Savona, Liguria) | Italy | 44.16 | 8.33 | right metatarsus V, proximal | -; cf. adult | | 10-11 ka (Epigravettian) | Paleolithic/Mesolithic | Felis silvestris clade I | Felis silvestris | Francesca Alhaique |
| ACcat02 | AC02 | cat02 | Arene Candide (Finale Ligure, Savona, Liguria) | Italy | 44.16 | 8.33 | right metatarsus V, proximal | -; cf. adult | | 10-11 ka (Epigravettian) | | | | Francesca Alhaique |
| AFcat03 | AF03 | cat02 | Alba Fucens -Cisterna (Fucino, Abruzzi) | Italy | 42.08 | 13.41 | left pelvis | all fused; adult | LAR 12.4; SH 11.8; SB 4.9; Lfo 19.6 | 210-340 AD (cal AMS) | Antiquity/Roman | Felis l. lybica/Felis catus clade IV-C | Felis l. lybica/Felis catus | Francesca Alhaique |
| AKRcat02 | AKR02 | cat02 | Akrai, Sicily | Italy | 37.06 | 14.89 | radius shaft | dist NF, prox NF | Bp 8.1 | late 3rd – mid 4th c. AD | | | | Anna Gręzak |
| ANGcat01 | ANG01 | cat01 | Grotta S.Angelo (Ostuni, Apulia) | Italy | 40.82 | 16.70 | right pelvis | -; cf adult | | "Pleistocene" (Upper Paleolithic?) | | | | Francesca Alhaique |
| CALAcat01 | CALA01 | cat01 | Grotta della Cala (Marina di Camerota - Salerno) | Italy | 40.00 | 15.37 | right tibia, proximal | prox NF | | 8370±80 BP; 8060±100 | | | | Paolo Boscato, Francesco Boschin |
| CALAcat02 | CALA02 | cat02 | Grotta della Cala (Marina di Camerota - Salerno) | Italy | 40.00 | 15.37 | left humerus, proximal | prox F | | 8370±80 BP; 8060±100 | | | | Paolo Boscato, Francesco Boschin |
| CALcat01 | CAL01 | cat01 | Calvatone, Lombardia, Italy | Italy | 45.13 | 10.44 | left femur, proximal | prox F | Bp 23.3; DC 11.1 | Roman, Augustean | | | | Barbara Wilkens |
| CASTcat02 | CAST02 | cat01 | Castiglione in Sabina | Italy | 42.11 | 12.78 | left radius | prox F, dist F | GL 90.5; Bp 8.2; SD 6.2; Bd 12.7 | 11-12th c AD | post-Roman | Felis l. lybica/Felis catus clade IV-A2 | Felis l. lybica/Felis catus | Claudia Minniti |
| CECcat03 | CEC03 | cat03 | Rome, S. Cecilia in Trastevere | Italy | 41.89 | 12.48 | right pelvis, ilium | cf. adult | | 12-13th c AD | post-Roman | Felis l. lybica/Felis catus clade IV-C | Felis l. lybica/Felis catus | Claudia Minniti, Jacopo De Grossi Mazzorin |
| COCcat01 | COC01 | cat01 | Grotta del Cocci (Narni, Umbria) | Italy | 42.52 | 12.52 | right femur, proximal | prox F | Bp 21.7; DC 9.7 | 5370BC (95.4%) 5210BC | Neolithic/Chalcolithic | Felis silvestris clade I | Felis silvestris | Francesca Alhaique |
| DOScat01 | DOS01 | cat01 | Galgenbühel/Dos de la Forca-Salorno | Italy | 11.08 | 46.24 | right metatarsus IV, proximal | -; cf. adult | | 8240BC (95.4%) 7940BC | Paleolithic/Mesolithic | Felis silvestris clade I | Felis silvestris | Ursula Wierer |
| DOScat02 | DOS02 | cat02 | Galgenbühel/Dos de la Forca-Salorno | Italy | 11.08 | 46.24 | left calcaneus | - | | 8,500-7,500 cal BC | | | | Ursula Wierer |
| DOScat03 | DOS03 | cat03 | Galgenbühel/Dos de la Forca-Salorno | Italy | 11.08 | 46.24 | radius, distal | dist F | | 8,500-7,500 cal BC | Paleolithic/Mesolithic | Felis silvestris clade I | Felis silvestris | Ursula Wierer |
| FIDcat01 | FID01 | cat01 | Fidene | Italy | 12.52 | 42.00 | long bone fragment (calcined; completely white) | | | calcined; completely white | | | | C. Minniti & J. De Grossi Mazzorin |
| FLUMcat02 | FLUM02 | cat02 | nuraghe Flumenelongu, Sardinia | Italy | 8.26 | 40.64 | right pelvis, ilium | all fused; adult | LA 11.7 SH 10.1 | 600AD (2.7%) 620AD / 630AD (86.2%) 690AD / 740AD ( 6.6%) 780AD | Antiquity/Roman | Felis l. lybica/Felis catus clade IV-C | Felis l. lybica/Felis catus | Barbara Wilkens |
| GAOcat11 | GAO11 | cat01 | Grotta dell'Onda | Italy | 10.34 | 43.98 | left tibia, shaft | cf. (sub)adult | | 40 Kya | | | | Marco Romboni |
| GAOcat13 | GAO13 | cat02 | Grotta dell'Onda | Italy | 10.34 | 43.98 | left radius, distal | dist NF | | 40 Kya | | | | Marco Romboni |
| GBcat01 | GB01 | cat01 | Grotta Breuil (Circeo, Latium) | Italy | 41.23 | 13.05 | left humerus, distal | dist F | BT:16.6; Bd:22.7 | middle Palaeolithic | | | | Francesca Alhaique |
| GMcat01 | GM01 | cat01 | Grotta Maritza (Fucino, Abruzzi) | Italy | 41.94 | 13.60 | left tibia, distal part | dist F | Bd 15.4 | 10-11 ka BP uncal / LUP (B2) | | | | Francesca Alhaique |
| GMcat02 | GM02 | cat02 | Grotta Maritza (Fucino, Abruzzi) | Italy | 41.94 | 13.60 | left tibia | prox F; dist F | GL 129.1; Bd 16.2 | 11-11 ka BP uncal / LUP (B3) | | | | Francesca Alhaique |
| GMcat03 | GM03 | cat03 | Grotta Maritza (Fucino, Abruzzi) | Italy | 41.94 | 13.60 | left tibia | prox NF | Bd 15.7 | 12-11 ka BP uncal / LUP (B3) | | | | Francesca Alhaique |
| GMcat04 | GM04 | cat04 | Grotta Maritza (Fucino, Abruzzi) | Italy | 41.94 | 13.60 | left tibia | prox F; dist F | GL 141.3; Bd 17.0 | 13-11 ka BP uncal / LUP (B4-B3) | | | | Francesca Alhaique |
| GMcat05 | GM05 | cat05 | Grotta Maritza (Fucino, Abruzzi) | Italy | 41.94 | 13.60 | left tibia | prox F; dist F | GL 124.8; Bd 14.4 | 14-11 ka BP uncal / LUP (B4-B3) | | | | Francesca Alhaique |
| GMcat06 | GM06 | cat06 | Grotta Maritza (Fucino, Abruzzi) | Italy | 41.94 | 13.60 | left tibia | prox NF, dist F | | 15-11 ka BP uncal / LUP (B4-B3) | | | | Francesca Alhaique |
| GRAcat01 | GRA01 | cat01 | Graffignano (Viterbo, Latium) | Italy | 42.57 | 12.20 | left humerus | prox F, dist F | GL 90.5 Dp 17.5 SD 6.2 Bd 16.5 | Late 15th-Early 16th cent AD | post-Roman | Felis l. lybica/Felis catus clade IV-C | Felis l. lybica/Felis catus | Francesca Alhaique |
| GSAcat01 | GSA01 | cat01 | Genoni Santu Antine, Sardinia | Italy | 39.80 | 9.00 | left femur | prox F, dist F | GL 120.5 Bp 20.8 DC 9.7 SD 8.7 Bd 19.5 | 340BC (3.9%) 320BC / 200BC (91.5%) 50BC | / | Felis l. lybica/Felis catus clade IV-E | Felis l. lybica/Felis catus | Barbara Wilkens |
| LATcat02 | LAT02 | cat02? | Latronico 3, Basilicata | Italy | 40.08 | 16.02 | left humerus, distal | dist F | Bd 21.5 | mesolithic; 7420+-90 BP | | | | Barbara Wilkens |
| LATcat03 | LAT03 | cat03 | Latronico 3, Basilicata | Italy | 40.08 | 16.02 | left humerus, proximal | prox F | Dp 24.6 | mesolithic | | | | Barbara Wilkens |
| LATcat04 | LAT04 | cat04? | Latronico 3, Basilicata | Italy | 40.08 | 16.02 | left tibia, distal | dist F | Bd 16.9; Dd 11.1 | mesolithic | | | | Barbara Wilkens |
| LATcat05 | LAT05 | cat05 | Latronico 3, Basilicata | Italy | 40.08 | 16.02 | | dist F | Bd 15.4 | mesolithic | | | | Barbara Wilkens |
| LATcat06 | LAT06 | cat06 | Latronico 3, Basilicata | Italy | 40.08 | 16.02 | left pelvis | all fused; adult | | mesolithic | | | | Barbara Wilkens |
| LATcat07 | LAT07 | cat07 | Latronico 3, Basilicata | Italy | 40.08 | 16.02 | left pelvis | all fused; adult | | mesolithic | | | | Barbara Wilkens |
| LATcat08 | LAT08 | cat08 | Latronico 3, Basilicata | Italy | 40.08 | 16.02 | right scapula | dist F | | mesolithic | | | | Barbara Wilkens |
| LATcat09 | LAT09 | cat09 | Latronico 3, Basilicata | Italy | 40.08 | 16.02 | left tibia, proximal | prox F | | mesolithic | | | | Barbara Wilkens |
| MARcat01 | MAR01 | cat01 | La Marmotta (Bracciano Lake, Latium) | Italy | 42.10 | 12.28 | left radius, proximal | prox F | Bp 7.2 | Middle Neolithic | | | | Francesca Alhaique |
| MLECcat01 | MLEC01 | cat01 | Muro Leccese | Italy | 40.10 | 18.33 | right metacarpus III | dist F | | 16-17th c AD | | | | Claudia Minniti, Jacopo De Grossi Mazzorin |
| MLECcat02 | MLEC02 | cat02 | Muro Leccese | Italy | 40.10 | 18.33 | right mandible with P3 | dentition fully erupted | (5) 19.1; (10) 8.7 | 18th c AD | | | | Claudia Minniti, Jacopo De Grossi Mazzorin |
| MLECcat05 | MLEC05 | cat03 | Muro Leccese | Italy | 40.10 | 18.33 | atlas | adult | H 16.8; BFcr 25.1; BFcd 19.0 | 16th c AD | | | | Claudia Minniti, Jacopo De Grossi Mazzorin |
| MUScat01 | MUS01 | cat01 | Musarna (Viterbo, Latium) | Italy | 42.41 | 11.98 | atlas | adult | GL 17.9; H 16.0; BFcr 24.4 | 360AD (95.4%) 540AD | Antiquity/Roman | Felis l. lybica/Felis catus clade IV-A1 | Felis silvestris | Francesca Alhaique |
| PADcat01 | PAD01 | cat01 | Padova Via Cesare Battisti (Veneto) | Italy | 45.41 | 11.87 | right scapula | dist F | GLP 13.1; LG 11.6; BG 8.8; SLC 12.4 | 1400AD (95.4%) 1440AD | post-Roman | Felis l. lybica/Felis catus clade IV-A2 | Felis l. lybica/Felis catus | Francesca Alhaique |
| PAZcat01 | PAZ01B | cat01 | Sassari, pozzo di Largo Pazzola, Sardinia | Italy | 40.73 | 8.55 | right mandible with C, P3 & P4 | dentition fully erupted | (5) 19.5; (6) 7.8/3.4; (7) 7.1; (9) 11.6; (10) 1 | 16th c AD | | | | Barbara Wilkens |
| PGLcat01 | PGL01 | cat01 | Grotta Paglicci (Rignano Garganico - Foggia) | Italy | 41.67 | 15.57 | right ulna | -; cf. adult | | 11.440 - 15.320 BP | | | | Paolo Boscato, Francesco Boschin |
| PGLcat02 | PGL02 | cat02 | Grotta Paglicci (Rignano Garganico - Foggia) | Italy | 41.67 | 15.57 | left humerus, distal | distal F | | 11.440 - 15.320 BP | | | | Paolo Boscato, Francesco Boschin |
| PGLcat03 | PGL03 | cat03 | Grotta Paglicci (Rignano Garganico - Foggia) | Italy | 41.67 | 15.57 | left humerus, proximal | prox F | | 11.440 - 15.320 BP | | | | Paolo Boscato, Francesco Boschin |
| PGLcat04 | PGL04 | cat04 | Grotta Paglicci (Rignano Garganico - Foggia) | Italy | 41.67 | 15.57 | left tibia, proximal | prox F | | 11.440 - 15.320 BP | | | | Paolo Boscato, Francesco Boschin |
| PIAcat01 | PIA01 | cat01 | Pianosa | Italy | 42.59 | 10.08 | canine | adult | | 650-770 AD (cal AMS) | | | | Paolo Boscato, Francesco Boschin |
| PNEcat01 | PNE01 | cat01 | Palatino North East slope (Rome) | Italy | 41.89 | 12.49 | left tibia, distal | dist F | Bd 14.6 | 2nd half 6th cent AD (Roman) | | | | Francesca Alhaique |
| PNEcat02 | PNE02 | cat02 | Palatino North East slope (Rome) | Italy | 41.89 | 12.49 | right humerus, distal | dist F | Bd 16.8 | 230AD (95.4%) 370AD | Antiquity/Roman | Felis l. lybica/Felis catus clade IV-C | Felis l. lybica/Felis catus | Francesca Alhaique |
| PNEcat03 | PNE03 | cat03 | Palatino North East slope (Rome) | Italy | 41.89 | 12.49 | right radius, proximal | prox F | Bp 8.4 | End 4th cent.-beg. 5th cent AD (Roman) | Antiquity/Roman | Felis l. lybica/Felis catus clade IV-A2 | Felis l. lybica/Felis catus | Francesca Alhaique |
| PRAcat01 | PRA01 | cat01 | Grotta della Madonna (Praia, Calabria) | Italy | 39.92 | 15.77 | left metatarsus V, proximal | cf. adult | | Mesolithic | | | | Francesca Alhaique |
| PRAcat02 | PRA02 | cat02 | Grotta della Madonna (Praia, Calabria) | Italy | 39.92 | 15.77 | right metacarpus IV | dist F | GL 38.8 | Upper Palaeolitic | | | | Francesca Alhaique |
| PRAcat03 | PRA03 | cat03 | Grotta della Madonna (Praia, Calabria) | Italy | 39.92 | 15.77 | right calcaneus | prox NF | GL 30.5 | Upper Palaeolitic | | | | Francesca Alhaique |
| PRAcat04 | PRA04 | cat04 | Grotta della Madonna (Praia, Calabria) | Italy | 39.92 | 15.77 | right calcaneus | prox NF | GL 31.7 | Upper Palaeolitic | | | | Francesca Alhaique |
| PRAcat05 | PRA05 | cat05 | Grotta della Madonna (Praia, Calabria) | Italy | 39.92 | 15.77 | right metacarpus IV | dist F | GL 36.3 | Upper Palaeolitic | | | | Francesca Alhaique |
| PRNcat01 | PRN01 | cat01 | Partanna C. da Capo d'acqua | Italy | 12.92 | 37.72 | right P3 superior | | | Early Bronze Age | | | | Roberto Micciché |
| PRNcat02 | PRN02 | cat01 | Partanna C. da Capo d'acqua | Italy | 12.92 | 37.72 | os petrosum | cf. adult | | Early Bronze Age | Bronze/Iron Age | Felis silvestris clade I | Felis silvestris | Roberto Micciché |
| PURcat01 | PUR01B | cat01 | La Purissima, Sardinia | Italy | 39.22 | 9.12 | left maxilla with P4 | dentition fully erupted | (12) 21.7; (13) 19.3; (14) 9.8 | roman imperial | | | | Barbara Wilkens |
| RIPcat01 | RIP01 | cat01 | Ripoli | Italy | 0.00 | 0.00 | left femur, proximal | prox F | Bp 23.8; DC 11.8 | Neolitico | | | | Marco Romboni |
| ROCcat01 | ROC01 | cat01 | Roca vecchia (Melendugno, Lecce) | Italy | 18.33 | 40.10 | left humerus, proximal | prox F | Dp 22.3; SD 7.4 | 1499 (84.4%) 1382 calBC / 1341 (11.1%) 1311 calBC | Bronze/Iron Age | Felis silvestris clade I | Felis silvestris | Michela Rugge |
| ROCcat02 | ROC02 | cat02 | Roca vecchia (Melendugno, Lecce) | Italy | 18.33 | 40.10 | right tibia, distal | dist F | SD 7.2; Bd 15.4; Dd 9.7 | 1401 (90.3%) 1256 calBC / 1248 (5.1%) 1227 calBC | Bronze/Iron Age | Felis silvestris clade I | Felis silvestris | Michela Rugge |
| ROCcat04 | ROC04 | cat03 | Roca vecchia (Melendugno, Lecce) | Italy | 18.33 | 40.10 | left mandible with C, P3, P4 & M1 | dentition fully erupted | | Medieval | post-Roman | Felis silvestris lybica/catus clade IV-C | Felis l. lybica/Felis catus | Michela Rugge |
| ROMcat01 | ROM01 | cat01 | Grotta Romanelli (Castro Marina, Apulia) | Italy | 40.02 | 18.43 | right metatarsus III | dist NF | | Upper Palaeolitic (Epigravettian) ca. 10ka | | | | Francesca Alhaique |
| RSScat01 | RSS01 | cat01 | Roccia San Sebastiano (Mondragone, Caserta, Camp | Italy | 41.10 | 13.88 | right humerus, distal | dist F | Bd 23.1 | Upper Palaeolitic (Gravettian), ca. 20ka | | | | Francesca Alhaique |
| SARcat01 | SAR01 | cat01 | Saracena (Cosenza, Calabria) | Italy | 39.77 | 16.15 | left metacarpus IV | dist F | GL 36.4 Bd 5.1 | Early Neolithic | Neolithic/Chalcolithic | Felis silvestris clade I | Felis silvestris | Francesca Alhaique |
| SCAScat01 | SCAS01 | cat01 | Sassari castle, Sardinia | Italy | 40.73 | 8.55 | left metatarsus V | dist F | | modern age | | | | Barbara Wilkens |
| SCATcat01 | SCAT01 | cat01 | Nardò "torre santa Caterina" | Italy | 40.14 | 17.99 | left tibia | prox NF, dist F | SD 6.6; Bd 12.6; Dd 8.3 | 17-18th c AD | post-Roman | Felis l. lybica/Felis catus clade IV-A2 | Felis l. lybica/Felis catus | Jacopo De Grossi Mazzorin |
| SCATcat03 | SCAT03 | cat02 | Nardò "torre santa Caterina" | Italy | 40.14 | 17.99 | left tibia | prox NF, dist F | Bd 12.3; Dd 8.4 | 17-18th c AD | | | | Jacopo De Grossi Mazzorin |
| SDUCcat01 | SDUC01 | cat01 | Sassari, palazzo ducale, Sardinia | Italy | 40.73 | 8.55 | right metatarsus V | adult, dist F | | 19th AD | | | | Barbara Wilkens |
| SDUCcat02 | SDUC02B | cat02 | Sassari, palazzo ducale, Sardinia | Italy | 40.73 | 8.55 | left mandible with P3, P4 & M1 | dentition fully erupted | (1) 59.0; (2) 55.3; (3)52.3; (4) 49.2; (5) 17.4; | 19th AD | post-Roman | Felis l. lybica/Felis catus clade IV-C | Felis l. lybica/Felis catus | Barbara Wilkens |
| SFBIcat01 | SFBI01 | cat01 | Sassari, via del Fiore Bianco, Sardinia | Italy | 40.73 | 8.55 | left humerus | prox NF, dist F | SD 6.5; Bd 17.8 | medieval or modern age | | | | Barbara Wilkens |
| SQCBcat01 | SQCB01 | cat01 | Monastero Ss.Quattro Coronati- Butto della Torre (Ro | Italy | 41.89 | 12.50 | phalanx 1 | prox F | | Mid 16th cent. AD | | | | Francesca Alhaique |
| SQCcat03 | SQC03 | cat02 | Monastero Ss.Quattro Coronati- ex-portico (Rome) | Italy | 41.89 | 12.49 | left calcaneum | prox F | GL 32.5 | end 16th- beg.17th cent AD | | | | Francesca Alhaique |
| SSATcat02 | SSAT02 | cat02 | Sassari, pozzo di via Satta, Sardinia | Italy | 40.73 | 8.55 | right femur, distal | dist F | Bd 16.6 | early 14th AD | | | | Barbara Wilkens |
| SSATcat03 | SSAT03 | cat02? | Sassari, pozzo di via Satta, Sardinia | Italy | 40.73 | 8.55 | right ulna proximal | prox F | SDO 9.2; BPC 10.3 | early 14th AD | | | | Barbara Wilkens |
| SSATcat05 | SSAT05 | cat02? | Sassari, pozzo di via Satta, Sardinia | Italy | 40.73 | 8.55 | right humerus | prox F, dist F | GL 91.1; Dp 18.3; SD 6.1; Bd 16.8 | early 14th AD | | | | Barbara Wilkens |
| SSPIcat01 | SSPI01 | cat01 | Sassari, infermeria San Pietro, Sardinia | Italy | 40.73 | 8.55 | right tibia, proximal | prox NF | Bp 18.9 | 18/19th c AD | | | | Barbara Wilkens |
| STURcat01 | STUR01B | cat01 | Sassari, via Turritana, Sardinia | Italy | 40.73 | 8.55 | right mandible with P3, P4 & M1 | dentition fully erupted | (1) 59.2; (2) 55.4; (3)52.2; (4) 48.9; (5) 17.7; | 17th AD | | | | Barbara Wilkens |
| TERTcat03 | TERT03 | cat01 | Tertenia, Sardinia | Italy | 39.69 | 9.58 | left humerus | prox F, dist F | GL 99.8; Dp 20.5; SD 6.3; Bd 18.5 | 4th-5th c AD | Antiquity/Roman | Felis l. lybica/Felis catus clade IV-A2 | Felis l. lybica/Felis catus | Barbara Wilkens |
| TUScat01 | TUS01 | cat01 | Tusculum, Frascati | Italy | 41.82 | 12.68 | left tibia, shaft | juv., prox NF, dist NF | GL without epiphyses 67.7 | 12th c AD | | | | Marta Moreno |
| UZcat01 | UZ01 | cat01 | Grotta dell'Uzzo (Trapani, Sicily) | Italy | 38.11 | 12.79 | left metacarpus V | dist F | GL 32.3; Bd 5.5 | 5000-4800BC (Neolithic I) | | | | Francesca Alhaique |
| UZcat02 | UZ02 | cat02 | Grotta dell'Uzzo (Trapani, Sicily) | Italy | 38.11 | 12.79 | left metacarpus V | adult, dist F | GL 30.9; Bd 4.6 | 5000-4800BC (Neolithic I) | | | | Francesca Alhaique |
| UZcat03 | UZ03 | cat03 | Grotta dell'Uzzo (Trapani, Sicily) | Italy | 38.11 | 12.79 | right astragalus | cf. adult | GL 16.0 | Neolithic II | | | | Francesca Alhaique |
| UZcat04 | UZ04 | cat04 | Grotta dell'Uzzo (Trapani, Sicily) | Italy | 38.11 | 12.79 | left astragalus | cf. adult | GL 16.3 | Neolithic II | | | | Francesca Alhaique |
| UZcat05 | UZ05 | cat05 | Grotta dell'Uzzo (Trapani, Sicily) | Italy | 38.11 | 12.79 | left astragalus | cf. adult | GL 15.9 | Neolithic I | | | | Francesca Alhaique |
| UZcat06 | UZ06 | cat06 | Grotta dell'Uzzo (Trapani, Sicily) | Italy | 38.11 | 12.79 | left astragalus | cf. adult | GL 17.1 | Mesolithic/Neolithic Transition | | | | Francesca Alhaique |
| ALJcat01 | ALJ01 | cat01 | Castelo de Aljezur (Algarve) | Portugal | 37.32 | -8.80 | left mandible | juvenile; M1 piercing | (1) 54.2; (2) 52.0; (8) 21.4; (9) 10.3; (10) 9.7 | 12-13th c | | | | Cleia Detry |
| BPcat01 | BP01 | cat01 | Bank of Portugal | Portugal | 38.71 | -9.14 | right femur | prox F, dist F | GL 107.2; Bp 20.0; DC 9.7; SD 8.5; Bd 19.0 | 890AD (32.4%) 930AD / 940AD (63.0%) 1020AD | post-Roman | Felis l. lybica/Felis catus clade IV-A1 | Felis l. lybica/Felis catus | Cleia Detry |
| BPcat02 | BP02 | cat02 | Bank of Portugal | Portugal | 38.71 | -9.14 | right femur | prox F, dist F | GL 110.5; Bp 21.5; DC 9.9; SD 9.0; Bd 20.0 | 900AD ( 4.6%) 920AD / 970AD (90.8%) 1040AD | post-Roman | Felis l. lybica/Felis catus clade IV-A2 | Felis l. lybica/Felis catus | Cleia Detry |
| BPcat03 | BP03 | cat03 | Bank of Portugal | Portugal | 38.71 | -9.14 | right humerus, proximal | prox F | Dp 19.4; SD 6.9 | 890AD (11.0%) 920AD / 950AD (84.4%) 1030AD | post-Roman | Felis l. lybica/Felis catus clade IV-A2 | Felis l. lybica/Felis catus | Cleia Detry |
| CSALcat03 | CSAL03 | cat03 | Castelo de Salir, Loulé, Faro | Portugal | 37.24 | -8.05 | left humerus shaft | prox NF, dist NF | GL without epiphyses 55.5 | 12th c AD | | | | Maria João Valente |
| MOScat01 | MOS01 | cat01 | Moita do Sebastião (Muge) | Portugal | 39.10 | -8.71 | left calcaneum | prox F | GL 29.7 | mesolithic | | | | Cleia Detry |
| MOScat02 | MOS02 | cat02 | Moita do Sebastião (Muge) | Portugal | 39.10 | -8.71 | right humerus, distal | dist F | Bd 20.8 | mesolithic | | | | Cleia Detry |
| MOScat03 | MOS03 | cat03 | Moita do Sebastião (Muge) | Portugal | 39.10 | -8.71 | left humerus, distal | dist F | Bd 22.2 | mesolithic | | | | Cleia Detry |
| PALcat08 | PAL08 | cat08 | Palmela Castle | Portugal | 38.57 | -8.90 | right tibia, shaft | juv., prox NF, dist NF | SD 6.5 | 12-13th c | | | | Cleia Detry |
| PALcat09 | PAL09 | cat09 | Palmela Castle | Portugal | 38.57 | -8.90 | right humerus, distal | dist F | Bd 15.5 | 12-13th c | post-Roman | Felis l. lybica/Felis catus clade IV-A2 | Felis l. lybica/Felis catus | Cleia Detry |
| PALcat10 | PAL10 | cat10 | Palmela Castle | Portugal | 38.57 | -8.90 | right tibia | prox F, dist F | GL 102; Bp 17.7; SD 6.6; Bd 12.3 | 11-12th c | post-Roman | Felis l. lybica/Felis catus clade IV-A2 | Felis l. lybica/Felis catus | Cleia Detry |
| PORcat01 | POR01B | cat01 | Portela 3 | Portugal | 37.66 | -8.35 | right mandible with C, P3, P4 & M1 | dentition fully erupted | (10) 9.4 | islamic (10-13 th c AD) | | | | Vera Pereira |
| PORcat02 | POR02B | cat02 | Portela 3 | Portugal | 37.66 | -8.35 | left mandible with C, P3, P4 & M1 | dentition fully erupted | (5) 18.9; (9) 11.1; (10) 10.5 | islamic (10-13 th c AD) | | | | Vera Pereira |
| PORcat03 | POR03B | cat03 | Portela 3 | Portugal | 37.66 | -8.35 | right mandible with P3, P4 & M1 | dentition fully erupted | (5) 21.5; (9) 10.9; (10) 8.0 | islamic (10-13 th c AD) | | | | Vera Pereira |
| PORcat04 | POR04B | cat04 | Portela 3 | Portugal | 37.66 | -8.35 | right mandible with P3, P4 & M1 | dentition fully erupted | (5) 18.1; (6) 7.3/3.9; (7) 7.0; (9) 11.0; (10)9. | islamic (10-13 th c AD) | | | | Vera Pereira |
| TOVcat01 | TOV01 | cat01 | Torre Velha 3 (Serpa) | Portugal | 37.21 | -8.32 | right humerus, distal | dist F | Bd 14.1 | Late Antiquity | | | | Cleia Detry |
| PTRcat01 | PTR01B | cat01 | Petrovac, Kladurovo | Serbia | 44.44 | 21.54 | left mandible with C, P3, P4 & M1 | dentition fully erupted | (1) 64.5; (2) 60.2; (3) 57.1; (4) 52.8; (5) 20.4 | recent | | | | Günther Karl Kunst |
| VIAcat01 | VIA01 | cat01 | Viminacium/Amphitheatre | Serbia | 44.74 | 21.23 | cranium | adult | | 4th century AD | | | | Sonja Vuković |
| VIAcat02 | VIA02 | cat02 | Viminacium/Amphitheatre | Serbia | 44.74 | 21.23 | left mandible with C, P3, P4 & M1 | dentition fully erupted | (1) 63.6; (2) 58.8; (3)56.2; (4) 52.1; (5) 19.2; | 2nd-3rd century AD | | | | Sonja Vuković |
| VIAcat03 | VIA03 | cat03 | Viminacium/Amphitheatre | Serbia | 44.74 | 21.23 | left mandible with C, P4 & M1 | dentition fully erupted | (1) 61.7; (2) 58.6; (3)54.3; (4) 51.2; (5) 19.3; | 4th century AD | | | | Sonja Vuković |
| VIAcat04 | VIA04 | cat04 | Viminacium/Amphitheatre | Serbia | 44.74 | 21.23 | left mandible with C, P3 & M1 | dentition fully erupted | (1) 58.9; (2) 55.0; (3) 51.3; (4) 48.1; (5) 18.9 | 250AD (16.8%) 280AD / 330AD (78.6%) 420AD | Antiquity/Roman | Felis l. lybica/Felis catus clade IV-C | Felis l. lybica/Felis catus | Sonja Vuković |
| VIAcat05 | VIA05 | cat05 | Viminacium/Amphitheatre | Serbia | 44.74 | 21.23 | left scapula | prox F | HS 55.5; GLP 13.6; LG 12.3; BG 8.9; SLC 1 | 4th century AD | | | | Sonja Vuković |
| VIAcat06 | VIA06 | cat06 | Viminacium/Amphitheatre | Serbia | 44.74 | 21.23 | right humerus, distal | dist F | Bd 21.7 | 20AD (95.4%) 210AD | Antiquity/Roman | Felis l. lybica/Felis catus clade IV-C | Felis l. lybica/Felis catus | Sonja Vuković |
| VIAcat07 | VIA07 | cat07 | Viminacium/Amphitheatre | Serbia | 44.74 | 21.23 | left humerus, distal | dist F | SD 7.6; Bd 20.0 | 2nd-3rd century AD | | | | Sonja Vuković |
| VIAcat08 | VIA08 | cat08 | Viminacium/Amphitheatre | Serbia | 44.74 | 21.23 | right humerus, distal | dist F | SD 5.5; Bd 17.6 | 4th century AD | | | | Sonja Vuković |
| VIAcat09 | VIA09 | cat09 | Viminacium/Amphitheatre | Serbia | 44.74 | 21.23 | left ulna | prox F | SDO 11.3; DPA 12.4; BPC 9.9 | 2nd century AD | | | | Sonja Vuković |
| VIAcat10 | VIA10 | cat10 | Viminacium/Amphitheatre | Serbia | 44.74 | 21.23 | right femur | prox F, dist F | GL 123.5; Bp 22.6; DC 10.6; SD 8.7; Bd 19.5 | 3rd century AD | | | | Sonja Vuković |
| VINcat04 | VIN04 | cat02 | Viminacium-Nad Klepačkom | Serbia | 44.74 | 21.23 | right humerus | prox NF, dist F | SD 7.0; Bd 20.9 | 990AD (49.5%) 1050AD / 1080AD (45.9%) 1160AD | post-Roman | Felis silvestris clade I | Felis silvestris | Sonja Vuković |
| VINcat06 | VIN06 | cat04 | Viminacium/Nad Klepačkom | Serbia | 44.74 | 21.23 | left humerus | prox NF, dist F | SD 6.8; Bd 20.7 | 9th-11th century AD (?) | | | | Sonja Vuković |
| VINcat07 | VIN07 | cat05 | Viminacium/Nad Klepačkom | Serbia | 44.74 | 21.23 | right femur | prox NF, dist NF | DC 10.5; SD 6.3 | 9th-11th century AD (?) | | | | Sonja Vuković |
| ARGcat01 | ARG01B/T | cat01 | Castillo de Arganzón | Spain | 42.88 | -2.80 | right mandible with C, P3, P4 & M1 | dentition fully erupted | (1) 56.7; (2) 53.5; (3) 50.3; (4) 47.1; (5) 17.0 | late medieval | | | | Idoia Grau |
| BORcat04 | BOR04 | cat04 | El Bordellet | Spain | 41.35 | 1.69 | mandible with all teeth in situ | dentition fully erupted | (1) 55.8; (2) 52.6; (3) 48.9; (4) 45.6; (5) 19.1 | cal AD 970-1025 | | | | Lluís Lloveras, Jordi Nadal |
| BORcat06 | BOR06 | cat06 | El Bordellet | Spain | 41.35 | 1.69 | mandible with all teeth in situ | dentition fully erupted | (1) 53.9; (2) 48.6; (3) 48.1; (4) 43.2; (5) 18.0 | cal AD 970-1025 | | | | Lluís Lloveras, Jordi Nadal |
| BORcat08 | BOR08 | cat08 | El Bordellet | Spain | 41.35 | 1.69 | mandible with all teeth in situ | dentition fully erupted | (1) 52.3; (2) 48.1; (3) 45.9; (4) 43.4; (5) 18.0 | cal AD 970-1025 | | | | Lluís Lloveras, Jordi Nadal |
| CETcat01 | CET01 | cat01 | Cova de Els Trocs, Huesca | Spain | 42.47 | 0.53 | right ulna | prox F, dist F | GL 126.5; BPC 10.6; DPA 12.8; SDO 10.7 | 5300BC (9.4%) 5250BC / 5230BC (86.0%) 5040BC (dated twice accidentally) | Neolithic/Chalcolithic | Felis silvestris clade I | Felis silvestris | Marta Moreno |
| LABcat01 | LAB01 | cat01 | Labastida | Spain | 42.58 | -2.78 | right metatarsus V | adult, dist F | | end 15th-early 16th c. | post-Roman | Felis l. lybica/Felis catus clade IV-A2 | Felis l. lybica/Felis catus | Idoia Grau |
| PENcat01 | PEN01 | cat01 | Peñalosa, Jaén | Spain | 38.17 | -3.80 | left tibia, shaft | juv., prox NF, dist NF | GL without epiphyses 104.5 | Bronze Age | | | | Marta Moreno |
| PENcat02 | PEN02 | cat02 | Peñalosa, Jaén | Spain | 38.17 | -3.80 | left M1 inferior | M1 functional | | Bronze Age | | | | Marta Moreno |
| PENcat04 | PEN04 | cat04 | Peñalosa, Jaén | Spain | 38.17 | -3.80 | left metatarsus V, proximal | cf. adult | | Bronze Age | | | | Marta Moreno |
| TABcat01 | TAB01 | cat01 | Tabernas, Spain | Spain | 37.05 | -2.39 | left mandible, teeth partially preserved | dentition fully erupted | | 2,700-2,400 BC (Phase III) | | | | Joris Peters, Bea De Cupere |
| TABcat02 | TAB02 | cat02 | Tabernas, Spain | Spain | 37.05 | -2.39 | right humerus, distal | dist F | Bd 22.9 | 2,700-2,400 BC (Phase III) | Bronze/Iron Age | Felis silvestris clade I | Felis silvestris | Joris Peters, Bea De Cupere |
| ZAPcat02 | ZAP02 | cat02 | Zapatari 33 | Spain | 42.85 | -2.38 | left humerus | prox F, dist F | GL 84.0; Dp 16.3; SD 5.6; Bd 15.3 | mid 13th-mid 16th c. | | | | Idoia Grau |
| ZAPcat03 | ZAP03 | cat03 | Zapatari 33 | Spain | 42.85 | -2.38 | left humerus | prox NF, dist F | SD 5.3; Bd 15.7 | mid 13th-mid 16th c. | | | | Idoia Grau |
| ZORcat02 | ZOR02 | cat01 | Zornoztegi | Spain | 42.88 | -2.39 | right humerus | prox NF, dist F | SD 5.4; Bd 16.1 | 8th-10th c. | | | | Idoia Grau |
| ASKcat01 | ASK01 | cat01 | Aşıklı Höyük | Turkey | 38.35 | 34.23 | right humerus, distal | prox F | Bd 20.6 | 7950BC (95.4%) 7610BC | Neolithic/Chalcolithic | Felis l. lybica/Felis catus clade IV-A0 | Felis silvestris | Hijlke Buitenhuis |
| BAcat02 | BA02 | cat02 | Bademağacı | Turkey | 37.21 | 30.47 | right mandible with P3, P4 & M1 | dentition fully erupted | (1) 62.8; (2) 60.7; (3) 54.6; (4) 53.5; (5) 22.6 | 6430BC (61.9%) 6330BC / 6320BC (33.5%) 6240BC | Neolithic/Chalcolithic | Felis l. lybica/Felis catus clade IV-A0 | Felis silvestris | Bea De Cupere |
| BAcat03 | BA03 | cat03 | Bademağacı | Turkey | 37.21 | 30.47 | left mandible, teeth lost | dentition fully erupted | (5) 23; (7) 9.1; (10) 10.5 | 6390BC (90.0%) 6210BC / 6140BC (5.4%) 6090BC | Neolithic/Chalcolithic | Felis l. lybica/Felis catus clade IV-A0 | Felis silvestris | Bea De Cupere |
| BAcat04 | BA04 | cat04 | Bademağacı | Turkey | 37.21 | 30.47 | right pelvis | -; cf. adult | | Neolithic 6500-6000 BC | | | | Bea De Cupere |
| BAYcat01 | BAY01 | cat01 | Bayraklı | Turkey | 38.46 | 27.17 | left femur, complete shaft | subadult, prox NF; dist NF | GL without epiphyses 102 | 1955AD (95.4%) 2014AD | | | | Vedat Onar |
| BAYcat02 | BAY02 | cat01? | Bayraklı | Turkey | 38.46 | 27.17 | lumbar vertebra | (sub)adult; NF | | 4th century BC | | | | Vedat Onar |
| BAYcat03 | BAY03 | cat02 | Bayraklı | Turkey | 38.46 | 27.17 | left ulna, proximal | prox NF? | BPC 9.1 | 4th century BC | | | | Vedat Onar |
| DDcat03 | DD03 | cat03 | Didyma | Turkey | 37.39 | 27.26 | left humerus | prox NF, dist F | SD 5.8; Bd 17.7 | 900AD ( 3.8%) 930AD / 940AD (73.6%) 1060AD / 1070AD (18.0%) 1160AD | | | | Joris Peters, Bea De Cupere |
| DHcat03 | DH03 | cat03 | Demircihüyük | Turkey | 40.17 | 29.98 | left radius, distal | dist F | SD 6.7; Bd 14,7 | 3000-2500 BC | | | | Joris Peters, Bea De Cupere |
| DHcat06 | DH06 | cat01 | Demircihüyük | Turkey | 40.17 | 29.98 | left ulna | prox NF, dist NF | SDO 11.4; DPA 12.6; BPC 9.6 | 1302 (74.8%) 1369calAD / 1380 (20.6%) 1401calAD | post-Roman | Felis l. lybica/Felis catus clade IV-A2 | Felis l. lybica/Felis catus | Joris Peters, Bea De Cupere |
| DHcat07 | DH07 | cat05 | Demircihüyük | Turkey | 40.17 | 29.98 | left tibia, distal | dist F | Bd 16.1 | 3000-2500 BC | | | | Joris Peters, Bea De Cupere |
| IRTcat01 | IRT01 | cat01 | Iznik - Roman theatre | Turkey | 40.42 | 29.72 | left humerus | prox NF, dist F | GL without epiphyses 84.6; SD 6.0; Bd 16.6 | 575AD (95.4%) 650AD | post-Roman | Felis l. lybica/Felis catus clade IV-C | Felis l. lybica/Felis catus | Vedat Onar |
| IRTcat02 | IRT02 | cat01? | Iznik - Roman theatre | Turkey | 40.42 | 29.72 | right ulna | prox NF, dist F | GL without epiphyses 80.6; BPC 8.5 | Roman-Byzantine? | | | | Vedat Onar |
| MENTcat01 | MENT01 | cat01 | Menteşe | Turkey | 40.28 | 29.52 | right ulna shaft | -; adult | BPC 9.0 | 6073 (95.4%) 5986 calBC | Neolithic/Chalcolithic | Felis l. lybica/Felis catus clade IV-A2 | Felis silvestris | Lionel Gourichon |
| MRYcat07 | MRY07 | cat07 | Yenikapı, Marmaray exc. | Turkey | 41.00 | 28.95 | left femur | prox F, dist F | GL 113.0; Bp 20.7; SD 7.8; Bd 18.7 | 4th-14th cent AD? | post-Roman | Felis l. lybica/Felis catus clade IV-A2 | Felis l. lybica/Felis catus | Vedat Onar |
| MRYcat08 | MRY08 | cat08 | Yenikapı, Marmaray exc. | Turkey | 41.00 | 28.95 | right femur | prox F, dist F | GL 119.3; Bp 21.7; DC 10.1; SD 8.5; Bd 20. | 4th-14th cent AD? | post-Roman | Felis l. lybica/Felis catus clade IV-C | Felis l. lybica/Felis catus | Vedat Onar |
| MRYcat09 | MRY09 | cat09 | Yenikapı, Marmaray exc. | Turkey | 41.00 | 28.95 | right femur | prox F, dist F | GL 100.7; Bp 18.5; DC 8.7; SD 6.4; Bd 17.0 | 650AD (48.9%) 710AD / 720AD (46.5%) 780AD (dated twice accedentally) | post-Roman | Felis l. ornata clade III | Felis l. lybica/Felis catus | Vedat Onar |
| MRYcat10 | MRY10 | cat10 | Yenikapı, Marmaray exc. | Turkey | 41.00 | 28.95 | right femur | prox F, dist F | GL 105.2; Bp 19.6; DC 8.7; SD 7.1; Bd 18.2 | 640AD (75.4%) 690AD / 740AD (20.0%) 780AD | post-Roman | Felis l. lybica/Felis catus clade IV-E | Felis l. lybica/Felis catus | Vedat Onar |
| SAcat09 | SA09 | cat09 | Sagalassos | Turkey | 37.68 | 30.52 | left femur | dist NF | Bp 20.0; DC 10.0 | Early Byzantine (7th cent. AD) | | | | Bea De Cupere |
| SAcat14 | SA14 | cat14 | Sagalassos | Turkey | 37.68 | 30.52 | left humerus | prox NF, dist F | SD 5.2; Bd 16.2 | Early Byzantine | post-Roman | Felis l. lybica/Felis catus clade IV-E | Felis l. lybica/Felis catus | Bea De Cupere |
| TRLcat01 | TRL01 | cat01 | Tralles | Turkey | 37.85 | 27.85 | right femur | prox F, dist F | GL 122.5; Bp 22.3; DC 11.1; SD 9.4; Bd 22.2 | Roman? | | | | Vedat Onar |
| TRLcat02 | TRL02 | cat01 | Tralles | Turkey | 37.85 | 27.85 | left mandible with M1 | dentition fully erupted | (1) 64.6; (2) 60.4; (3) 58.4; (4) 54.8; (5) 22.3 | Roman? | | | | Vedat Onar |

*F=fused; NF=not fused; prox: proximal; dist=distal

**measurements and abbreviations as in von den Driesch (1976).

**Supplementary table 2.** Results of the shallow shotgun sequencing conducted on genomic libraries from the archaeological samples.

| ID | Sample ID | ID-Lab | Individual | Site | Site | Chronology / C14 date | extraction batch coc | Library code | QC_passed_reads (w/o dup) | Full genome mapped (w/o dup) | Full genome mapped -q 1 (w/o dup) | DeDup duplicates | % duplicates | % spurious | % endogenous | %Nuclear 5' PMD | %Nuclear 3' PMD | mtDNA mapped -q 1 (w/o dup) | fold mt-cov | mt-3X |
|---|---|---|---|---|---|---|---|---|---|---|---|---|---|---|---|---|---|---|---|---|
| ACcat01 | AC01 | AC01 | cat01 | Italy | Arene Candide (Finale Ligure, Savona, Liguria) | 10-11 ka (Epigravettian) | FX46 | FX-CO | 8447954 | 5864584 | 4648560 | 667323.00 | 10.22 | 20.74 | 51.0 | 25.23 | 21.18 | 6168 | 21.6936 | 99.08 |
| ACcat02 | AC02 | AC02 | cat02 | Italy | Arene Candide (Finale Ligure, Savona, Liguria) | 10-11 ka (Epigravettian) | FX7 | FX-AG | 11404332 | 33861 | 24832 | 23283.00 | 40.74 | 26.66 | 0.2 | 23.18 | 24.5 | 18 | 0.05 | 0 |
| AFcat03 | AF03 | AF03 | cat02 | Italy | Alba Fucens -Cisterna (Fucino, Abruzzi) | 210-340 AD (cal AMS) | FX28 | FX-BI | 9305507 | 3786188 | 3460748 | 219359.00 | 5.48 | 8.60 | 36.3 | 28.55 | 29.3 | 8217 | 30.9039 | 99.37 |
| AKRcat02 | AKR02 | AKR02 | cat02 | Italy | Akrai, Sicily | late 3rd – mid 4th c. AD | FX31 | FX-BO | 12393949 | 184708 | 168638 | 10526.00 | 5.39 | 8.70 | 1.4 | 37.36 | 34.2 | 501 | 1.64 | 24.18 |
| AKRcat02 | AKR02 | AKR02 | cat02 | Italy | Akrai, Sicily | late 3rd – mid 4th c. AD | FX28 | FX-BI | 9855523 | 35593 | 32252 | 1057.00 | 2.88 | 9.39 | 0.3 | 35.61 | 36.9 | 58 | 0.19 | 0 |
| ANGcat01 | ANG01 | ANG01 | cat01 | Italy | Grotta S.Angelo (Ostuni, Apulia) | “Pleistocene” (Upper Paleolithic?) | FX5 | FX-AE | 17424402 | 40460 | 36247 | 13359.00 | 24.82 | 10.41 | 0.2 | 48.35 | 47.6 | 57 | 0.18 | 0.21 |
| ARGcat01 | ARG01B | ARG01B | cat01 | Spain | Castillo de Arganzón | late medieval | FX3 | FX-AC | 16721610 | 677080 | 567313 | 309361.00 | 31.36 | 16.21 | 3.3 | 16.03 | 16.1 | 1320 | 5.13 | 77.1 |
| ASKcat01 | ASK01 | ASK01 | wildcat1 | Turkey | Aşıklı Höyük | 7950BC (95.4%) 7610BC | FX42 | FXCI2 | 4978205 | 629042 | 419122 | 300461.00 | 32.32 | 33.37 | 7.9 | 44.57 | 42.25 | 2641 | 7.1076 | 89.08 |
| ASKcat01 | ASK01 | ASK01 | wildcat1 | Turkey | Aşıklı Höyük | 7950BC (95.4%) 7610BC | FX42 | FX-DO | 7322616 | 723297 | 466697 | 574627.00 | 44.27 | 35.48 | 5.9 | 42.61 | 38.5 | 2379 | 6.7486 | 89.83 |
| BAcat02 | BA02 | BA02 | wildcat2 | Turkey | Bademağacı | 6430BC (61.9%) 6330BC / 6320BC (33.5%) 6240BC | FX32 | FX-BS | 4062837 | 2211722 | 1995041 | 388145.00 | 14.93 | 9.80 | 44.8 | 45.62 | 46.3 | 1599 | 4.3294 | 70.9 |
| BAcat03 | BA03 | BA03 | wildcat3 | Turkey | Bademağacı | 6390BC (90.0%) 6210BC / 6140BC (5.4%) 6090BC | FX32 | FX-BS | 3918962 | 1341862 | 1162804 | 203685 | 13.18 | 13.34 | 28.2 | 46.39 | 47.17 | 3249 | 9.1089 | 96.26 |
| BAcat04 | BA04 | BA04 | wildcat4 | Turkey | Bademağacı | Neolithic 6500-6000 BC | FX32 | FX-BS | 4587442 | 9701 | 8395 | 1052 | 9.78 | 13.46 | 0.2 | 50.48 | 51.79 | 298 | 0.8942 | 6.56 |
| BAYcat01 | BAY01 | BAY01 | cat01 | Turkey | Bayraklı | 1955AD (95.4%) 2014AD | FX35 | FX-CB | 6122785 | 3726135 | 3461489 | 76422 | 2.01 | 7.10 | 55.8 | 4.43 | 4.56 | 13317 | 43.9976 | 99.94 |
| BAYcat02 | BAY02 | BAY02 | cat01? | Turkey | Bayraklı | 4th century BC | FX35 | FX-CB | 7229732 | 5363445 | 4919572 | 111311 | 2.03 | 8.28 | 67.0 | 2.07 | 2.14 | 15639 | 52.0775 | 99.97 |
| BAYcat03 | BAY03 | BAY03 | cat02 | Turkey | Bayraklı | 4th century BC | FX29 | FX-BN | 12969665 | 2703364 | 2426483 | 60230 | 2.18 | 10.24 | 18.6 | 1.01 | 1.5 | 1201 | 3.78 | 66.22 |
| BG01 | BG01 | BG01 | wild cat 1 | Bulgaria | #N/A | modern (museum collection) | FXM2 | FX-BU | 4683896 | 3190792 | 1093288 | 60010 | 1.85 | 65.74 | 23.1 | 4.08 | 4.02 | 297 | 1.7494 | 26.48 |
| BG07 | BG07 | BG07 | wild cat 7 | Bulgaria | #N/A | modern (museum collection) | FXM2 | FX-BU | 6579265 | 5175483 | 3700624 | 231626 | 4.28 | 28.50 | 54.3 | 2.18 | 2.25 | 1127 | 6.3177 | 47.65 |
| BG16 | BG16 | BG16 | wild cat 16 | Bulgaria | #N/A | modern (museum collection) | FXM2 | FX-BU | 6836917 | 4665932 | 1939741 | 426936 | 8.38 | 58.43 | 26.7 | 2.26 | 2.36 | 301 | 1.5404 | 17.78 |
| BG2 | BG2 | BG2 | wild cat 2 | Bulgaria | #N/A | modern (museum collection) | FXM2 | FX-BU | 5450179 | 4519346 | 1360286 | 547337 | 10.80 | 69.90 | 22.7 | 2.88 | 2.95 | 219 | 1.0243 | 12.63 |
| BG5 | BG5 | BG5 | wild cat 5 | Bulgaria | #N/A | modern (museum collection) | FXM2 | FX-BU | 6135340 | 3747563 | 1834819 | 60152 | 1.58 | 51.04 | 29.6 | 4.12 | 4.15 | 105 | 0.602 | 7.58 |
| BHHLcat01 | BHHL01 | BHHL01 | cat01 | Austria | Bernhardsthal | 770AD (95.4%) 980AD | FX41 | FX-CH | 8541381 | 5465889 | 4949060 | 529799 | 8.84 | 9.46 | 54.6 | 18.52 | 16.28 | 9238 | 31.3935 | 99.9 |
| BHHLcat01 | BHHL01 | BHHL01 | cat01 | Austria | Bernhardsthal | 770AD (95.4%) 980AD | FX29 | FX-BN | 10888878 | 2386728 | 2186188 | 86100 | 3.48 | 8.40 | 19.9 | 19.16 | 16.4 | 5170 | 17.3346 | 99.57 |
| BHHLcat03 | BHHL03 | BHHL03 | cat02? | Austria | Bernhardsthal | Roman | FX41 | FX-CH | 7562355 | 1468430 | 1300102 | 135620 | 8.45 | 11.46 | 16.9 | 17.19 | 13.82 | 1448 | 5.2602 | 84.29 |
| BHHLcat03 | BHHL03 | BHHL03 | cat02? | Austria | Bernhardsthal | Roman | FX29 | FX-BN | 10808192 | 535799 | 475931 | 27562 | 4.89 | 11.17 | 4.4 | 18.87 | 15.8 | 655 | 2.33 | 41.55 |
| BORcat04 | BOR04 | BOR04 | cat04 | Spain | El Bordellet | cal AD 970-1025 | FX28 | FX-BI | 10873456 | 152681 | 138622 | 4218 | 2.69 | 9.21 | 1.3 | 34.03 | 35.3 | 390 | 1.38 | 16.87 |
| BORcat04 | BOR04T | BOR04T | cat04 | Spain | El Bordellet | cal AD 970-1025 | FX59 | FX-DF | 9034961 | 4255 | 3524 | 420 | 8.98 | 17.18 | 0.0 | 24.04 | 21.17 | 57 | 0.1533 | 0.08 |
| BORcat06 | BOR06 | BOR06 | cat06 | Spain | El Bordellet | cal AD 970-1025 | FX28 | FX-BI | 10333116 | 266986 | 241936 | 4735 | 1.74 | 9.38 | 2.3 | 29.39 | 30 | 243 | 0.76 | 5.49 |
| BORcat06 | BOR06 | BOR06 | cat06 | Spain | El Bordellet | cal AD 970-1025 | FX44 | FX-CN | 6968053 | 59825 | 43439 | 285677 | 82.68 | 27.39 | 0.6 | 26.83 | 21.57 | 61 | 0.1903 | 0.45 |
| BORcat08 | BOR08 | BOR08 | cat08 | Spain | El Bordellet | cal AD 970-1025 | FX29 | FX-BN | 11616404 | 306436 | 281289 | 5182 | 1.66 | 8.21 | 2.4 | 22.36 | 17.9 | 351 | 1.21 | 12.69 |
| BORcat08 | BOR08 | BOR08 | cat08 | Spain | El Bordellet | cal AD 970-1025 | FX44 | FX-CN | 7952118 | 14697 | 11360 | 41357 | 73.78 | 22.71 | 0.1 | 22.29 | 20.15 | 24 | 0.0817 | 0.32 |
| BORcat08 | BOR08T | BOR08T | cat08 | Spain | El Bordellet | cal AD 970-1025 | FX59 | FX-DF | 7217888 | 3816 | 3118 | 381 | 9.08 | 18.29 | 0.0 | 18.56 | 17.29 | 38 | 0.1286 | 0 |
| BPcat01 | BP01 | BP01 | cat01 | Portugal | Bank of Portugal | Roman (Roman Imperial II) | FX62 | FX-DO | 6127743 | 1394596 | 1266765 | 73810 | 5.03 | 9.17 | 20.4 | 40.66 | 36 | 5734 | 20.6345 | 99.5 |
| BPcat02 | BP02 | BP02 | cat02 | Portugal | Bank of Portugal | Roman (Roman Imperial II) | FX62 | FX-DO | 6728241 | 1397897 | 1268196 | 38870 | 2.71 | 9.28 | 18.7 | 38.66 | 36.56 | 2346 | 7.6882 | 93.93 |
| BPcat03 | BP03 | BP03 | cat03 | Portugal | Bank of Portugal | Roman (Roman Imperial II) | FX62 | FX-DO | 6263845 | 968184 | 862995 | 41554 | 4.12 | 10.86 | 13.7 | 26.21 | 23.79 | 1051 | 3.5665 | 61.67 |
| BREMcat05 | BREM05 | BREM05 | cat04 | Germany | Bremen; 201-Altstadt, Marktplatz | 1440AD (65.0%) 1530AD / 1570AD (30.4%) 1630AD | FX29 | FX-BN | 9525173 | 2823599 | 2580374 | 145591 | 4.90 | 8.61 | 26.7 | 17.25 | 14 | 2658 | 9.3533 | 96.12 |
| BREMcat05 | BREM05 | BREM05 | cat04 | Germany | Bremen; 201-Altstadt, Marktplatz | 1440AD (65.0%) 1530AD / 1570AD (30.4%) 1630AD | FX42 | FX-CI | 9241362 | 2415655 | 2198785 | 139460 | 5.46 | 8.98 | 23.4 | 16.43 | 13.72 | 2047 | 7.0942 | 92.57 |
| BREMcat07 | BREM07 | BREM07 | cat06 | Germany | Bremen; 253-Altstadt, Am Wall | 16th | FX29 | FX-BN | 12008015 | 2260664 | 2085114 | 82830 | 3.53 | 7.77 | 17.3 | 14.16 | 11.5 | 4928 | 17.37 | 99.84 |
| BREMcat07 | BREM07 | BREM07 | cat06 | Germany | Bremen; 253-Altstadt, Am Wall | 16th | FX42 | FX-CI | 7841793 | 1388625 | 1276369 | 68797 | 4.72 | 8.08 | 16.1 | 14.18 | 11.5 | 3973 | 14.1186 | 99.58 |
| BRPAcat01 | BRPA01 | BRPA01 | cat01 | Belgium | Parking 58, Brussels | end 14th - 15th cent. AD | FX56 | FX-DE | 3814622 | 1361359 | 1233767 | 92525 | 6.36 | 9.37 | 31.6 | 8.1 | 7.88 | 6390 | 23.0713 | 99.89 |
| BRPAcat01 | BRPA01 | BRPA01 | cat01 | Belgium | Parking 58, Brussels | end 14th - 15th cent. AD | FX20 | FX-AV | 9000246 | 956487 | 848926 | 610664 | 38.97 | 11.25 | 8.8 | 6.5 | 4.7 | 4849 | 20.11 | 99.8 |
| BRPAcat11 | BRPA11 | BRPA11 | cat11 | Belgium | Parking 58, Brussels | mid 13th cent. AD | FX23 | FX-BB | 8518323 | 487083 | 429081 | 86142 | 15.03 | 11.91 | 5.0 | 9.7 | 9 | 1536 | 5.72 | 86.57 |
| BRPAcat16 | BRPA16 | BRPA16 | cat16 | Belgium | Parking 58, Brussels | 12th - 13th cent. AD | FX56 | FX-DE | 3833376 | 503707 | 460567 | 13164 | 2.55 | 8.56 | 12.0 | 20.63 | 17.87 | 810 | 2.7417 | 50.58 |
| BRPAcat16 | BRPA16 | BRPA16 | cat16 | Belgium | Parking 58, Brussels | 12th - 13th cent. AD | FX23 | FX-BB | 6647607 | 509258 | 464632 | 31799 | 5.88 | 8.76 | 7.0 | 19.5 | 15.7 | 756 | 2.78 | 50.93 |
| BRPAcat22 | BRPA22 | BRPA22 | cat02B | Belgium | Parking 58, Brussels | end 14th - 15th cent. AD | FX20 | FX-AV | 8818494 | 223213 | 186837 | 250475 | 52.88 | 16.30 | 2.1 | 12.4 | 9.3 | 1036 | 4.38 | 78.93 |
| BRPAcat24 | BRPA24_SS | BRPA24_SS | cat04B | Belgium | Parking 58, Brussels | end 14th - 15th cent. AD | FX20 | FX-AV | 6449610 | 906712 | 846326 | 15356 | 1.67 | 6.66 | 13.1 | 9.79 | 0.5 | 4555 | 18.84 | 99.81 |
| BRPAcat24 | BRPA24 | BRPA24 | cat04B | Belgium | Parking 58, Brussels | end 14th - 15th cent. AD | FX20 | FX-AV | 9573784 | 553413 | 487922 | 695957 | 55.70 | 11.83 | 4.8 | 10.6 | 7.8 | 2106 | 8.63 | 97.98 |
| BRPAcat27 | BRPA27 | BRPA27 | cat07B | Belgium | Parking 58, Brussels | end 14th - 15th cent. AD | FX56 | FX-DE | 4165807 | 1553394 | 1400445 | 102820 | 6.21 | 9.85 | 32.8 | 15.1 | 15.31 | 4356 | 13.3481 | 99.15 |
| BRPAcat27 | BRPA27 | BRPA27 | cat07B | Belgium | Parking 58, Brussels | end 14th - 15th cent. AD | FX20 | FX-AV | 9129704 | 451064 | 404442 | 211212 | 31.89 | 10.34 | 4.3 | 14.9 | 11.7 | 1458 | 5.8 | 90.68 |
| BUD01 | BUD01 | BUD01 | wildcat1 | Bulgaria | Budjaka (near Sozopol) | Late Neolithic | FX34 | FXCA | 5115089 | 58772 | 50485 | 3630 | 5.82 | 14.10 | 1.0 | 43.46 | 43.15 | 198 | 0.5511 | 2.25 |
| BUD01 | BUD01 | BUD01 | wildcat1 | Bulgaria | Budjaka (near Sozopol) | Late Neolithic | FX27 | FX-BF | 11600474 | 12790 | 10767 | 803 | 5.91 | 15.82 | 0.1 | 34.02 | 33 | 47 | 0.14 | 0 |
| CALAcat01 | CALA01 | CALA01 | cat01 | Italy | Grotta della Cala (Marina di Camerota - Salerno) | 8370±80 BP; 8060±100 | FX8 | FX-AH | 11601685 | 11709 | 10072 | 2099 | 15.20 | 13.98 | 0.1 | 39.01 | 38.7 | 85 | 0.3 | 0.94 |
| CALAcat02 | CALA02 | CALA02 | cat02 | Italy | Grotta della Cala (Marina di Camerota - Salerno) | 8370±80 BP; 8060±100 | FX8 | FX-AH | 15964428 | 14052 | 12100 | 3510 | 19.99 | 13.89 | 0.1 | 37.91 | 38.9 | 27 | 0.12 | 0.25 |
| CALcat01 | CAL01 | CAL01 | cat01 | Italy | Calvatone, Lombardia, Italy | Roman, Augustean | FX4 | FX-AD | 13776757 | 674495 | 557675 | 226356 | 25.13 | 17.32 | 4.0 | 23.3 | 23.3 | 616 | 2.37 | 39.11 |
| CALcat01 | CAL01 | CAL01 | cat01 | Italy | Calvatone, Lombardia, Italy | Roman, Augustean | FX52 | FX-DB | 4333054 | 2789 | 2308 | 73 | 2.55 | 17.25 | 0.1 | 28.17 | 23.35 | 1 | 0.0026 | 0 |
| CASTcat02 | CAST02 | CAST02 | cat01 | Italy | Castiglione in Sabina | 11-12th c AD | FX44 | DPs-AH | 4621027 | 3008508 | 2616641 | 2034676 | 40.35 | 13.03 | 39.3 | 23.69 | 21.21 | 4461 | 15.3941 | 98.67 |
| CASTcat02 | CAST02 | CAST02 | cat01 | Italy | Castiglione in Sabina | 11-12th c AD | FX44 | FXCN | 3736109 | 1638814 | 1314672 | 3706915 | 69.34 | 19.78 | 17.7 | 23.41 | 19.57 | 2134 | 7.6597 | 94.67 |
| CASTcat02 | CAST02 | CAST02 | cat01 | Italy | Castiglione in Sabina | 11-12th c AD | FX3 | FX-AC | 14342034 | 649917 | 566324 | 210490 | 24.46 | 12.86 | 3.9 | 24.71 | 24.7 | 1053 | 4.53 | 80.41 |
| CECcat03 | CEC03 | CEC03 | cat03 | Italy | Rome, S. Cecilia in Trastevere | 12-13th c AD | FX52 | FX-DB | 4781842 | 806012 | 721326 | 34957 | 4.16 | 10.51 | 15.0 | 30.47 | 27.45 | 1925 | 6.9063 | 90.66 |
| CECcat03 | CEC03b | CEC03b | cat03 | Italy | Rome, S. Cecilia in Trastevere | 12-13th c AD | FX3 | FX-AC | 16084019 | 737523 | 635092 | 219060 | 22.90 | 13.89 | 3.9 | 28.02 | 27.8 | 1324 | 5.18 | 82.62 |
| CECcat03 | CEC03a | CEC03a | cat03 | Italy | Rome, S. Cecilia in Trastevere | 12-13th c AD | FX3 | FX-AC | 12536647 | 417771 | 353231 | 130240 | 23.77 | 15.45 | 2.8 | 29.72 | 29.7 | 804 | 3.15 | 57.08 |
| CETcat01 | CET01 | CET01 | cat01 | Spain | Cova de Els Trocs, Huesca | 5300BC (9.4%) 5250BC / 5230BC (86.0%) 5040BC (datec | FX34 | FXCA | 5952893 | 4926003 | 4452949 | 271942 | 5.23 | 9.60 | 71.5 | 24.16 | 24.77 | 3619 | 13.9113 | 98.42 |
| CETcat01 | CET01 | CET01 | cat01 | Spain | Cova de Els Trocs, Huesca | 5300BC (9.4%) 5250BC / 5230BC (86.0%) 5040BC (datec | FX25 | FX-BD | 4362241 | 620656 | 538355 | 649615 | 51.14 | 13.26 | 10.7 | 27.27 | 28 | 1769 | 9.03 | 96.18 |
| CETcat01 | CET01 | CET01 | cat01 | Spain | Cova de Els Trocs, Huesca | 5300BC (9.4%) 5250BC / 5230BC (86.0%) 5040BC (datec | FX25 | FX-BD | 6300764 | 786649 | 676380 | 753054 | 48.91 | 14.02 | 9.6 | 27.2 | 18.9 | 2304 | 9.84 | 96.8 |
| CLWHcat01 | CLWH01 | CLWH01 | cat01 | Germany | Calw-Hirsau | 12th/13rd century CE | FX54 | FX-DD | 3861995 | 476375 | 429788 | 17994 | 3.64 | 9.78 | 11.1 | 23.32 | 21.06 | 1035 | 3.5235 | 66.11 |
| CLWHcat02 | CLWH02 | CLWH02 | cat02 | Germany | Calw-Hirsau | 12th/13rd century CE | FX54 | FX-DD | 11397247 | 902174 | 818892 | 73410 | 7.52 | 9.23 | 7.1 | 17.79 | 16.6 | 1632 | 5.5739 | 90.79 |
| COCcat01 | COC01 | COC01 | cat01 | Italy | Grotta del Cocci (Narni, Umbria) | 5370BC (95.4%) 5210BC | FX31 | FX-BO | 9571251 | 2463838 | 2079219 | 750192 | 23.34 | 15.61 | 20.1 | 34.34 | 30.7 | 2748 | 7.8019 | 95.82 |
| COCcat01 | COC01 | COC01 | cat01 | Italy | Grotta del Cocci (Narni, Umbria) | 5370BC (95.4%) 5210BC | FX8 | FX-AH | 14315144 | 176715 | 138526 | 62820 | 26.23 | 21.61 | 1.0 | 30.96 | 30.9 | 151 | 0.47 | 1.53 |
| CSALcat03 | CSAL03 | CSAL03 | cat03 | Portugal | Castelo de Salir, Loulé, Faro | 12th c AD | FX44 | FXCN | 6822124 | 256722 | 238435 | 21971 | 7.88 | 7.12 | 3.5 | 30.21 | 24.7 | 970 | 3.6119 | 68.34 |
| CSALcat03 | CSAL03 | CSAL03 | cat03 | Portugal | Castelo de Salir, Loulé, Faro | 12th c AD | FX29 | FX-BN | 14428833 | 221174 | 205718 | 4481 | 1.99 | 6.99 | 1.4 | 31.22 | 26.6 | 774 | 2.68 | 47.59 |
| DDcat03 | DD03 | DD03 | cat3 | Turkey | Didyma | 900AD ( 3.8%) 930AD / 940AD (73.6%) 1060AD / 1070AD | FX31 | FX-BO | 12382030 | 1044881 | 942248 | 340361 | 24.57 | 9.82 | 7.4 | 38.64 | 33.3 | 1750 | 5.41 | 86.9 |
| DDcat03 | DD03 | DD03 | cat3 | Turkey | Didyma | 900AD ( 3.8%) 930AD / 940AD (73.6%) 1060AD / 1070AD | FX17 | FX-AP | 7348145 | 146352 | 130976 | 8056 | 5.22 | 10.51 | 1.8 | 37.8 | 32.8 | 139 | 0.49 | 1.6 |
| DHcat03 | DH03 | DH03 | wildcat3 | Turkey | Demircihüyük | 3000-2500 BC | FX45 | FX-DC | 3801057 | 238426 | 192495 | 148675 | 38.41 | 19.26 | 4.9 | 37.02 | 34.81 | 121 | 0.3582 | 0.83 |
| DHcat05 | DH05 | DH05 | wildcat1 | Turkey | Demircihüyük | 2500-2000 BC | FX58 | FX-DI | 5881909 | 4060980 | 3771528 | 196332 | 4.61 | 7.13 | 62.1 | 15.58 | 11.54 | 11869 | 42.512 | 99.86 |
| DHcat06 | DH06 | DH06 | wildcat1 | Turkey | Demircihüyük | 2500-2000 BC | FX45 | FX-DC | 3183463 | 2737945 | 2484844 | 811777 | 22.87 | 9.24 | 62.2 | 18.11 | 13.94 | 6099 | 22.1845 | 99.64 |
| DHcat07 | DH07 | DH07 | wildcat5 | Turkey | Demircihüyük | 3000-2500 BC | FX45 | FX-DC | 4017916 | 8039 | 4149 | 1253 | 13.48 | 48.39 | 0.1 | 18.61 | 19.64 | 101 | 0.27 | 3.92 |
| DOLcat01 | DOL01 | DOL01 | wildcat1 | Bulgaria | Dolnoslav | 4510-4330 BC (cal AMS) | FX34 | FXCA | 4896622 | 6109 | 5126 | 265 | 4.16 | 16.09 | 0.1 | 40.83 | 40.76 | 44 | 0.1422 | 0 |
| DOLcat01 | DOL01 | DOL01 | wildcat1 | Bulgaria | Dolnoslav | 4510-4330 BC (cal AMS) | FX27 | FX-BF | 12035682 | 7442 | 5855 | 733 | 8.97 | 21.32 | 0.1 | 31.96 | 29.7 | 34 | 0.1 | 0.25 |
| DOLcat02 | DOL02 | DOL02 | cat01 | Bulgaria | Dolnoslav | Chalcolithic | FX50 | FX-CV | 5776171 | 6079 | 5229 | 293 | 4.60 | 13.98 | 0.1 | 37.4 | 37.12 | 30 | 0.0875 | 0 |
| DOLcat03 | DOL03 | DOL03 | cat02 (exDOLcat0 | Bulgaria | Dolnoslav | Chalcolithic | FX50 | FX-CV | 6507576 | 6218 | 5118 | 638 | 9.31 | 17.69 | 0.1 | 35.64 | 33.12 | 30 | 0.0852 | 0 |
| DOScat01 | DOS01 | DOS01 | wild1 | Italy | Galgenbühel/Dos de la Forca-Salorno | 8240BC (95.4%) 7940BC | FX42 | FX-CI | 5036671 | 884376 | 790497 | 135461 | 13.28 | 10.62 | 15.3 | 23.97 | 19.53 | 799 | 2.8 | 51.72 |
| DOScat01 | DOS01 | DOS01 | wild1 | Italy | Galgenbühel/Dos de la Forca-Salorno | 8240BC (95.4%) 7940BC | FX25 | FX-BD | 9388414 | 686157 | 626639 | 96277 | 12.30 | 8.67 | 6.6 | 22.3 | 15.6 | 450 | 1.82 | 27.02 |
| DOScat03 | DOS03 | DOS03 | wild3 | Italy | Galgenbühel/Dos de la Forca-Salorno | 8,500-7,500 cal BC | FX42 | FX-CI | 6842851 | 5888715 | 5404520 | 306502 | 4.95 | 8.22 | 75.6 | 24.76 | 18.94 | 3706 | 13.6016 | 98.88 |
| DOScat03 | DOS03 | DOS03 | wild3 | Italy | Galgenbühel/Dos de la Forca-Salorno | 8,500-7,500 cal BC | FX25 | FX-BD | 8868990 | 6206918 | 5701869 | 404294 | 6.12 | 8.14 | 61.5 | 26.1 | 20.2 | 4108 | 16.189 | 98.94 |
| DURcat01 | DUR01 | DUR01 | wildcat1 | Bulgaria | Durankulak | 4450BC (95.4%) 4330BC | FX35 | FXCB | 5598064 | 3297899 | 2959379 | 200167 | 5.72 | 10.26 | 51.0 | 40.81 | 41.67 | 5292 | 16.9163 | 99.51 |
| DURcat02 | DUR02 | DUR02 | wildcat2 | Bulgaria | Durankulak | Aeneolithic 4200-4100 cal BC | FX35 | FXCB | 6929461 | 4750680 | 4315155 | 161990 | 3.30 | 9.17 | 60.9 | 44.63 | 45.47 | 25597 | 91.0273 | 99.98 |
| DURcat03 | DUR03 | DUR03 | wildcat3 | Bulgaria | Durankulak | Aeneolithic 4200-4100 cal BC | FX35 | FXCB | 5971551 | 4605496 | 4108664 | 558929 | 10.82 | 10.79 | 62.9 | 42.96 | 43.78 | 44475 | 150.3814 | 99.98 |
| DURcat06 | DUR06 | DUR06 | cat2 | Bulgaria | Durankulak | Aeneolithic 4200-4100 cal BC | FX35 | FXCB | 3528306 | 1196591 | 1104514 | 27662 | 2.26 | 7.69 | 31.1 | 8.75 | 9.1 | 3559 | 11.6544 | 99.72 |
| DURcat07 | DUR07 | DUR07 | wildcat5 | Bulgaria | Durankulak | Aeneolithic 4200-4100 cal BC | FX35 | FXCB | 5665278 | 4425192 | 4044336 | 440929 | 9.06 | 8.61 | 66.2 | 30.94 | 31.83 | 2013 | 6.3597 | 89.81 |
| ECLYcat01 | ECLY01 | ECLY01 | cat01 | France | Ecly | 20AD (2.1%) 50AD / 60AD (93.3%) 210AD | FX35 | FXCB | 7177745 | 5909649 | 5476756 | 204266 | 3.34 | 7.33 | 74.2 | 30.57 | 31.34 | 27552 | 89.2556 | 99.89 |
| FLUMcat02 | FLUM02 | FLUM02 | cat02 | Italy | nuraghe Flumenelongu, Sardinia | roman imperial | FX31 | FX-BO | 11942596 | 2278041 | 2058428 | 353201 | 13.42 | 9.64 | 16.7 | 22.72 | 21.6 | 2504 | 8.19 | 97.92 |
| FLUMcat02 | FLUM02 | FLUM02 | cat02 | Italy | nuraghe Flumenelongu, Sardinia | roman imperial | FX28 | FX-BI | 10482364 | 489253 | 440229 | 15196 | 3.01 | 10.02 | 4.2 | 27.1 | 27.5 | 661 | 2.18 | 36.42 |
| GAOcat11 | GAO11 | GAO11 | cat01 | Italy | Grotta dell'Onda | 40 Kya | FX46 | FXCO | 4789475 | 7276 | 4825 | 3759 | 34.06 | 33.69 | 0.1 | 29.71 | 26.39 | 36 | 0.096 | 0.94 |
| GAOcat11 | GAO11 | GAO11 | cat01 | Italy | Grotta dell'Onda | 40 Kya | FX12 | FX-AL | 13975734 | 3947 | 2502 | 1104 | 21.86 | 36.61 | 0.0 | 23.34 | 22.7 | 14 | 0.04 | 0.09 |
| GAOcat13 | GAO13 | GAO13 | cat02 | Italy | Grotta dell'Onda | 40 Kya | FX46 | FXCO | 8644015 | 18243 | 12377 | 5658 | 23.67 | 32.15 | 0.1 | 16.01 | 15.1 | 153 | 0.4474 | 6.05 |
| GAOcat13 | GAO13 | GAO13 | cat02 | Italy | Grotta dell'Onda | 40 Kya | FX12 | FX-AL | 12989498 | 1166 | 655 | 647 | 35.69 | 43.83 | 0.0 | 3.48 | 3.8 | 0 | 0 | 0 |
| GBcat01 | GB01 | GB01 | cat01 | Italy | Grotta Breuil (Circeo, Latium) | middle Palaeolithic | FX4 | FX-AD | 21886814 | 963 | 428 | 5488 | 85.07 | 55.56 | 0.0 | 12.08 | 53.2 | 1 | 0 | 0 |
| GLCAcat01 | GLCA01 | GLCA01 | cat01 | Ireland | Glencurran Cave, Co. Clare | 4778±24 BP (UBA-45133), 3634-3526 cal BC (Neolithic) | FX50 | FX-CV | 6610551 | 5718412 | 5321595 | 511526 | 8.21 | 6.94 | 74.7 | 28.93 | 24.1 | 14432 | 48.7986 | 99.9 |
| GMcat01 | GM01 | GM01 | cat01 | Italy | Grotta Maritza (Fucino, Abruzzi) | 10-11 ka BP uncal / LUP (B2) | FX9 | FX-AI | 13163666 | 5216 | 4331 | 2736 | 34.41 | 16.97 | 0.0 | 37.82 | 39.3 | 9 | 0.04 | 0 |
| GMcat02 | GM02 | GM02 | cat02 | Italy | Grotta Maritza (Fucino, Abruzzi) | 11-11 ka BP uncal / LUP (B3) | FX4 | FX-AD | 13981619 | 43134 | 39344 | 8807 | 16.96 | 8.79 | 0.3 | 30.96 | 31.2 | 31 | 0.14 | 0.14 |
| GMcat03 | GM03 | GM03 | cat03 | Italy | Grotta Maritza (Fucino, Abruzzi) | 12-11 ka BP uncal / LUP (B3) | FX4 | FX-AD | 15464186 | 142809 | 123741 | 54575 | 27.65 | 13.35 | 0.8 | 30.3 | 30.5 | 314 | 0.94 | 8.12 |
| GMcat04 | GM04 | GM04 | cat04 | Italy | Grotta Maritza (Fucino, Abruzzi) | 13-11 ka BP uncal / LUP (B4-B3) | FX4 | FX-AD | 19310608 | 64089 | 50877 | 192703 | 75.04 | 20.62 | 0.3 | 35.96 | 36 | 171 | 0.64 | 3.23 |
| GMcat05 | GM05 | GM05 | cat05 | Italy | Grotta Maritza (Fucino, Abruzzi) | 14-11 ka BP uncal / LUP (B4-B3) | FX4 | FX-AD | 15104752 | 244613 | 198874 | 295629 | 54.72 | 18.70 | 1.3 | 29.08 | 28.9 | 709 | 2.57 | 46.12 |
| GMcat06 | GM06 | GM06 | cat06 | Italy | Grotta Maritza (Fucino, Abruzzi) | 15-11 ka BP uncal / LUP (B4-B3) | FX4 | FX-AD | 16541053 | 225727 | 169933 | 594170 | 72.47 | 24.72 | 1.0 | 33.29 | 32.7 | 2290 | 8.42 | 95.66 |
| GRAcat01 | GRA01 | GRA01 | cat01 | Italy | Graffignano (Viterbo, Latium) | Late 15th-Early 16th cent AD | FX4 | FX-AD | 11938945 | 8694784 | 7944434 | 2784667 | 24.26 | 8.63 | 54.0 | 21.9 | 21.8 | 22581 | 100.1189 | 100 |
| GSAcat01 | GSA01 | GSA01 | cat01 | Italy | Genoni Santu Antine, Sardinia | 340BC (3.9%) 320BC / 200BC (91.5%) 50BC | FX28 | FX-BI | 9310555 | 2714715 | 2388164 | 837165 | 23.57 | 12.03 | 23.5 | 24.56 | 25.2 | 5707 | 22.0618 | 98.98 |
| HAIScat01 | HAIS01 | HAIS01 | cat01 | Germany | Haithabu, settlement area | 800-1050 AD | FX41 | FX-CH | 8644279 | 3273440 | 2927697 | 295968 | 8.29 | 10.56 | 32.8 | 24.87 | 24.03 | 6566 | 20.9234 | 99.46 |
| HAIScat01 | HAIS01 | HAIS01 | cat01 | Germany | Haithabu, settlement area | 800-1050 AD | FX26 | FX-BG | 10550443 | 1333909 | 1203077 | 116370 | 8.02 | 9.81 | 11.3 | 24.96 | 25.2 | 3033 | 11.3 | 98.76 |
| HAIScat03 | HAIS03 | HAIS03 | cat03 | Germany | Haithabu, settlement area | 800-1050 AD | FX41 | FX-CH | 9523842 | 1153262 | 1033335 | 47996 | 4.00 | 10.40 | 10.8 | 21.55 | 21.68 | 6507 | 21.0023 | 99.57 |
| HAIScat03 | HAIS03 | HAIS03 | cat03 | Germany | Haithabu, settlement area | 800-1050 AD | FX26 | FX-BG | 10964830 | 407625 | 367034 | 10010 | 2.40 | 9.96 | 3.3 | 21.95 | 22 | 3006 | 11.08 | 98.77 |
| HAIScat04 | HAIS04 | HAIS04 | cat04 | Germany | Haithabu, settlement area | 800-1050 AD | FX52 | FX-DB | 4972434 | 931511 | 830850 | 48182 | 4.92 | 10.81 | 16.6 | 12.99 | 12.75 | 5094 | 18.969 | 99.57 |
| HAIScat04 | HAIS04 | HAIS04 | cat04 | Germany | Haithabu, settlement area | 800-1050 AD | FX26 | FX-BG | 10940755 | 1025339 | 912557 | 38982 | 3.66 | 11.00 | 8.3 | 15.47 | 15.8 | 5459 | 19.79 | 99.96 |
| HAIScat10 | HAIS10 | HAIS10 | cat10 | Germany | Haithabu, settlement area | 800-1050 AD | FX52 | FX-DB | 4337691 | 605839 | 530624 | 64865 | 9.67 | 12.42 | 12.1 | 18.98 | 19.31 | 4579 | 15.7324 | 99.29 |
| HAIScat10 | HAIS10 | HAIS10 | cat10 | Germany | Haithabu, settlement area | 800-1050 AD | FX30 | FX-BL | 12073921 | 1377777 | 1220702 | 122303 | 8.15 | 11.40 | 10.0 | 20.06 | 21.4 | 8705 | 28.87 | 99.95 |
| HDRcat01 | HDR01 | HDR01 | cat01 | Germany | Hochdorf-Reps | 450 to 250 BCE (La Tène Period Early) | FX54 | FX-DD | 12017105 | 95151 | 86278 | 3179 | 3.23 | 9.33 | 0.7 | 6.25 | 6.04 | 304 | 1.0012 | 9.5 |
| HDRcat02 | HDR02 | HDR02 | cat02 | Germany | Hochdorf-Reps | 450 to 250 BCE (La Tène Period Early) | FX54 | FX-DD | 95229918 | 1598991 | 1429602 | 411883 | 20.48 | 10.59 | 1.5 | 29.87 | 24.56 | 6268 | 23.8145 | 99.18 |
| HTBcat01 | HTB01SS | HTB01SS | cat01 | Austria | Hinterbrühl | Holocene unspecified | FX14 | FX-BM | 6584756 | 192995 | 173488 | 2843 | 1.45 | 10.11 | 2.6 | 9.67 | 1.3 | 506 | 1.56 | 21.35 |
| HTBcat01 | HTB01 | HTB01 | cat01 | Austria | Hinterbrühl | Holocene unspecified | FX14 | FX-AM | 11484514 | 181637 | 144504 | 331774 | 64.62 | 20.44 | 1.2 | 16.55 | 16.6 | 330 | 1.65 | 22.98 |
| IRTcat01 | IRT01 | IRT01 | cat01 | Turkey | Iznik - Roman theatre | Roman-Byzantine? | FX50 | FX-CV | 4908068 | 2055334 | 1882275 | 41729 | 1.99 | 8.42 | 38.0 | 23.86 | 19.43 | 1711 | 6.2238 | 89.68 |
| IRTcat02 | IRT02 | IRT02 | cat01? | Turkey | Iznik - Roman theatre | Roman-Byzantine? | FX50 | FX-CV | 4590725 | 475953 | 437223 | 17716 | 3.59 | 8.14 | 9.5 | 29.08 | 22.01 | 2822 | 10.9342 | 98.33 |
| KAcat01 | KA01 | KA01 | cat1 | Greece | Kastanas | ca. 1250 BC | FX27 | FX-BF | 10073281 | 707529 | 601224 | 109639 | 13.42 | 15.02 | 5.9 | 27.93 | 26.2 | 1304 | 4.09 | 72.8 |
| KAcat01 | KA01 | KA01 | cat1 | Greece | Kastanas | ca. 1250 BC | FX41 | FX-CH | 7173301 | 287039 | 244119 | 50789 | 15.03 | 14.95 | 3.4 | 27.88 | 26.25 | 777 | 2.4225 | 41.75 |
| KAcat02 | KA02 | KA02 | cat2 | Greece | Kastanas | ca. 1250 BC | FX42 | FX-CI | 6649958 | 66935 | 55138 | 19943 | 22.96 | 17.62 | 0.8 | 32.12 | 32.33 | 81 | 0.2588 | 0.09 |
| KAcat02 | KA02 | KA02 | cat2 | Greece | Kastanas | ca. 1250 BC | FX27 | FX-BF | 11153009 | 70459 | 58425 | 18876 | 21.13 | 17.08 | 0.5 | 26.57 | 24.6 | 100 | 0.32 | 0.05 |
| KAcat03 | KA03T | KA03T | cat3 | Greece | Kastanas | ca 200-400 BC | FX27 | FX-BF | 11753392 | 20002 | 15515 | 2440 | 10.87 | 22.43 | 0.1 | 21.57 | 21.8 | 66 | 0.21 | 0.05 |
| KAcat06 | KA06 | KA06 | wildcat2 | Greece | Kastanas | late phase of the Late Bronze Age (c. 1300 BC) | FX27 | FX-BF | 11067015 | 60936 | 49990 | 2574 | 4.05 | 17.96 | 0.5 | 30.19 | 29.1 | 183 | 0.56 | 2.7 |
| KAcat06 | KA06 | KA06 | wildcat2 | Greece | Kastanas | late phase of the Late Bronze Age (c. 1300 BC) | FX41 | FXCH | 7786666 | 19536 | 16465 | 578 | 2.87 | 15.72 | 0.2 | 32.55 | 30.26 | 41 | 0.1333 | 0.25 |
| KASPcat01 | KASP01 | KASP01 | wildcat1 | Greece | Kassope | 400BC (33.6%) 340BC / 310BC (61.8%) 200BC | FX32 | FXBS | 6379764 | 4879180 | 4442392 | 123584 | 2.47 | 8.95 | 68.3 | 29.78 | 30.6 | 4207 | 13.8376 | 97.51 |
| KASPcat02 | KASP02 | KASP02 | wildcat2 | Greece | Kassope | 4th-1st cent BC | FX32 | FXBS | 4635753 | 1050099 | 938158 | 67616 | 6.05 | 10.66 | 20.0 | 34.11 | 34.78 | 1796 | 5.7222 | 88.55 |
| KOPcat02 | KOP02 | KOP02 | wildcat2 | Bulgaria | Koprivec | Neolithic B 5800-4800 cal BC | FX34 | FXCA | 5331330 | 588874 | 525558 | 79086 | 11.84 | 10.75 | 9.7 | 32.51 | 33.02 | 480 | 1.4233 | 20.5 |
| KOPcat02 | KOP02_SS | KOP02_SS | wildcat2 | Bulgaria | Koprivec | Neolithic B 5800-4800 cal BC | FX25 | FX-BM | 9006273 | 138831 | 119369 | 2242 | 1.59 | 14.02 | 1.3 | 20.46 | 1.2 | 125 | 0.34 | 1.61 |
| KOPcat02 | KOP02 | KOP02 | wildcat2 | Bulgaria | Koprivec | Neolithic B 5800-4800 cal BC | FX25 | FX-BD | 10165025 | 57352 | 50922 | 3250 | 5.36 | 11.21 | 0.5 | 29.8 | 22.4 | 41 | 0.17 | 0.39 |
| KOPcat03 | KOP03 | KOP03 | wildcat3 | Bulgaria | Koprivec | 6050-5900 BC (cal AMS) | FX34 | FXCA | 4982334 | 335121 | 298168 | 25813 | 7.15 | 11.03 | 6.0 | 47.6 | 47.73 | 2778 | 8.6757 | 97.09 |
| KOPcat03 | KOP03 | KOP03 | wildcat3 | Bulgaria | Koprivec | 6050-5900 BC (cal AMS) | FX25 | FX-BD | 8535402 | 5896 | 4915 | 632 | 9.68 | 16.64 | 0.1 | 39 | 29.1 | 38 | 0.16 | 0.2 |
| KOPcat04 | KOP04 | KOP04 | wildcat4 | Bulgaria | Koprivec | 5480BC (95.4%) 5360BC | FX34 | FXCA | 4242966 | 1865118 | 1497628 | 341097 | 15.46 | 19.70 | 32.7 | 30.69 | 31.43 | 4178 | 13.1734 | 99.07 |
| KOPcat04 | KOP04 | KOP04 | wildcat4 | Bulgaria | Koprivec | 5480BC (95.4%) 5360BC | FX25 | FX-BD | 5635784 | 205005 | 154531 | 18751 | 8.38 | 24.62 | 2.7 | 33.9 | 26 | 885 | 3.63 | 67.87 |
| LABcat01 | LAB01 | LAB01 | cat01 | Spain | Labastida | end 15th-early 16th c. | FX46 | FX-CO | 7699429 | 6838564 | 6201637 | 1044081 | 13.25 | 9.31 | 70.9 | 14.48 | 10.65 | 10642 | 40.2702 | 99.89 |
| LABcat01 | LAB01 | LAB01 | cat01 | Spain | Labastida | end 15th-early 16th c. | FX29 | FX-BN | 11040026 | 6786613 | 6175272 | 479314 | 6.60 | 9.01 | 53.6 | 17.14 | 14.4 | 24124 | 88.4217 | 99.99 |
| LATcat02 | LAT02_SS | LAT02_SS | cat02? | Italy | Latronico 3, Basilicata | mesolithic; 7420+-90 BP | FX12 | FX-BC | 6084082 | 33874 | 30572 | 1320 | 3.75 | 9.75 | 0.5 | 41.5 | 1.4 | 58 | 0.16 | 0 |
| LATcat02 | LAT02 | LAT02 | cat02? | Italy | Latronico 3, Basilicata | mesolithic; 7420+-90 BP | FX12 | FX-AL | 9859775 | 20032 | 17170 | 3513 | 14.92 | 14.29 | 0.2 | 48.25 | 50.8 | 36 | 0.13 | 0.13 |
| LATcat03 | LAT03 | LAT03 | cat03 | Italy | Latronico 3, Basilicata | mesolithic | FX5 | FX-AE | 25172625 | 2507053 | 2130563 | 1112704 | 30.74 | 15.02 | 8.1 | 34.58 | 33.3 | 2768 | 10.84 | 97.08 |
| LATcat04 | LAT04 | LAT04 | cat04? | Italy | Latronico 3, Basilicata | mesolithic | FX46 | FXCO | 4111798 | 4874 | 3616 | 2109 | 30.20 | 25.81 | 0.1 | 45.52 | 36.5 | 17 | 0.0553 | 0 |
| LATcat04 | LAT04 | LAT04 | cat04? | Italy | Latronico 3, Basilicata | mesolithic | FX12 | FX-AL | 12654029 | 5228 | 2846 | 2891 | 35.61 | 45.56 | 0.0 | 26.37 | 27.1 | 9 | 0.03 | 0 |
| LATcat05 | LAT05 | LAT05 | cat05 | Italy | Latronico 3, Basilicata | mesolithic | FX5 | FX-AE | 12762783 | 3281 | 2346 | 1017 | 23.66 | 28.50 | 0.0 | 33.15 | 34.8 | 3 | 0.01 | 0 |
| LATcat06 | LAT06 | LAT06 | cat06 | Italy | Latronico 3, Basilicata | mesolithic | FX5 | FX-AE | 11523954 | 50558 | 44274 | 38552 | 43.26 | 12.43 | 0.4 | 48.5 | 47.3 | 89 | 0.31 | 0.49 |
| LATcat07 | LAT07 | LAT07 | cat07 | Italy | Latronico 3, Basilicata | mesolithic | FX5 | FX-AE | 17734998 | 2793 | 2323 | 332 | 10.62 | 16.83 | 0.0 | 18.01 | 20.1 | 4 | 0.02 | 0 |
| LATcat08 | LAT08 | LAT08 | cat08 | Italy | Latronico 3, Basilicata | mesolithic | FX5 | FX-AE | 21510177 | 9749 | 8067 | 4379 | 31.00 | 17.25 | 0.0 | 46.25 | 42.8 | 8 | 0.02 | 0 |
| LATcat09 | LAT09 | LAT09 | cat09 | Italy | Latronico 3, Basilicata | mesolithic | FX5 | FX-AE | 22187207 | 17559 | 15233 | 8077 | 31.51 | 13.25 | 0.1 | 40.31 | 39.9 | 13 | 0.04 | 0 |
| LHMcat01 | LHM01 | LHM01 | cat01 | Germany | Lauchheim-Mittelhofen | 6th/7th century CE | FX52 | FX-DB | 4153863 | 8153 | 7310 | 96 | 1.16 | 10.34 | 0.2 | 19.49 | 16.68 | 9 | 0.0332 | 0 |
| LHMcat02 | LHM02 | LHM02 | cat02 | Germany | Lauchheim-Mittelhofen | 8th/9th century CE | FX52 | FX-DB | 4968307 | 1298609 | 1191657 | 27911 | 2.10 | 8.24 | 23.9 | 29.36 | 26.42 | 6625 | 22.3108 | 99.58 |
| LHMcat03 | LHM03 | LHM03 | cat03 | Germany | Lauchheim-Mittelhofen | 8th/9th century CE | FX52 | FX-DB | 4516835 | 743520 | 679322 | 20165 | 2.64 | 8.63 | 15.0 | 26.94 | 20.02 | 1940 | 7.4093 | 96.1 |
| MARcat01 | MAR01 | MAR01 | cat01 | Italy | La Marmotta (Bracciano Lake, Latium) | Middle Neolithic | FX9 | FX-AI | 13401889 | 1339 | 603 | 420 | 23.88 | 54.97 | 0.0 | 4.47 | 5.8 | 0 | 0 | 0 |
| MENTcat01 | MENT01 | MENT01 | wild? cat01 | Turkey | Menteşe | 6200-6100 cal. BC | FX56 | FX-DE | 4473455 | 336033 | 278511 | 10681 | 3.08 | 17.12 | 6.2 | 42.11 | 39.63 | 1207 | 3.7592 | 69.81 |
| MLECcat01 | MLEC01 | MLEC01 | cat01 | Italy | Muro Leccese | 16-17th c AD | FX6 | FX-AF | 12283595 | 198045 | 125034 | 1653817 | 89.31 | 36.87 | 0.9 | 15.41 | 15.7 | 97 | 0.38 | 1.19 |
| MLECcat02 | MLEC02 | MLEC02 | cat02 | Italy | Muro Leccese | 18th c AD | FX6 | FX-AF | 13911019 | 920467 | 838157 | 487389 | 34.62 | 8.94 | 5.8 | 18.97 | 18.9 | 2277 | 9.11 | 98.44 |
| MLECcat05 | MLEC5b | MLEC5b | cat03 | Italy | Muro Leccese | 16th c AD | FX1 | FX-AA | 14921652 | 5434 | 3383 | 1271 | 18.96 | 37.74 | 0.0 | 9.26 | 11 | 1 | 0 | 0 |
| MOScat01 | MOS01 | MOS01 | cat01 | Portugal | Moita do Sebastião (Muge) | mesolithic | FX12 | FX-AL | 13883197 | 2463 | 756 | 1230 | 33.31 | 69.31 | 0.0 | 10.19 | 7.7 | 2 | 0.01 | 0 |
| MOScat01 | MOS01_SS | MOS01_SS | cat01 | Portugal | Moita do Sebastião (Muge) | mesolithic | FX12 | FX-BE | 9174149 | 2144 | 1137 | 543 | 20.21 | 46.97 | 0.0 | 7.2 | 3.6 | 1 | 0 | 0 |
| MOScat02 | MOS02 | MOS02 | cat02 | Portugal | Moita do Sebastião (Muge) | mesolithic | FX6 | FX-AF | 10344126 | 6618 | 4794 | 14092 | 68.04 | 27.56 | 0.1 | 33.2 | 32.2 | 30 | 0.1 | 0 |
| MOScat03 | MOS03 | MOS03 | cat03 | Portugal | Moita do Sebastião (Muge) | mesolithic | FX6 | FX-AF | 14473196 | 849 | 668 | 293 | 25.66 | 21.32 | 0.0 | 18.52 | 23.5 | 2 | 0.01 | 0 |
| MRYcat07 | MRY07 | MRY07 | cat7 | Turkey | Yenikapı, Marmaray exc. | 4th-14th cent AD? | FX33 | FXBZ | 6031746 | 3679347 | 3376868 | 90694 | 2.41 | 8.22 | 55.2 | 32.17 | 32.98 | 4314 | 14.9447 | 98.68 |
| MRYcat08 | MRY08 | MRY08 | cat8 | Turkey | Yenikapı, Marmaray exc. | 4th-14th cent AD? | FX33 | FXBZ | 5826906 | 3907858 | 3548427 | 189860 | 4.63 | 9.20 | 59.0 | 40.64 | 41.49 | 5319 | 17.2306 | 99.09 |
| MRYcat09 | MRY09 | MRY09 | cat9 | Turkey | Yenikapı, Marmaray exc. | 650AD (48.9%) 710AD / 720AD (46.5%) 780AD (dated twi | FX17 | FX-AP | 6198276 | 1111159 | 989556 | 128803 | 10.39 | 10.94 | 15.6 | 19.54 | 20.2 | 583 | 2.21 | 36.26 |
| MRYcat09 | MRY09 | MRY09 | cat9 | Turkey | Yenikapı, Marmaray exc. | 650AD (48.9%) 710AD / 720AD (46.5%) 780AD (dated twi | FX17 | FX-AP | 7012917 | 1224709 | 1088265 | 132106 | 9.74 | 11.14 | 15.2 | 19.7 | 16.9 | 695 | 2.48 | 41.77 |
| MRYcat10 | MRY10 | MRY10 | cat10 | Turkey | Yenikapı, Marmaray exc. | 640AD (75.4%) 690AD / 740AD (20.0%) 780AD | FX33 | FXBZ | 5526064 | 4429215 | 4099442 | 99754 | 2.20 | 7.45 | 72.9 | 36.93 | 37.9 | 5811 | 21.8153 | 99.81 |
| MTRcat01 | MTR01B | MTR01B | cat01 | Austria | Mautern - vicus Ost | 50BC (95.4%) 80AD | FX22 | FX-AZ | 6277438 | 5094545 | 4572503 | 1136596 | 18.24 | 10.25 | 61.7 | 25.29 | 25.5 | 11423 | 46.9017 | 99.77 |
| MTRcat01 | MTR01T | MTR01T | cat01 | Austria | Mautern - vicus Ost | 50BC (95.4%) 80AD | FX22 | FX-AZ | 8853383 | 85407 | 75273 | 26299 | 23.54 | 11.87 | 0.9 | 30.36 | 30 | 460 | 1.8 | 27.31 |
| MUScat01 | MUS01 | MUS01 | cat01 | Italy | Musarna (Viterbo, Latium) | 360AD (95.4%) 540AD | FX46 | FXCO | 7420936 | 2562747 | 2161388 | 1619039 | 38.72 | 15.66 | 23.9 | 15.76 | 14.27 | 3569 | 12.6494 | 99.22 |
| MUScat01 | MUS01 | MUS01 | cat01 | Italy | Musarna (Viterbo, Latium) | 360AD (95.4%) 540AD | FX14 | FX-AM | 10264249 | 310343 | 221282 | 401313 | 56.39 | 28.70 | 2.1 | 14.7 | 15.1 | 398 | 1.78 | 26.09 |
| NBLcat01 | NBL01 | NBL01 | cat01 | Austria | Nebellöcher, Baden district | Holocene unspecified, prob. medieval/modern | FX14 | FX-AM | 7949619 | 911093 | 727400 | 805021 | 46.91 | 20.16 | 8.3 | 6.06 | 6.3 | 1480 | 5.63 | 79.58 |
| NBLcat02 | NBL02 | NBL02 | cat02 | Austria | Nebellöcher, Baden district | Holocene unspecified, prob. medieval/modern | FX14 | FX-AM | 9704757 | 363305 | 276451 | 921508 | 71.72 | 23.91 | 2.6 | 8.34 | 8.4 | 625 | 3.06 | 56.61 |
| PADcat01 | PAD01 | PAD01 | cat01 | Italy | Padova Via Cesare Battisti (Veneto) | 1400AD (95.4%) 1440AD | FX42 | FX-CI | 8791303 | 1370055 | 1244327 | 52675 | 3.70 | 9.18 | 14.1 | 19.39 | 16.53 | 892 | 3.1897 | 58.94 |
| PADcat01 | PAD01 | PAD01 | cat01 | Italy | Padova Via Cesare Battisti (Veneto) | 1400AD (95.4%) 1440AD | FX12 | FX-AL | 16304996 | 12587 | 8790 | 5888 | 31.87 | 30.17 | 0.1 | 17.99 | 19.4 | 39 | 0.19 | 0 |
| PALcat08 | PAL08 | PAL08 | cat08 | Portugal | Palmela Castle | 12-13th c | FX56 | FX-DE | 3548879 | 10816 | 9848 | 164 | 1.49 | 8.95 | 0.3 | 21.48 | 18.51 | 20 | 0.0687 | 0 |
| PALcat09 | PAL09 | PAL09 | cat09 | Portugal | Palmela Castle | 12-13th c | FX56 | FX-DE | 3992598 | 944516 | 859313 | 20317 | 2.11 | 9.02 | 21.4 | 20.99 | 18.08 | 1797 | 6.4876 | 94.08 |
| PALcat10 | PAL10 | PAL10 | cat10 | Portugal | Palmela Castle | 11-12th c | FX56 | FX-DE | 3334705 | 542223 | 493059 | 12699 | 2.29 | 9.07 | 14.7 | 26.49 | 20.92 | 1245 | 4.6424 | 82.97 |
| PAZcat01 | PAZ01B_SS | PAZ01B_SS | cat01 | Italy | Sassari, pozzo di Largo Pazzola, Sardinia | 16th c AD | FX21 | FX-BM | 5158531 | 53906 | 48221 | 783 | 1.43 | 10.55 | 0.9 | 24.3 | 0.9 | 73 | 0.2 | 0.02 |
| PAZcat01 | PAZ01B | PAZ01B | cat01 | Italy | Sassari, pozzo di Largo Pazzola, Sardinia | 16th c AD | FX21 | FX-AT | 8940269 | 47102 | 42394 | 9310 | 16.50 | 10.00 | 0.5 | 30.49 | 29.6 | 52 | 0.18 | 0.33 |
| PENcat01 | PEN01 | PEN01 | cat01 | Spain | Peñalosa, Jaén | Bronze Age | FX6 | FX-AF | 482 | 0 | 0 | 0 | / | / | 0.0 | 0 | 0 | 0 | 0 | 0 |
| PENcat02 | PEN02 | PEN02 | cat02 | Spain | Peñalosa, Jaén | Bronze Age | FX18 | FX-AR | 9714743 | 3083 | 1615 | 548 | 15.09 | 47.62 | 0.0 | 8.63 | 10.4 | 27 | 0.1 | 0 |
| PENcat04 | PEN04 | PEN04 | cat04 | Spain | Peñalosa, Jaén | Bronze Age | FX18 | FX-AR | 9310671 | 3146 | 2608 | 120 | 3.67 | 17.10 | 0.0 | 13.98 | 15.9 | 2 | 0.01 | 0 |
| PETRcat03 | PETR03 | PETR03 | cat01 | Austria | Petronell-Carnuntum | Roman | FX14 | FX-AM | 10447417 | 353356 | 294883 | 139469 | 28.30 | 16.55 | 2.8 | 21.07 | 21 | 1412 | 7.48 | 93.54 |
| PETRcat04 | PETR04 | PETR04 | cat02 | Austria | Petronell-Carnuntum | Roman | FX16 | FX-AN | 6375121 | 113059 | 66662 | 997861 | 89.82 | 41.04 | 0.9 | 17.52 | 17.5 | 398 | 1.82 | 26.2 |
| PETRcat06 | PETR06 | PETR06 | cat03 | Austria | Petronell-Carnuntum | 120AD (87.6%) 260AD / 290AD (7.8%) 320AD | FX16 | FX-AN | 8515973 | 5530866 | 4991148 | 1977248 | 26.33 | 9.76 | 47.6 | 17.1 | 16.8 | 2849 | 10.4948 | 94.9 |
| PETRcat07 | PETR07B_SS | PETR07B_SS | cat04 | #N/A | #N/A | Roman | FX22 | FX-BC | 6653747 | 573460 | 525713 | 9023 | 1.55 | 8.33 | 7.9 | 18.7 | 1.3 | 1280 | 3.84 | 71.12 |
| PETRcat07 | PETR07B | PETR07B | cat04 | Austria | Petronell-Carnuntum | Roman | FX22 | FX-AZ | 9992573 | 615108 | 556702 | 141365 | 18.69 | 9.50 | 5.5 | 25.71 | 24.4 | 1102 | 4.41 | 76.97 |
| PETRcat07 | PETR07T | PETR07T | cat04 | Austria | Petronell-Carnuntum | Roman | FX22 | FX-AZ | 11554418 | 11017 | 5847 | 2233 | 16.85 | 46.93 | 0.1 | 15.92 | 17.8 | 10 | 0.04 | 0 |
| PETRcat08 | PETR08B | PETR08B | cat05 | Austria | Petronell-Carnuntum | Roman | FX22 | FX-AZ | 8146315 | 5010139 | 4497016 | 1515887 | 23.23 | 10.24 | 46.5 | 22.64 | 22.6 | 8845 | 32.7489 | 99.91 |
| PETRcat08 | PETR08T | PETR08T | cat05 | Austria | Petronell-Carnuntum | Roman | FX22 | FX-AZ | 10879031 | 97130 | 84573 | 20029 | 17.10 | 12.93 | 0.8 | 21.53 | 21 | 103 | 0.36 | 0.44 |
| PGLcat01 | PGL01 | PGL01 | cat01 | Italy | Grotta Paglicci (Rignano Garganico - Foggia) | 11.440 - 15.320 BP | FX9 | FX-AI | 16349118 | 289 | 187 | 57 | 16.47 | 35.29 | 0.0 | 6.52 | 6.8 | 1 | 0 | 0 |
| PGLcat02 | PGL02 | PGL02 | cat02 | Italy | Grotta Paglicci (Rignano Garganico - Foggia) | 11.440 - 15.320 BP | FX9 | FX-AI | 10205674 | 4148 | 2212 | 25538 | 86.03 | 46.67 | 0.0 | 25.9 | 19.3 | 2 | 0.01 | 0 |
| PGLcat03 | PGL03 | PGL03 | cat03 | Italy | Grotta Paglicci (Rignano Garganico - Foggia) | 11.440 - 15.320 BP | FX9 | FX-AI | 13822663 | 2181 | 1720 | 2656 | 54.91 | 21.14 | 0.0 | 31.88 | 33.3 | 1 | 0 | 0 |
| PGLcat04 | PGL04 | PGL04 | cat04 | Italy | Grotta Paglicci (Rignano Garganico - Foggia) | 11.440 - 15.320 BP | FX9 | FX-AI | 11893850 | 74126 | 51728 | 223805 | 75.12 | 30.22 | 0.4 | 41.39 | 41.8 | 31 | 0.1 | 0 |
| PIAcat01 | PIA01 | PIA01 | cat01 | Italy | Pianosa | 650-770 AD (cal AMS) | FX17 | FX-AP | 6560499 | 60141 | 52780 | 11482 | 16.03 | 12.24 | 0.8 | 35.2 | 30.1 | 594 | 2.29 | 41.48 |
| PNEcat01 | PNE01 | PNE01 | cat01 | Italy | Palatino North East slope (Rome) | 2nd half 6th cent AD (Roman) | FX54 | FX-DD | 8875706 | 32621 | 29220 | 809 | 2.42 | 10.43 | 0.3 | 21.52 | 18.88 | 51 | 0.1756 | 0 |
| PNEcat02 | PNE02 | PNE02 | cat02 | Italy | Palatino North East slope (Rome) | 2nd half 5th cent AD (Roman) | FX54 | FX-DD | 9353308 | 1460146 | 1337834 | 62187 | 4.08 | 8.38 | 14.2 | 33.53 | 29.47 | 4887 | 16.806 | 99.61 |
| PNEcat03 | PNE03 | PNE03 | cat03 | Italy | Palatino North East slope (Rome) | End 4th cent.-beg. 5th cent AD (Roman) | FX54 | FX-DD | 4036275 | 3374207 | 2935230 | 237350 | 6.57 | 13.01 | 68.7 | 21.75 | 20.86 | 1723 | 5.4757 | 80.49 |
| PORcat01 | POR01T | POR01T | cat01 | Portugal | Portela 3 | islamic (10-13 th c AD) | FX17 | FX-AP | 6190280 | 4990 | 3893 | 848 | 14.53 | 21.98 | 0.1 | 36.4 | 27.5 | 44 | 0.15 | 0 |
| PORcat01 | POR01B | POR01B | cat01 | Portugal | Portela 3 | islamic (10-13 th c AD) | FX17 | FX-AP | 7200733 | 4611 | 3738 | 429 | 8.51 | 18.93 | 0.1 | 33 | 28.2 | 3 | 0.01 | 0 |
| PORcat02 | POR02B | POR02B | cat02 | Portugal | Portela 3 | islamic (10-13 th c AD) | FX17 | FX-AP | 6247825 | 65367 | 59091 | 3245 | 4.73 | 9.60 | 1.0 | 27.2 | 25.5 | 115 | 0.37 | 0.79 |
| PORcat02 | POR02B_SS | POR02B_SS | cat02 | Portugal | Portela 3 | islamic (10-13 th c AD) | FX17 | FX-BM | 9440811 | 94140 | 83856 | 1392 | 1.46 | 10.92 | 0.9 | 19.07 | 0.9 | 211 | 0.55 | 2.61 |
| PORcat03 | POR03B | POR03B | cat03 | Portugal | Portela 3 | islamic (10-13 th c AD) | FX18 | FX-AR | 10877311 | 30686 | 27205 | 10008 | 24.59 | 11.34 | 0.3 | 39.29 | 39.5 | 170 | 0.67 | 4.12 |
| PORcat03 | POR03T | POR03T | cat03 | Portugal | Portela 3 | islamic (10-13 th c AD) | FX18 | FX-AR | 13255706 | 6095 | 5410 | 1111 | 15.42 | 11.24 | 0.0 | 30.92 | 31.6 | 19 | 0.07 | 0 |
| PORcat04 | POR04B | POR04B | cat04 | Portugal | Portela 3 | islamic (10-13 th c AD) | FX18 | FX-AR | 8720061 | 32619 | 28670 | 8974 | 21.58 | 12.11 | 0.3 | 32.17 | 31.7 | 74 | 0.3 | 0.39 |
| PRAcat01 | PRA01 | PRA01 | cat01 | Italy | Grotta della Madonna (Praia, Calabria) | Mesolithic | FX8 | FX-AH | 14606807 | 109901 | 84277 | 204883 | 65.09 | 23.32 | 0.6 | 37.89 | 37.8 | 64 | 0.27 | 0.25 |
| PRAcat02 | PRA02_SS | PRA02_SS | cat02 | Italy | Grotta della Madonna (Praia, Calabria) | Upper Palaeolitic | FX8 | FX-BE | 8233740 | 743718 | 686972 | 14609 | 1.93 | 7.63 | 8.3 | 23.8 | 2.6 | 392 | 1.18 | 13.64 |
| PRAcat02 | PRA02 | PRA02 | cat02 | Italy | Grotta della Madonna (Praia, Calabria) | Upper Palaeolitic | FX8 | FX-AH | 14357793 | 421518 | 365475 | 130537 | 23.65 | 13.30 | 2.5 | 35.3 | 35.6 | 182 | 0.67 | 4.92 |
| PRAcat03 | PRA03 | PRA03 | cat03 | Italy | Grotta della Madonna (Praia, Calabria) | Upper Palaeolitic | FX8 | FX-AH | 15680885 | 80404 | 69553 | 23779 | 22.82 | 13.50 | 0.4 | 26 | 26.4 | 66 | 0.23 | 0.47 |
| PRAcat04 | PRA04 | PRA04 | cat04 | Italy | Grotta della Madonna (Praia, Calabria) | Upper Palaeolitic | FX8 | FX-AH | 15809865 | 77394 | 66115 | 28521 | 26.93 | 14.57 | 0.4 | 39.15 | 38.8 | 26 | 0.09 | 0 |
| PRAcat05 | PRA05 | PRA05 | cat05 | Italy | Grotta della Madonna (Praia, Calabria) | Upper Palaeolitic | FX8 | FX-AH | 14909466 | 115106 | 91058 | 93016 | 44.69 | 20.89 | 0.6 | 37.5 | 38 | 35 | 0.11 | 0 |
| PRNcat01 | PRN01 | PRN01 | cat01 | Italy | Partanna C. da Capo d'acqua | Early Bronze Age | FX12 | FX-AL | 12989491 | 1161 | 656 | 654 | 36.03 | 43.50 | 0.0 | 3.47 | 3.5 | 0 | 0 | 0 |
| PRNcat02 | PRN02 | PRN02 | cat01 | Italy | Partanna C. da Capo d'acqua | Early Bronze Age | FX41 | FX-DO | 6539358 | 5048841 | 4598121 | 960158 | 15.98 | 8.93 | 61.3 | 46.02 | 41.57 | 4338 | 13.7827 | 98.02 |
| PRNcat02 | PRN02 | PRN02 | cat01 | Italy | Partanna C. da Capo d'acqua | Early Bronze Age | FX41 | FX-CH | 7883213 | 6130373 | 5529938 | 1640381 | 21.11 | 9.79 | 58.1 | 46.93 | 42.93 | 5304 | 16.0998 | 98.79 |
| PURcat01 | PUR01T | PUR01T | cat01 | Italy | La Purissima, Sardinia | roman imperial | FX18 | FX-AR | 9852245 | 18279 | 15972 | 4771 | 20.70 | 12.62 | 0.2 | 36.93 | 38.5 | 73 | 0.25 | 0.31 |
| PURcat01 | PUR01B | PUR01B | cat01 | Italy | La Purissima, Sardinia | roman imperial | FX18 | FX-AR | 6636042 | 1936 | 1688 | 391 | 16.80 | 12.81 | 0.0 | 36.87 | 41.1 | 8 | 0.04 | 0 |
| RIPcat01 | RIP01 | RIP01 | cat01 | Italy | Ripoli | Neolitico | FX34 | FXCA | 4472190 | 291009 | 259983 | 23865 | 7.58 | 10.66 | 5.8 | 41.95 | 42.48 | 329 | 0.9219 | 7.04 |
| RIPcat01 | RIP01 | RIP01 | cat01 | Italy | Ripoli | Neolitico | FX25 | FX-BD | 7316391 | 36116 | 32409 | 7975 | 18.09 | 10.26 | 0.4 | 39.8 | 35.4 | 28 | 0.1 | 0 |
| ROCcat01 | ROC01 | ROC01 | cat01 | Italy | Roca vecchia (Melendugno, Lecce) | Bronze Age (Middle) | FX62 | FXDO | 7059804 | 3738926 | 3337617 | 675816 | 15.31 | 10.73 | 43.2 | 33.6 | 30.67 | 6052 | 20.4329 | 98.7 |
| ROCcat02 | ROC02 | ROC02 | cat02 | Italy | Roca vecchia (Melendugno, Lecce) | Bronze Age (Late) | FX62 | FXDO | 7950155 | 2206823 | 1962164 | 190186 | 7.93 | 11.09 | 24.1 | 32.71 | 29.1 | 3924 | 13.3061 | 98.42 |
| ROCcat04 | ROC04 | ROC04 | cat03 | Italy | Roca vecchia (Melendugno, Lecce) | Medieval | FX62 | FX-DO | 5762677 | 3620794 | 3299560 | 162793 | 4.30 | 8.87 | 55.7 | 31.57 | 28.8 | 10090 | 35.256 | 99.78 |
| ROMcat01 | ROM01 | ROM01 | cat01 | Italy | Grotta Romanelli (Castro Marina, Apulia) | Upper Palaeolitic (Epigravettian) ca. 10ka | FX9 | FX-AI | 19131354 | 2380 | 1795 | 887 | 27.15 | 24.58 | 0.0 | 31.8 | 31.7 | 9 | 0.02 | 0 |
| RSScat01 | RSS01 | RSS01 | cat01 | Italy | Roccia San Sebastiano (Mondragone, Caserta, Camp | Upper Palaeolitic (Gravettian), ca. 20ka | FX21 | FX-AT | 7520699 | 6838 | 5622 | 2155 | 23.96 | 17.78 | 0.1 | 50.9 | 51.6 | 2 | 0.01 | 0 |
| SAcat09 | SA09 | SA09 | cat9 | Turkey | Sagalassos | Early Byzantine (7th cent. AD) | FX32 | FXBS | 7708864 | 28760 | 26196 | 491 | 1.68 | 8.92 | 0.3 | 16.3 | 17.77 | 60 | 0.1747 | 0 |
| SAcat14 | SA14 | SA14 | cat14 | Turkey | Sagalassos | Early Byzantine | FX32 | FXBS | 5314106 | 4814550 | 4423287 | 220371 | 4.38 | 8.13 | 79.9 | 21.96 | 22.47 | 17467 | 59.6916 | 100 |
| SARcat01 | SAR01 | SAR01 | cat01 | Italy | Saracena (Cosenza, Calabria) | Early Neolithic | FX34 | FXCA | 6896042 | 4694946 | 4016099 | 142652 | 2.95 | 14.46 | 57.1 | 32.84 | 33.38 | 8019 | 27.5581 | 99.41 |
| SBRcat01 | SBR01 | SBR01 | cat01 | Austria | Salzburg - Residenz | 15/16th centuries | FX32 | FXBS | 4830900 | 3231449 | 2953529 | 210794 | 6.12 | 8.60 | 58.6 | 17.08 | 17.51 | 6586 | 21.9483 | 99.71 |
| SCAScat01 | SCAS01 | SCAS01 | cat01 | Italy | Sassari castle, Sardinia | modern age | FX1 | FX-AA | 16748968 | 32722 | 28709 | 8399 | 20.43 | 12.26 | 0.2 | 19.01 | 18.5 | 33 | 0.15 | 0 |
| SCATcat01 | SCAT01 | SCAT01 | cat01 | Italy | Nardò “torre santa Caterina” | 17-18th c AD | FX3 | FX-AC | 10859114 | 3806843 | 3412955 | 1175674 | 23.60 | 10.35 | 28.4 | 23.28 | 23.2 | 8085 | 39.1192 | 99.99 |
| SCATcat03 | SCAT03 | SCAT03 | cat02 | Italy | Nardò “torre santa Caterina” | 17-18th c AD | FX6 | FX-BF | 10674962 | 8105264 | 7350806 | 823470 | 9.22 | 9.31 | 63.9 | 17.25 | 16 | 14364 | 45.9055 | 99.85 |
| SCATcat03 | SCAT03 | SCAT03 | cat02 | Italy | Nardò “torre santa Caterina” | 17-18th c AD | FX6 | FX-AF | 5819750 | 1275448 | 907573 | 6560539 | 83.72 | 28.84 | 7.3 | 20.35 | 20.5 | 1587 | 5.77 | 88.65 |
| SDUCcat01 | SDUC01 | SDUC01 | cat01 | Italy | Sassari, palazzo ducale, Sardinia | 19th AD | FX14 | FX-AM | 8173477 | 33801 | 22269 | 168829 | 83.32 | 34.12 | 0.3 | 21.62 | 21.6 | 19 | 0.11 | 0 |
| SDUCcat02 | SDUC02B | SDUC02B | cat02 | Italy | Sassari, palazzo ducale, Sardinia | 19th AD | FX21 | FX-AT | 8478297 | 2551365 | 2270475 | 914374 | 26.38 | 11.01 | 24.2 | 21.53 | 21.4 | 4807 | 16.147 | 99.65 |
| SDUCcat02 | SDUC02T | SDUC02T | cat02 | Italy | Sassari, palazzo ducale, Sardinia | 19th AD | FX21 | FX-AT | 9828887 | 499906 | 453181 | 104479 | 17.29 | 9.35 | 4.6 | 24.27 | 24 | 3611 | 16.14 | 99.85 |
| SFBIcat01 | SFBI01 | SFBI01 | cat01 | Italy | Sassari, via del Fiore Bianco, Sardinia | medieval or modern age | FX1 | FX-AA | 16052810 | 190916 | 137375 | 253103 | 57.00 | 28.04 | 0.8 | 24.73 | 22.7 | 206 | 1.06 | 10.03 |
| SQCcat03 | SQC03 | SQC03 | cat02 | Italy | Monastero Ss.Quattro Coronati- ex-portico (Rome) | end 16th- beg.17th cent AD | FX14 | FX-AM | 9438278 | 16082 | 12354 | 19551 | 54.87 | 23.18 | 0.1 | 20.24 | 19.4 | 38 | 0.17 | 0.05 |
| SSATcat05 | SSAT05 | SSAT05 | cat02? | Italy | Sassari, pozzo di via Satta, Sardinia | early 14th AD | FX1 | FX-AA | 14851652 | 8213 | 4536 | 14288 | 63.50 | 44.77 | 0.0 | 14.63 | 13.4 | 3 | 0.01 | 0 |
| SSPIcat01 | SSPI01 | SSPI01 | cat01 | Italy | Sassari, infermeria San Pietro, Sardinia | 18/19th c AD | FX1 | FX-AA | 17895633 | 72744 | 29414 | 332554 | 82.05 | 59.57 | 0.2 | 12.58 | 12.3 | 48 | 0.18 | 0.52 |
| STURcat01 | STUR01B | STUR01B | cat01 | Italy | Sassari, via Turritana, Sardinia | 17th AD | FX21 | FX-AT | 11069454 | 481384 | 428533 | 128376 | 21.05 | 10.98 | 3.8 | 21.41 | 21.3 | 1121 | 3.8 | 70 |
| STURcat01 | STUR01T | STUR01T | cat01 | Italy | Sassari, via Turritana, Sardinia | 17th AD | FX21 | FX-AT | 8759683 | 311947 | 276731 | 96564 | 23.64 | 11.29 | 3.1 | 22.91 | 23 | 1182 | 5.22 | 87.51 |
| TABcat01 | TAB01B | TAB01B | wildcat1 | Spain | Tabernas, Spain | 2,700-2,400 BC (Phase III) | FX33 | FXBZ | 5579709 | 35732 | 30415 | 1113 | 3.02 | 14.88 | 0.5 | 26.95 | 28.26 | 13 | 0.0438 | 0 |
| TABcat02 | TAB02B | TAB02B | wildcat2 | Spain | Tabernas, Spain | 2,700-2,400 BC (Phase III) | FX33 | FXBZ | 6626940 | 1102272 | 936259 | 51001 | 4.42 | 15.06 | 14.0 | 35.17 | 35.91 | 1317 | 4.1465 | 66.98 |
| TBScat01 | TBS01 | TBS01 | cat01 | Germany | Tauberbischofsheim-Dittigheim | 26th/25th century BCE, Chalcolithic (Corded Ware Culture | FX50 | FX-CV | 5701064 | 42239 | 37759 | 2143 | 4.83 | 10.61 | 0.7 | 32.47 | 31.15 | 71 | 0.2022 | 0.15 |
| TERTcat03 | TERT03 | TERT03 | cat01? | Italy | Tertenia, Sardinia | 4th-5th c AD | FX31 | FX-BO | 10096080 | 938624 | 852958 | 21137 | 2.20 | 9.13 | 8.4 | 40.16 | 36.6 | 576 | 1.71 | 27.2 |
| TERTcat03 | TERT03 | TERT03 | cat01? | Italy | Tertenia, Sardinia | 4th-5th c AD | FX21 | FX-AT | 6630417 | 254628 | 218481 | 119963 | 32.03 | 14.20 | 3.2 | 39.36 | 39.4 | 203 | 0.6 | 2.4 |
| TERTcat03 | TERT03 | TERT03 | cat01? | Italy | Tertenia, Sardinia | 4th-5th c AD | FX31 | FX-DM | 6826865 | 235696 | 201715 | 33427 | 12.42 | 14.42 | 2.9 | 31.35 | 27.4 | 106 | 0.4105 | 1.52 |
| TONcat03 | TON03 | TON03 | cat02 | Belgium | Tongeren, Industrie Oost | 40BC (4.0%) 10BC / AD (91.4%) 130AD | FX41 | FXCH | 8010754 | 6608230 | 6032789 | 114305 | 1.70 | 8.71 | 74.3 | 35.16 | 30.67 | 6389 | 20.9071 | 99.37 |
| TOVcat01 | TOV01 | TOV01 | cat01 | Portugal | Torre Velha 3 (Serpa) | Late Antiquity | FX6 | FX-AF | 11926402 | 92591 | 79043 | 195272 | 67.84 | 14.63 | 0.7 | 46.96 | 46.7 | 427 | 1.87 | 30.01 |
| TRLcat01 | TRL01 | TRL01 | cat01 | Turkey | Tralles | Roman? | FX50 | FX-CV | 12498671 | 1155989 | 1047622 | 75206 | 6.11 | 9.37 | 8.3 | 31.47 | 27.07 | 990 | 3.5492 | 65.42 |
| TRLcat02 | TRL02 | TRL02 | cat01 | Turkey | Tralles | Roman? | FX50 | FX-CV | 4918349 | 40349 | 36417 | 1292 | 3.10 | 9.74 | 0.7 | 41.09 | 38 | 76 | 0.2604 | 0.21 |
| TRMRcat01 | TRMR01B | TRMR01B | cat01 | Austria | Traismauer | 60AD (95.4%) 220AD | FX46 | FXCO | 6388079 | 3811454 | 3437947 | 2526116 | 39.86 | 9.80 | 38.6 | 23.31 | 18.34 | 12107 | 45.6678 | 99.96 |
| TRMRcat02 | TRMR02 | TRMR02 | cat01? | Austria | Traismauer | Roman | FX14 | FX-AM | 6934853 | 135047 | 113482 | 66163 | 32.88 | 15.97 | 1.6 | 23.48 | 23.4 | 91 | 0.35 | 1.15 |
| UZcat02 | UZ02 | UZ02 | cat02 | Italy | Grotta dell'Uzzo (Trapani, Sicily) | 5000-4800BC (Neolithic I) | FX7 | FX-AG | 11568224 | 734426 | 315904 | 2508754 | 77.35 | 56.99 | 2.2 | 30.49 | 30.3 | 195 | 0.54 | 3.08 |
| UZcat03 | UZ03 | UZ03 | cat03 | Italy | Grotta dell'Uzzo (Trapani, Sicily) | Neolithic II | FX31 | FX-BO | 11085230 | 2587 | 1853 | 298 | 10.33 | 28.37 | 0.0 | 28.1 | 22.4 | 3 | 0.01 | 0 |
| UZcat03 | UZ03 | UZ03 | cat03 | Italy | Grotta dell'Uzzo (Trapani, Sicily) | Neolithic II | FX7 | FX-AG | 16258558 | 1786 | 1052 | 665 | 27.13 | 41.10 | 0.0 | 18.42 | 17.7 | 1 | 0 | 0 |
| UZcat04 | UZ04 | UZ04 | cat04 | Italy | Grotta dell'Uzzo (Trapani, Sicily) | Neolithic II | FX7 | FX-AG | 13473805 | 88225 | 67422 | 78879 | 47.20 | 23.58 | 0.5 | 33.87 | 34.1 | 99 | 0.33 | 1.49 |
| UZcat05 | UZ05 | UZ05 | cat05 | Italy | Grotta dell'Uzzo (Trapani, Sicily) | Neolithic I | FX31 | FX-BO | 10418195 | 783979 | 706223 | 34341 | 4.20 | 9.92 | 6.8 | 45.93 | 46 | 435 | 1.2 | 14.1 |
| UZcat06 | UZ06 | UZ06 | cat06 | Italy | Grotta dell'Uzzo (Trapani, Sicily) | Mesolithic/Neolithic Transition | FX31 | FX-BO | 10874120 | 886279 | 804235 | 42277 | 4.55 | 9.26 | 7.4 | 44.1 | 42.6 | 731 | 2.1 | 35.46 |
| VIAcat01 | VIA01 | VIA01 | cat01 | Serbia | Viminacium/Amphitheatre | 4th century AD | FX2 | FX-AB | 17779433 | 115654 | 77832 | 374713 | 76.41 | 32.70 | 0.4 | 16.51 | 16.8 | 148 | 0.68 | 2.6 |
| VIAcat02 | VIA02 | VIA02 | cat02 | Serbia | Viminacium/Amphitheatre | 2nd-3rd century AD | FX2 | FX-AB | 23406525 | 206289 | 168515 | 114486 | 35.69 | 18.31 | 0.7 | 30.53 | 30.6 | 1115 | 3.79 | 66.05 |
| VIAcat03 | VIA03 | VIA03 | cat03 | Serbia | Viminacium/Amphitheatre | 4th century AD | FX2 | FX-AB | 16376404 | 92394 | 78359 | 30512 | 24.83 | 15.19 | 0.5 | 29.18 | 29.2 | 197 | 0.77 | 4.73 |
| VIAcat04 | VIA04 | VIA04 | cat04 | Serbia | Viminacium/Amphitheatre | 250AD (16.8%) 280AD / 330AD (78.6%) 420AD | FX2 | FX-AB | 12647551 | 6940080 | 5945571 | 4935780 | 41.56 | 14.33 | 33.8 | 24.01 | 24.1 | 9663 | 39.5688 | 99.81 |
| VIAcat06 | VIA06a | VIA06a | cat06 | Serbia | Viminacium/Amphitheatre | 20AD (95.4%) 210AD | FX2 | FX-AB | 13986149 | 4671882 | 4106412 | 1283445 | 21.55 | 12.10 | 26.9 | 31.91 | 31.8 | 8187 | 36.5289 | 99.99 |
| VIAcat06 | VIA06b | VIA06b | cat06 | Serbia | Viminacium/Amphitheatre | 20AD (95.4%) 210AD | FX2 | FX-AB | 16722340 | 3980499 | 3459518 | 1033805 | 20.62 | 13.09 | 19.5 | 30.63 | 30.8 | 5732 | 25.18 | 99.81 |
| VIAcat07 | VIA07 | VIA07 | cat07 | Serbia | Viminacium/Amphitheatre | 2nd-3rd century AD | FX2 | FX-AB | 17786761 | 758911 | 659494 | 206196 | 21.37 | 13.10 | 3.7 | 36.54 | 36.6 | 940 | 3.77 | 67.55 |
| VIAcat10 | VIA10 | VIA10 | cat10 | Serbia | Viminacium/Amphitheatre | 3rd century AD | FX6 | FX-AF | 9079097 | 76955 | 37155 | 1244350 | 94.18 | 51.72 | 0.4 | 29.43 | 26 | 50 | 0.21 | 0.39 |
| VINcat04 | VIN04 | VIN04 | cat02 | Serbia | Viminacium-Nad Klepačkom | 990AD (49.5%) 1050AD / 1080AD (45.9%) 1160AD | FX33 | Dps-AG | 7013927 | 1803122 | 1657392 | 271064 | 13.07 | 8.08 | 22.8 | 24.39 | 22.44 | 12204 | 38.4053 | 99.81 |
| VINcat04 | VIN04 | VIN04 | cat02 | Serbia | Viminacium-Nad Klepačkom | 990AD (49.5%) 1050AD / 1080AD (45.9%) 1160AD | FX33 | FZBZ | 4363113 | 1032399 | 941301 | 325140 | 23.95 | 8.82 | 20.1 | 24.47 | 25.19 | 6899 | 22.4377 | 99.1 |
| ZAPcat02 | ZAP02 | ZAP02 | cat02 | Spain | Zapatari 33 | mid 13th-mid 16th c. | FX44 | FXCN | 8425337 | 103293 | 93785 | 26090 | 20.16 | 9.20 | 1.1 | 20.17 | 17.89 | 190 | 0.6264 | 3.45 |
| ZAPcat02 | ZAP02 | ZAP02 | cat02 | Spain | Zapatari 33 | mid 13th-mid 16th c. | FX28 | FX-BI | 8089272 | 27029 | 24084 | 528 | 1.92 | 10.90 | 0.3 | 19 | 18.1 | 29 | 0.08 | 0 |
| ZAPcat03 | ZAP03 | ZAP03 | cat03 | Spain | Zapatari 33 | mid 13th-mid 16th c. | FX28 | FX-BI | 11347819 | 740411 | 679564 | 25470 | 3.33 | 8.22 | 6.0 | 17.57 | 17.8 | 1686 | 5.8 | 90.14 |
| ZAPcat03 | ZAP03 | ZAP03 | cat03 | Spain | Zapatari 33 | mid 13th-mid 16th c. | FX44 | FXCN | 7540890 | 162010 | 130164 | 466556 | 74.23 | 19.66 | 1.6 | 16.22 | 13.36 | 272 | 0.9667 | 11.16 |
| ZORcat02 | ZOR02 | ZOR02 | cat01 | Spain | Zornoztegi | 8th-10th c. | FX54 | FX-DD | 5482686 | 75577 | 68040 | 1223 | 1.59 | 9.97 | 1.2 | 18.69 | 19.03 | 57 | 0.1731 | 0.11 |

**Supplementary table 3.** Results of the deep shotgun sequencing conducted on selected archaeological samples. To increase the coverage, genomic libraries of some samples were sequenced twice (indicated by an asterisk).

| Sample ID | Cat ID | Individual | Country | Site | element | Library code | QC_passed_reads (w/o dup) | Full genome mapped (w/o dup) | Full genome mapped -q 1 (w/o dup) | DeDup duplicates | % duplicates | % spurious | % endogenous | Nuclear 5' PMD | Nuclear 3' PMD | fold nu-cov | stDev | combined fold nu-cov | mtDNA mapped -q 1 (w/o dup) | mt 5' PMD | mt 3' PMD | fold mt-cov | %mt-3X |
|---|---|---|---|---|---|---|---|---|---|---|---|---|---|---|---|---|---|---|---|---|---|---|---|
| AC01 | ACcat01 | cat01 | Italy | Arene Candide (Finale Ligure, Savona, Liguria) | right MT V | DpS_AH-2 | 20,754,310 | 13,755,009 | 10,658,823 | 2,972,897 | 17.77 | 22.51 | 44.92 | 25.96% | 25.85% | 0.363 | 1.5603X | - | 14,239 | 30.74% | 29.04% | 54.6272X | 99.97% |
| AC01* | ACcat01 | cat01 | Italy | Arene Candide (Finale Ligure, Savona, Liguria) | right MT V | DpS_AH | 11,263,259 | 7,772,897 | 6,073,530 | 1,190,396 | 13.28 | 21.86 | 48.77 | 25.94% | 26.01% | 0.201 | 0.933X | - | 13,897 | 28.86% | 30.74% | 30.5905X | 97.18% |
| AF03 | AFcat03 | cat02 | Italy | Alba Fucens -Cisterna (Fucino, Abruzzi) | left pelvis | DpS_AC | 44,470,430 | 13,848,063 | 12,412,779 | 3,048,249 | 18.04 | 10.36 | 26.12 | 29.20% | 29.11% | 0.351 | 0.9267X | - | 29,291 | 29.24% | 29.52% | 107.2794X | 99.43% |
| ASK01 | ASKcat01 | wildcat1 | Turkey | Aşıklı Höyük | distal humurus right (adult) | FX-DO | 171,141,343 | 3,356,775 | 482,966 | 24,620,556 | 88 | 85.61 | 0.25 | 35.18% | 34.75% | 0.071 | 0.9177X | - | 2,525 | 58.38% | 48.24% | 7.3455X | 91.18% |
| BA02 | BAcat02 | wildcat2 | Turkey | Bademağacı | mandible right with P3, P4, M1 | DpS_AE | 31,380,104 | 14,167,505 | 12,462,350 | 6,353,947 | 30.96 | 12.04 | 33.03 | 45.55% | 45.65% | 0.275 | 0.6363X | - | 10,147 | 52.13% | 52.56% | 27.3956X | 99.58% |
| BA03 | BAcat03 | wildcat3 | Turkey | Bademağacı | mandible left | DpS_AE | 35,315,498 | 9,158,014 | 7,634,118 | 4,296,881 | 31.94 | 16.64 | 19.27 | 44.64% | 44.60% | 0.175 | 0.5797X | - | 21,102 | 53.97% | 53.83% | 58.5338X | 99.94% |
| BA03* | BAcat03 | wildcat3 | Turkey | Bademağacı | mandible left | DpS_AE-2 | 24,863,269 | 7,243,224 | 6,103,326 | 2,535,349 | 25.93 | 15.74 | 22.28 | 46.51% | 46.71% | 0.138 | 0.4881X | 0.313 | 4,087 | 55.19% | 55.65% | 47.8708X | 99.36% |
| BHHL01 | BHHLcat01 | cat01 | Austria | Bernhardsthal | tibia L compl (prox NF; dist NF) | DpS_AH-2 | 21,692,488 | 13,158,786 | 11,770,566 | 2,976,471 | 18.45 | 10.55 | 47.71 | 19.05% | 19.05% | 0.343 | 0.8788X | 0.498 | 21,764 | 17.96% | 19.04% | 82.1572X | 100.00% |
| BHHL01* | BHHLcat01 | cat01 | Austria | Bernhardsthal | tibia L compl (prox NF; dist NF) | DpS_AH | 9,579,237 | 6,086,902 | 5,466,666 | 937,597 | 13.35 | 10.19 | 51.98 | 19.02% | 19.07% | 0.155 | 0.4907X |  | 14,599 | 17.80% | 18.25% | 37.1146X | 99.41% |
| BHHL03 | BHHLcat03 | cat02? | Austria | Bernhardsthal | radius R dist (dist F) | DpS_AH | 48,675,737 | 7,339,734 | 6,300,082 | 2,383,381 | 24.51 | 14.16 | 12.34 | 17.47% | 17.61% | 0.215 | 0.7957X | - | 3,968 | 21.86% | 21.02% | 30.3349X | 98.18% |
| BP01 | BPcat01 | cat01 | Portugal | Bank of Portugal | femur R compl (prox F; dist F) | DpSAN | 41,358,255 | 7,120,433 | 6,308,642 | 1,911,972 | 21.17 | 11.4 | 14.58 | 41.18% | 41.37% | 0.176 | 0.6068X | - | 27,245 | 34.73% | 34.84% | 116.5058X | 100.00% |
| BP02 | BPcat02 | cat02 | Portugal | Bank of Portugal | femur R compl (prox F; dist F) | DpSAN | 43,069,855 | 7,353,376 | 6,534,817 | 1,281,212 | 14.84 | 11.13 | 14.73 | 39.96% | 40.02% | 0.172 | 0.5331X | - | 11,296 | 37.32% | 37.78% | 41.3055X | 99.84% |
| BP03 | BPcat03 | cat03 | Portugal | Bank of Portugal | humerus R prox+shaft (prox F) | DpSAN | 49,864,910 | 6,464,010 | 5,624,494 | 1,425,111 | 18.06 | 12.99 | 10.97 | 27.50% | 27.14% | 0.168 | 0.544X | - | 7,102 | 29.32% | 28.55% | 28.0161X | 99.99% |
| BREM05 | BREMcat05 | cat04 | Germany | Bremen; 201-Altstadt, Marktplatz | femur L compl (prox F; dist F) | DpS_AH | 39,787,046 | 10,519,370 | 9,402,848 | 1,966,204 | 15.75 | 10.61 | 22.52 | 17.83% | 17.78% | 0.281 | 0.7198X | - | 18,789 | 23.76% | 25.74% | 36.3804X | 99.06% |
| BREM07 | BREMcat07 | cat06 | Germany | Bremen; 253-Altstadt, Am Wall | femur R compl (prox F; dist F) | DpS_AI | 56,655,502 | 9,292,431 | 8,424,310 | 1,512,581 | 14 | 9.34 | 14.48 | 14.43% | 14.40% | 0.248 | 0.655X | - | 38,641 | 16.43% | 16.50% | 76.3155X | 99.91% |
| BRPA24 | BRPAcat24 | cat04B | Belgium | Parking 58, Brussels | mandible compl R, with C, P3 | DpS_AC | 64,049,390 | 6,761,996 | 6,241,225 | 1,121,139 | 14.22 | 7.7 | 9.58 | 10.11% | 0.51% | 0.181 | 0.6001X |  | 34,047 | 7.97% | 1.06% | 147.3184X | 99.95% |
| BRPA24* | BRPAcat24 | cat04B | Belgium | Parking 58, Brussels | mandible compl R, with C, P3 | DpS_AC-2 | 35,588,360 | 4,168,982 | 3,852,511 | 526,865 | 11.22 | 7.59 | 10.67 | 10.08% | 0.58% | 0.11 | 0.4159X | 0.291 | 62,212 | 8.30% | 1.41% | 88.3991X | 99.88% |
| BRPA27 | BRPAcat27 | cat07B | Belgium | Parking 58, Brussels | mandible compl L, with P3, M1 | DpS_AL-2 | 28,092,226 | 6,641,965 | 5,970,323 | 1,312,875 | 16.5 | 10.11 | 20.3 | 16.29% | 16.24% | 0.158 | 0.5007X | 0.311 | 19,302 | 16.40% | 14.24% | 74.7459X | 100.00% |
| BRPA27* | BRPAcat27 | cat07B | Belgium | Parking 58, Brussels | mandible compl L, with P3, M1 | DpS_AL | 25,488,678 | 6,521,320 | 5,867,421 | 1,125,541 | 14.72 | 10.03 | 22.05 | 16.16% | 16.14% | 0.153 | 0.4886X |  | 17,972 | 15.40% | 14.51% | 71.7525X | 99.93% |
| CAST02 | CASTcat02 | cat01 | Italy | Castiglione in Sabina | radius | DpSAH | 12,495,045 | 4,790,531 | 3,426,191 | 18,021,761 | 79 | 28.48 | 11.23 | 23.17% | 23.05% | 0.12 | 0.6564X | - | 5,626 | 24.40% | 25.84% | 21.6364X | 99.46% |
| CEC03 | CECcat03 | cat03 | Italy | Rome, S. Cecilia in Trastevere | pelvis | DpS_AL | 47,358,706 | 6,727,476 | 5,872,594 | 1,511,083 | 18.34 | 12.71 | 12.02 | 31.30% | 31.27% | 0.162 | 0.5704X |  | 7,053 | 32.35% | 33.78% | 53.2088X | 53.2088X |
| CEC03* | CECcat03 | cat03 | Italy | Rome, S. Cecilia in Trastevere | pelvis | DpS_AL-2 | 42,829,045 | 5,802,304 | 5,077,756 | 1,307,596 | 18.39 | 12.49 | 11.5 | 31.04% | 31.47% | 0.143 | 0.5207X | 0.305 | 12,378 | 32.67% | 31.99% | 48.9385X | 99.98% |
| CET01 | CETcat01 | cat01 | Spain | Cova de Els Trocs, Huesca | ulna R compl (prox F; dist F) | DpS_AD | 27,173,756 | 21,372,811 | 18,982,895 | 4,717,957 | 18.08 | 11.18 | 59.52 | 24.36% | 24.50% | 0.619 | 1.5725X | - | 15,169 | 27.55% | 27.74% | 60.7292X | 99.77% |
| COC01 | COCcat01 | cat01 | Italy | Grotta del Cocci (Narni, Umbria) | right femur | DpS_AE | 44,729,592 | 9,406,998 | 7,647,311 | 4,839,402 | 33.97 | 18.71 | 15.43 | 32.71% | 32.56% | 0.196 | 0.6189X |  | 10,278 | 40.92% | 38.76% | 29.0957X | 99.10% |
| COC01* | COCcat01 | cat01 | Italy | Grotta del Cocci (Narni, Umbria) | right femur | DpS_AE-2 | 32,217,119 | 7,463,502 | 6,146,146 | 3,010,207 | 28.74 | 17.65 | 17.45 | 35.04% | 35.04% | 0.155 | 0.5151X | 0.351 | 9,838 | 43.13% | 40.82% | 23.2333X | 96.23% |
| DH06 | DHcat06 | wildcat1 | Turkey | Demircihüyük | complete ulna left (subadult) | DpS_AL | 7,767,757 | 6,583,845 | 5,894,459 | 2,759,179 | 29.53 | 10.47 | 55.99 | 18.40% | 18.16% | 0.191 | 0.5702X | - | 37,317 | 18.51% | 17.14% | 59.2015X | 99.52% |
| DOS01 | DOScat01 | wild1 | Italy | Galgenbühel/Dos de la Forca-Salorno | MT IV right | DpS_AG-2 | 66,348,385 | 6,756,274 | 5,661,646 | 5,770,923 | 46.07 | 16.2 | 7.85 | 24.24% | 24.32% | 0.186 | 0.6195X | 0.313 | 5,525 | 38.58% | 39.77% | 21.0571X | 98.55% |
| DOS01* | DOScat01 | wild1 | Italy | Galgenbühel/Dos de la Forca-Salorno | MT IV right | DpS_AG | 35,144,913 | 4,676,783 | 4,023,677 | 2,366,474 | 33.6 | 13.96 | 10.73 | 24.48% | 24.67% | 0.127 | 0.4535X |  | 8,163 | 37.20% | 39.95% | 14.5845X | 79.22% |
| DOS03 | DOScat03 | wild3 | Italy | Galgenbühel/Dos de la Forca-Salorno | radio distal | DpS_AC | 33,553,427 | 19,528,320 | 17,947,650 | 4,440,478 | 18.53 | 8.09 | 47.24 | 26.71% | 26.78% | 0.677 | 1.5604X | - | 12,420 | 26.31% | 26.36% | 58.9183X | 99.27% |
| DUR01 | DURcat01 | wildcat1 | Bulgaria | Durankulak | complete mandible right (adult) | DpS_AE | 35,522,907 | 19,295,115 | 16,955,686 | 4,499,594 | 18.91 | 12.12 | 42.37 | 41.46% | 41.48% | 0.404 | 1.0168X | - | 29,639 | 42.94% | 41.86% | 94.0912X | 99.93% |
| DUR02 | DURcat02 | wildcat2 | Bulgaria | Durankulak | fragment mandible right (adult) | DpS_AE | 41,203,584 | 27,961,540 | 24,868,994 | 5,696,665 | 16.93 | 11.06 | 53.03 | 44.72% | 44.68% | 0.589 | 1.6981X | - | 130,730 | 41.29% | 41.12% | 444.7587X | 100.00% |
| DUR03 | DURcat03 | wildcat3 | Bulgaria | Durankulak | complete mandible left (juvenile) | DpS_AD | 36,927,977 | 27,248,481 | 23,569,987 | 11,297,941 | 29.31 | 13.5 | 48.87 | 43.22% | 43.26% | 0.572 | 2.2952X | - | 216,202 | 44.04% | 44.34% | 730.5261X | 100.00% |
| DUR07 | DURcat07 | wildcat5 | Bulgaria | Durankulak | complete MT III (adult) | DpS_AD | 28,456,217 | 27,248,481 | 18,669,676 | 6,070,578 | 18.22 | 31.48 | 54.07 | 31.45% | 31.55% | 0.469 | 0.9375X | - | 9,270 | 40.39% | 36.16% | 29.4284X | 99.18% |
| ECLY01 | ECLYcat01 | cat01 | France | Ecly | left mandible | DpS_AD | 26,650,065 | 20,801,388 | 19,221,057 | 3,827,793 | 15.54 | 7.6 | 63.07 | 31.25% | 31.50% | 0.532 | 1.2236X | - | 88,638 | 32.21% | 32.04% | 317.8382X | 99.98% |
| FLUM02 | FLUMcat02 | cat02 | Italy | nuraghe Flumenelongu, Sardinia | right pelvis | DpS_AE-2 | 46,388,202 | 7,400,212 | 6,564,190 | 1,975,676 | 21.07 | 11.3 | 13.57 | 23.51% | 23.50% | 0.166 | 0.486X | 0.328 | 4,078 | 26.30% | 24.10% | 26.2594X | 98.99% |
| FLUM02* | FLUMcat02 | cat02 | Italy | nuraghe Flumenelongu, Sardinia | right pelvis | DpS_AE | 46,203,382 | 20,817,657 | 6,296,590 | 2,248,746 | 9.75 | 69.75 | 13 | 23.25% | 23.28% | 0.162 | 0.4823X |  | 7,736 | 24.26% | 24.29% | 26.2795X | 99.98% |
| GLCA01 | GLCAcat01 | cat01 | Ireland | Glencurran Cave, Co. Clare | mandible T fragm (with P3) | DpS_AI | 11,868,327 | 10,005,762 | 9,259,855 | 1,679,002 | 14.37 | 7.45 | 68.35 | 29.37% | 29.24% | 0.269 | 0.6867X | - | 9,144 | 31.75% | 29.86% | 94.2698X | 99.34% |
| GRA01 | GRAcat01 | cat01 | Italy | Graffignano (Viterbo, Latium) | left humerus | DpS_AA | 44,365,248 | 33,828,579 | 30,747,034 | 9,366,789 | 21.68 | 9.11 | 57.22 | 21.42% | 21.45% | 0.973 | 2.1013X | - | 81,731 | 23.53% | 23.89% | 326.661X | 100.00% |
| GSA01 | GSAcat01 | cat01 | Italy | Genoni Santu Antine, Sardinia | femur | DpS_AC | 35,905,458 | 7,834,171 | 6,606,202 | 5,499,224 | 41.24 | 15.67 | 15.96 | 24.80% | 25.00% | 0.188 | 0.7397X |  | 15,010 | 28.21% | 28.00% | 55.3569X | 99.06% |
| GSA01* | GSAcat01 | cat01 | Italy | Genoni Santu Antine, Sardinia | femur | DpS_AC-2 | 15,912,909 | 4,800,088 | 4,145,751 | 1,818,921 | 27.48 | 13.63 | 23.38 | 25.25% | 25.41% | 0.111 | 0.4564X | 0.299 | 17,712 | 26.71% | 28.32% | 32.7251X | 98.51% |
| HAIS01 | HAIScat01 | cat01 | Germany | Haithabu, settlement area | mandible R compl (with C, P3, P4, M1) | DpS_AI | 41,967,562 | 14,886,722 | 13,061,360 | 3,767,403 | 20.2 | 12.26 | 28.56 | 25.42% | 25.46% | 0.336 | 0.9193X | - | 12,094 | 27.10% | 27.74% | 96.4352X | 99.92% |
| HAIS04 | HAIScat04 | cat04 | Germany | Haithabu, settlement area | mandible R compl (with P3, P4, M1) | DpS_AL | 50,258,791 | 9,039,912 | 7,810,836 | 2,422,931 | 21.14 | 13.6 | 14.83 | 13.44% | 13.46% | 0.215 | 0.8484X | - | 24,273 | 14.85% | 14.53% | 172.7177X | 100.00% |
| HAIS10 | HAIScat10 | cat10 | Germany | Haithabu, settlement area | mandible R compl (with P3, P4) | DpS_AL-2 | 82,209,207 | 8,108,700 | 6,613,341 | 5,917,262 | 42.19 | 18.44 | 7.5 | 19.15% | 18.84% | 0.164 | 0.7708X | 0.291 | 48,294 | 14.84% | 14.63% | 176.4954X | 99.99% |
| HAIS10* | HAIScat10 | cat10 | Germany | Haithabu, settlement area | mandible R compl (with P3, P4) | DpS_AL | 54,202,734 | 6,306,409 | 5,262,950 | 3,093,873 | 32.91 | 16.55 | 9.19 | 19.22% | 19.00% | 0.127 | 0.6098X |  | 17,372 | 15.19% | 14.91% | 138.5495X | 99.94% |
| IRT01 | IRTcat01 | cat01 | Turkey | Iznik - Roman theatre | humerus L compl (prox NF; dist F; SA) | DpS_AL | 30,308,219 | 11,823,170 | 10,695,865 | 1,764,295 | 12.98 | 9.53 | 33.35 | 24.53% | 24.61% | 0.318 | 0.7816X | - | 3,052 | 30.20% | 29.66% | 36.1339X | 98.47% |
| KASP01 | KASPcat01 | wildcat1 | Greece | Kassope | mandible left (adult) | DpS_AD | 28,323,239 | 20,033,521 | 17,893,106 | 3,556,318 | 15.08 | 10.68 | 56.13 | 30.33% | 30.31% | 0.496 | 1.0022X | - | 16,226 | 37.91% | 38.18% | 52.9183X | 99.41% |
| KASP02 | KASPcat02 | wildcat2 | Greece | Kassope | distal femur left (adult) | DpS_AE | 53,986,169 | 9,178,225 | 7,921,750 | 2,758,075 | 23.11 | 13.69 | 13.96 | 34.41% | 34.34% | 0.2 | 0.6087X |  | 14,563 | 38.21% | 39.84% | 44.954X | 99.64% |
| KASP02* | KASPcat02 | wildcat2 | Greece | Kassope | distal femur left (adult) | DpS_AE-2 | 39,281,762 | 7,464,604 | 6,487,823 | 1,531,328 | 17.02 | 13.09 | 15.9 | 34.53% | 34.55% | 0.161 | 0.5144X | 0.361 | 9,444 | 40.66% | 39.34% | 37.1676X | 98.66% |
| KOP04 | KOPcat04 | wildcat4 | Bulgaria | Koprivec | pelvis (adult) | DpS_AE | 33,064,649 | 12,249,032 | 9,219,256 | 5,746,540 | 31.93 | 24.73 | 23.75 | 29.98% | 30.08% | 0.261 | 0.8926X | - | 24,629 | 36.69% | 35.91% | 78.9112X | 99.95% |
| KOP04* | KOPcat04 | wildcat4 | Bulgaria | Koprivec | pelvis (adult) | DpS_AE-2 | 15,280,726 | 6,362,711 | 4,976,441 | 1,548,337 | 19.57 | 21.79 | 29.57 | 30.84% | 30.90% | 0.134 | 0.5089X | 0.395 | 8,218 | 37.44% | 37.92% | 42.0867X | 99.16% |
| LAB01 | LABcat01 | cat01 | Spain | Labastida | metatarsus | DpS_AH | 9,779,098 | 8,491,807 | 7,652,965 | 1,621,355 | 16.03 | 9.88 | 67.13 | 15.05% | 15.21% | 0.26 | 0.7141X | - | 7,878 | 18.61% | 18.72% | 60.3819X | 99.54% |
| LHM02 | LHMcat02 | cat02 | Germany | Lauchheim-Mittelhofen | humerus R ~compl (prox NF; dist NF) | DpS_AL | 49,913,589 | 12,206,676 | 11,136,976 | 1,751,438 | 12.55 | 8.76 | 21.56 | 30.25% | 30.54% | 0.315 | 0.9242X | - | 16,073 | 31.11% | 30.14% | 215.3602X | 99.89% |
| LHM03 | LHMcat03 | cat03 | Germany | Lauchheim-Mittelhofen | ulna L prox (prox F) | DpS_AL | 52,868,619 | 7,417,137 | 6,616,472 | 1,356,547 | 15.46 | 10.79 | 12.2 | 26.95% | 27.13% | 0.202 | 0.6839X | - | 13,199 | 27.32% | 28.89% | 71.6168X | 99.36 |
| MENT01 | MENTcat01 | wild? cat01 | Turkey | Menteşe | ulna R shaft | DpSAN | 143,029,599 | 7,770,213 | 6,028,199 | 3,235,742 | 29.4 | 22.42 | 4.12 | 42.40% | 42.30% | 0.175 | 0.801X | - | 25,910 | 49.59% | 50.16% | 82.3212X | 99.95% |
| MRY07 | MRYcat07 | cat7 | Turkey | Yenikapı, Marmaray exc. | complete femur right (adult) | DpS_AE | 44,867,324 | 25,127,717 | 22,619,516 | 4,561,400 | 15.36 | 9.98 | 45.76 | 32.04% | 32.27% | 0.65 | 1.3842X | - | 27,802 | 36.73% | 36.53% | 95.4478X | 99.99% |
| MRY08 | MRYcat08 | cat8 | Turkey | Yenikapı, Marmaray exc. | complete femur right (adult) | DpS_AD | 30,381,815 | 18,375,164 | 16,322,495 | 4,125,850 | 18.34 | 11.17 | 47.3 | 40.96% | 41.25% | 0.424 | 0.9494X | - | 24,323 | 43.80% | 45.77% | 78.4028X | 99.95% |
| MRY09 | MRYcat09 | cat9 | Turkey | Yenikapı, Marmaray exc. | complete femur right (adult) | DpS_AC | 101,283,645 | 10,480,240 | 8,816,084 | 11,514,635 | 52.35 | 15.88 | 7.82 | 19.66% | 19.75% | 0.306 | 0.8919X | - | 5,300 | 26.98% | 26.76% | 22.6896X | 93.53% |
| MRY10 | MRYcat10 | cat10 | Turkey | Yenikapı, Marmaray exc. | complete femur right (adult) | DpS_AD | 34,387,900 | 27,985,847 | 25,109,350 | 5,288,361 | 15.89 | 10.28 | 63.29 | 37.74% | 37.93% | 0.673 | 1.3085X | - | 37,847 | 47.98% | 48.65% | 124.5823X | 100.00% |
| MTR01B | MTRcat01 | cat01 | Austria | Mautern - vicus Ost | mandible L compl (with C, P3, P4, M1) | DpS_AA | 63,571,620 | 48,491,268 | 43,668,193 | 7,409,760 | 13.26 | 9.95 | 61.52 | 25.52% | 25.53% | 1.398 | 3.0075X | - | 102,975 | 25.51% | 25.11% | 442.3856X | 100.00% |
| MUS01 | MUScat01 | cat01 | Italy | Musarna (Viterbo, Latium) | atlas | DpS_AG-2 | 29,757,070 | 8,016,376 | 6,463,297 | 9,349,199 | 53.84 | 19.37 | 16.53 | 16.12% | 16.22% | 0.207 | 0.7745X | 0.403 | 10,735 | 22.93% | 23.56% | 42.9026X | 99.97% |
| MUS01* | MUScat01 | cat01 | Italy | Musarna (Viterbo, Latium) | atlas | DpS_AG | 25,568,544 | 7,679,524 | 6,235,497 | 8,068,698 | 51.24 | 18.8 | 18.54 | 16.16% | 16.34% | 0.196 | 0.7209X | - | 7,658 | 22.97% | 24.22% | 40.777X | 99.49% |
| PAD01 | PADcat01 | cat01 | Italy | Padova Via Cesare Battisti (Veneto) | right scapula | DpS_AH | 57,161,267 | 8,199,114 | 7,280,699 | 1,680,010 | 17.01 | 11.2 | 12.37 | 19.42% | 19.82% | 0.206 | 0.6536X | - | 29,295 | 16.64% | 18.86% | 18.173X | 89.42% |
| PAL09 | PALcat09 | cat09 | Portugal | Palmela Castle | humerus dist; NF prox NF dist | DpS_AL | 32,601,670 | 7,269,050 | 6,512,794 | 1,023,037 | 12.34 | 10.4 | 19.37 | 21.25% | 21.24% | 0.183 | 0.5497X | - | 10,299 | 25.59% | 25.63% | 47.9362X | 98.82% |
| PAL09* | PALcat09 | cat09 | Portugal | Palmela Castle | humerus dist; NF prox NF dist | DpS_AL-2 | 29,007,328 | 6,066,967 | 5,443,056 | 866,957 | 12.5 | 10.28 | 18.22 | 21.39% | 21.34% | 0.156 | 0.4915X | 0.339 | 10,495 | 25.86% | 25.96% | 41.2898X | 99.99% |
| PAL10 | PALcat10 | cat10 | Portugal | Palmela Castle | tibia | DpS_AL | 47,052,348 | 7,270,516 | 6,496,248 | 1,092,755 | 13.07 | 10.65 | 13.49 | 26.99% | 27.08% | 0.197 | 0.6339X | - | 4,945 | 29.58% | 27.69% | 63.8083X | 99.48% |
| PAL10* | PALcat10 | cat10 | Portugal | Palmela Castle | tibia | DpS_AL-2 | 45,982,026 | 6,246,561 | 5,580,402 | 1,049,132 | 14.38 | 10.66 | 11.87 | 27.02% | 27.09% | 0.174 | 0.5824X | 0.371 | 13,291 | 29.90% | 29.83% | 55.2295X | 99.95% |
| PETR06 | PETRcat06 | cat03 | Austria | Petronell-Carnuntum | femur L prox (prox F) | DpS_AA | 55,822,199 | 19,849,944 | 17,868,392 | 2,946,432 | 12.93 | 9.98 | 30.4 | 16.58% | 16.64% | 0.532 | 1.0807X | - | 9,359 | 18.55% | 19.37% | 34.9812X | 99.59% |
| PETR08B | PETRcat08 | cat05 | Austria | Petronell-Carnuntum | mandible L compl (with C, P3, P4, M1) | DpS_AA | 55,695,495 | 31,709,856 | 28,737,732 | 7,888,038 | 19.92 | 9.37 | 45.2 | 22.97% | 23.00% | 0.929 | 1.9133X | - | 51,931 | 22.94% | 23.26% | 199.6175X | 100.00% |
| PNE02 | PNEcat02 | cat02 | Italy | Palatino North East slope (Rome) | right humerus, distal | DpS_AL | 50,786,328 | 7,185,412 | 6,501,613 | 1,242,599 | 14.74 | 9.52 | 12.5 | 34.16% | 34.15% | 0.177 | 0.5491X | - | 12,515 | 34.69% | 34.91% | 84.2307X | 99.90% |
| PNE02* | PNEcat02 | cat02 | Italy | Palatino North East slope (Rome) | right humerus, distal | DpS_AL-2 | 49,438,537 | 6,442,746 | 5,823,455 | 1,281,963 | 16.6 | 9.61 | 11.48 | 34.19% | 34.15% | 0.161 | 0.5165X | 0.338 | 20,578 | 34.63% | 32.97% | 77.0441X | 99.95% |
| PNE03 | PNEcat03 | cat03 | Italy | Palatino North East slope (Rome) | right radius, proximal | DpS_AL-2 | 22,209,289 | 17,760,061 | 15,252,735 | 4,115,133 | 18.81 | 14.12 | 57.94 | 22.50% | 22.52% | 0.445 | 1.1625X | 0.642 | 8,787 | 27.81% | 28.50% | 31.7881X | 99.71% |
| PNE03* | PNEcat03 | cat03 | Italy | Palatino North East slope (Rome) | right radius, proximal | DpS_AL | 9,802,295 | 8,020,193 | 6,940,601 | 1,184,984 | 12.87 | 13.46 | 63.17 | 22.54% | 22.53% | 0.197 | 0.6117X | - | 15,678 | 28.14% | 28.95% | 14.241X | 73.21% |
| PRN02 | PRNcat02 | cat01 | Italy | Partanna C. da Capo d'acqua | Petrous | DpSAN | 16,795,568 | 12,333,507 | 11,044,902 | 4,574,339 | 27.05 | 10.45 | 51.68 | 47.43% | 47.55% | 0.299 | 0.6839X | - | 10,016 | 50.05% | 50.80% | 33.7306X | 98.99% |
| ROC01 | ROCcat01 | cat01 | Italy | Roca vecchia (Melendugno, Lecce) | humerus L prox+shaft (prox F) | DpSAN | 18,640,784 | 8,727,217 | 7,572,098 | 2,940,437 | 25.2 | 13.24 | 35.09 | 34.91% | 34.85% | 0.21 | 0.6351X | - | 13,784 | 37.59% | 38.51% | 48.9812X | 99.49% |
| ROC02 | ROCcat02 | cat02 | Italy | Roca vecchia (Melendugno, Lecce) | tibia R dist+shaft (dist F) | DpSAN | 36,271,328 | 8,078,071 | 6,968,933 | 2,242,729 | 21.73 | 13.73 | 18.09 | 34.09% | 34.03% | 0.202 | 0.6151X | - | 13,133 | 45.22% | 44.69% | 46.8848X | 99.18% |
| ROC04 | ROCcat04 | cat03 | Italy | Roca vecchia (Melendugno, Lecce) | skull with mandible (to sample) (skull with teeth, mandible with teeth) | DpSAN | 15,318,202 | 8,762,070 | 7,881,320 | 1,333,798 | 13.21 | 10.05 | 47.33 | 32.48% | 32.62% | 0.226 | 0.6954X | - | 24,512 | 30.90% | 31.07% | 100.2672X | 100.00% |
| SA14 | SAcat14 | cat14 | Turkey | Sagalassos | complete humerus left, subadult | DpS_AD | 24,724,209 | 21,680,052 | 19,589,765 | 4,776,254 | 18.05 | 9.64 | 66.4 | 22.29% | 22.34% | 0.571 | 1.2425X | - | 73,903 | 22.91% | 23.18% | 260.5526X | 100.00% |
| SAR01 | SARcat01 | cat01 | Italy | Saracena (Cosenza, Calabria) | left MC IV | DpS_AD | 34,223,938 | 21,369,665 | 17,664,762 | 4,288,866 | 16.72 | 17.34 | 45.87 | 33.04% | 32.74% | 0.508 | 1.2803X | - | 34,438 | 38.74% | 38.66% | 117.6453X | 99.99% |
| SBR01 | SBRcat01 | cat01 | Austria | Salzburg - Residenz | humerus L compl (prox & dist F) | DpS_AD | 36,101,973 | 21,227,561 | 18,889,991 | 6,555,848 | 23.6 | 11.01 | 44.28 | 17.06% | 17.04% | 0.495 | 1.0529X | - | 40,118 | 17.58% | 18.32% | 129.8365X | 100.00% |
| SCAT01 | SCATcat01 | cat01 | Italy | Nardò “torre santa Caterina” | tibia, dist F; prox NF | DpS_AA | 63,189,285 | 23,223,134 | 20,380,691 | 10,861,201 | 31.87 | 12.24 | 27.52 | 22.75% | 22.37% | 0.662 | 1.9709X | - | 45,763 | 22.41% | 22.57% | 187.2319X | 99.99% |
| SDUC02B | SDUCcat02 | cat02 | Italy | Sassari, palazzo ducale, Sardinia | right mandible with P3, P4 & M1 | DpS_AG | 29,815,535 | 9,529,272 | 8,567,016 | 3,979,817 | 29.46 | 10.1 | 25.35 | 22.38% | 22.24% | 0.246 | 0.6567X | - | 20,939 | 21.97% | 22.71% | 63.76X | 99.91% |
| TAB02B | TABcat02 | wildcat2 | Spain | Tabernas, Spain | distal humerus right (adult) | DpS_AH-2 | 141,799,212 | 21,309,869 | 17,675,885 | 4,955,168 | 18.87 | 17.05 | 12.04 | 33.95% | 32.54% | 0.442 | 1.1111X | 0.61 | 24,457 | 34.22% | 32.82% | 74.3898X | 99.88% |
| TAB02B* | TABcat02 | wildcat2 | Spain | Tabernas, Spain | distal humerus right (adult) | DpS_AH | 56,283,673 | 8,067,655 | 6,638,768 | 1,507,039 | 15.74 | 17.71 | 11.49 | 35.26% | 35.02% | 0.168 | 0.574X | - | 42,135 | 36.57% | 35.48% | 29.4017X | 95.99% |
| TERT03 | TERTcat03 | cat01? | Italy | Tertenia, Sardinia | humerus L compl (prox F; dist F) | DpSAN | 68,095,421 | 5,371,577 | 4,825,241 | 723,010 | 11.86 | 10.17 | 7.01 | 41.61% | 41.35% | 0.129 | 0.4202X | - | 3,228 | 51.26% | 46.80% | 10.2972X | 96.72% |
| TON03 | TONcat03 | cat02 | Belgium | Tongeren, Industrie Oost | os petrosum | DpS_AI | 10,539,033 | 8,343,614 | 7,565,368 | 959,695 | 10.32 | 9.33 | 65.79 | 35.96% | 36.18% | 0.214 | 0.6446X | - | 59,306 | 38.84% | 38.48% | 28.0209X | 97.70% |
| TRMR01B | TRMRcat01 | cat01 | Austria | Traismauer | mandible L compl, with I1, P3 & M1 | DpS_AG | 18,505,333 | 9,841,654 | 8,711,532 | 10,286,384 | 51.1 | 11.48 | 30.26 | 23.27% | 23.50% | 0.293 | 0.7999X | - | 18,950 | 23.34% | 23.04% | 140.9287X | 100% |
| VIA04 | VIAcat04 | cat04 | Serbia | Viminacium/Amphitheatre | mandible | DpS_AA | 34,398,125 | 20,304,057 | 17,665,197 | 12,791,806 | 38.65 | 13 | 37.43 | 24.35% | 25.14% | 0.556 | 1.3844X | - | 28,034 | 23.48% | 24.12% | 117.5081X | 100.00% |
| VIA06a | VIAcat06 | cat06 | Serbia | Viminacium/Amphitheatre | humerus | DpS_AB | 54,097,215 | 20,842,236 | 18,625,487 | 3,234,099 | 13.43 | 10.64 | 32.49 | 32.44% | 32.16% | 0.578 | 1.4172X | - | 37,230 | 30.00% | 29.50% | 173.9741X | 100.00% |
| VIN04 | VINcat04 | cat02 | Serbia | Viminacium-Nad Klepačkom | humerus; NF prox | DpSAG | 54,966,751 | 6,672,323 | 5,635,542 | 10,913,013 | 62.06 | 15.54 | 8.55 | 24.34% | 24.33% | 0.16 | 0.6064X | - | 39,718 | 26.93% | 27.15% | 132.379X | 99.96% |

**Supplementary table 4.** Results of the deep shotgun sequencing conducted on modern samples.

| ID ISPRA | date | Country | Site | source | Library code | QC_passed_reads (w/o dup) | Full genome mapped (w/o dup) | fold nu-cov | stDev | SNPs Panel preparation | modern nuclear genome analysis | PCA Fig. 1C cluster | mt-DNA analyzed | mtDNA |
|---|---|---|---|---|---|---|---|---|---|---|---|---|---|---|
| 46_Fsl | modern | Italy | Sardinia | DNA extracts from ISPRA | FX-CT | 265,077,211 | 237,053,796 | 10.7328X | 7.1353X | included | included | *Felis libica lybica/ Felis catus* | included | Felis lybica lybica/catus clade IV-E |
| 52_Fsl | modern | Italy | Sardinia, Monte Arcosu | DNA extracts from ISPRA | FX-CT | 297,471,369 | 288,081,670 | 12.7835X | 22.8118X | included | included | *Felis libica lybica/ Felis catus* | included | Felis lybica lybica/catus clade IV-E |
| 53_Fsl | modern | Italy | Sardinia, Monte Arcosu | DNA extracts from ISPRA | FX-CT | 337,786,146 | 328,134,277 | 14.4809X | 16.1764X | included | included | *Felis libica lybica/ Felis catus* | included | Felis lybica lybica/catus clade IV-C |
| 55_Fsl | modern | Italy | Sardinia, Nuoro - around Bitti | DNA extracts from ISPRA | FX-CT | 255,482,342 | 243,639,495 | 8.8466X | 38.4242X | excluded | excluded | *Felis libica lybica/ Felis catus* | included | Felis lybica lybica/catus clade IV-C |
| 66_Fss | modern | Italy | Tuscany - Maremma - Bivio Gavorrano | DNA extracts from ISPRA | FX-CT | 178,447,879 | 173,566,316 | 5.0639X | 33.0114X | excluded | excluded | *Felis silvestris* | included | Felis silvestris clade I |
| 78_Fss | modern | Italy | Probabilly Sicily | DNA extracts from ISPRA | FX-CS | 317,463,029 | 276,365,926 | 12.3302X | 15.7897X | included | included | *Felis silvestris* | included | Felis silvestris clade I |
| 79_Fss | modern | Italy | Sicily, monti Nebrodi | DNA extracts from ISPRA | FX-CS | 319,282,726 | 278,548,890 | 13.0231X | 9.6725X | included | included | *Felis silvestris* | included | Felis silvestris clade I |
| 99_Fss | modern | Italy | Tuscany | DNA extracts from ISPRA | FX-CT | 291,211,152 | 252,552,310 | 10.1288X | 11.4089X | included | included | *Felis silvestris* | included | Felis silvestris clade I |
| 997_Fss | modern | Italy | Rugo Storto, Fanna, PN | DNA extracts from ISPRA | FX-CS | 342,871,492 | 315,623,942 | 17.4632X | 9.8853X | included | included | *Felis silvestris* | included | Felis silvestris clade I |
| 1294_Fss | modern | Italy | Umbria - Perugia - Cascia | DNA extracts from ISPRA | FX-CS | 322,490,102 | 309,415,806 | 16.3347X | 23.4815X | included | included | *Felis silvestris* | included | Felis silvestris clade I |
| 1302_Fss | modern | Italy | Umbria - Perugia - Vallo di Nera | DNA extracts from ISPRA | FX-CS | 332,971,531 | 306,835,593 | 15.4634X | 10.7704X | included | included | *Felis silvestris* | included | Felis lybica lybica/catus clade IV-C |
| 1611_Fss | modern | Italy | Parco delle Dolomiti Lucane | DNA extracts from ISPRA | FX-CS | 307,438,766 | 293,977,732 | 15.5276X | 11.2534X | included | included | *Felis silvestris* | included | Felis silvestris clade I |
| 778_Fss | modern | Marocco | na | DNA extracts from ISPRA | FX-CR | 280,321,210 | 262,668,432 | 14.1321X | 8.3864X | included | included | *Felis libica lybica/ Felis catus* | included | Felis lybica lybica/catus clade IV-C |
| 779_Fss | modern | Tunisia | na | DNA extracts from ISPRA | FX-CR | 302,564,434 | 285,574,181 | 13.2867X | 13.1496X | included | included | *Felis libica lybica/ Felis catus* | included | Felis lybica lybica/catus clade IV-A2 |
| BG05 | 20th century | Bulgaria | village Chelopechene, Sofia district, died in the royal zoo | skin + hair | DpS_AF | 37,324,550 | 23,086,927 | 0.854X | 8.1508X | excluded | included | *Felis silvestris* | excluded | NA |
| BG07 | 20th century | Bulgaria | village Belene, Svishtov district (с. Белене, Свищовско) | claw fragment | DpS_AF | 41,128,557 | 31,785,173 | 1.219X | 9.6961X | excluded | included | *Felis silvestris* | excluded | NA |
| BG16 | 20th century | Bulgaria | Pomorie, Forestry station (Поморие, горско стопанство) | claw fragment | DpS_AF | 35,342,601 | 24,004,079 | 0.736X | 7.917X | excluded | included | *Felis silvestris* | excluded | NA |

**Supplementary table 5.** Newly generated radiocarbon dates.

| ID | ID-Lab | Individual | Site | Country | ID_kikirpa | uncalibrated date BP | 94% probability calibrated date |
|---|---|---|---|---|---|---|---|
| BHHLcat01 | BHHL01 | cat01 | Bernhardsthal | Austria | RICH-34515 | 1165±26BP | 770AD (95.4%) 980AD |
| MTRcat01 | MTR01B | cat01 | Mautern - vicus Ost | Austria | RICH-34526 | 2007±25BP | 50BC (95.4%) 80AD |
| PETRcat08 | PETR08B | cat05 | Petronell-Carnuntum | Austria | RICH-35370 | 1919±24BP | 20AD (2.6%) 50AD / 60AD (92.8%) 210AD |
| PETRcat06 | PETR06 | cat03 | Petronell-Carnuntum | Austria | RICH-34523 | 1830±26BP | 120AD (87.6%) 260AD / 290AD ( 7.8%) 320AD |
| TRMRcat01 | TRMR01B | cat01 | Traismauer | Austria | RICH-34530 | 1897±27BP | 60AD (95.4%) 220AD |
| TONcat03 | TON03 | cat02 | Tongeren, Industrie Oost | Belgium | RICH-34531 | 1953±27BP | 40BC (4.0%) 10BC / 10BC (91.4%) 130AD |
| DURcat01 | DUR01 | cat01 | Durankulak | Bulgaria | RICH-34522 | 5535±32BP | 4450BC (95.4%) 4330BC |
| KOPcat04 | KOP04 | cat04 | Koprivec | Bulgaria | RICH-34528 | 6452±31BP | 5480BC (95.4%) 5360BC |
| ECLYcat01 | ECLY01 | cat01 | Ecly | France | RICH-34507 | 1915±25BP | 20AD (2.1%) 50AD / 60AD (93.3%) 210AD |
| BREMcat05 | BREM05 | cat04 | Bremen; 201-Altstadt, Marktplatz | Germany | RICH-34505 | 381±24BP | 1440AD (65.0%) 1530AD / 1570AD (30.4%) 1630AD |
| KASPcat01 | KASP01 | cat01 | Kassope | Greece | RICH-34520 | 2253±25BP | 400BC (33.6%) 340BC / 310BC (61.8%) 200BC |
| AFcat03 | AF03 | cat02 | Alba Fucens -Cisterna (Fucino, Abruzzi) | Italy | RICH-29170 | 1783±24BP | 210AD (95.4%) 340AD |
| DOScat01 | DOS01 | cat01 | Galgenbühel/Dos de la Forca-Salorno | Italy | RICH-34534 | 8899±35BP | 8240BC (95.4%) 7940BC |
| GSAcat01 | GSA01 | cat01 | Genoni Santu Antine, Sardinia | Italy | RICH-34524 | 2121±25BP | 340BC (3.9%) 320BC / 200BC (91.5%) 50BC |
| COCcat01 | COC01 | cat01 | Grotta del Cocci (Narni, Umbria) | Italy | RICH-34517 | 6324±31BP | 5370BC (95.4%) 5210BC |
| MUScat01 | MUS01 | cat01 | Musarna (Viterbo, Latium) | Italy | RICH-34516 | 1644±25BP | 360AD (95.4%) 540AD |
| FLUMcat02 | FLUM02 | cat02 | nuraghe Flumenelongu, Sardinia | Italy | RICH-34518 | 1365±25BP | 600AD (2.7%) 620AD / 630AD (86.2%) 690AD / 740AD (6.6%) |
| PADcat01 | PAD01 | cat01 | Padova Via Cesare Battisti (Veneto) | Italy | RICH-34527 | 515±23BP | 1400AD (95.4%) 1440AD |
| PNEcat02 | PNE02 | cat02 | Palatino North East slope (Rome) | Italy | RICH-35364 | 1764±24BP | 230AD (95.4%) 370AD |
| ROCcat01 | ROC01 | cat01 | Roca vecchia (Melendugno, Lecce) | Italy | RICH-35362 | 3145±29BP | 1500BC (84.2%) 1380BC / 1350BC (11.2%) 1310BC |
| ROCcat02 | ROC02 | cat02 | Roca vecchia (Melendugno, Lecce) | Italy | RICH-35363 | 3051±23BP | 1410BC (95.4%) 1220BC |
| BPcat01 | BP01 | cat01 | Bank of Portugal | Portugal | RICH-35365 | 1084±23BP | 890AD (32.4%) 930AD / 940AD (63.0%) 1020AD |
| BPcat02 | BP02 | cat02 | Bank of Portugal | Portugal | RICH-35366 | 1049±23BP | 900AD ( 4.6%) 920AD / 970AD (90.8%) 1040AD |
| BPcat03 | BP03 | cat03 | Bank of Portugal | Portugal | RICH-35367 | 1059±23BP | 890AD (11.0%) 920AD / 950AD (84.4%) 1030AD |
| VINcat04 | VIN04 | cat02 | Viminacium-Nad Klepačkom | Serbia | RICH-34510 | 992±25BP | 990AD (49.5%) 1050AD / 1080AD (45.9%) 1160AD |
| VIAcat04 | VIA04 | cat04 | Viminacium/Amphitheatre | Serbia | RICH-34521 | 1697±25BP | 250AD (16.8%) 280AD / 330AD (78.6%) 420AD |
| VIAcat06 | VIA06 | cat06 | Viminacium/Amphitheatre | Serbia | RICH-34512 | 1923±25BP | 20AD (95.4%) 210AD |
| CETcat01 | CET01 | cat01 | Cova de Els Trocs, Huesca | Spain | RICH-34519 | 6207±32BP | 5300BC ( 9.4%) 5250BC / 5230BC (86.0%) 5040BC |
| ASKcat01 | ASK01 | wildcat01 | Aşıklı Höyük | Turkey | RICH-25375 | 8745±37BP | 7950BC (95.4%) 7610BC |
| BAcat02 | BA02 | cat02 | Bademağacı | Turkey | RICH-34506 | 7496±32BP | 6430BC (61.9%) 6330BC / 6320BC (33.5%) 6240BC |
| BAcat03 | BA03 | cat03 | Bademağacı | Turkey | RICH-34508 | 7398±32BP | 6390BC (90.0%) 6210BC / 6140BC ( 5.4%) 6090BC |
| BAYcat01 | BAY01 | cat01 | Bayraklı | Turkey | RICH-34525 | 103.26±0.27pMC | [cal AD 1955.82:cal AD 1956.47]0.215 / [cal AD 2012.05:cal AD |
| DHcat05 | DH05 | cat01 | Demircihüyük | Turkey | RICH-35360 | 608±22BP | 1300AD (74.5%) 1370AD / 1375AD (20.9%) 1400AD |
| IRTcat01 | IRT01 | cat01 | Iznik - Roman theatre | Turkey | RICH-35359 | 1448±22BP | 575AD (95.4%) 650AD |
| MENTcat01 | MENT01 | cat01 | Menteşe | Turkey | RICH-35361 | 7163±30BP | 6075BC (95.4%) 5980BC |
| MRYcat09 | MRY09 | cat09 | Yenikapı, Marmaray exc. | Turkey | RICH-34529 | 1313±26BP | 650AD (48.9%) 710AD / 720AD (46.5%) 780AD |
| MRYcat10 | MRY10 | cat10 | Yenikapı, Marmaray exc. | Turkey | RICH-34509 | 1350±25BP | 640AD (75.4%) 690AD / 740AD (20.0%) 780AD |

**Supplementary table 6.** Sequence data downloaded from literature and used for comparative analyses in this study.

| Species-Subspecies | ID | breed/status | Country - Region | sample IDs in plots | SRA | NCBI_BIOSAMPLE | BioProject | nuclear genome analysis | mitochondrial genome analysis |
|---|---|---|---|---|---|---|---|---|---|
| *Felis catus* (Domestic) | felCat.CVB6166.Morpheus | Maine Coon | *NA* | CGAE6166_MaineCoon | SRR11392458 | SAMN14425477 | PRJNA614458 | INCLUDED | INCLUDED |
| *Felis catus* (Domestic) | felCat.Fcat20907.BlueJasmine | Selkirk Rex | Canada | Fcat20907_SelkirRex | SRR11392498 | SAMN14425584 | PRJNA614458 | INCLUDED | INCLUDED |
| *Felis catus* (Domestic) | felCat.Fcat20784.Imortantum | American Wirehair | USA | Fcat20784_AmericanWirehair | SRR11392553 | SAMN14425422 | PRJNA614458 | INCLUDED | INCLUDED |
| *Felis catus* (Domestic) | felCat.CVB18351.Charlie | Devon Rex | *NA* | CGAE18351_DevonRex | SRR11392588 | SAMN14425449 | PRJNA614458 | INCLUDED | INCLUDED |
| *Felis catus* (Domestic) | felCat.Fcat16274.Shiya | Ragdoll | *NA* | Fcat16274_Ragdoll | SRR11392597 | SAMN14425547 | PRJNA614458 | INCLUDED | INCLUDED |
| *Felis catus* (Domestic) | felCat.Fcat22335.Selena | Turkish Angora | *NA* | Fcat22335_TurkishAngora | SRR11392600 | SAMN14425597 | PRJNA614458 | INCLUDED | INCLUDED |
| *Felis catus* (Domestic) | felCat.Fcat22361.Anna | Persian | *NA* | Fcat22361_Persian | SRR11392604 | SAMN14425544 | PRJNA614458 | INCLUDED | INCLUDED |
| *Felis catus* (Domestic) | felCat.Fcat22054.Chester | Siberian | *NA* | Fcat22054_Siberian | SRR11392605 | SAMN14425590 | PRJNA614458 | INCLUDED | INCLUDED |
| *Felis catus* (Domestic) | felCat.CVB15266.20680 | Egyptian Mau | USA | Fcat20680_EgyptianMau | SRR11392607 | SAMN14425454 | PRJNA614458 | INCLUDED | INCLUDED |
| *Felis catus* (Domestic) | felCat.CVB6197.Putney | Domestic Longhair | USA | CGAE6197_DomesticLonghair | SRR11392635 | SAMN14425452 | PRJNA614458 | INCLUDED | INCLUDED |
| *Felis catus* (Domestic) | felCat.CVB2613.Osiris | Sphynx | *NA* | CGAE2613_Sphynx | SRR11392636 | SAMN14425593 | PRJNA614458 | INCLUDED | INCLUDED |
| *Felis catus* (Domestic) | felCat.Fcat20822.Stradivarious | Ocicat | Europe | Fcat20822_Ocicat | SRR11392640 | SAMN14425505 | PRJNA614458 | INCLUDED | INCLUDED |
| *Felis catus* (Domestic) | felCat.Hills1.Sizzy | Random bred | Western | Sizzle_DomesticShorthair | SRR5055407 | SAMN05980362 | PRJNA343392 | INCLUDED | INCLUDED |
| *Felis catus* (Domestic) | felCat.Fcat18666.Portugal | Random bred | Portugal | Fcat18666_RandomBred | SRR5040114/SRR5040108 | SAMN05980325 | PRJNA343389 | INCLUDED | INCLUDED |
| *Felis catus* (Domestic) | felCat.Fcat18801.Italy | Random bred | Italy | Fcat18801_RandomBred | SRR5040117/SRR5040124 | SAMN05980326 | PRJNA343389 | INCLUDED | INCLUDED |
| *Felis catus* (Domestic) | felCat.DS05501.Mateo | Random bred | UK | DS05501_RandomBred | SRR11392568 | SAMN14425581 | PRJNA614458 | INCLUDED | INCLUDED |
| *Felis catus* (Domestic) | felCat.Katze169.Moischily | Random bred | Germany | Katze169_RandomBred | SRR11392572 | SAMN14425570 | PRJNA614458 | INCLUDED | INCLUDED |
| *Felis catus* (Domestic) | felCat.Hills2.Question | Random bred | Western | Hills2_RandomBred | SRR5055383 | SAMN05980363 | PRJNA343392 | INCLUDED | INCLUDED |
| *Felis lybica lybica* | FSI47 | wild | Israel | FSI47_Fsl | ERR10494332 | SAMEA112127403 | PRJEB57412 | INCLUDED | INCLUDED |
| *Felis lybica lybica* | FSI48 | wild | Israel | FSI48_Fsl | ERR10494333-ERR104943[illegible] | SAMEA112127404 | PRJEB57412 | INCLUDED | INCLUDED |
| *Felis lybica lybica* | FSI51 | wild | Israel | FSI51_Fsl | ERR10494336-ERR104943[illegible] | SAMEA112127405 | PRJEB57412 | INCLUDED | INCLUDED |
| *Felis silvestris* | FSX360 | wild | Portugal | FSX360_Fss | ERR10494342, ERR104943[illegible] | SAMEA112127406 | PRJEB57412 | INCLUDED | INCLUDED |
| *Felis silvestris* | FSX392 | wild | Scotland | FSX392_Fss | ERR10494345 | SAMEA112127407 | PRJEB57412 | INCLUDED | INCLUDED |
| *Felis silvestris* | FSX405 | wild | Scotland | FSX405_Fss | ERR10494346-ERR104943[illegible] | SAMEA112127408 | PRJEB57412 | INCLUDED | INCLUDED |
| *Felis bieti* | FBI4 | wild | China | FBI4_Fsb | ERR10494327 | SAMEA112127400 | PRJEB57412 | INCLUDED | INCLUDED |
| *Felis bieti* | FBIP0004 | wild | China | FBIP0004_Fsb | SRR7621226 | SAMN09509265 | PRJNA478778 | INCLUDED | INCLUDED |
| *Felis bieti* | FBIP0008 | wild | China | FBIP0008_Fsb | SRR7621227 | SAMN09509266 | PRJNA478778 | INCLUDED | INCLUDED |
| *Felis bieti* | FBIP0003 | wild | China | FBIP0003_Fsb | SRR7621229 | SAMN0950926 | PRJNA478778 | INCLUDED | INCLUDED |
| *Felis bieti* | FBIP0026 | wild | China | FBIP0026_Fsb | SRR7621235 | SAMN09509267 | PRJNA478778 | INCLUDED | INCLUDED |
| *Felis lybica ornata* | FSIP0006 | wild | China | FSIP0006_Fso | SRR7621238 | SAMN09509272 | PRJNA478778 | INCLUDED | INCLUDED |
| *Felis lybica ornata* | FSI204 | wild | Kazakhstan | FSI204_Fso | ERR10494331 | SAMEA112127402 | PRJEB57412 | INCLUDED | INCLUDED |
| *Felis lybica ornata* | FLI3 | wild | Tajikistan | FLI3_Fso | SRR15116526 | SAMN20192906 | PRJNA746237 | INCLUDED | INCLUDED |
| *Felis lybica ornata* | FLI4 | wild | Kazakhstan | FLI4_Fso | SRR15116525 | SAMN20192907 | PRJNA746237 | INCLUDED | INCLUDED |
| *Felis silvestris* | FA699 | wild | Germany | FA699_Fss | ERR4671936, ERR4777100 | SAMEA7324340 | PRJEB40421 | INCLUDED | INCLUDED |
| *Felis silvestris* | FA807 | wild | Germany | FA807_Fss | ERR4671937, ERR4777101 | SAMEA7324341 | PRJEB40421 | INCLUDED | INCLUDED |
| *Felis silvestris* | FB139 | wild | Germany | FB139_Fss | ERR4671940, ERR4777104 | SAMEA7324346 | PRJEB40421 | INCLUDED | INCLUDED |
| *Felis silvestris* | FB434 | wild | Germany | FB434_Fss | ERR4777106, ERR4671942 | SAMEA7324348 | PRJEB40421 | INCLUDED | INCLUDED |
| *Felis silvestris* | FC779 | wild | Germany | FC779_Fss | ERR4671948, ERR4777112 | SAMEA7324355 | PRJEB40421 | INCLUDED | INCLUDED |
| *Felis silvestris* | FD582 | wild | Germany | FD582_Fss | ERR4671955, ERR4777119 | SAMEA7324363 | PRJEB40421 | INCLUDED | INCLUDED |
| *Felis silvestris* | FE384 | wild | Germany | FE384_Fss | ERR4777121, ERR4671957 | SAMEA7324365 | PRJEB40421 | INCLUDED | INCLUDED |
| *Felis silvestris* | FF567 | wild | Germany | FF567_Fss | ERR4777130, ERR4671966 | SAMEA7324378 | PRJEB40421 | INCLUDED | INCLUDED |
| *Felis silvestris* | FF975 | wild | Germany | FF975_Fss | ERR4671967, ERR4777131 | SAMEA7324379 | PRJEB40421 | INCLUDED | INCLUDED |
| *Felis silvestris* | FG711 | wild | Germany | FG711_Fss | ERR4777135, ERR4671971 | SAMEA7324383 | PRJEB40421 | INCLUDED | INCLUDED |
| *Felis margarita* | FCH | wild | *NA* | Felis_chaus | SRR2062187 | SAMN03771691 | PRJNA286908 | INCLUDED | NOT INCLUDED |
| *Felis chaus* | FMA08 | wild | UAE | Felis_margarita | ERR10494328-ERR104943[illegible] | SAMEA112127401 | PRJEB57412 | NOT INCLUDED | INCLUDED |

**Supplementary table 7.** Ratio between reads aligned to the reference genomes of the mtDNA (NC_001700.1) and the nuDNA (F.catus_Fca126_mat1.0, GCF_018350175.1) of *F. catus* . The number of reads were normalized by the respective reference genome lengths. Length mtDNA: 17.009 bp. Length nuDNA: 2.425.000.000 bp

| **Sample ID** | **ratio_mt/fg** |
|---|---|
| BG5 | 7.96 |
| BG16 | 25.57 |
| BG07 | 33.39 |
| IRT01 | 40.68 |
| PETR06 | 74.68 |
| PNE03 | 82.13 |
| MRY09 | 85.71 |
| FLUM02 | 88.57 |
| BHHL03 | 89.8 |
| TERT03 | 95.38 |
| BA03 | 95.47 |
| DOS03 | 98.66 |
| PAL10 | 108.53 |
| CET01 | 113.93 |
| BA02 | 116.08 |
| KASP01 | 129.29 |
| PRN02 | 129.29 |
| HAIS01 | 132.01 |
| DOS01 | 139.13 |
| GLCA01 | 140.79 |
| LAB01 | 146.76 |
| CEC03 | 171.23 |
| MUS01 | 175.1 |
| FLUM02 | 175.16 |
| MRY07 | 175.24 |
| BP03 | 180.02 |
| AC01 | 190.46 |
| COC01 | 191.62 |
| TAB02B | 197.27 |
| LHM02 | 205.76 |
| KASP02 | 207.53 |
| MRY08 | 212.45 |
| MRY10 | 214.9 |
| PAL09 | 225.46 |
| VIA04 | 226.26 |
| COC01 | 228.21 |
| CAST02 | 234.11 |
| KOP04 | 235.44 |
| MUS01 | 236.8 |
| BP02 | 246.45 |
| DUR01 | 249.22 |
| PETR08B | 257.64 |
| ROC01 | 259.53 |
| KASP02 | 262.1 |
| BHHL01 | 263.62 |
| ROC02 | 268.68 |
| PNE02 | 274.44 |
| PAL09 | 274.9 |
| SAR01 | 277.95 |
| LHM03 | 284.41 |
| BREM05 | 284.89 |
| VIA06a | 284.98 |
| DOS01 | 289.24 |
| SBR01 | 302.79 |
| TRMR01B | 310.13 |
| SCAT01 | 320.13 |
| PNE03 | 322.05 |
| GSA01 | 323.94 |
| AC01 | 326.22 |
| MTR01B | 336.2 |
| AF03 | 336.43 |
| PAL10 | 339.57 |
| CEC03 | 347.55 |
| SDUC02B | 348.47 |
| GRA01 | 378.98 |
| BHHL01 | 380.74 |
| KOP04 | 380.88 |
| BA03 | 394.09 |
| BRPA27 | 436.7 |
| HAIS04 | 443.06 |
| ROC04 | 443.42 |
| BRPA27 | 460.93 |
| HAIS10 | 470.6 |
| PNE02 | 503.8 |
| SA14 | 537.86 |
| PAD01 | 573.66 |
| GSA01 | 609.11 |
| MENT01 | 612.79 |
| BP01 | 615.72 |
| BREM07 | 653.95 |
| ECLY01 | 657.47 |
| ASK01 | 745.38 |
| DUR02 | 749.46 |
| BRPA24 | 777.75 |
| DH06 | 902.6 |
| TAB02B | 904.87 |
| VIN04 | 1004.81 |
| HAIS10 | 1041.13 |
| TON03 | 1117.64 |
| DUR03 | 1307.78 |
| BRPA24 | 2302.31 |

**Supplementary table 8.** Complete mtDNA turnover model at varying F. l. lybica introgression proportions in Neolithic Anatolian wildcats at different effective population sizes (Ne).

| | European wildcat Ne (mtDNA) | | | | | | |
|---|---|---|---|---|---|---|---|
| **Introgressing mtDNA proportion (p)** | 1,000 | 2,000 | 2,500 | 5,000 | 7,500 | 10,000 | 20,000 |
| 1.00E-05 | 5,999.90 | 11,999.90 | 14,999.90 | 29,999.80 | 44,999.80 | 59,999.70 | 119,999 |
| 1.00E-04 | 5,999.70 | 11,999.40 | 14,999.20 | 29,998.50 | 44,997.70 | 59,997.00 | 119,994 |
| 1.00E-03 | 5,997.00 | 11,994.00 | 14,992.50 | 29,985.00 | 44,977.50 | 59,970.00 | 119,940 |
| 5.00E-03 | 5,984.90 | 11,969.90 | 14,962.40 | 29,924.90 | 44,887.30 | 59,849.70 | 119,699 |
| 0.01 | 5,969.90 | 11,939.80 | 14,924.70 | 29,849.50 | 44,774.20 | 59,699.00 | 119,398 |
| 0.02 | 5,939.60 | 11,879.20 | 14,849.00 | 29,698.00 | 44,547.00 | 59,396.00 | 118,792 |
| 0.05 | 5,847.40 | 11,694.90 | 14,618.60 | 29,237.20 | 43,855.80 | 58,474.40 | 116,949 |
| 0.1 | 5,689.40 | 11,378.90 | 14,223.70 | 28,447.30 | 42,671.00 | 56,894.70 | 113,789 |
| 0.15 | 5,525.60 | 11,051.30 | 13,814.10 | 27,628.20 | 41,442.30 | 55,256.40 | 110,513 |
| 0.2 | 5,355.40 | 10,710.90 | 13,388.60 | 26,777.20 | 40,165.80 | 53,554.50 | 107,109 |
| 0.3 | 4,993.40 | 9,986.90 | 12,483.60 | 24,967.20 | 37,450.90 | 49,934.50 | 99,869 |
